# The Interplay of Electrostatics and Chemical Positioning in the Evolution of Antibiotic Resistance in TEM β-Lactamases

**DOI:** 10.1101/2021.05.27.446023

**Authors:** Samuel H. Schneider, Jacek Kozuch, Steven G. Boxer

**Author notes:** These authors contributed equally to this work (Samuel H. Schneider, Jacek Kozuch). Present address: Department of Pharmaceutical Chemistry, University of California San Francisco, San Francisco, CA, 94158, USA. Present address: Experimental Molecular Biophysics, Department of Physics, Freie Universitat Berlin, Arnimallee 14, 14195 Berlin, Germany (Jacek Kozuch).

## Abstract

The interplay of enzyme active site electrostatics and chemical positioning are important for understanding the origin(s) of enzyme catalysis and the design of novel catalysts. We reconstruct the evolutionary trajectory of TEM-1 β-lactamase to TEM-52 towards extended-spectrum activity to better understand the emergence of antibiotic resistance and to provide insights into the structure-function paradigm and non-covalent interactions involved in catalysis. Utilizing a detailed kinetic analysis and the vibrational Stark effect, we quantify the changes in rates and electric fields in the Michaelis and acyl-enzyme complexes for penicillin G and cefotaxime to ascertain the evolutionary role of electric fields to modulate function. These data are combined with MD simulations to interpret and quantify the substrate-dependent structural changes during evolution. We observe that this evolutionary trajectory utilizes a large preorganized electric field and substrate-dependent chemical positioning to facilitate catalysis. This governs the evolvability, substrate promiscuity, and protein fitness landscape in TEM β-lactamase antibiotic resistance.

## Introduction

The origins of the enormous catalytic capacity of enzymes remains a long-standing question. Unravelling these origins is essential to our ability to design and engineer proteins with diverse functions, as well as rationalize the evolutionarily-acquired mutations that lead to altered functions such as novel substrate scopes and chemistry, solvent adaptation, thermal-stability, and improved rates, as demonstrated with the many successes of directed evolution.^1–2^ A physical picture of enzyme evolution requires new observables that can link evolutionary changes in catalysis to local interactions arising from mutations.

Recent experimental work, based on the electrostatic catalysis model and the vibrational Stark effect (VSE), has quantified the electrostatic contribution to catalysis in the model enzyme, ketosteroid isomerase (KSI),^3–4^ and more generally for enzymes such as serine proteases and dehalogenases.^5–6^ The VSE method utilizes high-frequency and local vibrational reporters, such as carbonyls (C=O), to quantify the electric field in enzyme active sites. Carbonyls are the functional moieties of many enzymatic reactions, including KSI and the β-lactamases discussed in detail in the following, making them ideal for investigating electrostatic catalysis.

Measurement of the active site electric field is achieved through calibration of the carbonyl-containing molecule by VSE spectroscopy, using a well-defined external electric field, vibrational solvatochromism and MD simulations.^5, 7^ This approach provides an electric field-frequency calibration curve from which the environment’s electric field projected onto the probe bond can be directly quantified using vibrational (IR or Raman) spectroscopy.^6^ By combining the free-energy barrier of activation, *ΔG*^*‡*^, from steady-state kinetics and transition-state (TS) theory with measurements of the active site electric field 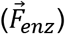 for a series of wild-type (WT) and mutant enzymes, the contribution of electrostatic catalysis to accelerate a reaction (*ΔΔG*^*‡*^) can be modeled according to,^6, 8^

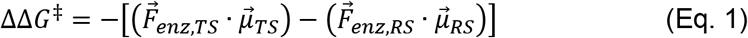

where 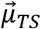 and 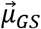 are the TS and reactant-state (RS) dipole moment for bonds that change charge distribution over the course of the reaction coordinate.

All previous studies using the VSE approach have been performed on wild-type and site-directed mutant enzymes,^6^ where the latter were all deleterious to the overall enzymatic function and do not necessarily reflect those acquired over the course of evolution. However, the multi-dimensional evolutionary fitness landscape can modulate favorable enthalpic and entropic effects by improving the electrostatic environment and chemical positioning within the enzyme active site. Chemical positioning refers to both entropic and enthalpic effects on catalysis of placing reactive groups in close proximity and correct orientations relative to the bound substrate. While both electrostatic catalysis and chemical positioning are important, we start with the hypothesis that, similar to KSI,^6^ if electrostatic TS-stabilization is the primary driving force for navigating the evolutionary fitness landscape, then an enzyme’s evolution towards improved function (i.e. lower *ΔG*^*‡*^) would correspond to a proportional increase in the active site electric field (note that negative electric fields correspond to stabilizing interactions).

The enzymes we have chosen to test this hypothesis are β-lactamases, which play a clinical role in the emergence of antibiotic resistance.^9–10^ TEM-1, a class A and the first isolated and genetically-characterized β-lactamase, has served as a model enzyme for understanding protein evolution towards extended-spectrum β-lactamase (ESBL) activity, one of the mechanisms for achieving antibiotic resistance.^11–22^ There are now over 240 TEM β-lactamase clinical variants (and over 7100 identified β-lactamases in total^23^) that endow extended-spectrum activity against multiple β-lactams (including penicillins, cephalosporins, and monobactams) as well as inhibitor-resistance (e.g. clavulanic acid).^24^ TEM-1 has narrow spectrum penicillinase activity, i.e. against penicillin G (PenG) and ampicillin, and its evolution towards new function against other classes of β-lactam antibiotics, such as cephalosporins, has been observed both in clinical isolates and directed evolution.^25–26^

One of the most well-studied ESBL evolutionary trajectories is from TEM-1 over the course of three mutations, E104K, G238S, and M182T, to TEM-52 (E104K/G238S/M182T Ambler notation;^27^ Figure 1a), resulting in both decreased penicillinase activity and increased hydrolysis of the third-generation cephalosporin, cefotaxime (CTX). We use the functionally critical β-lactam C=O (Figure 1b)^25, 28–29^ as an IR probe of the active site electric field. The general mechanism and reaction coordinate for class A β-lactamases are shown in Figure 1c. Upon binding, the β-lactam antibiotic forms the non-covalent enzyme-substrate complex (ES) which is first acylated via ring-opening of the β-lactam by the active site serine (S70), forming an acyl-enzyme intermediate (AE). The AE complex is subsequently deacylated via a nucleophilic water molecule activated by a nearby glutamate (E166), followed by product release.^10^ Note that in contrast to KSI which uses amino-acid side-chains to H-bond with the substrate/inhibitor, TEM β-lactamases interact with the bound β-lactam C=O through backbone amides of S70 and A237. These backbone amides serve as hydrogen-bond donors and are maintained throughout the course of natural evolution, although slight changes in their position may accompany mutations elsewhere in the protein.

**Figure 1.**
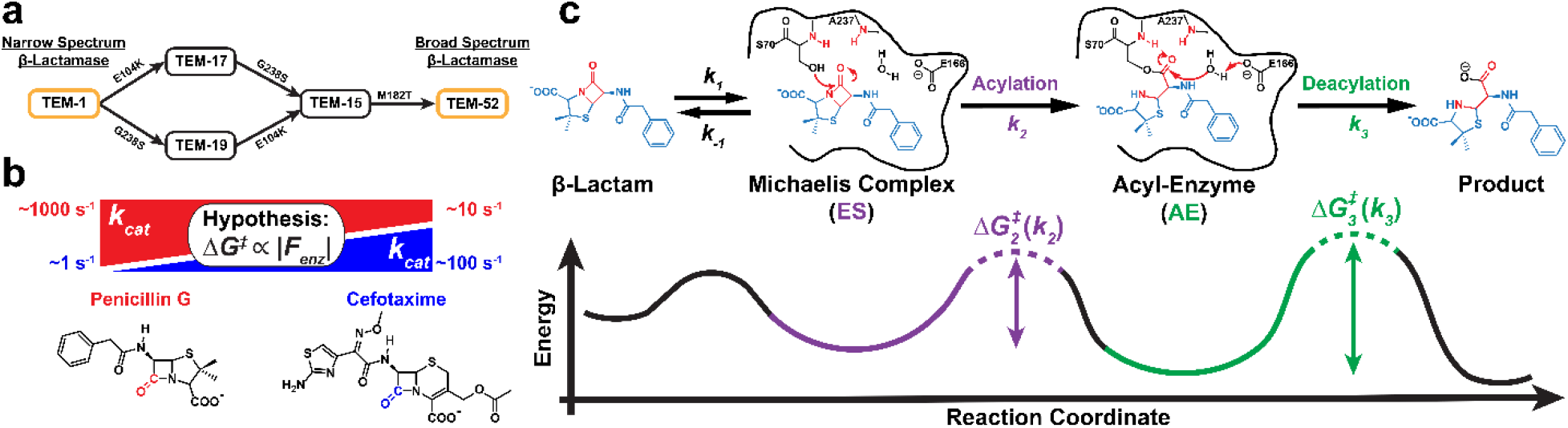
Evolutionary trajectory and reaction mechanism of TEM-1 to TEM-52 β-lactamase. **(a)** The most-probable evolutionary trajectory from TEM-1 to TEM-52 with the acquisition of three mutations (E104K, G238S, and M182T) which alters the substrate scope of TEM β-lactamase from narrow spectrum to broad spectrum activity against β-lactam antibiotics. **(b)** Order of magnitude change in rate-constant for the hydrolysis (k_cat_) of penicillin G (PenG; red) and cefotaxime (CTX; blue) over the course of evolution from TEM-1 to TEM-52^30^ with the hypothesized change in free energy barrier of activation (ΔG^‡^ from TS-theory) proportional to the active site electric field, |F_enz_|. Reactive carbonyls of the β-lactams are highlighted for clarity. **(c)** Mechanism of class A β-lactamases and reaction coordinate diagram for PenG hydrolysis. The Michaelis complex (ES; non-covalent) and acyl-enzyme (AE; covalent) are illustrated in reference to functionally important active site residues formed via the amide backbone nitrogens of S70 and A237 as well as key catalytic residues, S70 and E166, necessary for the acylation and deacylation reactions, respectively. The functional β-lactam ring and oxyanion hole are shown in red and all other substrate moieties in blue, where chemical functionalization is altered across antibiotic classes (e.g. penicillins, cephalosporins, monobactams, etc.). The two chemical steps, acylation (k_2_; purple) and deacylation (k_3_; green) can be related to their respective activation free energy barriers through transition-state theory which proceed through tetrahedral intermediates at each step (dotted lines, see Scheme S1 for details). Barrier heights not drawn to scale.

In the following, we use two substrates, PenG and CTX, to monitor the evolutionary changes to catalytic rates, the electric fields sensed by the reactive β-lactam, and chemical positioning throughout the active sites. In order to understand the mechanistic implications of these evolutionary changes we measured the Michaelis-Menten kinetic parameters and the corresponding acylation (*k*_*2*_) and deacylation (*k*_*3*_) rate constants for all mutants utilizing both PenG and CTX as substrates. Using the VSE to monitor local catalysis-relevant electric fields and molecular dynamics (MD) simulations to evaluate changes in chemical positioning, we expand upon the physical origins of antibiotic resistance and catalysis in TEM β-lactamases.

## Results and Discussion

### Kinetic Decomposition of β-Lactamase Activity

Given the mechanism depicted in Figure 1c for class A β-lactamases, the Michaelis-Menten parameters, *k*_*cat*_ and *K*_*M*_, correspond to:

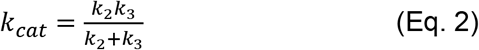

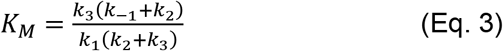

The acylation (*k*_*2*_) and deacylation (*k*_*3*_) rate constants were determined following the previously described steady-state mass spectrometry approach (Figures S1-2).^31^ At steady-state conditions, the relative concentrations of the AE and ES complexes can be directly related to the corresponding rate constants according to (Figures S3-4):

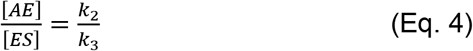

The kinetic results are presented pictorially in Figure 2 and summarized in Table S1 (see Methods and Figure S1-2).

**Figure 2.**
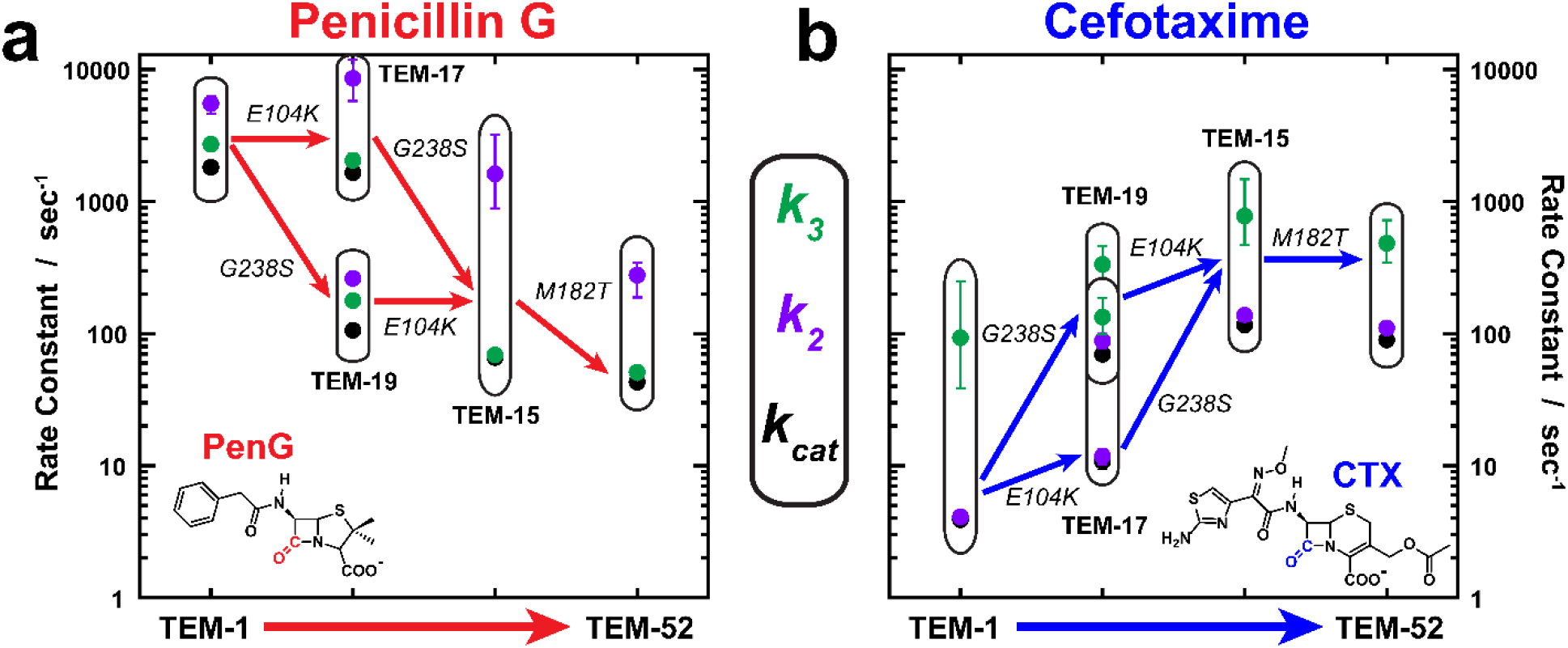
Evolution of the catalytic rates of β-lactam hydrolysis from TEM-1 to TEM-52. The absolute rate constants for **(a)** PenG (red) and **(b)** CTX (blue) hydrolysis are shown for k_cat_ (black), k_2_ (purple), and k_3_ (green), for each TEM β-lactamase mutant along the evolutionary trajectory from TEM-1 to TEM-52 as denoted by directional arrows. Error bars represent the equivalent of 1 standard deviation (68.25% confidence interval (CI)), see the methods section,Table S1, and Figures S3-4 for details.

As seen in Figure 2, the rate-constants all change in accordance with the observed ESBL phenotype as TEM-1 evolves towards TEM-52. In the case of PenG hydrolysis (Figure 2a), the mutations generally decrease the rate-constants in accordance with the loss of function as a narrow-spectrum penicillinase; however, the *K*_*M*_ decreases significantly going from TEM-1 to TEM-52, primarily as a result of the G238S mutation (Table S1). As has been previously observed with TEM-1 and PenG,^31–32^ the rate-limiting step for all variants is deacylation (*k*_*3*_). The extent to which acylation and deacylation change (i.e. *k*_*2*_/*k*_*3*_) is differentially affected by the mutations. All of the mutations, with the exception of TEM-19 (G238S), lead to *k*_*2*_/*k*_*3*_ perturbed away from that of TEM-1 (*k*_*2*_ ≈ *k*_*3*_),^32^ with the double-mutant TEM-15 being the most perturbed (*k*_*2*_/*k*_*3*_ > 10). The ratio is then partially recovered with the global suppressor mutation, M182T, which is distant from the active site but thermodynamically stabilizing,^16^ leading to TEM-52 (*k*_*2*_/*k*_*3*_ ≈ 5). This kinetic evaluation deconvolves some of the epistatic relationship of these mutations on function, supplementing previous studies.^12–13^

In the case of the evolutionarily improved CTX hydrolysis (Figure 2b), we observe an improvement in both binding and turnover with the two distal active site mutations, E104K and G238S, and minimal effect of M182T, relative to TEM-1. Significantly, and in contrast to that of PenG hydrolysis, the rate-limiting step for each variant is the acylation (*k*_*2*_) step, such that *k*_*2*_ ≈ *k*_*cat*_, as previously suspected by several authors.^10, 31, 33^ Furthermore, the relative ratio of *k*_*2*_/*k*_*3*_ generally improves (i.e. towards unity) as TEM-1 evolves towards the ESBL TEM-52, primarily as a result of the G238S mutation. In terms of TEM-52’s phenotype as an ESBL, i.e. increased promiscuity, the final corresponding rate-constants for both PenG and CTX converge towards the same order of magnitude, consistent with the broad-spectrum activity profile.

### VSE Measurements of β-Lactamase Complexes

We utilized isotope-edited ^12^C – ^13^C difference spectroscopy to determine the active site electric fields using the VSE, made possible by site-specific isotope incorporation of ^13^C into common β-lactam antibiotics.^34^ This method enables detection of a single C=O stretching frequency even amid the extensive protein backbone signal (Figure S5 & 7).^35^ In order to trap the ES and AE species for the intrinsically slow IR measurements, we introduce the S70G and E166N mutations, respectively, to monitor the mechanistic intermediates. These kinetically incompetent background S70G or E166N mutations are denoted “TEM-# S70G” or “TEM-# E166N” relative to the evolutionary and clinically observed WT proteins TEM-1, −17, −19, −15, and −52. We infer the projected electric fields based on the observed frequency shifts using the electric-field/frequency calibration of model compounds.^36^ We use the smallest and chemically simplest rigid functional moieties of the β-lactam substrate’s C=O, namely the bicyclic penam/cephem core and the acylated ester for the ES and AE complexes, respectively (see ref. ^36^ and Figure S6).

Isotope-edited difference IR measurements of PenG and CTX in the S70G ES and E166N AE complexes with TEM β-lactamases are shown in Figure 3 (see Figure S8 for further details of spectral processing and Figure S9 for additional S70 and E166N variant spectra). Across all of the enzyme-substrate complexes we observe multiple sets of narrow (FWHM 5-16 cm^−1^) vibrational features, with corresponding sets of ^12^C (positive) and ^13^C (negative) features (shown only on the bottom spectrum in each panel for clarity, Fig 3 c-f) that exhibit the expected isotope frequency shift of 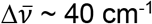 (Tables S1-4). These peaks are significantly narrower than the β-lactam C=O frequency in aqueous buffer (D2O, Figure S5), which is observed at 1762 cm^−1^ with FWHM ~ 35 cm^−1^. The multiple features are indicative of conformational heterogeneity upon binding. Previous studies with β-lactamases have indicated the possibility of multiple peaks upon substrate binding, but they could not be confidently assigned in the absence of isotopic labels.^37–38^

**Figure 3.**
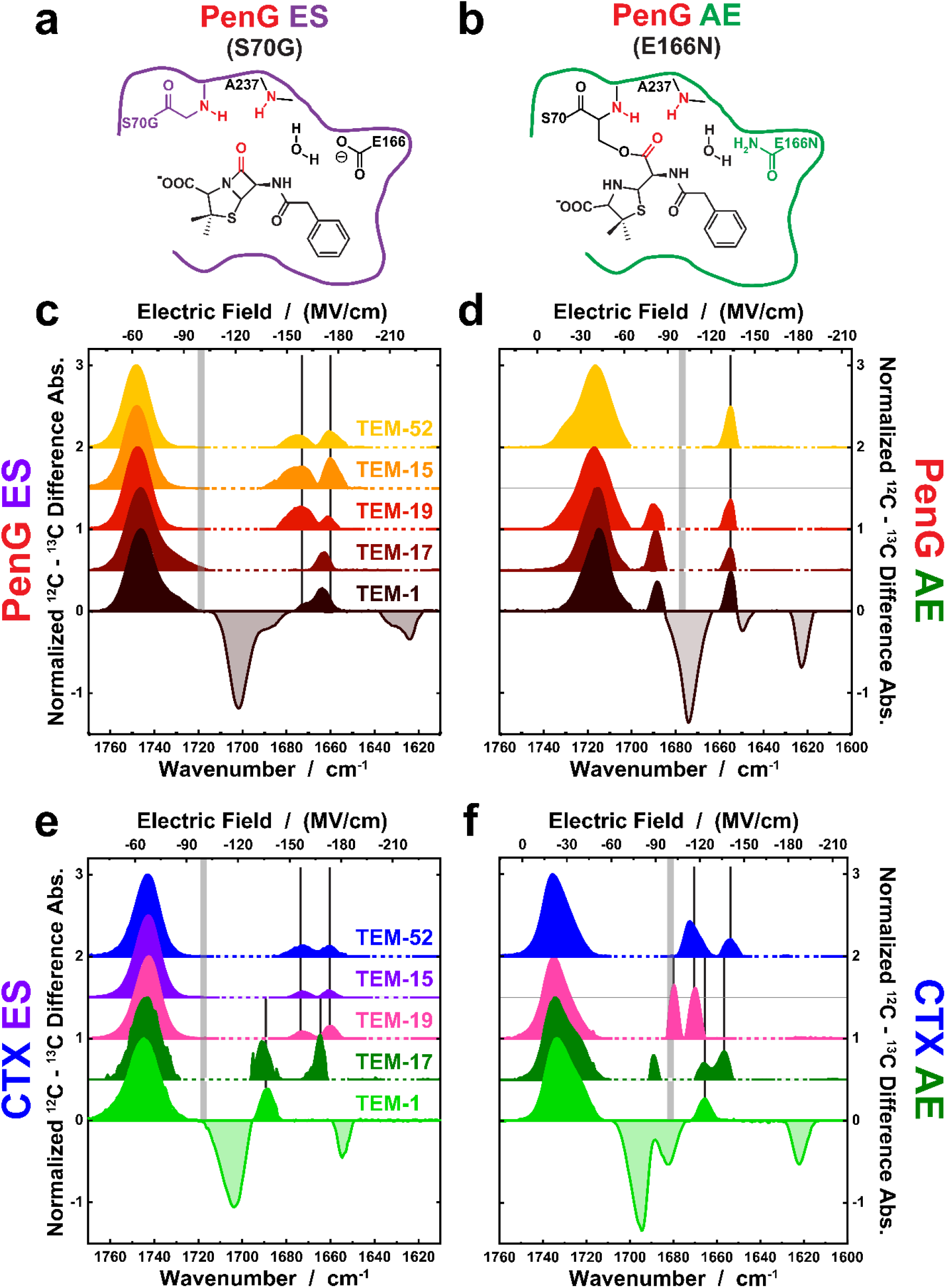
Isotope-edited IR difference spectroscopy of the TEM β-lactamases. **(a,b)** Schematic illustration of the active site environment of TEM β-lactamases in the **(a)** ES and **(b)** AE complexes with PenG and the background mutations S70G and E166N required for FTIR spectroscopy. The oxyanion hole, composed of backbone amides from S70 and A237, relative to the substrate C=O are shown in red. **(c-f)** The vibrational frequencies and corresponding electric fields for **(c, d)** PenG and **(e, f)** CTX in the **(c, e)** ES (S70G) and **(d, f)** AE (E166N) complexes across evolutionary variants from TEM-1 to TEM-52. IR spectra are shown as a 12C-normalized difference absorbance with an example ^12^C – ^13^C spectra shown for TEM-1 complexes and otherwise depicted with only the positive (^12^C) features for clarity with dashed horizontal axes indicating negative features that have been removed for simplicity (see Figure S7 for full ^12^C – ^13^C difference spectra). The top axis is the electric field corresponding to the ^12^C-isotope of the corresponding VSE calibrated probe (Figure S6). The panels are sub-divided between those C=O species corresponding to small or large electric fields (−100 MV/cm; vertical gray bars) to broadly classify the results. Vertical black lines indicate large electric field species changes between evolutionary mutations. The TEM-15 E166N mutant was not measured due to the protein’s instability and likely aggregation, observed as increasing protein-specific spectral changes and loss of 280 nm absorbance over time.

As shown in Figure 3 c-f, where the upper axis translates the observed frequencies into electric fields (Figure S6), we observe predominantly two distinct electric field environments for both PenG and CTX: a major population (by integrated area) in a small electric field conformation (ca. −30 to −70 MV/cm), and minor populations at significantly larger and more stabilizing (i.e. more negative) electric fields (ca. −140 – −175 MV/cm). The lineshapes of the lower electric field species are broad compared to the minor species experiencing much larger electric fields. The simplest interpretation of linewidth variations in proteins is that they reflect the heterogeneity of the environment around the probe.^39^ For PenG, we observe minimal changes in the peak position/electric field over the course of (de)evolution for either the small or large electric field species, with the largest electric field species consistently observed at ca. −175 and −140 MV/cm (Figure 3a,b) for the ES and AE complexes, respectively. However, there is an increase in conformational heterogeneity as evidenced by additional populations with fields of ca. −160 and −30 MV/cm with introduction of G238S (i.e. TEM-19, TEM-15, and TEM-52). For PenG the negligible effect on the electric field sensed by the reactive C=O over evolution contradicts the hypothesized changes expected based on the observed rate decrease; the implications of this are discussed below. For CTX (Figure 3c,d), we observe minimal changes in the small electric field species but significantly larger electric fields detected over the course of evolution towards TEM-52, with the largest improvement in field (ΔF(TEM-52 – TEM-1) ≈ −40 MV/cm) corresponding to the ES complex (*k*_*2*_ is rate-limiting). Interestingly, the largest electric fields projected onto the CTX C=O converge towards the maximum fields experienced by PenG’s C=O of ca. −175 and −145 MV/cm for the ES and AE complexes, respectively; the corresponding conformational heterogeneities converge in a similar way. The magnitude of the electric field in the high-field population in TEM is consistent with or even considerably larger than that of other proteins with similar oxyanion holes comprised of backbone amides (ca. −90 to −120 MV/cm).^5^

### MD Simulations of β-Lactamase Complexes – Effects of S70G and E166N Mutations

We utilized MD simulations to provide a structural basis for interpreting the IR and kinetic results as well as monitoring changes in chemical positioning between the start and end points of the evolutionary trajectory (TEM-1 and −52; 1 μs MD runs of each). The MD simulations provide two primary results: (1) they compare and contrast the extent to which the necessary background mutations, S70G and E166N, bias the conformational sampling relative to their WT variants and assess the functional relevance of the species detected by IR; and (2) they help elucidate the structural origins and effects of chemical positioning on catalysis and evolution, which cannot be fully described by electrostatics as evidenced by the VSE results.

Protein-ligand complexes were simulated starting from an initial ligand pose based on crystallographic coordinates of PenG-acylated TEM-1 E166N (PDB ID: 1FQG) and CTX-acylated Toho-1 E166A (PDB ID: 1IYO), which were modelled into the corresponding variants in both the non-covalent ES and covalent AE states (see Methods). We focus on the oxyanion hole interactions that are important for function and most directly relevant to interpreting the VSE results,^40^ namely, the O – N distances between the β-lactam C=O and the backbone amide nitrogens of S70 and A237. To compare the background mutants with WT, the O – N distances were measured for the ES complexes of WT and S70G, and the AE complexes of the WT and E166N; the comparison is made for both TEM-1 and TEM-52 (Figure 4a-d). Based on these MD trajectories, 2D-correlation plots of the O_βlactam C=O_ – N_A237_ (“O – N237”) versus the O_βlactam C=O_ – N_S70_ (“O – N70”) distances were generated for each protein-ligand complex (Figure 4e-h), and the average structures for each conformation were compared between the mutants and WT protein complexes (Figure S10-12). Comparison of the commonly observed conformations, for both the S70G and E166N mutants and the respective WT proteins, enables a structure-guided approach to interpret the IR and kinetic results as discussed below. While it is tempting to directly calculate the active site electric field as experienced by the β-lactam C=O in the MD simulations, accurate calculation of electric fields when short H-bonds are involved in catalysis requires more advanced simulation forcefields and levels of theory.^41–46^

**Figure 4.**
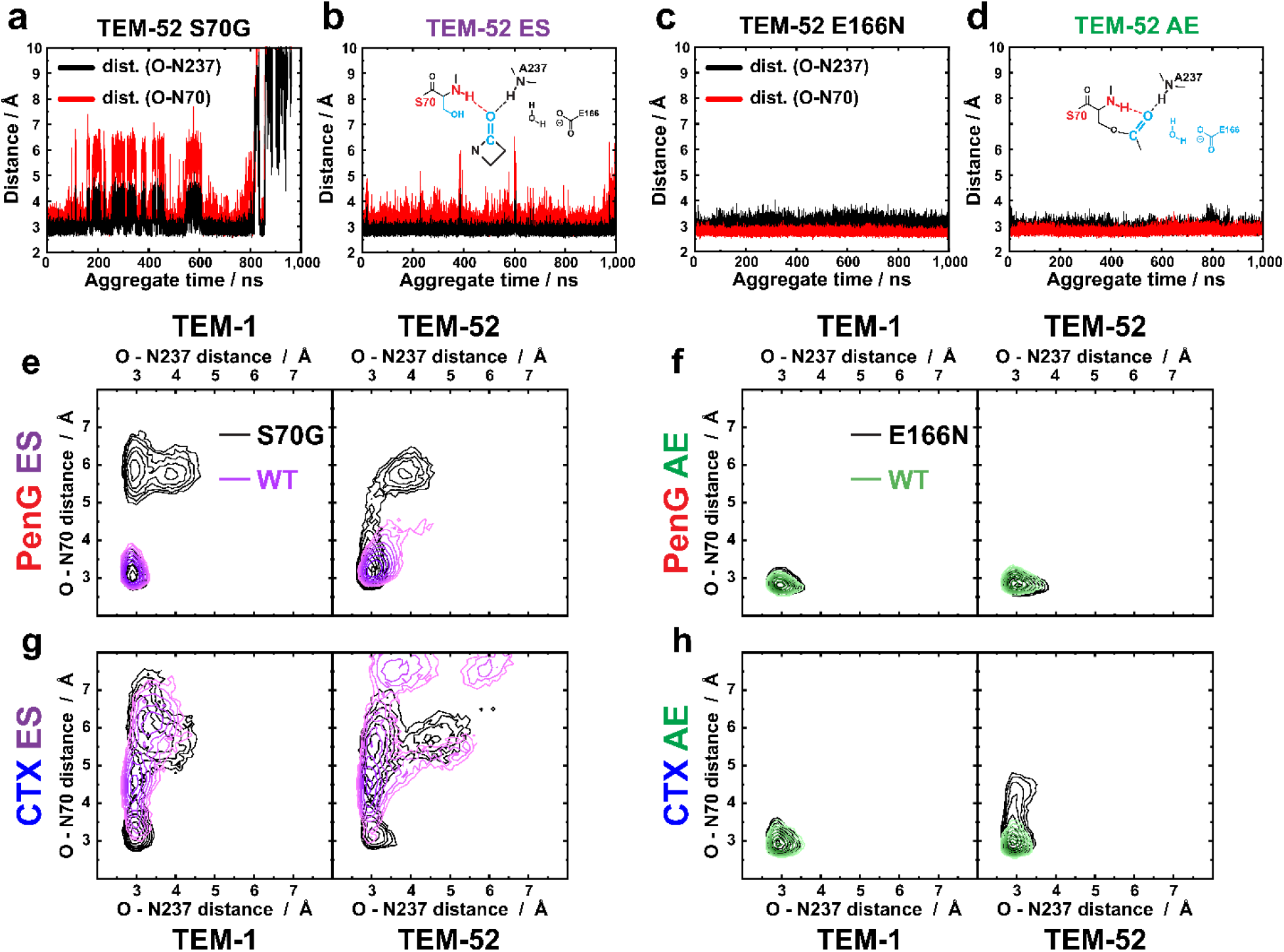
MD structure-guided approach to assess the effect of background and evolutionary mutations on the ES and AE complexes. **(a-d)** Example MD trajectories of the TEM-52 O_βlactam C=O_ – N distances between the oxyanion hole comprised of the backbone N-H of S70 (red) and A237(black), and the β-lactam C=O of PenG in the ES and AE complexes. Comparison of the background mutations **(a)** TEM-52 S70G ES and **(c)** TEM-52 E166N AE relative to the **(b, d)** TEM-52 WT ES and AE complexes, respectively, provides a means to assess perturbations to the active site environment, e.g. conformational sampling, structural fluctuations, and effects of evolutionary mutations. **(e-h)** 2D correlation plots of the O – N distances as a function of background mutants (black contours) relative to the WT for the **(e, g)** ES (purple) and **(f, h)** AE (green) complexes of TEM-1 and −52 with **(e, f)** PenG and **(g, h)** CTX. Contours represent dwell times ≥400 ps on a log-scale. Average structures of overlapping conformational species were evaluated for structural similarity (see main text).

A comparison of MD simulations of the S70G and E166N background mutants with WT complexes, which are inaccessible to our IR measurements, provides insights into states that are due to the required background mutations versus those that are catalytically permissive (Figure 4). In these simulations, the background mutations introduce exaggerated conformational heterogeneity relative to the WT, primarily evident in the ES complexes, where the functional β-lactam C=O is displaced from the oxyanion hole due to increased water accessibility (Figure S12). This results in the C=O forming H-bonds to structured water located in the active site, which would be expected to result in smaller electric fields than those observed in the oxyanion hole, consistent with the VSE measurements and vibrational solvatochromism in water (Figure 3, Figure S6,12-13).^36^ When we biased the WT ES complexes to start from structures resembling these long O – N states in the S70G mutant, we find that these conformations are not maintained, and generally converge towards the shortest O – N distance species (Figure S14). This suggests that the removal of the S70 sidechain can induce additional substrate binding modes within the active site oxyanion hole, as previously suggested in the TEM-1 S70G crystal structure.^47^ Importantly, the species with the shortest oxyanion hole distances (< 4 Å), likely the catalytically permissive states, are commonly observed in both the background mutants and WT TEM-1 and −52 simulations for the ES and AE complexes with PenG and CTX. The average structures are similar to the crystallographic binding geometry as observed for the TEM-1 S70G and E166N complexes with PenG (Figure S11,12).^47–48^ We consider only those species shared in common as the biologically relevant and catalytically permissive species.

### MD Simulations of β-Lactamase Complexes – Evolutionary Changes and Structural Plasticity

Based on the MD results, we quantify the significant (p < 0.05 in two-tailed t-test) and substrate-dependent structural changes from TEM-1 to TEM-52 in terms of the changes in inter-atom distances and root-mean square fluctuations (RMSF) of key atoms and residues in the active site (Figure 5, Scheme S2, Table S7-12). We emphasize in Figure 5 only those residues directly involved in the extended H-bond network to nucleophilic groups, i.e. the first coordination sphere, with other potentially important residues reported in Tables S7-12 and Figures S15 & 16. These structural and dynamic changes are generally involved in modulating interactions with the catalytic residues and/or water, binding of the β-lactams’ amide side-chains or carboxylates, or the extensive H-bond network in the active site. Alignment of the reactive substrate C=O in TEM-1 and TEM-52 for the corresponding ES and AE complexes provides a functional and structural glimpse into how the evolutionarily-acquired mutations modulate catalytic turnover of PenG and CTX (Figure 5a,b,e,f), complementary to the VSE measurements. The changes in structural heterogeneity of key active site atoms and residues, in terms of RMSFs, are graphically presented in Figure 5,c,d,g,h to identify key differences between TEM-1 and TEM-52 in a substrate-dependent and ES- and AE-dependent manner.

**Figure 5.**
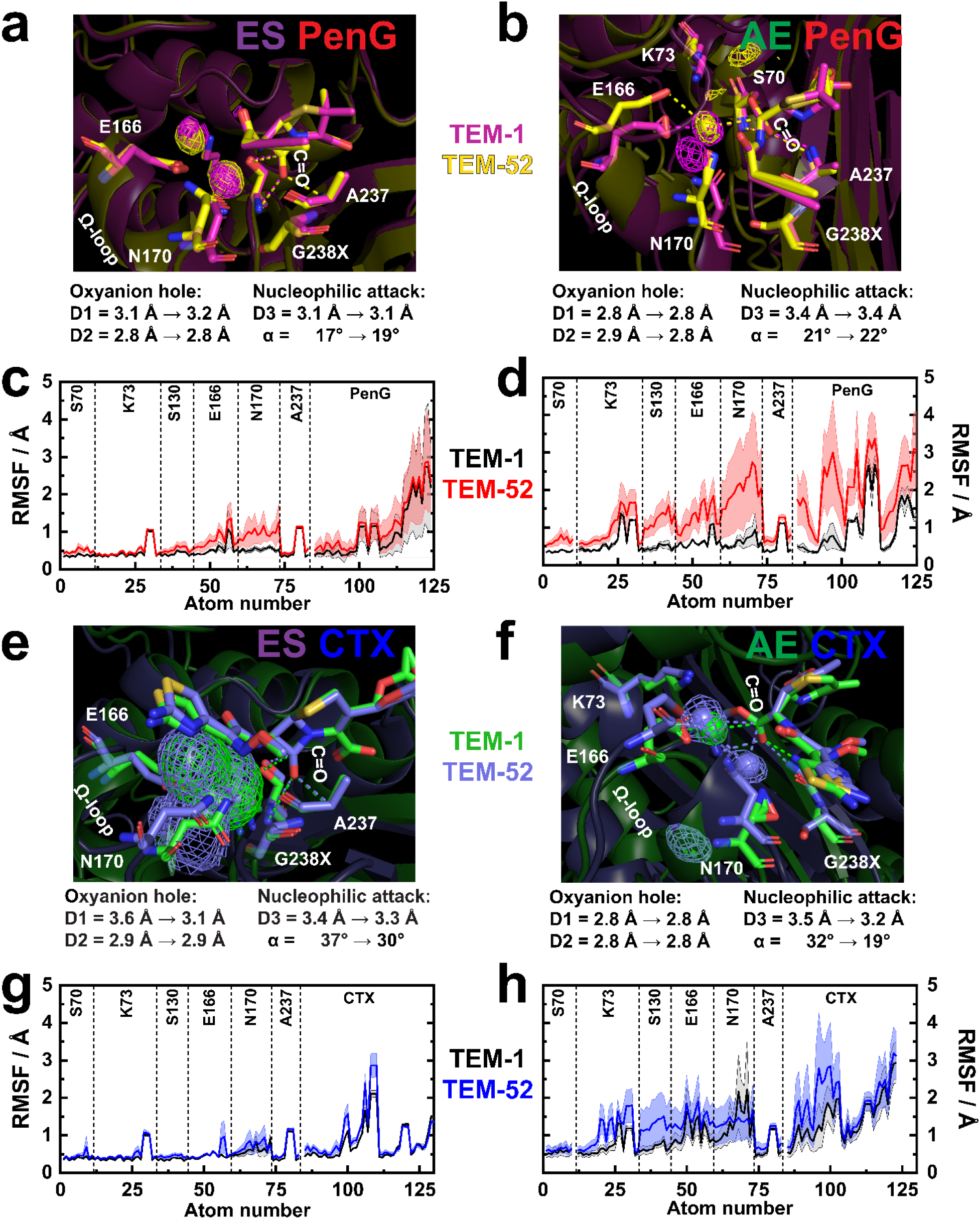
Catalytic interactions and active site structural heterogeneity in TEM β-lactamases as quantified via MD simulations. **(a,b,e,f)** Average active site structural overlay of **(a,b)** PenG and **(e,f)** CTX in complex with TEM-1 (pink or green) and TEM-52 (yellow or purple) in the ES and AE complexes, respectively. D1 and D2 refer to the N_backbone_ – O_βlactam C=O_ distances of the oxyanion hole between the β-lactam C=O and the S70 and A237 backbone amides, respectively. D3 and α refer to the distance and angle of the nucleophilic S70 side chain O atom or catalytic water O atom in the ES- or AE-complex, respectively, the latter relative to the substrate C=O (the plane of the β-lactam and ester C=O in the ES and AE complexes, respectively, Scheme S3). Structures are aligned according to the substrate’s C=O to indicate evolutionary changes from the perspective of the reactive functional group. Averaged structures as well as extracted observables/values refer to MD aggregate times of 1 μs; except for CTX ES complexes where a total of 200 ns was used. Comparison of catalytic interactions with the oxyanion hole and nucleophilic S70 or reactive water are indicated by dotted lines and values. Solid spheres in the AE complexes represent the average water positions, and isomeshed density represents the active site regions where waters are observed in the vicinity of E166 and the substrate. **(c,d,g,h)** Absolute RMSF of active site residues and substrate atoms with respect to TEM-1 (black) and TEM-52 (red or blue) for **(c,d)** PenG and **(g,h)** CTX in the ES and AE complexes. Solid lines represent the average RMSF of a given atom across the MD trajectories with the 95% CI shown with dotted lines and shaded regions. See SI Scheme S2, Table S7-12, Figure S15,16 for further details on active site RMSF and distance changes between TEM-1 and TEM-52 with PenG and CTX.

The introduction of the evolutionarily-acquired mutations E104K and G238S leads to substrate-dependent structural changes across the active site. These changes primarily affect H-bonding interactions in the Ω-loop^10, 49^ (E166 and N170) and surrounding H-bond network, which modulate the positioning of S70, E166, and the catalytic water with respect to the reactive C=O (Figure 5). For example, the average structures from MD simulations illustrate how the G238S mutation contributes to disrupting the backbone interaction of N170 – G238 in TEM-1, which results in the Ω-loop becoming more flexible and expanding the active site to accommodate larger substrates such as CTX (Figure 5).^10^ We generally observe an increase in structural heterogeneity for interactions with PenG and CTX in TEM-52 relative to TEM-1 as evident in the RMSF, which is also associated with additional active site water molecules (Figure 5b,e,f). For CTX, interactions in ES and AE are differently affected. In the ES complex we observe minimal changes in the active site RMSFs, but there is a significant shortening of the distance of the β-lactam C=O to the S70 backbone amide (ΔD1 = −0.5 Å) as well as improved positioning of the S70 side-chain hydroxyl group for the nucleophilic attack (α = 37° → 30°; Figure 5e, Scheme S3), which approaches those observed in the TEM-1 ES-complex with PenG (Figure 5a). The shortened distance of the β-lactam C=O to the backbone amide would be predicted to lead to an increase in the electric field of the highest-field (most red-shifted) component, consistent with the VSE results (Fig. 3). In the AE complex with CTX, there is increased solvent accessibility as evident from the presence of an additional water molecule in the vicinity of E166, N170 and CTX, which exhibits improved positioning in terms of attack distance and angle relative to those found in TEM-1 (Figure 5f), again approaching those observed in the AE-complex with PenG (Figure 5b). There are additional significant structural changes, in terms of both inter-atom distances and RMSFs that are observed more globally around the active site, consistent with the aforementioned observations and, thus, likely support the positioning of reactive groups as well as modulating substrate binding into the oxyanion hole (Scheme S2, Table S7-14, Figure S15,16). This is a result of the substrate-specific geometry of the bicyclic core (i.e. penam versus cephem), including the carboxylate geometry relative to the β-lactam C=O, and the amide side-chain, which together differentially influence, and are influenced by the evolutionary trajectory (Table S13,14, Figure S18).

### Role of Electric Fields and Structural Changes for Catalysis

Nearly all crystallographic studies of TEM β-lactamases with inhibitors and substrates exhibit only a single conformation,^47–48, 50–51^ while MD simulations indicate conformational sampling and biasing as a mechanism for β-lactamase evolution.^37–38, 52–57^ In order to rationalize the observed changes over the evolutionary trajectory from TEM-1 to TEM-52 we combine the kinetic, VSE, and MD results to assess the role of electrostatic, chemical positioning, and conformational effects in modulating the activation free energy barrier (*ΔG*^*‡*^ from *k*_*cat*_ and TS-theory) (Figure 6, Table S2).

**Figure 6.**
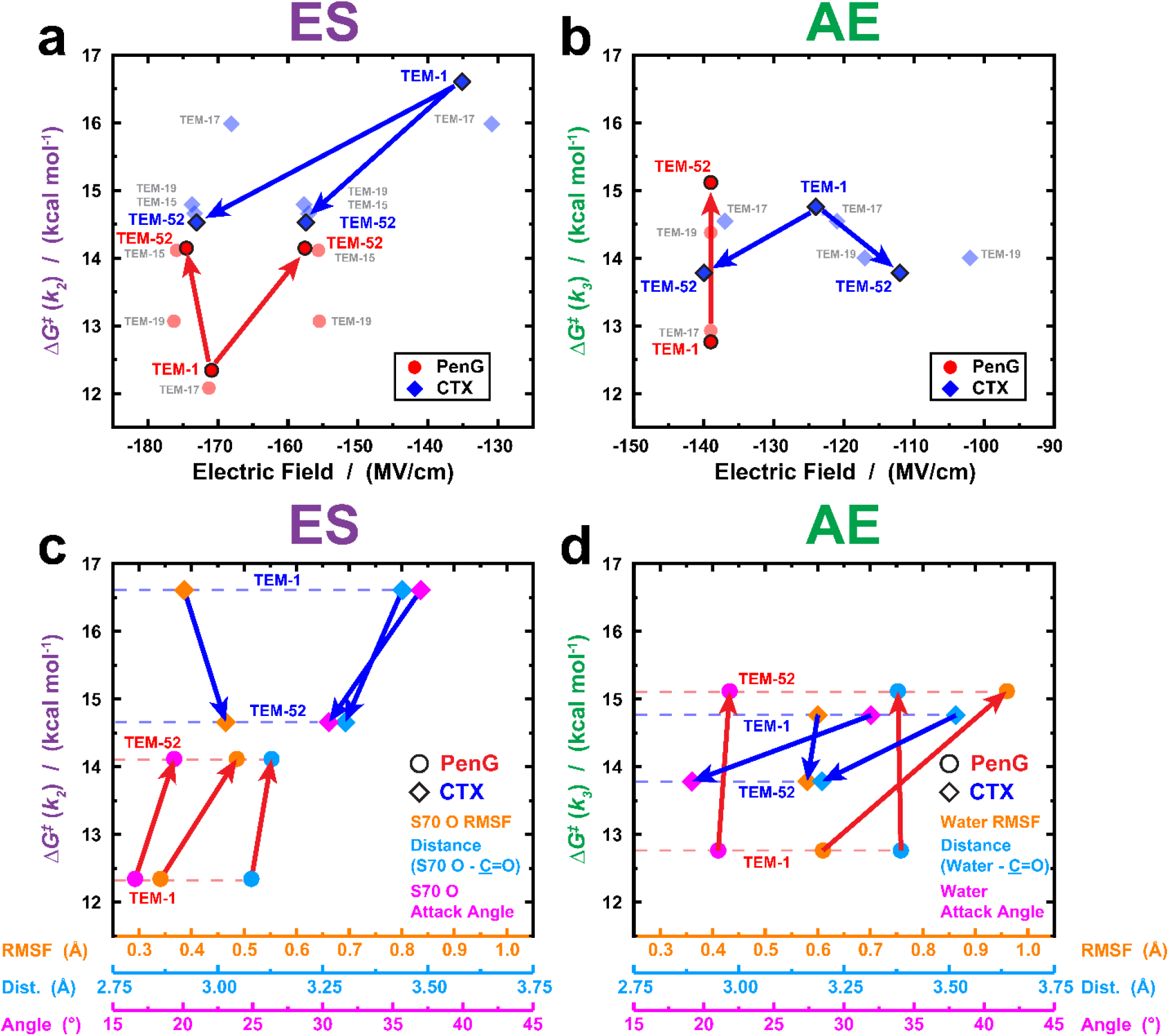
Substrate-dependent evolution of active site electric fields and chemical positioning in the catalysis of TEM β-lactamase. **(a,b)** The non-monotonic correlation between the large catalytically relevant electric field species and free energy barriers of activation from steady-state kinetics and TS-theory for **(a)** acylation (ES) and **(b)** deacylation (AE) with respect to PenG (red) and CTX (blue). **(c,d)** Correlation between the free energy barriers of activation and absolute changes in chemical positioning (RMSF, distances to β-lactam <u>C</u>=O, and attack angle) for nucleophilic attack of **(c)** S70 or **(d)** active site water from MD simulations of the ES and AE complex, respectively, for hydrolysis of PenG (circles) and CTX (diamonds). The RMSF (orange), distance to β-lactam <u>C</u>=O (light blue), and nucleophilic attack angle (pink; defined according to the plane of the C=O, Scheme S3) represent averages across the MD trajectories as discussed in Figure 4–5. Catalytic waters are assigned according to their proximity to E166, K73, N170, and the substrate C=O as described according to Figure 5. Arrows denote the direction of evolution from TEM-1 to TEM-52 for either PenG (red) or CTX (blue).

In the case of the evolved improvements in cefotaximase activity of TEM-52, relative to TEM-1, we observe changes in *both* the catalytically relevant active site electric field and positioning of the nucleophilic atoms, i.e. the side-chain of S70 and active site water involved in acylation (ES state) and deacylation (AE state), respectively, as summarized in Figure 6. Contrary to our expectations based on the kinetics, even in TEM-1 we observe in the IR spectra a catalytically relevant population with a high electric field projected onto the β-lactam C=O of CTX; MD simulations indicate that the catalytic inefficiency can be ascribed to mispositioned catalytic residues. Over the course of evolution, in the rate-limiting ES state, the three mutations result in non-monotonic increases in the magnitude of the largest electric fields experienced by the C=O (Figure 6a, blue arrows). This is consistent with E104K and G238S leading to an expansion of the active site to accommodate CTX,^10^ allowing an improved alignment of the β-lactam C=O with the backbone amide N-H’s of S70 and A237 as observed using MD and expected to generate large electric fields, consistent with what is observed by the VSE (Figure 6a, blue arrows). In addition, we observe small increases in the RMSFs of the S70 side-chain that are compensated by shortened distances and a better attack angle, i.e. improved chemical positioning, approaching those observed in the PenG complexes (Figure 6c, blue arrows). In the deacylation step, we detect improved *and* worsened electric fields over the course of evolution (Figure 6b, blue arrows), which is in line with the overall increased RMSFs of the active site (Figure 5d,h). At the same time, better positioning of the nucleophilic water, in terms of distance and attack angle with respect to the acyl ester C=O (Figure 6d, blue arrows), is achieved through increased solvent accommodation and entry of a second water molecule (Figure 5e,f).

Based on previous investigations of enzymes with the VSE that employ similar reaction mechanisms (e.g. serine proteases),^6^ a reduction of the activation free energy barrier by 1 kcal/mol corresponds to more stabilizing electric fields by 10 – 20 MV/cm if modulated primarily by electrostatic TS-stabilization. The observed improvements of electric field are consistent with this relative change in activation free energy barrier for acylation (rate-limiting) and deacylation for CTX hydrolysis, so that increased RMSFs are compensated by improved distances and attack angle of nucleophilic atoms via global structural changes throughout the enzyme active site (Figure S15-17). For deacylation, the divergence of electric fields over the course of evolution may suggest that improved chemical positioning is the major contribution to improved rates.

In the case of the PenG hydrolysis, where deacylation is rate-limiting (Figure 2a), there is no obvious selective pressure to preserve the catalytic efficiency of TEM-1 during the evolutionary trajectory to TEM-52. We observe that the E104K mutation greatly alters the *k*_*2*_/*k*_*3*_ ratio, resulting in *k*_*3*_ ≈ *k*_*cat*_, consistent with either a reduced electric field magnitude at the β-lactam C=O or mispositioning of E166 and the nucleophilic water molecule for deacylation. The β-lactam C=O vibration is only sensitive to the electric field projected onto its bond axis, and therefore changes occurring orthogonal to or far from the reactive C=O probe, such as changes in the positioning of E166, S70, or the catalytic water, can not be spectroscopically observed. Therefore, the apparent absence of a change in electric field for the catalytically relevant high-field population in the IR spectra (Figure 6a,b) suggests that the changes in chemical positioning of active site residues are a more likely basis for the observed rate changes. This is consistent with the changes in the RMSFs of the nucleophilic atoms (Figure 6c,d, orange, red arrows), such as the acylating serine, in line with decreased *k*_*2*_, and overall increased active site heterogeneity as observed in the MD that accompany the evolution from TEM-1 to TEM-52 (e.g. K73, E166, N170; Figure 5c,d, S15-17, Table S7-12). This is also consistent with the observation that G238S is primarily responsible for the reduction in both *k*_*2*_ and *k*_*3*_, which are the primary contribution to lowering *K*_*M*_ (Figure S19 and Table S15). Recently reported studies using time-resolved electrospray ionization with H/D exchange on TEM-1 with ampicillin and cephalexin are consistent with these structural changes and implications in the AE complex, with efficient deacylation (i.e. TEM-1 with PenG) being associated with tighter active site interactions as evident from decreased deuterium uptake.^58^ This increased active site heterogeneity results in PenG becoming more inhibitor-like for TEM-52 relative to TEM-1, i.e. a high on-rate (*k*_*1*_) but lower turnover (*k*_*cat*_ ≈ *k*_*3*_). The increased structural displacement and heterogeneity of the S70 and E166 side-chains and nucleophilic water are expected to break one essential aspect of the preorganized active site for acylation and deacylation, namely proper chemical positioning for catalysis.

Taken together, we see that changes in the chemical positioning of the side-chains of S70 and the catalytic water have significant kinetic consequences (Figure 6c,d) without necessarily perturbing the electric field as sensed by the β-lactam C=O (Figure 6a,b). This observation is reminiscent of the D40N mutation in KSI, which ablates the catalytic base from proton abstraction (i.e. changing the mechanism), thereby lowering the catalytic rate by 6-orders of magnitude, but has a minimal effect on the electric field projected onto the functionally important carbonyl for TS-stabilization relative to the WT enzyme.^3^

### Implications for the Role of Electric Fields, the Protein Fitness Landscape, and Antibiotic Resistance

We observe that the natural evolution from TEM-1 to TEM-52 in response to new antibiotics utilizes both electrostatic effects directed at the reactive β-lactam C=O and chemical positioning to modulate catalysis in a substrate-dependent manner. This is in contrast to prior studies of KSI, serine proteases, and 4-chlorobenzoyl-CoA dehalogenase,^6^ where the observed catalytic effect could be ascribed largely to mutations that were in close contact (e.g. H-bonding) with the reactive C=O bonds. These laboratory mutations do not necessarily recapitulate the natural evolutionary trajectory of these enzymes. The observed linear free energy relationship with the active site electric field in these enzymes indicates that there is a minimal (or constant) effect of each mutation on the overall active site structural heterogeneity, so the dominant effect is through electrostatic interaction with the reactive C=O bond. In contrast, the mutations along the evolutionary trajectory from TEM-1 to TEM-52 are not directly involved in the dominant non-covalent interactions between the β-lactam C=O and backbone amides of S70 and A237 which are difficult to perturb except as a secondary consequence of changes in backbone positioning. Rather, these mutations modulate binding interactions across the active site in a substrate-dependent manner, indicative of the structural plasticity of the TEM scaffold. These can propagate to changes in the alignment of the reactive C=O with respect to the oxyanion hole electric field, as observed using the VSE with CTX (i.e. partially attributed to the 4,6-bicyclic ring), as a result of increased active site accommodation for improved binding.

Substrate promiscuity and structural plasticity enable a starting point for evolutionary improvement,^1–2, 57, 59^ which in the case of TEM-1 takes three mutations to achieve the robust cefotaximase activity in the ESBL TEM-52. As demonstrated across all the TEM variants with both PenG and CTX, the maximum observed electric fields experienced by the C=O of the functional β-lactam in both the ES and AE complexes are surprisingly large throughout the enzyme’s evolution. This might be expected given that the backbone amides of S70 and A237 are difficult to perturb without deformation of the active site’s secondary and tertiary structure, and only a single mutation, either E104K or G238S, is necessary for the β-lactam C=O of CTX to achieve the largest electric fields observed with PenG of −140 to −175 MV/cm. If we were not starting with the evolved oxyanion hole of TEM-1 for penicillinase activity, mutations would be required to both position the C=O of CTX into the oxyanion hole (i.e. if it could not fit or bind correctly) and to increase the overall magnitude of the electric field experienced by the carbonyl group of the β-lactam ring. This hypothesis could be explored further in ancestral variants of TEM using ancestral sequence reconstruction (ASR)^60^ to compare and contrast more long-term evolutionary changes in structure, function, and fields. During the evolution of CTX hydrolysis from TEM-1 to TEM-52, the large pre-existing oxyanion hole electric field simplifies the mutational search along the fitness landscape for improved catalytic rates – via improved alignment of the reactive C=O into the active site and correct positioning of the reactive water, S70, and E166 residues – thereby enabling the ESBL phenotype to be acquired over the course of three mutations and driving the emergence of antibiotic resistance. This implies that for a new β-lactam antibiotic to be effective against a given class of β-lactamases, i.e. to minimize the risk of resistance, it must bind in a manner that is able to simultaneously avoid the large active site electric field and unproductively position the catalytic side-chains. Across class A β-lactamases, the overall active site architectures, including the oxyanion hole and active site amino acids, are well conserved and therefore it is not surprising that many families of β-lactamase have been observed (and/or have the potential) to rapidly evolve activity towards hydrolysis of penicillins, cephalosporins, and monobactams.

Our findings of large electric fields in the oxyanion hole of TEM β-lactamases may help to rationalize how structural plasticity leads to substrate promiscuity.^57, 60^ The evolutionary history leading to TEM-1 resulted in an enzyme scaffold and active site that is capable of exerting large stabilizing electric fields necessary for TSS of chemical reactions with similar substrate geometries and mechanisms. This may be the case for hydrolases and many other enzymes,^61^ which utilize pre-existing catalytic architectures, such as oxyanion holes, to enable catalysis of a large substrate scope for further improvements via directed evolution.^62^ As evident in the evolution of cefotaximase activity from TEM-1 to TEM-52, there is some basal level of binding and catalysis in the inherited protein scaffold (as required for evolution), which is supported by the large electric fields observed even in TEM-1 with CTX. Instead of electric field optimization, subsequent mutations are focused on accommodating and chemical re-positioning of reactive side-chains and the substrate, with only small perturbations to the electric field sensed by the reactive C=O, to fully utilize these naturally evolved electric fields for catalysis. This interplay of electrostatic and chemical positioning to modulate the protein fitness landscape may be a general observation, as evident in the changes in active site heterogeneity in the directed evolution of *de novo* Kemp eliminases and ancestrally reconstructed β-lactamases.^57, 63–64^ Combining the complementary approaches of MD simulations and the VSE with kinetic studies enables a more comprehensive molecular picture of the parameters modulating an enzyme’s evolution towards new and/or improved function with implications in protein design, enzyme evolution, and antibiotic resistance.

## Supporting Information

Experimental details and methods for protein expression and purification, kinetics, steady-state MS, FTIR spectroscopy, and MD simulations. Additional discussions of the solvatochromic calibration for the VSE, difference spectra processing, IR peak determination and evaluation, choice of mutations for VSE experiments, and utilization of MD simulations to guide experimental findings. Additional figures, tables, and schemes relating to the kinetics, activation free energy barriers, FTIR spectral processing, FTIR lineshape parameters, structural comparisons from MD simulations, analysis of MD simulations, and alternative representations of evolutionary changes from MD simulations.

## Acknowledgement

We acknowledge helpful discussions with Peter Kasson at the UVA and the Vincent Coates Foundation Mass Spectrometry Laboratory at Stanford University. We thank extensive feedback from Dr. Zhe Ji, Jacob Kirsh, and Jared Weaver during the drafting process. J.K. acknowledges the Deutsche Forschungsgemeinschaft for a Research Fellowship (KO5464/1). This work is supported in part by NIH Grant GM118044 (to S.G.B.).

