## Supplementary Information for "The Interplay of Electrostatics and Chemical Positioning in the Evolution of Antibiotic Resistance in TEM β-Lactamases"

### Extended Data and Supplementary Information for
### The Interplay of Electrostatics and Chemical Positioning in the Evolution of Antibiotic Resistance in TEM β-Lactamases

Samuel H. Schneider, Jacek Kozuch, and Steven G. Boxer

#### Table of Contents:

##### *Materials & Methods:*

**Nucleotide sequence for TEM-1 β-lactamase.**

**Protein expression and purification.**

**UV-vis kinetics.**

**Steady-state mass spectrometry.**

**FTIR spectroscopy.**

**MD simulation parameterization.**

**MD simulations.**

##### *Extended Figures, Tables, and Schemes:*

**Scheme S1.** Expanded reaction mechanism for TEM β-lactamase with PenG.

**Table S1.** Kinetics of TEM β-lactamase-catalyzed hydrolysis of PenG and CTX from steady-state kinetics and mass spectrometry.

**Table S2.** Activation free energy barriers for the TEM β-lactamase reaction with PenG and CTX.

**Figure S1.** Absorbance spectra and extinction coefficient determination for PenG and CTX.

**Figure S2.** Representative time-trace data for PenG hydrolysis by TEM-1 S70A β-lactamase.

**Figure S3.** Representative steady-state MS charge envelope of TEM-1 with PenG.

**Figure S4.** Example kinetic parameter distributions determined using bootstrapping algorithm for TEM-1 β-lactamase with CTX.

**Figure S5.** FTIR spectra of $^{12}C_{CO}$ and $^{13}C_{CO}$ β-lactams in aqueous buffer.

**Figure S6.** Electric field-frequency calibration for model compounds and their β-lactam antibiotic mechanistic equivalents.

**Figure S7.** Full isotope-edited IR difference spectroscopy of the TEM β-lactamases with PenG and CTX.

**Figure S8.** Representative example of FTIR data processing and assignment with TEM-52 E166N and CTX.

**Table S3.** TEM-# S70G PenG vibrational frequencies and lineshape parameters from curve-fitting.

**Table S4.** TEM-# S70G CTX vibrational frequencies and lineshape parameters from curve-fitting.

**Table S5.** TEM-# E166N PenG vibrational frequencies and lineshape parameters from curve-fitting.

**Table S6.** TEM-# E166N CTX vibrational frequencies and lineshape parameters from curve-fitting.

**Figure S9.** Mutational effect on IR spectra of TEM-1 β-lactamase with PenG.

**Figure S10.** MD Methodology for Structure-Guided Interpretation of Kinetic and VSE Results.

**Figure S11.** Structural comparison of tight-binding PenG complexes with S70G and E166N background mutants and WT TEM β-lactamases.

**Figure S12.** Structural comparison of weak-binding PenG complexes with S70G background mutations and WT TEM β-lactamases.

**Figure S13.** Structure-guided approach to inferring catalytically relevant electric fields.

**Figure S14.** Biased MD simulations of WT TEM β-lactamases as informed from kinetically compromised variants.

**Scheme S2.** Atoms and Interactions Considered in the MD Analysis.

**Table S7.** PenG-dependent average changes in active site RMSFs due to evolution.

**Table S8.** CTX-dependent average changes in active site RMSFs due to evolution.

**Table S9.** Collective RMSF changes in the evolution of TEM-1 to TEM-52 for PenG and CTX.

**Table S10.** PenG-dependent average changes in active site distances due to evolution.

**Table S11.** CTX-dependent average changes in active site distances due to evolution.

**Table S12.** Collective distance changes in the evolution of TEM-1 to TEM-52 for PenG and CTX.

**Scheme S3.** Angle of nucleophilic attack in the ES and AE complexes.

**Figure S15.** Alternative probability density representation of significant active site distance changes over the course of evolution from TEM-1 to TEM-52 for PenG and CTX.

**Figure S16.** Alternative probability density representation of significant active site RMSF changes over the course of evolution from TEM-1 to TEM-52 for PenG and CTX.

**Table S13.** Average changes in active site RMSFs between TEM-1 and TEM-52 with PenG and CTX, respectively.

**Table S14.** Average changes in active site distances between TEM-1 and TEM-52 with PenG and CTX, respectively.

**Figure S17.** Alternative representation of significant active site RMSF and distances changes over the course of evolution from TEM-1 to TEM-52 for PenG and CTX.

**Figure S18.** Structural comparison of TEM-1 PenG versus TEM-52 CTX complexes.

**Figure S19.** Viscosity-dependence of PenG hydrolysis by TEM β-lactamases.

**Table S15.** Evolutionary change in on- and off-rates of PenG hydrolysis with TEM-1 and TEM-52.

#### Supplementary References:

#### *Extended Data and Scripts for Analysis:*

**MD Files and Parameters:**

**MATLAB Code:**

- MATLAB script for kinetic fitting of β-lactam hydrolysis using the Lambert function.
- MATLAB script for quantifying the [ES]/[AE] ratios from steady-state MS.
- MATLAB script for kinetic determination of $k_2$ and $k_3$ using boot-strapping algorithm.

#### Methods:

##### Nucleotide sequence for TEM-1 β-Lactamase:

The pBAD plasmid containing TEM-1 (pBAD-TEM-1) β-lactamase was kindly given to us by Patrice Soumillion at the Université Catholique de Louvain.[1-2] Standard PCR site-directed mutagenesis was performed on the plasmid using commercial QuikChange Lightning (Agilent) kits and protocols, which was transformed into DH10B *E. coli* cells with 15 µg/mL tetracycline HCl selection on agar plates. This was used to introduce the evolutionarily derived mutations (E104K, G238S, M182T) and kinetically incompetent variants (S70G or E166N) in addition to a cleavable His-tag in TEM-1 β-lactamase.

ATGGGTAGTCAACATTTCCGTGTCGCCCTTATTCCCTTTTTTGCGGCATTTTGCCTTCCTGT
TTTTGCTCACCCAGAAACGCTGGTGAAAGTAAAAGATGCTGAAGATCAGTTGGGTGCACGA
GTGGGTTACATCGAACTGGATCTCAACAGCGGTAAGATCCTTGAGAGTTTTCGCCCCGAAG
AACGTTTTCCAATGATGAGCACTTTTAAAGTTCTGCTATGTGGCGCGGTATTATCCCGTGTT
GACGCCGGGCAAGAGCAACTCGGTCGCCGCATACACTATTCTCAGAATGACTTGGTTGAG
TACTCACCAGTCACAGAAAAGCATCTTACGGATGGCATGACAGTAAGAGAATTATGCAGTG
CTGCCATAACCATGAGTGATAACACTGCGGCCAACTTACTTCTGACAACGATCGGAGGACC
GAAGGAGCTAACCGCTTTTTTGCACAACATGGGGGATCATGTAACTCGCCTTGATCGTTGG
AATCCGGAGCTGAATGAAGCCATACCAAACGACGAGCGTGACACCACGATGCCTGTAGCA
ATGGCAACAACGTTGCGCAAACTATTAACTGGCGAACTACTTACTCTAGCTTCCCGGCAAC
AATTAATAGACTGGATGGAGGCGGATAAAGTTGCAGGACCACTTCTGCGCTCGGCCCTTC
CGGCTGGCTGGTTTATTGCTGATAAATCTGGAGCCGGTGAGCGTGGGTCTCGCGGTATCA
TTGCAGCACTGGGGCCAGATGGTAAGCCCTCCCGTATCGTAGTTATCTACACGACGGGGA
GTCAGGCAACTATGGATGAACGAAATAGACAGATCGCTGAGATAGGTGCCTCACTGATTAA

GCATTGGGCTCTAGTACCAAGAGGCAGCTCAGAAGAGGATCTGAATAGCGCCGTCGACCA

TCATCATCATCATCAT

**Protein Expression and Purification:**

The pBAD-TEM plasmids were transformed into One Shot TOP10 Chemically Competent *E. coli* cells (Thermo-Fisher) using selection with 15 µg/mL Tetracycline HCl on Luria Broth (Fisher) agar plates. Unless otherwise specified, all variants of TEM β-lactamase, wild-type (S70S, E166E), acylation-impaired (S70G), and deacylation-impaired (E166N), were prepared and purified identically. A single-colony of transformed cells was grown into 5 mL overnight cultures using Terrific Broth (Fisher) with 15 µg/mL Tetracycline HCl at 37C. Overnight cultures were then inoculated into 1 L of Terrific Broth (Fisher) media with 10 µg/mL Tetracycline HCl shaking at 200 rpm and 37C until they reached an OD600 ~ 0.6 at which point protein expression was induced with 2 g per L of L-(+)-Arabinose (Sigma), and grown for 4-5 hrs at 27 C. Cells were harvested by centrifugation at 6000x g's for 30 mins and resuspended in lysis buffer (50 mM potassium phosphate, 20 mM imidazole, 500 mM sodium chloride, 10% (v/v) glycerol, pH 7.4). Cells were lysed by homogenization, and the lysate centrifuged twice for 90 mins each at 15,000x g's before sterile filtering the supernatant. The crude protein was then purified using a Ni-NTA affinity resin column, washing with 50 mM potassium phosphate (pH 7.4), 50 mM imidazole, 500 mM sodium chloride and eluted using 50 mM potassium phosphate (pH 7.4), 200 mM imidazole, 500 mM sodium chloride. Further purification was performed using anion exchange chromatography on a 5 mL HiTrap-Q HP (GE Healthcare) column and eluted using a 0-40% gradient of Buffer A (25 mM Tris (pH 8.4) 25 mM sodium chloride) to Buffer B (25 mM Tris (pH 8.4) 1 M sodium chloride) over 25 column volumes. Purified protein was exchanged into a cryoprotectant-containing storage buffer (50 mM $KP_i$ (pH 7.4), 100 mM NaCl, 10% (v/v) glycerol) for long-term storage at -80C.

TEM-1 (WT) β-Lactamase with all permutation sites (Ambler numbering[3]) highlighted for clarity based on the key.

Key: Cyan = Signal Peptide (removed *in situ*); Red = S70; Pink = E166; Green = E104; Blue = G238; Yellow = M182.

MGSQHFRVALIPFFAAFCLPVFAHPETLVKVKDAEDQLGARVGYIELDLNSGKILESFRPEERFP
MMSTFKVLLCGAVLSRVDAGQEQLGRRIHYSQNDLVEYSPVTEKHLTDGMTVRELCSAAITMS
DNTAANLLLTTIGGPKELTAFLHNMGDHVTRLDRWEPELNEAIPNDERDTTMPVAMATTLRKLL
TGELLTLASRQQLIDWMEADKVAGPLLRSALPAGWFIADKSGAGERGSRGIIAALGPDGKPSRI
VVIYTTGSQATMDERNRQIAEIGASLIKHWALVPRGSSEEDLNSAVDHHHHHH

**UV-vis kinetics:**

All TEM kinetics with either PenG or CTX were performed using identical buffers and preparations unless otherwise mentioned. Before kinetic measurements, the concentration of the original purified TEM β-lactamase was determined using the absorbance at 280 nm ($\varepsilon_{280}$ = 28,085 $M^{-1}$ $cm^{-1}$)[4] in 50 mM potassium phosphate (pH 7.0). All buffer solutions for kinetic measurements contained 50 – 200 mM potassium phosphate (pH 7.0) and bovine serum albumin (BSA; Sigma) at 0.02 – 0.05% (w/v) in order to assist with passivation of all cuvette surfaces for the low TEM concentrations used, typically between 10 – 50 nM for PenG and 10 – 500 nM for CTX hydrolysis. Prior to use, all cuvettes and rapid-mixing tubing were passivated for at least 30 mins with a 1% (w/v) BSA solution. We assume the simplest mechanism of class A β-lactamase-catalyzed hydrolysis according to (c.f. Figure 1):[5-6]

$$\mathbf{E + S} \underset{k_{-1}}{\overset{k_1}{\rightleftharpoons}} \mathbf{ES} \xrightarrow{k_2} \mathbf{EA} \xrightarrow{k_3} \mathbf{E + P}$$

E, S, ES, AE, and P correspond to the free enzyme, substrate, non-covalent Michaelis complex, covalent acyl-enzyme complex, and product, respectively. The apparent rate constant from steady-state kinetics corresponds to the microscopic rate constants according to Eq. 1 and 2:

Hydrolysis of PenG by TEM β-lactamases was monitored on a Cary6000i UV-vis-NIR spectrometer equipped with a stopped-flow accessory (SFA-20 Rapid Mixing Accessory from TgK Scientific). Substrate loss was monitored at 232 nm ($\Delta\varepsilon_{232}$ = -940 $M^{-1}$ $cm^{-1}$)[7] every 0.2 secs until completion of the reaction (i.e. $\Delta A/\Delta t = 0$). A full kinetic decomposition into $k_{cat}$ and $K_M$ was determined using the closed-form solution to the Michaelis-Menten equation (Eq. S1), which utilizes the Lambert function (W) to solve for the time-dependent change in substrate concentration ($S_t$ and $S_0$),[8-9]

$$S_t = K_M * W\left[\frac{S_0}{K_M} exp\left(\frac{S_0 - k_{cat}t}{K_M}\right)\right] \qquad \text{(Eq. S1)}$$

Kinetic parameters $k_{cat}$ and $K_M$ were evaluated using a MATLAB script (see SI) and measurements were repeated 7 – 9 times and for at least two solute concentrations. Kinetic parameters presented in Table 1 represent the average and standard deviation across all measurements and solute concentrations.

Hydrolysis of CTX by TEM β-lactamases was monitored on a Lambda 25 (Perkin-Elmer) UV-vis spectrometer at 262 nm, corresponding to ring-opening of the β-lactam ($\Delta\varepsilon_{262}$ = -7250 $M^{-1} cm^{-1}$),[7] for 16 substrate concentrations (15-1500 μM). Initial rates were determined for the first 10-30 secs of the single-wavelength kinetic trace for all substrate concentrations. Standard Michaelis-Menten plots were utilized to determine kinetic parameters for TEM-19, -15, and -52. Kinetic constants for TEM-1 and TEM-17 were determined using Lineweaver-Burke analysis because of the high $K_M$ (ca. 1-2 mM), such that rates corresponding to saturating substrate concentrations (i.e. rate approaches $k_{cat}$) could not be measured. Kinetic parameters were averaged from three separate fits and errors represent the standard deviations across measurements.

**Steady-state mass spectrometry:**

Mass spectrometry (MS) was performed in the Stanford University Mass Spectrometry facility on a Waters Single Quadrupole LC-ESI/MS instrument. All protein LC-MS for assessing purity was performed at ~ 20 μM concentrations in water following ion-exchange chromatography and assessed using either the Intact Mass (Protein Metrics) or MassLynx (Waters) software suites.

Measurement of the proportions of non-covalent and covalent TEM-substrate complexes were performed similarly to that of Saves *et al.*[10] TEM β-lactamase was added to a solution containing an excess of substrate in 50 – 200 mM potassium phosphate buffer (pH 7.0), mixed gently, and then the reaction was quenched through addition of one volume equivalent of methanol. Concentrations were chosen to ensure that the steady-state reaction would be observed for at least 20 – 40 secs and final enzyme concentrations would be approximately 20-50 μM in at least 50 μL for sufficient LC-MS signal-to-noise. After quenching the reaction, the solution was buffer exchanged using a 3,000 Da MWCO Amicon (Millipore) spin filter to achieve a 1:1 (v/v) mixture of water/methanol before quantification using LC-MS on a reverse-phase C8 column (2.1 x 30 mm Agilent Zorbax columns with 3.5 μm Stablebond stationary phases; ESI+, 50-2000 m/z, standard cone voltage). The HPLC mobile phases consists of water and acetonitrile with 0.1% formic acid.

The relative abundance of non-covalent (ES) and covalent (AE) complexes (+ 334 or +413 Da, respectively) was determined from analysis of the unprocessed m/z data extracted using the MassLynx software and the mass envelope intensities quantified using a MATLAB script (see Figure S3 and SI).

While quantification of the [AE]/[ES] ratios may be normally distributed, the distributions of rate constants are not necessarily since both $k_2$ and $k_3 > k_{cat}$, i.e. $0 \leq [AE]/[ES] \leq$ infinity. As such, we utilize the uncertainty in both $k_{cat}$ and [AE]/[ES] ratios to bootstrap a distribution of possible $k_2$ and $k_3$ values and determine the median and appropriate confidence intervals. For comparison to normally distributed statistics we report the 68% (corresponding to 1 standard deviation in a normal distribution) and 95% confidence intervals (CI) to determine $k_2$ and $k_3$ and corresponding uncertainties. Example distributions are shown Figure S4 and the MATLAB scripts used to perform the bootstrapping is presented in the SI.

Control experiments to determine the ionization efficiency of the ES and AE complexes were performed by mixing an equal amount of kinetically incompetent protein TEM-1 E166N (which is not observed to deacylate) in the presence or absence of PenG or CTX, but otherwise treated identically as above. An equal concentration of protein treated with and without β-lactam were mixed and measured via LC-MS in order to quantify the ratio of ionization efficiencies ([AE]/[ES]), which was observed to be 1.0 ± 0.1 (n=3) based on the ratio of integrated peak intensities, as quantified above (see Figure S3,4). Similarly, no hydrolysis of the acylated TEM-1 E166N complex with either PenG or CTX was observed in the water:methanol mixtures, as no detectable non-covalent TEM-1 E166N was observed via LC-MS.

Note that the less thermodynamically stable TEM variants, i.e. TEM-19 and TEM-15, were more prone to unfolding in water/methanol solutions as determined via loss of concentration after buffer exchange. This could be ameliorated through lower methanol concentrations, thereby reducing the extent of unfolding, and no change in [AE]/[ES] ratios as determined from LC-MS were observed with water:methanol ratios (v/v) of 3:1 or 7:1.

**FTIR spectroscopy:**

FTIR spectra were recorded on a Bruker Vertex 70 spectrometer with a liquid nitrogen-cooled mercury cadmium telluride (MCT) detector; all data processing was performed using the OPUS program (Bruker). A liquid cell was prepared using two $CaF_2$ optical windows (19.05 mm diameter, 3 mm thickness, Lambda Research Optics, Inc.), separated by two semicircular Teflon spacers (25 and 50 μm thickness). FTIR spectra of $^{12}C$- and $^{13}C$-PenG and CTX, and all TEM-substrate complexes were acquired in a $D_2O$ buffer (50 mM $KP_i$, 100 mM NaCl, pD 7.0) through averaging 256-512 scans from 4000-1000 $cm^{-1}$ with 1 $cm^{-1}$ resolution after 600 secs of dry air purging.[11]

All protein-substrate measurements were performed similarly with a final concentration of ca. 2.3 mM TEM and a slight excess of substrate (ca. 2.5 – 3.0 mM). In the case of the TEM S70G mutants with PenG, where slow hydrolysis can be observed on the timescale of the FTIR measurements, an excess of substrate was added (ca. 10 – 25 mM PenG). Before FTIR measurements, proteins were exchanged overnight into the $D_2O$ buffer at 4 C and were exchanged again the following day into fresh $D_2O$ buffer to achieve optimal H/D exchange of the protein.

In order to obtain isotope-edited difference spectra ($^{12}C$ – $^{13}C$), measurements were performed on the same day from a single batch of doubly-$D_2O$-exchanged protein, which ensures minimal difference between the two measurements. A brief description of the process for generating the final isotope-edited difference spectra is described below. First, the $^{13}C$ spectrum was scaled and subtracted from the $^{12}C$ spectrum to remove the protein absorption. Second, residual unbound $^{12}C$- and $^{13}C$-β-lactam was subtracted from the difference spectra using either PenG or CTX in $D_2O$ buffer as a reference. Third, residual changes due to H/D back-exchange of either the bound or unbound protein complex was subtracted using either ($^{12}C$ – $^{12}C$) or ($^{13}C$ – $^{13}C$) difference spectrum in the former and a protein-only reference spectrum in $D_2O$ buffer in the latter. Finally, based on $2^{nd}$-derivative analysis and the $^{12}C$ – $^{13}C$

β-lactam C=O isotope frequency shift of ~45 $cm^{-1}$, sets of peaks (a positive and a negative feature) were identified and baselined conservatively. This process was repeated for multiple difference spectra and subsequent baselined spectra were averaged ($n \geq 5$) to generate those presented in Figure 3 (see Table S3-6). For further details and specific examples of the spectral processing see Figure S8.

**MD simulation parameterization:**

The initial structures for all simulations were based on the crystal structure of the TEM1(E166N)-Penicillin G acyl-enzyme complex (PDB id: 1FQG)[12]; mutants of interest were generated via the mutation option in PyMOL 2.3.2. Then, the PenG-acylated S70 was exchanged to a serine residue and amber99-ffSBildn parameters for the resulting 6 structures (TEM-1, TEM1-S70G, TEM-1 E166N, TEM-52, TEM-52 S70G, TEM-52 E166N) containing the nucleophilic active site water were obtained using the pbd2gmx option in GROMACS 2018.[13-14] Parameters for the ligands PenG and CTX were generated using AmberTools18 (ff99SB-ildn force field and AM1-BCC) based on optimized structures from DFT simulations in Gaussian09.A (b3lyp/6-311g**(2d,2p)).[15] The PenG-acylated and CTX-acylated serine residues, needed for the simulations of the acyl-enzyme states, were parameterized analogously using acetyl and methyl amine caps on the N- and C-termini of the serine group. In order to import the parameters into the protein topology files the caps were removed to obtain the acylated PenG-serine and CTX-serine building blocks. Then, backbone partial charges were adjusted to those of the serine parameters, and finally the charges of $C_\beta$ and its H atoms were minimally adjusted to ensure a correct net charge of the acylated serines. Finally, when implementing the new acylated serines into the respective protein topology file, the original structural parameters of serine (besides those containing the hydroxyl H) were maintained and all additional parameters that include the PenG or CTX moiety, but exclude the serine fragment, were added to the topology file.

**MD simulations:**

MD simulations were performed using GROMACS 2018,[16] and were closely related to the work of the Kasson lab.[17-19] Protein structures of all variants were based on the structure of TEM-1 E166N PenG acyl-enzyme complex (PDB ID: 1FQG) as explained above. For all acyl-enzyme simulations, the starting structure for the PenG-acylated S70 was taken as is from the PDB file. PenG's starting structure in the ES complex was adapted to the same conformation: starting from the DFT optimized geometry, PenG's core was positioned such that the β-lactam C=O was located in the oxyanion hole, similar to the acyl-enzyme, and torsion angles of side-chains were changed to resemble those in the PenG-acylated case. The CTX-acylated S70 geometry was taken from the acyl-enzyme structure of Toho-1 E166A (PDB ID: 1IYO).[20] CTX's ES geometry was adapted from 1IYO using the same approach as for PenG. In this way, in all structures the β-lactam C=O was located at H-bond length of ca. 3 Å to both nitrogens of A237 and S/G70. Each protein was positioned in the center of a box with a dodecahedral unit cell of a size corresponding to a cube with walls located at distance of at least 2 nm from the protein surface. The box was filled with ca. 20500 water molecules (tip3p) and an ion strength of 0.175 M (similar to experiments) was generated using sodium and chloride ions, such that the system is net charge neutral. Each system was energy minimized using the steepest descent algorithm until all forces were below 1000 kJ $mol^{-1}$ $nm^{-1}$, and then equilibrated (NVT and NPT steps, 100 ps each with protein and ligand heavy atoms restrained; second NPT over 100 ps with Cα atoms and ligand heavy atoms restrained). Here, the velocity-rescaling theromostat and Berendsen barostat were used (300K and 1 bar) with short range interactions truncated to 1.2 nm, long range electrostatics treated with Particle Mesh Ewald, and hydrogen bonds constrained used LINCS. MD production runs (using the Parrinello-Rahman barostat) were carried out over a total of 1 µs for each of the 16 structures. If the substrate left the active site (in simulations of ES states), the simulation was aborted and continued from a structure with bound ligand. In order to test (a) the stability of each of the observed conformations of the substrate (assessed via the 2D correlation plots of the oxyanion H-bond distances; Fig 4) and (b) the possibility to access additional conformations due to possible sampling of a local minimum, for example starting WT TEM-1 in a conformation only observed in TEM-1 S70G, representative structures were extracted, and ligand conformations were implanted into proteins for additional MD simulations. Each structure was then newly minimized and equilibrated, and then underwent five 50 ns MD simulations. Average structures, rms fluctuations, distances, etc. were extracted using the tools present in the GROMACS package and additional add-ons (e.g. GROmaps[21]). 2D correlation plots were generated in OriginLabs2018.

The quantification of changes in structural displacement, either in terms of the root mean square fluctuations (RMSF) or distances between atoms, were quantified from representative trajectories and average structures (n = 3-5, each 50-200 ns), respectively, from the WT MD simulations of TEM-1 and TEM-52 with PenG and CTX in the ES and AE states (see Scheme S2 and Tables S5-8 for further details). Across each comparison, either RMSF or distance, values that were statistically significant ($p < 0.05$) were determined from a two-tailed student's t-test (see t-values for given degrees of freedom (df). Significant values were then utilized in the corresponding averages for each set of regions or interactions in the active site: K73/N132/E166/N170, oxyanion hole, β-lactam carboxylate, or β-lactam (Figure 5, 6, S12). This approach highlights the complexity of evaluating structural perturbations arising from both mutational changes over evolution and those that are substrate specific.

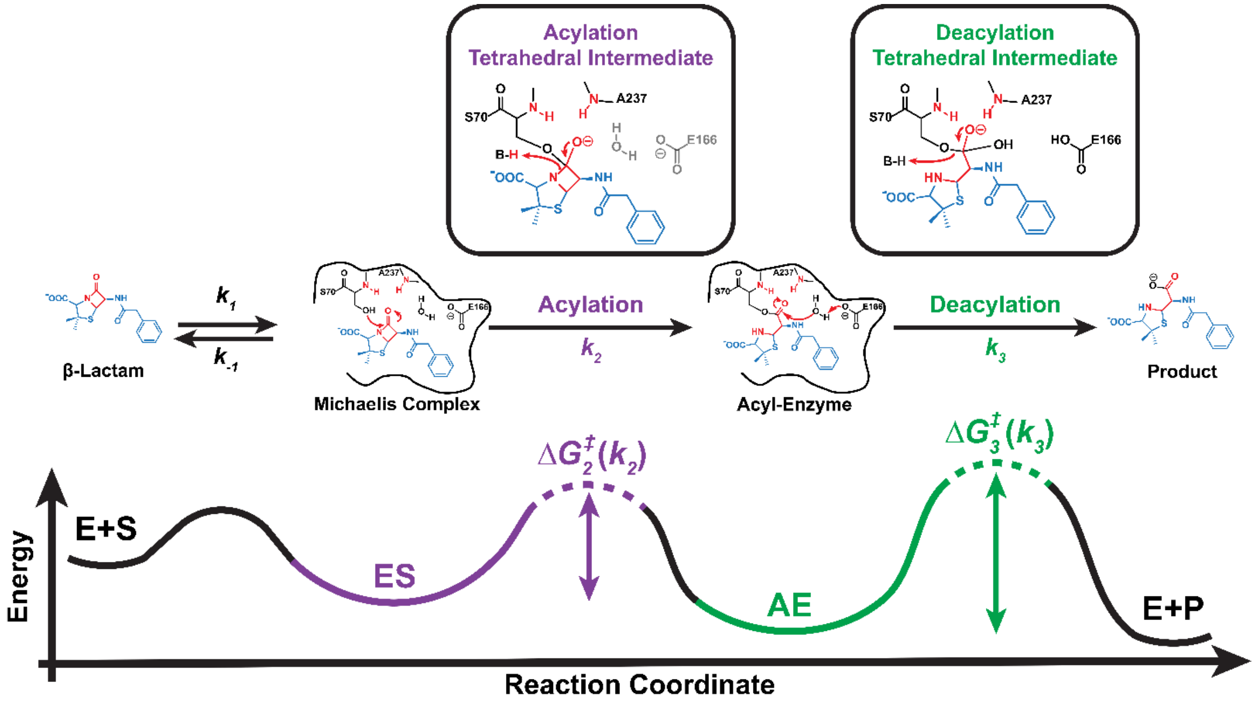

***Scheme S1. Expanded reaction mechanism for TEM β-lactamase with PenG.*** *The mechanism, proposed tetrahedral intermediates, and reaction coordinate involved in the acylation and deacylation processes.*[1] *The alkoxide tetrahedral intermediates are presumed to be similar, i.e. in terms of both chemical and electronic structure, to the TS for each chemical transformation (dotted lines) and illustrate the mechanism by which an electric field and the oxyanion hole can preferentially stabilize the TS relative to the reactant-state via electrostatic preorganization (Eq. 1). Barrier heights are not drawn to scale.*

***Table S1. Summary of kinetic data for TEM β-lactamase-catalyzed hydrolysis of PenG and CTX based on steady-state kinetics and mass spectrometry.***

| | PenG | | | | CTX | | | |
|---|---|---|---|---|---|---|---|---|
| **Variant** | $k_{cat}$ ($s^{-1}$) | $K_M$ (μM) | $k_2$ ($s^{-1}$)[b] | $k_3$ ($s^{-1}$)[b] | $k_{cat}$ ($s^{-1}$) | $K_M$ (μM) | $k_2$ ($s^{-1}$)[b] | $k_3$ ($s^{-1}$)[b] |
| **TEM-1** | 1816.2 ± 18.0 | 25.6 ± 8.0 | 5514.7 [4641.0, 6318.2] | 2707.5 [2545.4, 2985.6] | 3.9 ± 0.2 | 2051.3 ± 168.2 | 4.1 [3.8, 4.4] | 93.1 [38.7, 248.9] |
| **TEM-17** | 1650.5 ± 36.9 | 30.6 ± 0.3 | 8559.5 [5779.5, 11898.6] | 2047.0 [1908.3, 2314.7] | 10.8 ± 1.4 | 1189.0 ± 214.9 | 11.8 [10.2, 13.3] | 133.3 [99.0, 187.2] |
| **TEM-19** | 105.9 ± 4.4 | 2.5 ± 0.5 | 262.1 [236.6, 291.3] | 177.7 [165.2, 192.4] | 69.7 ± 7.1 | 281.2 ± 40.4 | 88.1 [77.7, 99.4] | 334.1 [259.1, 462.7] |
| **TEM-15** | 65.8 ± 2.6 | 1.9 ± 0.7 | 1620.1 [891.0, 3218.2][a] | 68.9 [65.7, 72.8][a] | 116.1 ± 3.6 | 76.3 ± 8.2 | 137.6 [125.7, 154.8] | 777.2 [467.2, 1477.6] |
| **TEM-52** | 42.9 ± 0.2 | 1.3 ± 0.2 | 276.8 [188.2, 342.4] | 50.9 [49.1, 55.6] | 89.6 ± 2.5 | 71.8 ± 11.5 | 110.4 [101.9, 121.9] | 483.1 [345.7, 721.4] |

[a] *Minimal detectable non-covalent species, such that $k_3 \approx k_{cat}$, only values of [AE]/[ES] < 50 were considered in rate-constant determination due to the signal-to-noise of LC-MS data.*

[b] *Rate-constants and uncertainties (68.25% confidence interval (CI) equivalent to 1σ) for $k_2$ and $k_3$ are the median value as determined through propagation of $k_{cat}$ values as well as the distribution of [AE]/[ES] values for each charge state of the protein charge envelope via LC-MS (see Figure S3-4).*

**Table S2. Activation free energy barriers for the TEM β-lactamase reaction with PenG and CTX.**

| | PenG | | | CTX | | |
|---|---|---|---|---|---|---|
| **Variant** | **Overall** $\Delta G^{\ddagger}_{kcat}$ **(kcal/mol)**[a] | **Acylation** $\Delta G^{\ddagger}_{k2}$ **(kcal/mol)**[a,b] | **Deacylation** $\Delta G^{\ddagger}_{k3}$ **(kcal/mol)**[a,b] | **Overall** $\Delta G^{\ddagger}_{kcat}$ **(kcal/mol)**[a] | **Acylation** $\Delta G^{\ddagger}_{k2}$ **(kcal/mol)**[a,b] | **Deacylation** $\Delta G^{\ddagger}_{k3}$ **(kcal/mol)**[a,b] |
| **TEM-1** | **13.00 ± 0.01** | **12.34 [12.44, 12.26]** | **12.76 [12.80, 12.70]** | **16.64 ± 0.04** | **16.61 [16.65, 16.57]** | **14.76 [15.28, 14.18]** |
| **TEM-17** | **13.06 ± 0.02** | **12.08 [12.31, 11.89]** | **12.93 [12.97, 12.86]** | **16.04 ± 0.04** | **15.98 [16.07, 15.91]** | **14.55 [14.72, 14.34]** |
| **TEM-19** | **14.68 ± 0.02** | **14.15 [14.21, 14.08]** | **14.38 [14.42, 14.33]** | **14.93 ± 0.06** | **14.79 [14.87, 14.72** | **14.00 [14.15, 13.81]** |
| **TEM-15** | **14.96 ± 0.02** | **13.07 [13.42, 12.66]** | **14.94 [14.96, 14.90]** | **14.63 ± 0.02** | **14.53 [14.58, 14.46]** | **13.50 [13.80, 13.12]** |
| **TEM-52** | **15.22 ± 0.01** | **14.11 [14.34, 13.99]** | **15.12 [15.14, 16.06]** | **14.78 ± 0.02** | **14.66 [14.70, 14.60]** | **13.78 [13.98, 13.55]** |

[a] *Activation free energy barriers are calculated using TS-theory at 298 K according to* $\Delta G^{\ddagger} = -RT\ln\left(k\frac{h}{k_BT}\right)$*, where k is the observed rate-constant,* $k_B$ *is the Boltzmann constant, h is Planck's constant, T is temperature (K), R is the gas constant.*

[b] *Activation free energy barrier values and uncertainties (68.25% confidence interval (CI) equivalent to 1σ) for* $k_2$ *and* $k_3$ *are the median value as determined through propagation of* $k_{cat}$ *values as well as the distribution of [AE]/[ES] values for each charge state of the protein charge envelope via LC-MS (see Figure S3-4, Table S1).*

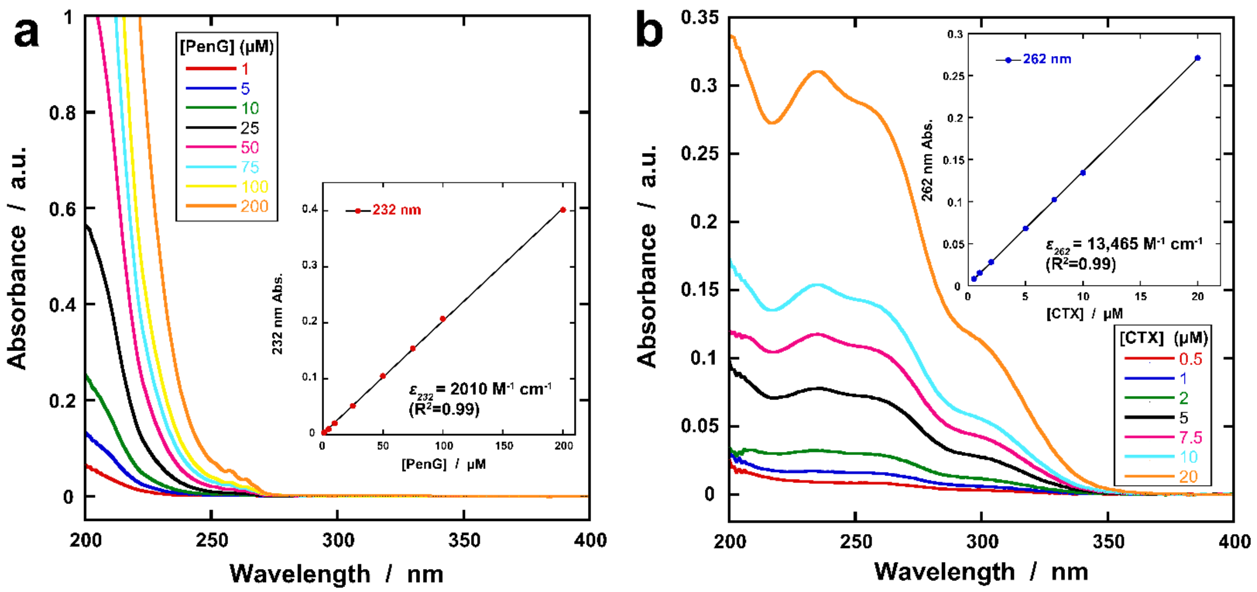

***Figure S1. Absorbance spectra and extinction coefficient determination for PenG and CTX.*** *Increasing concentrations of* ***(a)*** *PenG and* ***(b)*** *CTX in phosphate buffer (pH 7.0). Insets contain extinction coefficient determination information as determined from single-wavelength analysis as a function of concentration. The extinction coefficients at 232 and 262 nm, for PenG and CTX, respectively, are approximately 2-fold higher than those utilized for monitoring hydrolysis (i.e. substrate loss is quantified based on* $\Delta\varepsilon$ *= -940 and -7250* $M^{-1}$ $cm^{-1}$*, respectively).*[2]

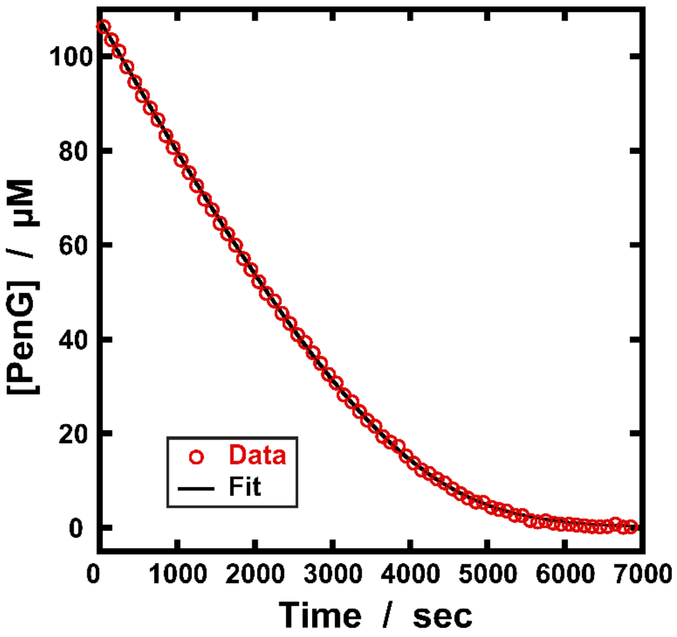

***Figure S2. Representative time-trace data for PenG hydrolysis by TEM-1 S70A β-lactamase.*** *Data (red circles) is shown relative to the analytical fit (black line) based on the closed-form solution to the Michaelis-Menten equation. Kinetic parameters:* $k_{cat}$ *= 0.0105 ± 0.0001* $sec^{-1}$*,* $K_M$ *= 23.35 ± 0.04 µM were determined using analytical solution (see MATLAB scripts) and errors are determined from the Jacobian and covariance matrix.*

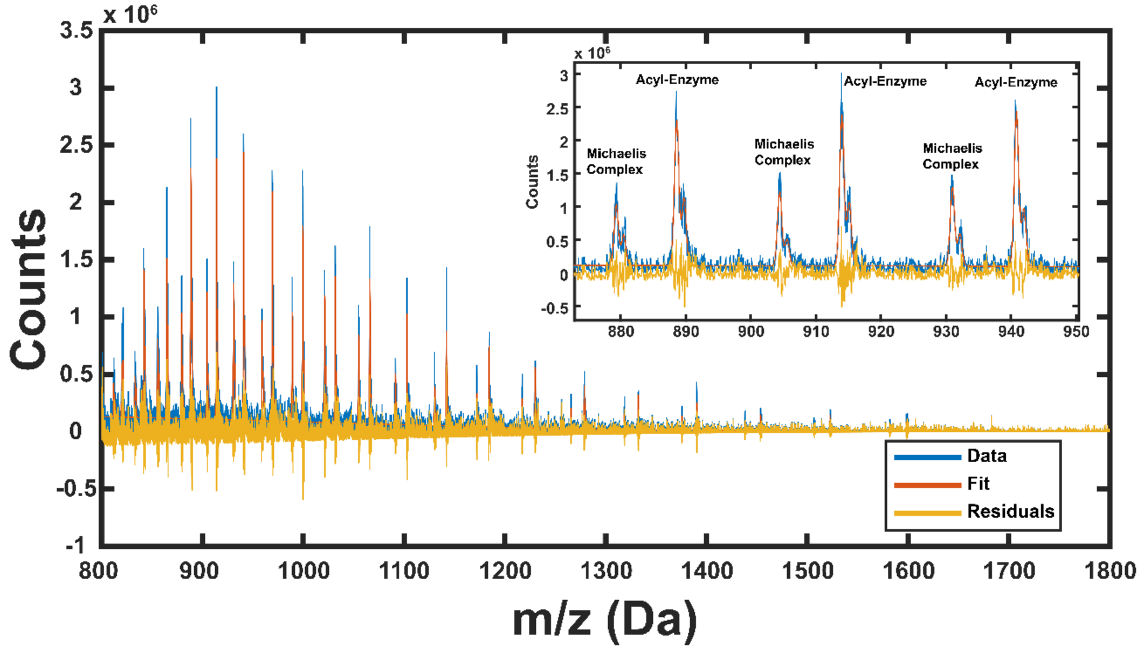

***Figure S3. Representative steady-state MS charge envelope of TEM-1 with PenG for kinetic deconvolution.*** *The relative abundance of the ES and AE complexes can be followed in MS following the general approach and methods of Saves et al.[3] The corresponding raw data (blue), overall fit (orange), and fitting residuals (yellow) for the steady-state quenched TEM-1 with PenG. Inset displays distribution pattern for Michaelis complex and acyl-enzyme for a given charge state of the overall protein charge envelope. The relative abundance of non-covalent (ES) and covalent (AE) complexes (+ 334 or +413 Da for PenG and CTX, respectively) was determined similarly to Saves et al.[3] from analysis of the unprocessed m/z data extracted using the MassLynx software and the mass envelope intensities quantified using a MATLAB script (See SI MATLAB code) for each charge state (z=20 – 39) of the charge envelope (also referred to as charge state distribution). For each charge state in the protein charge envelope, a series of gaussian bands corresponding to the non-covalent and covalent complexes were fit with equal width but adjustable intensities in order to compare the integrated peak areas within a given charge state for [AE]/[ES] determination. A broad gaussian baseline was applied throughout to improve the fits. Ratios of [AE]/[ES] were acquired across 3-5 independent sample preparations, and [AE]/[ES] values from each charge state were utilized to determine the overall [AE]/[ES] ratios and subsequently $k_2$ and $k_3$ using a bootstrapping algorithm as described in Figure S4. The relative ionization efficiencies of the complexes ([AE]/[ES]), was determined to be 1.0 ± 0.1 (n=3) by mixing an equal amount of kinetically incompetent protein TEM-1 E166N (which is not observed to deacylate) both with and without PenG or CTX, but*

*otherwise treated as described in the Methods, and measured via LC-MS. There was no apparent hydrolysis of the acylated TEM-1 E166N complex with either PenG or CTX in the water:methanol mixtures, as no detectable non-covalent TEM-1 E166N was observed via LC-MS. Note that the less thermodynamically stable TEM variants, i.e. TEM-19 and TEM-15, were more prone to unfolding in water/methanol solutions as determined via loss of concentration after buffer exchange, indicated by transfer through the 3k molecular weight cut-off (MWCO) filter. This could be ameliorated through lower methanol concentrations, thereby reducing the extent of unfolding, and no change in [AE]/[ES] ratios as determined from LC-MS were observed with water:methanol ratios of 3:1 or 7:1.*

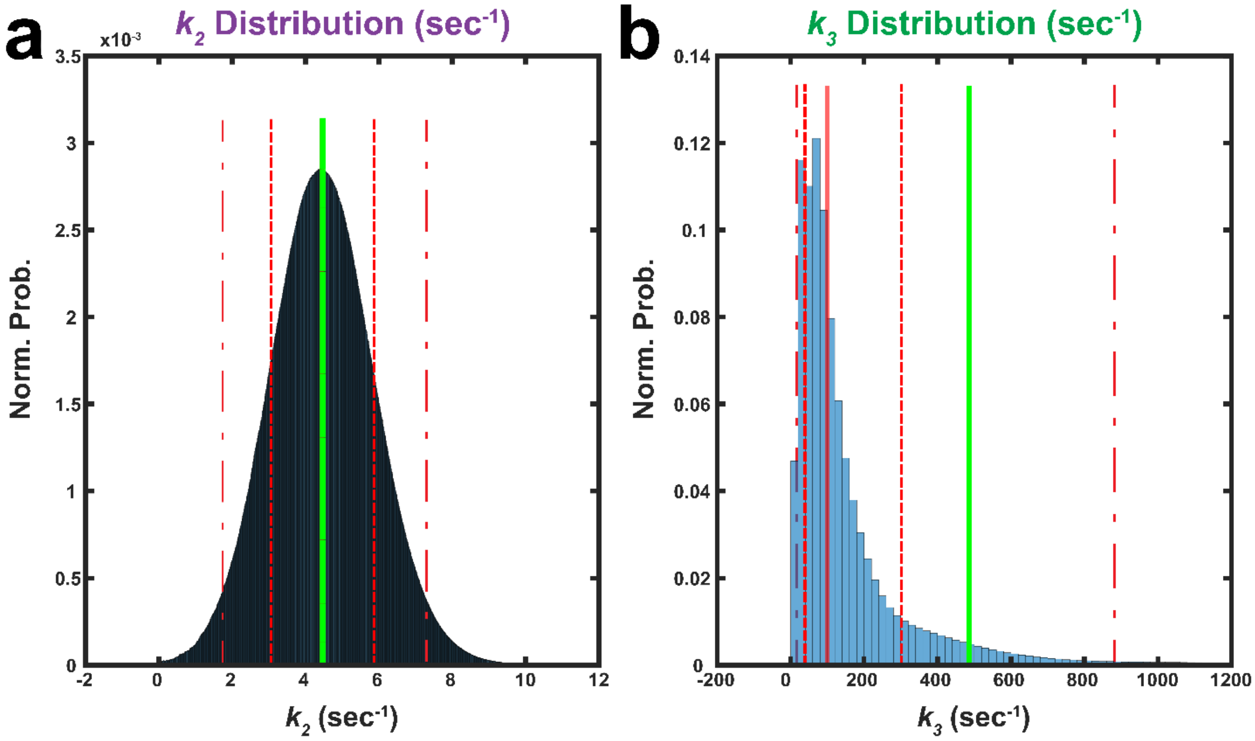

***Figure S4. Example kinetic parameter distributions determined using bootstrapping algorithm for TEM-1 β-lactamase with CTX.*** *Normalized probability distribution of* ***(a)*** *$k_2$ and* ***(b)*** *$k_3$ values from analytical bootstrapping using Michaelis-Menten parameters and measured [AE]/[ES ]ratios. While quantification of the [AE]/[ES] ratios may be normally distributed, the distributions of rate constants are not necessarily since both $k_2$ and $k_3$ > $k_{cat}$, i.e. $0 \leq [AE]/[ES] \leq$ infinity. As such, we utilize the uncertainty in both $k_{cat}$ and [AE]/[ES] ratios to bootstrap a distribution of possible $k_2$ and $k_3$ values and determine the median and appropriate confidence intervals. The solid green and red lines indicate the average and median, respectively. The 68% CI (red dashed lines; 15.865 and 84.135 quantiles) and 95% CI (red double dashed lines; 2.5 and 95 quantiles), equivalent to 1σ and 2σ for normal distributions, respectively, are shown in red dashed lines. See SI MATLAB code.*

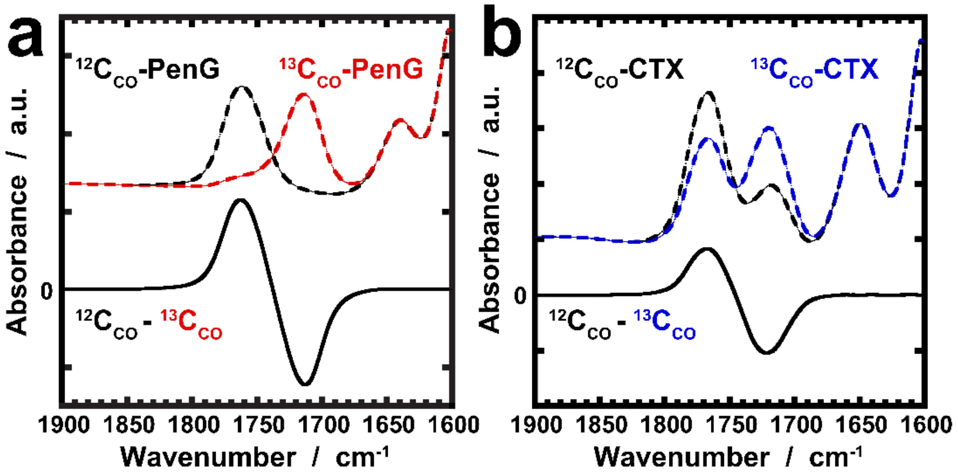

***Figure S5. FTIR spectra of $^{12}C_{CO}$ and $^{13}C_{CO}$ β-lactams in aqueous buffer.[4] (a)*** *PenG (black and red dashed lines, respectively) and **(b)** CTX (black and blue dashed lines, respectively) and corresponding isotope-edited difference spectra ($^{12}C_{CO}$ − $^{13}C_{CO}$; solid black line) in deuterated aqueous buffer (50 mM $KP_i$, 100 mM NaCl, pD 7.0), offset for clarity. Note that the $^{12}$C-component of the 50:50 $^{12/13}$C-CTX basis spectrum is first removed using a reference $^{12}$C-CTX spectrum in aqueous buffer. Figure adapted from Kozuch et al.[4] Note that the IR features for both PenG and CTX are much narrower when associated with the protein (Fig. 3 and S7) so that the positive ($^{12}$C=O) and negative ($^{13}$C=O) features are well separated.*

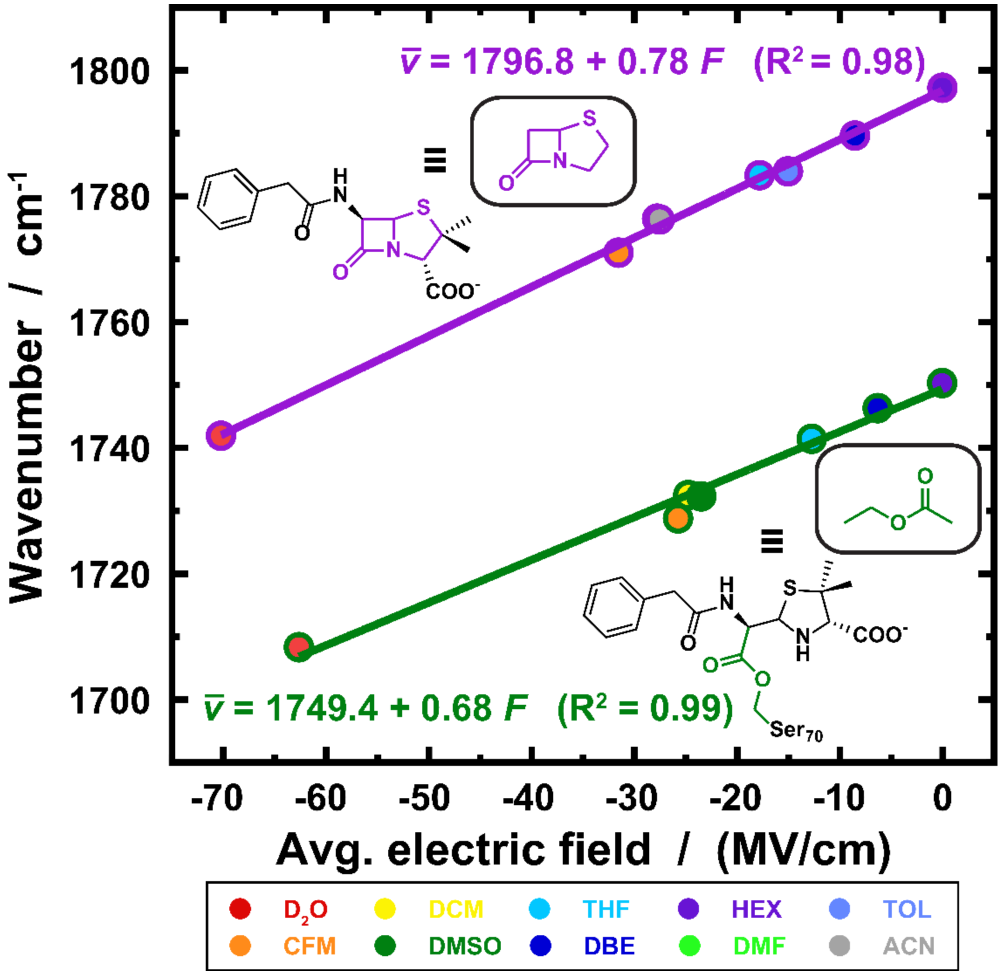

***Figure S6. Electric field-frequency calibration for model compounds and their β-lactam antibiotic mechanistic equivalents.*** [5] *Top line (purple) corresponds to the penam core (inset) and bottom line (green) to ethyl acetate, which correspond to the Michaelis complex and acyl-enzyme, respectively. Best fit lines and equations are shown in terms of the VSE based on fixed-charge MD simulations. Solvent legend: CFM = chloroform, DCM = dichloromethane, DMSO = dimethylsulfoxide, THF = tetrahydrofuran, DBE = dibutyl ether, HEX = hexanes, DMF = dimethylformamide, TOL = toluene, ACN = acetonitrile. In order to establish an analogous comparison for CTX's cephem-core (ES complex), DFT vibrational frequency calculations were used to predict the harmonic frequencies for the penam and cephem β-lactam C=O, which were scaled according to the scaling factor of 0.9679 for B3LYP/6-311(p,d) to 6-311G**(3pd,3df), resulting in corrected frequencies of 1781.1 and 1779.8 cm$^{-1}$. We utilize this frequency offset ($\Delta\bar{v} = 1.3$ cm$^{-1}$) and the Stark tuning rate for the penam-core (|Δμ| = 0.78 cm$^{-1}$/(MV/cm); purple line), based on their similar bicyclic structures, to determine the VSE field-frequency calibration of intact CTX as $\bar{v} = 0.78F + 1795.5$. Figure adapted from Kozuch et al.*[5]

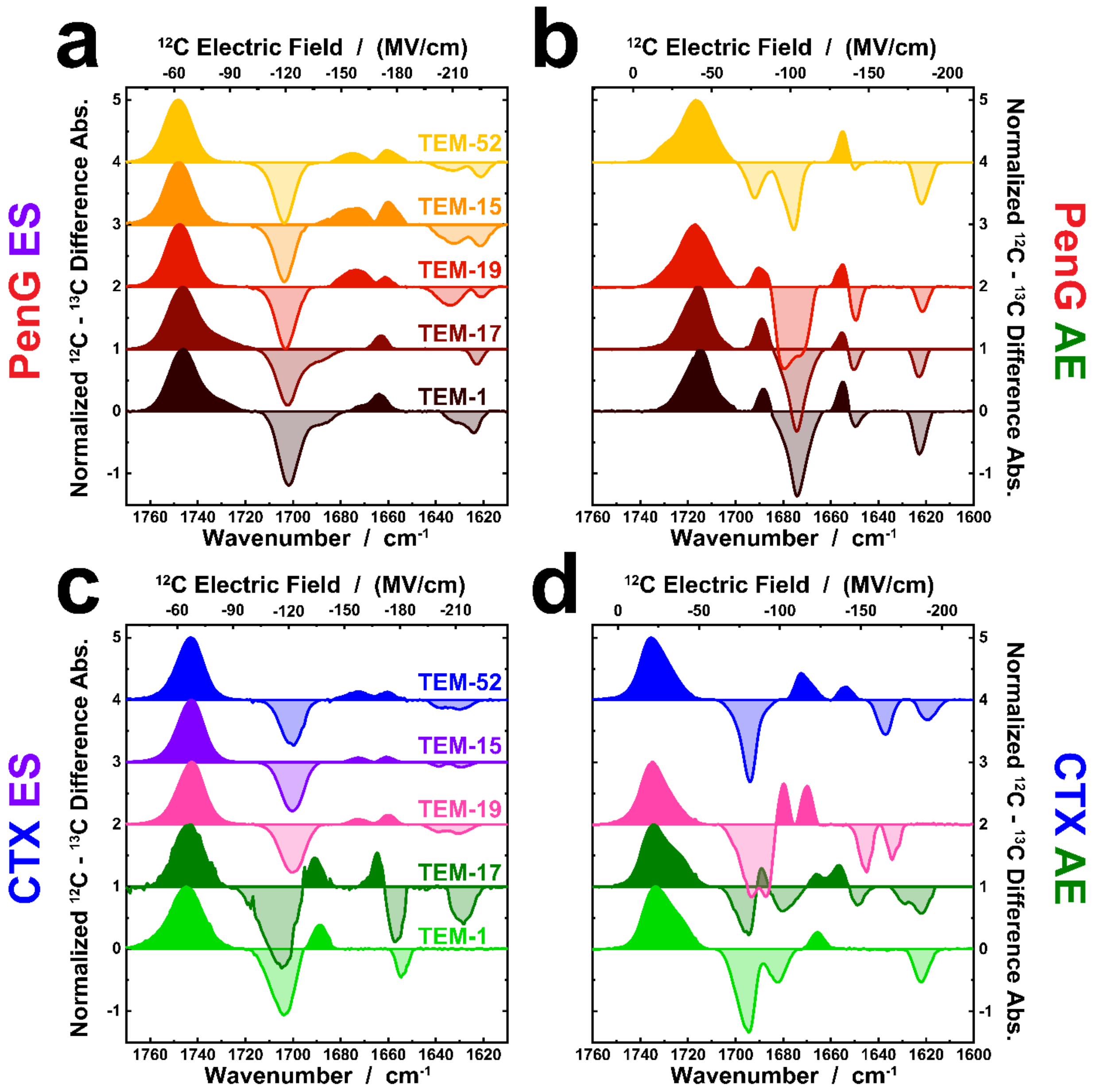

**Figure S7.** ***Full isotope-edited IR difference spectroscopy of the TEM β-lactamases with PenG and CTX. (a-d)*** *The vibrational frequencies and corresponding $^{12}$C-electric fields for* ***(a, b)*** *PenG and* ***(c, d)*** *CTX in the* ***(a,c)*** *ES (S70G) and* ***(b,d)*** *AE (E166N) complexes across evolutionary variants from TEM-1 to TEM-52. IR spectra are shown as a $^{12}$C-normalized $^{12}$C-$^{13}$C difference spectra (see Figure 3) with $^{13}$C spectral components (negative absorbance) being shown with faded color. The top axis is the electric field corresponding to the $^{12}$C-isotope of the corresponding VSE calibrated probe (Figure S6). The TEM-15 E166N mutant was not measured due to the protein's instability and likely aggregation, observed as increasing protein-specific spectral changes and loss of 280 nm absorbance over time.*

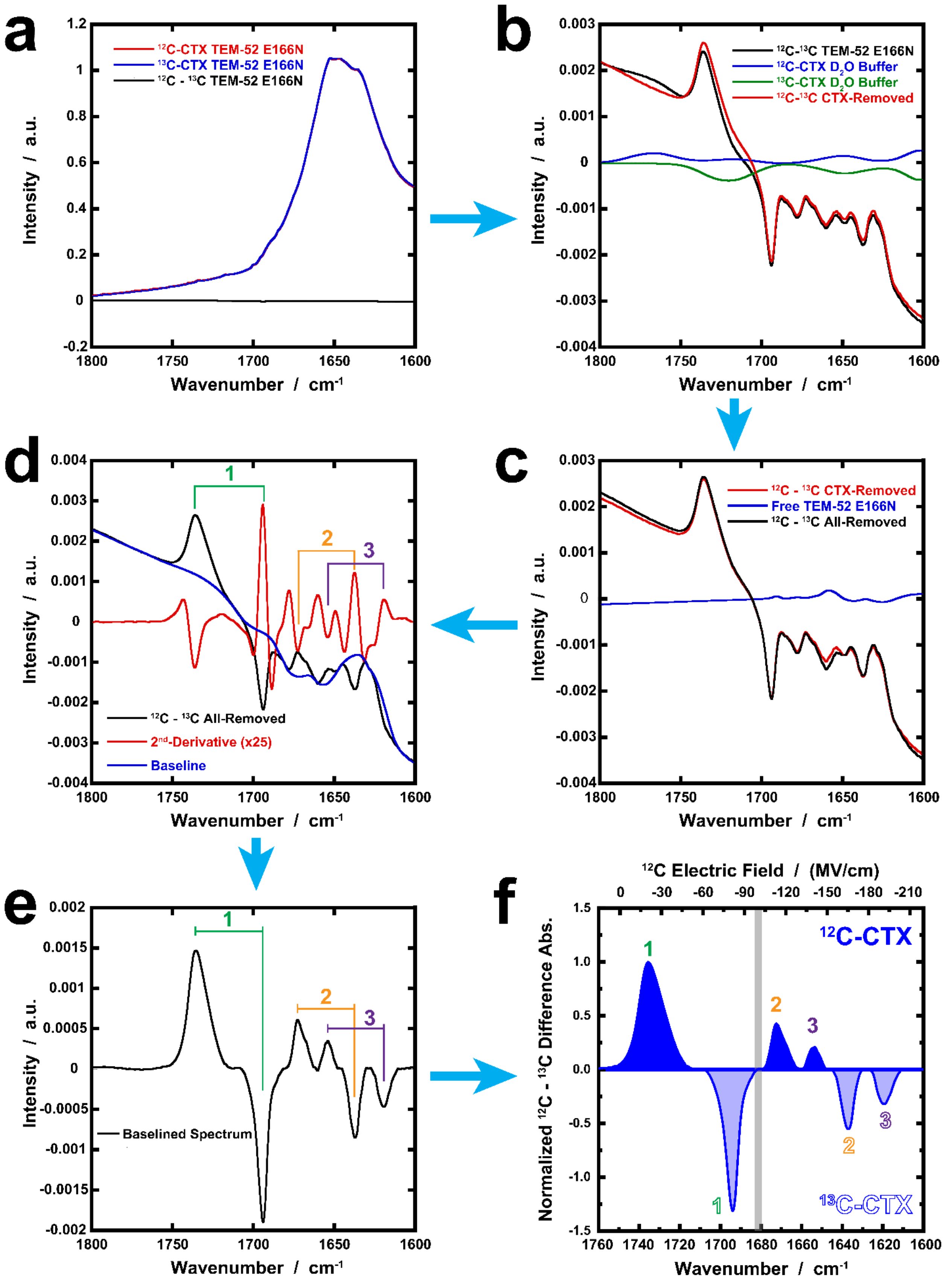

a
$^{12}$C-CTX TEM-52 E166N
$^{13}$C-CTX TEM-52 E166N
$^{12}$C - $^{13}$C TEM-52 E166N
Intensity / a.u.
Wavenumber / cm$^{-1}$
b
$^{12}$C-$^{13}$C TEM-52 E166N
$^{12}$C-CTX D$_2$O Buffer
$^{13}$C-CTX D$_2$O Buffer
$^{12}$C-$^{13}$C CTX-Removed
c
$^{12}$C - $^{13}$C CTX-Removed
Free TEM-52 E166N
$^{12}$C - $^{13}$C All-Removed
d
$^{12}$C - $^{13}$C All-Removed
2$^{nd}$-Derivative (x25)
Baseline
e
Baselined Spectrum
f
$^{12}$C Electric Field / (MV/cm)
$^{12}$C-CTX
$^{13}$C-CTX
Normalized $^{12}$C - $^{13}$C Difference Abs.

***Figure S8. Representative example of FTIR data processing and assignment with TEM-52 E166N and CTX. (a)*** *$^{12}C$- and $^{13}C$- spectra are scaled to minimize all protein absorbances and then subtracted to produce the isotope-edited difference spectrum (black).* ***(b)*** *All unbound CTX ($^{12}C$ in blue, $^{13}C$- in green) is then removed from the preliminary difference spectra using basis spectra of free CTX acquired in aqueous ($D_2O$) buffer (Fig. S5).* ***(c)*** *Additional features due to H/D exchange of the protein backbone and side-chains are also removed using basis spectra of the unbound protein acquired in identical aqueous buffer.* ***(d)*** *2$^{nd}$-derivative analysis (red) is utilized to determine the sets of positive and negative features and then the spectrum is baselined resulting in the* ***(e)*** *final spectrum.* ***(f)*** *Multiple difference spectra were conservatively baselined and averaged to generate the final spectra shown in Figure 3 and S7. There was no evidence of spectral features corresponding to the β-lactam C=O below 1600 $cm^{-1}$. In the TEM-# S70G mutants with PenG, as previously observed,[6] these mutants exhibit a residual slow level of hydrolytic activity leading to time-dependent changes on the timescale of the FTIR measurements. Measurements were acquired until nearly all PenG was hydrolyzed and difference spectra were taken between an early and late spectrum (timescale of ~ 1-3 hrs apart) of identical isotopes, i.e. $^{12}C_{early}$ – $^{12}C_{late}$ and $^{13}C_{early}$ – $^{13}C_{late}$. An isotope-edited difference spectrum was then achieved through scaled subtraction, i.e. ($^{12}C_{early}$ – $^{12}C_{late}$) – ($^{13}C_{early}$ – $^{13}C_{late}$), which was further processed according to the protocol described above. Hydrolysis leads to formation of the $^{12}C$ and $^{13}C$-penicilloic acid product species (1599.8 and 1554.5 $cm^{-1}$), which have opposite signs relative to the peaks assigned to the Michaelis complex due to the inverse correlation between substrate and product concentrations as a function of time. An increasing amount of bound product is observed over time with sharp peaks at ca. 1582 and 1545 $cm^{-1}$ across all the TEM variants, corresponding to the $^{12}C$- and $^{13}C$-species, respectively, and assigned to the asymmetric carboxylate stretching frequency.*

**Table S3. TEM-# S70G PenG vibrational frequencies and lineshape parameters from curve-fitting.**

| TEM-# S70G **PenG** | Frequency ($cm^{-1}$)[a,b] | FWHM ($cm^{-1}$)[a,b] | Integral ($OD/cm^{-1}$) | Lineshape (%L+G)[c] |
|---|---|---|---|---|
| 1 | 1746.4 (1701.8) | 13.4 (9.6) | 0.0142 (-0.0115) | 5 (0) |
| | 1732.2 (1689.3) | 13.3 (14.3) | 0.0028 (-0.0052) | 20 (100) |
| | 1671.2 (1631.0) | 14.2 (8.1) | 0.0024 (-0.0016) | 99 (0) |
| | 1663.5 (1624.0) | 6.8 (5.7) | 0.0017 (-0.0019) | 0 (0) |
| | 1582.8 (1546.1) | 9.2 (9.9) | -0.0080 (0.0053) | 0 (0) |
| 17 | 1746.6 (1702.5) | 14.1 (10.1) | 0.0189 (-0.0142) | 4 (28) |
| | 1731.7 (1690.3) | 15.9 (12.2) | 0.0057 (-0.0030) | 86 (0) |
| | 1663.2 (1622.8) | 5.9 (5.2) | 0.0019 (-0.0017) | 8 (0) |
| | 1585.0 (1546.4) | 8.7 (10.1) | -0.0075 (0.0072) | 0 (3) |
| 19 | 1747.7 (1703.1) | 13.3 (9.6) | 0.0482 (-0.0345) | 10 (12) |
| | 1673.9 (1634.2) | 12.5 (10.9) | 0.0121 (-0.0113) | 0 (0) |
| | 1660.7 (1620.5) | 5.0 (5.2) | 0.0024 (-0.0029) | 0 (0) |
| | 1582.3 (1544.5) | 8.5 (11.4) | -0.0182 (0.0240) | 0 (3) |
| 15 | 1748.0 (1703.8) | 13.7 (9.1) | 0.0495 (-0.0319) | 17 (26) |
| | 1675.5 (1633.0) | 14.6 (12.5) | 0.0177 (-0.0125) | 64 (0) |
| | 1659.3 (1620.8) | 6.7 (7.6) | 0.0073 (-0.0085) | 0 (12) |
| | 1582.8 (1545.7) | 9.0 (12.5) | -0.0207 (0.0264) | 0 (2) |
| 52 | 1748.3 (1703.8) | 13.7 (9.6) | 0.0796 (-0.0566) | 10 (20) |
| | 1675.4 (1633.5) | 10.3 (11.3) | 0.0101 (-0.0082) | 30 (0) |
| | 1659.6 (1621.1) | 7.7 (6.2) | 0.0081 (-0.0079) | 0 (0) |

| | | | | |
|---|---|---|---|---|
| | 1582.6 (1544.8) | 9.7 (13.0) | -0.0304 (0.0439) | 0 (4) |

[a] *Uncertainties in peak positions are ca. ± 1 $cm^{-1}$ based on comparison of multiple spectral comparisons within a given protein-ligand complex. There are greater uncertainties in the FWHM, integrations, and lineshapes due to curve-fitting of multiple overlapping peaks in many protein spectra.*

[b] *Values in parentheses correspond to those of the $^{13}C$-substrate peaks. Note that the $^{13}C$ peaks are generally narrower than the corresponding $^{12}C$ peak and that as the frequencies shift to lower frequencies, the isotope frequency offset ($^{12}C$-$^{13}C$) gets smaller. These changes may arise due to differential effects of normal mode coupling, slight variations in the Stark tuning rate, and homogeneous broadening, respectively.*

[c] *The pseudo-Voigt lineshape was used to fit all peaks, where the % corresponds to the linear combination of Lorentzian (L) and Gaussian (G) lineshapes.*

**Table S4. TEM-# S70G CTX vibrational frequencies and lineshape parameters from curve-fitting.**

| TEM-# S70G **CTX** | Frequency ($cm^{-1}$)[a,b] | FWHM ($cm^{-1}$)[a,b] | Integral (OD/$cm^{-1}$) | Lineshape (%L+G)[c] |
|---|---|---|---|---|
| 1 | 1744.9 (1704.2) | 16.3 (11.9) | 0.0035 (-0.0028) | 32 (33) |
| | 1690.1 (1654.3) | 10.0 (5.0) | 0.0007 (-0.0004) | 0 (0) |
| 17 | 1743.7 (1703.6) | 14.8 (18.8) | 0.0040 (-0.0072) | 0 (12) |
| | 1693.4 (1657.0) | 11.2 (6.5) | 0.0028 (-0.0017) | 0 (0) |
| | 1664.4 (1628.7) | 5.0 (8.9) | 0.0010 (-0.0014) | 63 (0) |
| 19 | 1751.2 (1704.9) | 11.6 (8.4) | 0.0039 (-0.0079) | 0 (43) |
| | 1742.4 (1699.2) | 12.4 (9.8) | 0.0448 (-0.0234) | 18 (0) |
| | 1672.5 (1639.4) | 7.4 (6.9) | 0.0023 (-0.0026) | 9 (0) |
| | 1660.0 (1630.1) | 6.1 (9.1) | 0.0032 (-0.0049) | 0 (0) |
| 15 | 1749.5 (1704.9) | 13.9 (7.7) | 0.0061 (-0.0071) | 0 (38) |
| | 1742.5 (1699.5) | 12.1 (9.4) | 0.0451 (-0.0255) | 13 (0) |
| | 1672.7 (1639.0) | 6.8 (5.1) | 0.0019 (-0.0013) | 8 (0) |
| | 1660.5 (1629.2) | 5.6 (7.4) | 0.0018 (-0.0022) | 0 (0) |
| 52 | 1750.1 (1704.2) | 14.2 (7.6) | 0.0076 (-0.0153) | 4 (58) |
| | 1742.7 (1699.0) | 12.2 (8.0) | 0.0552 (-0.0210) | 17 (0) |
| | 1673.0 (1638.6) | 11.0 (6.5) | 0.0071 (-0.0032) | 21 (0) |
| | 1660.3 (1629.9) | 7.0 (9.6) | 0.0038 (-0.0065) | 5 (0) |

[a] *Uncertainties in peak positions are ca. ± 1 $cm^{-1}$ based on comparison of multiple spectral comparisons within a given protein-ligand complex. There are greater uncertainties in the FWHM, integrations, and lineshapes due to curve-fitting of multiple overlapping peaks in many protein spectra.*

[b] *Values in parentheses correspond to those of the $^{13}$C-substrate peaks. Note that the $^{13}$C peaks are generally narrower than the corresponding $^{12}$C peak and that as the frequencies shift to lower frequencies, the isotope frequency offset ($^{12}$C-$^{13}$C) gets smaller. These changes may arise due to differential effects of normal mode coupling, slight variations in the Stark tuning rate, and homogeneous broadening, respectively.*

[c] *The pseudo-Voigt lineshape was used to fit all peaks, where the % corresponds to the linear combination of Lorentzian (L) and Gaussian (G) lineshapes.*

**Table S5. TEM-# E166N PenG vibrational frequencies and lineshape parameters from curve-fitting.**

| TEM-# E166N **PenG** | Frequency ($cm^{-1}$)[a,b] | FWHM ($cm^{-1}$)[a,b] | Integral ($OD/cm^{-1}$) | Lineshape (%L+G)[c] |
|---|---|---|---|---|
| 1 | 1722.3 (***)[d] | 9.1 | 0.0046 | 0 |
| | 1714.6 (1673.9) | 11.2 (9.0) | 0.0320 (-0.0384) | 49 (66) |
| | 1688.2 (1650.6) | 5.0 (6.6) | 0.0051 (-0.0063) | 0 (0) |
| | 1654.5 (1622.6) | 5.5 (5.4) | 0.0130 (-0.0089) | 100 (0) |
| 17 | 1722.5 (***)[d] | 10.0 | 0.0119 | 23 |
| | 1715.1 (1674.2) | 10.7 (8.7) | 0.0443 (-0.0566) | 31 (60) |
| | 1688.9 (1650.6) | 6.1 (5.2) | 0.0126 (-0.0102) | 0 (18) |
| | 1654.4 (1622.9) | 6.6 (4.7) | 0.0143 (-0.0084) | 100 (0) |
| 19 | 1729.2 (***)[b, d] | 10.7 | 0.0060 | 0 |
| | 1717.1 (1680.4)[b] | 14.0 (9.9) | 0.0469 (-0.0588) | 0 (4) |
| | 1708.1 (1671.5)[b] | 8.2 (6.4) | 0.0038 (-0.0195) | 0 (0) |
| | 1685.2 (1649.7) | 10.1 (4.5) | 0.0277 (-0.0090) | 1 (0) |
| | 1655.1 (1621.5) | 5.0 (5.0) | 0.0075 (-0.0068) | 35 (0) |
| 52 | 1730.6 (1691.9) | 7.4 (7.4) | 0.0026 (-0.0161) | 0 (100) |
| | 1716.3 (1676.3) | 16.1 (7.5) | 0.0437 (-0.0190) | 19 (0) |
| | *** (1650.5)[a, d] | (4.5) | (-0.0018) | (0) |
| | 1655.0 (1621.6) | 5.0 (6.4) | 0.0071 (-0.0108) | 26 (0) |

[a] *Uncertainties in peak positions are ca. ± 1 $cm^{-1}$ based on comparison of multiple spectral comparisons within a given protein-ligand complex. There are greater uncertainties in the FWHM, integrations, and lineshapes due to curve-fitting of multiple overlapping peaks in many protein spectra.*

[b] *Values in parentheses correspond to those of the $^{13}C$-substrate peaks. Note that the $^{13}C$ peaks are generally narrower than the corresponding $^{12}C$ peak and that as the frequencies shift to lower frequencies, the isotope frequency offset ($^{12}C$-$^{13}C$) gets smaller. These changes may arise due to differential effects of normal mode coupling, slight variations in the Stark tuning rate, and homogeneous broadening, respectively.*

[c] *The pseudo-Voigt lineshape was used to fit all peaks, where the % corresponds to the linear combination of Lorentzian (L) and Gaussian (G) lineshapes.*

[d] *"***" corresponds to either a $^{12}C$ or $^{13}C$ feature that cannot be resolved due to spectral overlap and cancellation.*

**Table S6. TEM# E166N CTX vibrational frequencies and lineshape parameters from curve-fitting.**

| TEM-# E166N **CTX** | Frequency ($cm^{-1}$)[a,b] | FWHM ($cm^{-1}$)[a,b] | Integral ($OD/cm^{-1}$) | Lineshape (%L+G)[c] |
|---|---|---|---|---|
| 1 | 1734.3 (1695.6) | 11.2 (9.0) | 0.0139 (-0.0168) | 6 (18) |
| | 1724.4 (1682.1) | 12.0 (7.0) | 0.0074 (-0.0048) | 0 (0) |
| | 1665.6 (1621.9) | 6.2 (6.8) | 0.0024 (-0.0047) | 12 (0) |
| 17[d] | 1735.6 (1694.7) | | | |
| | 1722.5 (1681.3) | | | |
| | 1689.0 (1649.1) | | | |
| | 1667.0 (1630.3) | | | |
| | 1655.8 (1621.0) | | | |
| 19 | 1735.1 (1693.0) | 11.8 (11.1) | 0.0188 (-0.0232) | 16 (32) |
| | 1725.5 (1686.5) | 10.8 (4.2) | 0.0056 (-0.0045) | 77 (0) |
| | 1680.0 (1645.5) | 4.7 (5.6) | 0.0054 (-0.0067) | 0 (0) |
| | 1670.0 (1633.9) | 5.0 (5.2) | 0.0051 (-0.0044 | 0 (0) |
| 52 | 1735.9 (1694.3) | 10.6 (7.2) | 0.0168 (-0.0195) | 29 (66) |
| | 1727.8 (***)[e] | 11.4 | 0.0080 | 41 |
| | 1672.7/1668.2 (1637.3) | 5.7/5.7 (6.6) | 0.0053/0.0014 (-0.0058) | 86/0 (0) |
| | 1653.8 (1619.2) | 5.9 (7.5) | 0.0019 (-0.0039) | 0 (0) |

[a] *Uncertainties in peak positions are ca. ± 1 $cm^{-1}$ based on comparison of multiple spectral comparisons within a given protein-ligand complex. There are greater uncertainties in the FWHM, integrations, and lineshapes due to curve-fitting of multiple overlapping peaks in many protein spectra.*

[b] *Values in parentheses correspond to those of the $^{13}C$-substrate peaks. Note that the $^{13}C$ peaks are generally narrower than the corresponding $^{12}C$ peak and that as the frequencies shift to lower frequencies, the isotope frequency offset ($^{12}C$-$^{13}C$) gets smaller. These changes may arise due to differential effects of normal mode coupling, slight variations in the Stark tuning rate, and homogeneous broadening, respectively.*

[c] *The pseudo-Voigt lineshape was used to fit all peaks, where the % corresponds to the linear combination of Lorentzian (L) and Gaussian (G) lineshapes.*

[d] *Peak positions determined based on 2nd-derivative maxima and minima positions. Uncertainties will be increased for peaks that overlap due to cancellation.*

[e] *"***" corresponds to either a $^{12}C$ or $^{13}C$ feature that cannot be resolved due to spectral overlap and cancellation.*

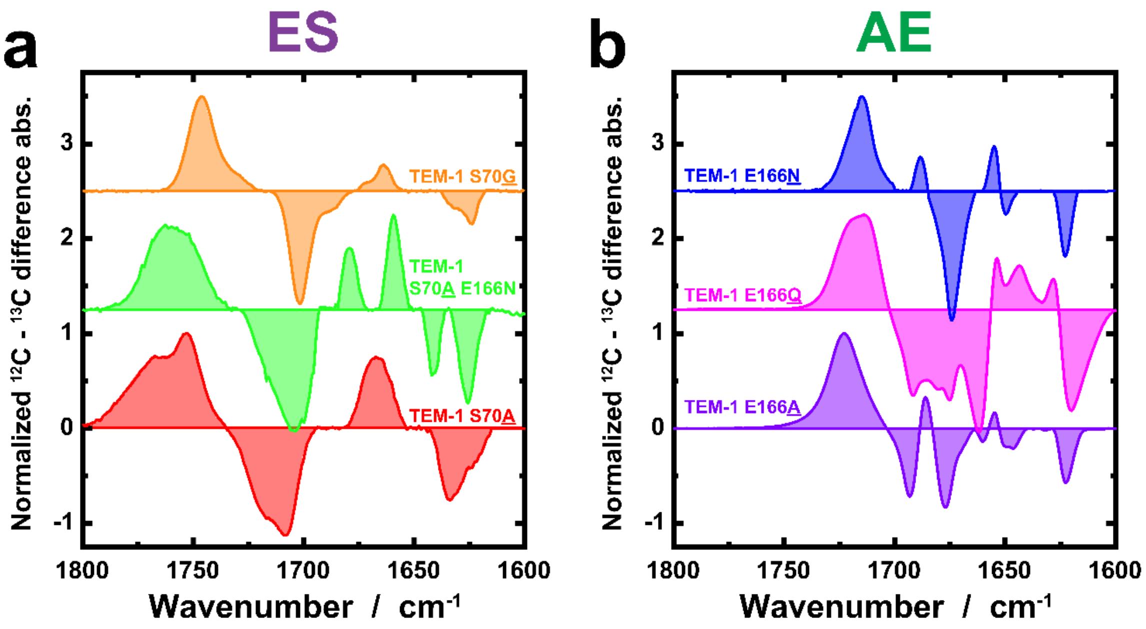

***Figure S9. Mutational effect on IR spectra of TEM-1 β-lactamase with PenG.*** *The mutational choice for the S70X and E166X positions as related to the WT TEM for the* ***(a)*** *ES and* ***(b)*** *AE complex with PenG. Positive peaks correspond to $^{12}$C-PenG with corresponding negative peaks for the $^{13}$C-PenG with an offset of ca. 35-45 cm$^{-1}$. Spectra are normalized and offset for clarity. The highest frequency peaks (1800-1700 cm$^{-1}$) are generally more mutation-dependent than the lower frequency peak(s) (<1675 cm$^{-1}$), with the simplest spectra corresponding to those of the S70G and E166N, as supported by crystallographic results.[7] Additional variants S70(C/T) and E166D were all considered but either did not express or had rates too fast for FTIR measurements.[8]*

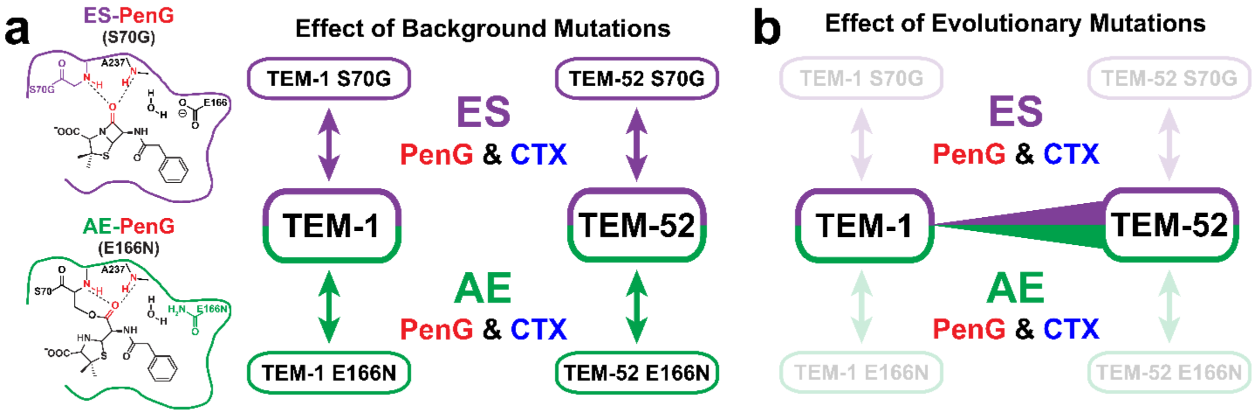

***Figure S10. MD Methodology for Structure-Guided Interpretation of Kinetic and VSE Results.*** *The MD methodology to assess the structural effect of* ***(a)*** *introduction of background mutations S70G and E166N, relative to the WT TEM-1 and TEM-52, on conformational sampling and* ***(b)*** *evolutionary mutations to alter the active site environment in the ES (purple) and AE (green) complexes with PenG (red) and CTX (blue).*

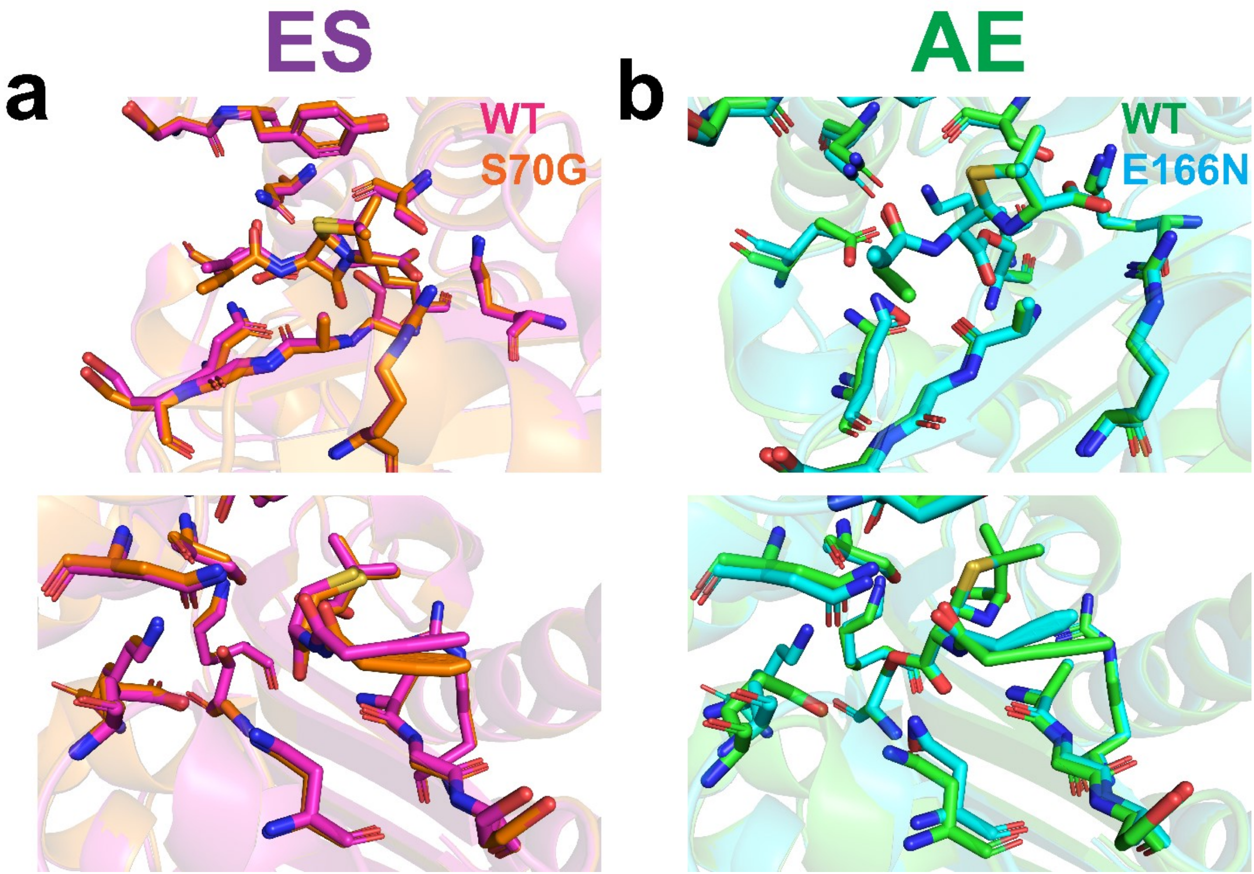

***Figure S11. Structural comparison of tight-binding PenG complexes with S70G and E166N background mutants and WT TEM β-lactamases from MD simulations.*** *Representative structural alignment examples of WT TEM-1 (WT; pink, green) relative to the TEM-1 S70G (orange) and E166N (cyan) shortest oxyanion hole complexes for the **(a)** ES and **(b)** AE states with PenG. Global alignment between variants (WT vs. S70G and WT vs. E166N) exhibit a RMSD of 0.3 Å for the average structures of those observed with the shortest distances to the S70 andA237 backbone amide N-H.*

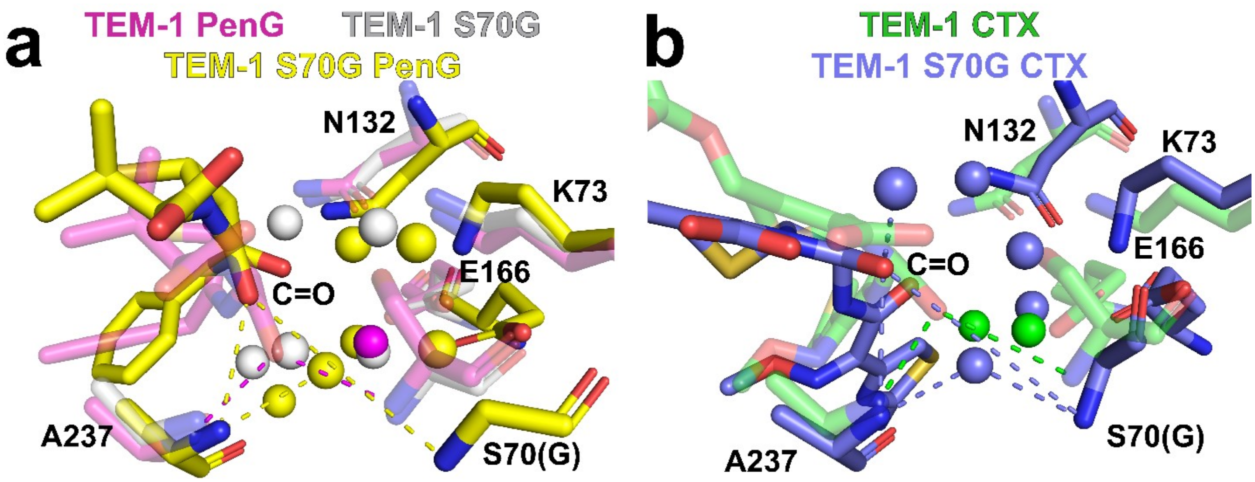

***Figure S12. Structural comparison of weak-binding PenG complexes with S70G background mutations and WT TEM β-lactamases from MD simulations.*** *Representative structural alignment examples of the strong- (WT; pink, green) and weak-binding TEM-1 (S70G; yellow, purple) complexes with* ***(a)*** *PenG (pink, yellow) and* ***(b)*** *CTX (green, purple) relative to the apo crystal structure of TEM-1 S70G (white; PDBID: 1ZG6 ). Alignments are made on the entire protein structure.*

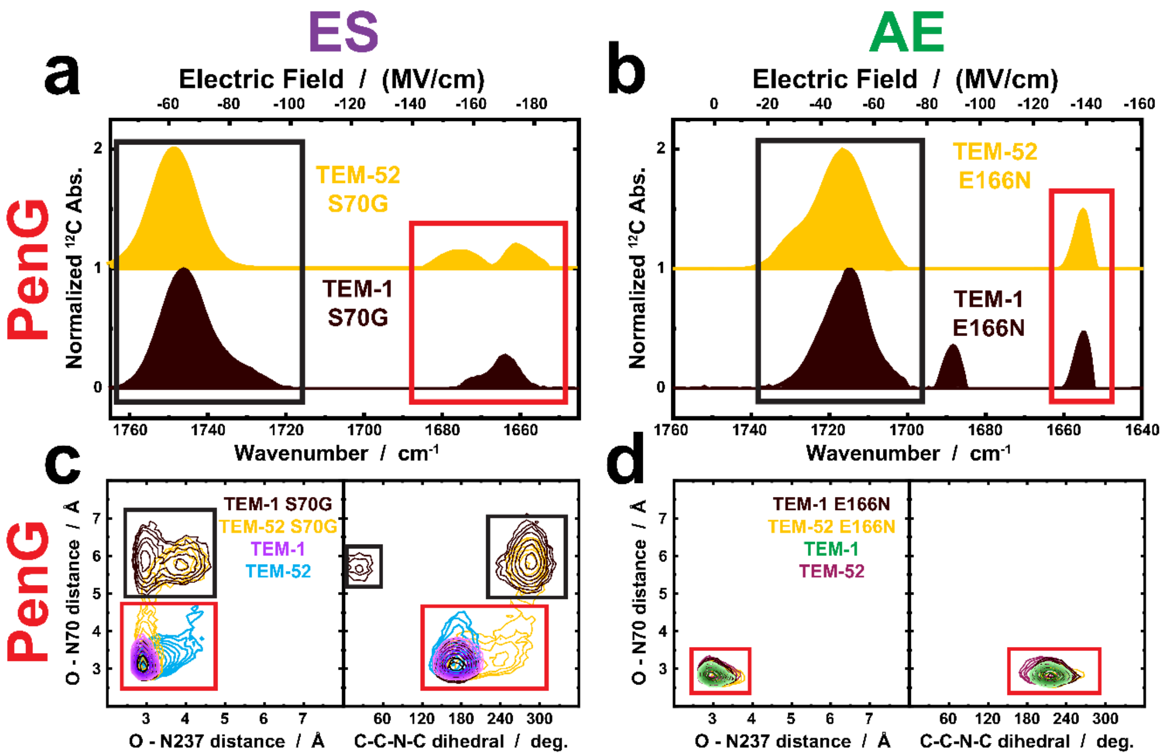

***Figure S13. Structure-guided approach to inferring catalytically relevant electric fields.*** *Representative examples of TEM-1 and TEM-52 with PenG for comparison between the IR spectra of* ***(a)*** *S70G and* ***(b)*** *E166N background mutants relative to MD simulations of the* ***(c)*** *ES and* ***(d)*** *AE complexes.* $^{13}$*C PenG (negative intensity) signal removed for clarity. MD simulations are depicted with either the oxyanion hole C=O – N distances (see main text) or dihedral angle (C-C-N-C) of the bound PenG substrate. Background mutants are shown consistently for TEM-1 (brown) and TEM-52 (yellow), relative to their WT species in the ES (pink, cyan) and AE (green, maroon) complexes. Black and red boxes indicate presumed assignment of populations between the VSE results and MD simulations.*

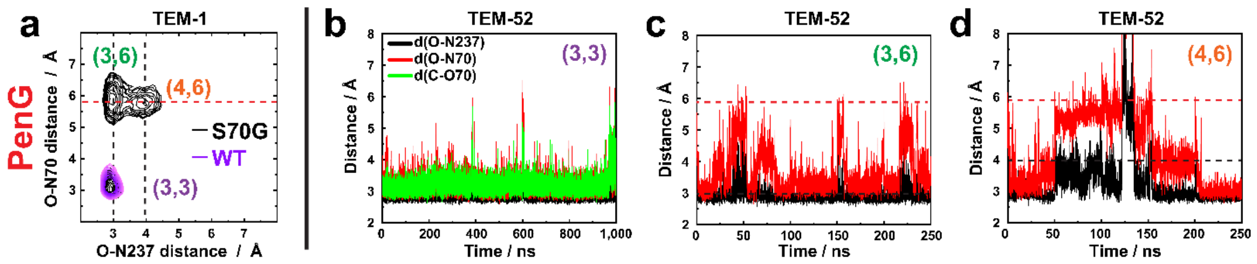

***Figure S14. Biased MD simulations of WT TEM β-lactamases as informed from kinetically compromised variants.*** *A representative* ***(a)*** *overlaid 2D correlation plot of the ES complex of PenG with WT TEM-1 (purple) and S70G TEM-1 to identify additional conformations (cross peaks of black and red dotted lines) relative to those in the shortest oxyanion hole geometry* ***(b)*** *(3, 3). Using these conformations, five 50 ns MD simulations of the WT TEM-52 complex were biased to start from the* ***(c)*** *(3,6) and* ***(d)*** *(4,6) conformations (starting distances indicated with dotted lines), which converge back towards the (3,3) conformation.*

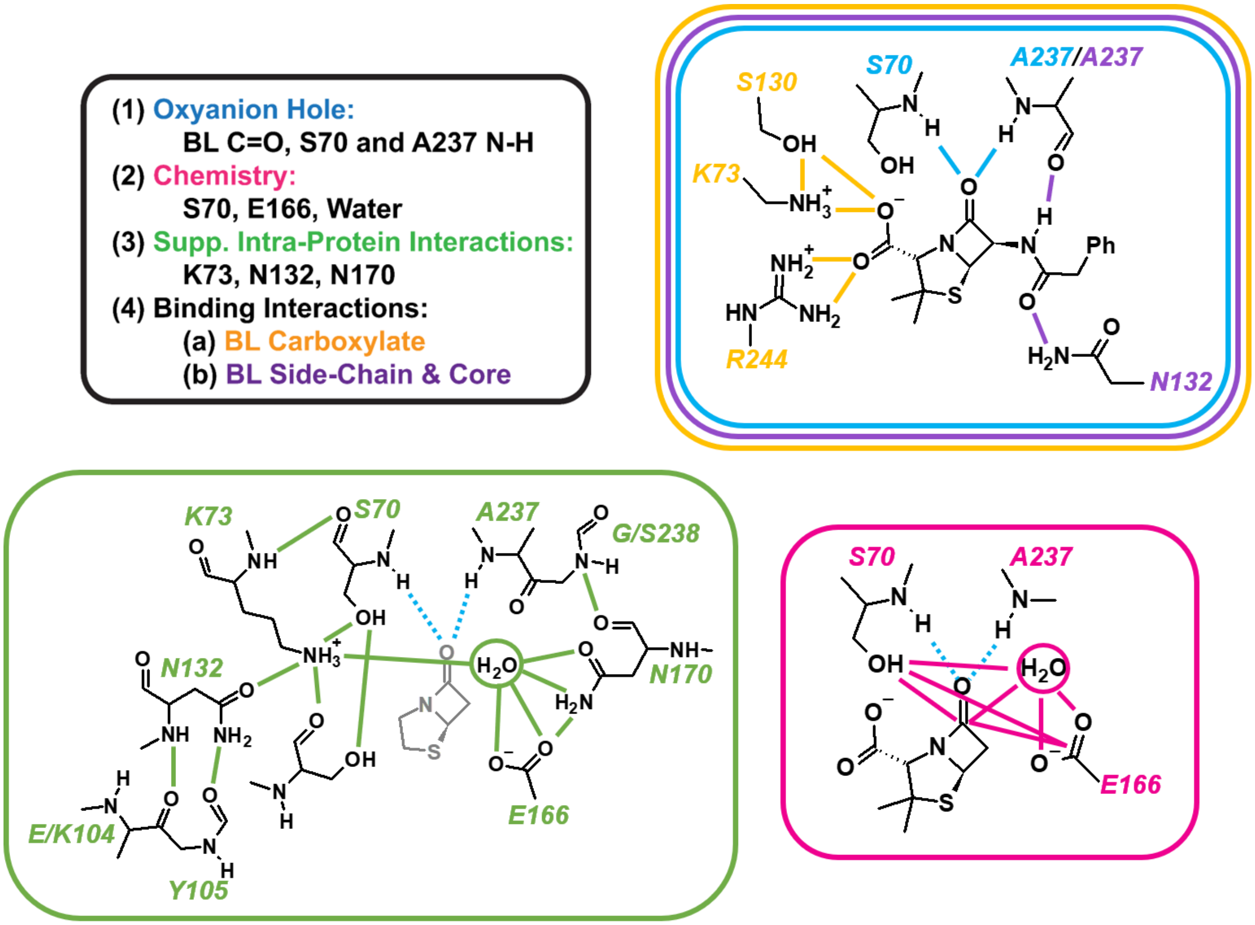

***Scheme S2. Atoms and Interactions Considered in the MD Analysis.*** *Atoms and distances quantified in terms of RMSF and distance changes are indicated by color-coded solid lines and bolded atoms. Colors correspond to the categorization scheme shown in Figure S15b,c, Tables S7-12, and Figure S17. The same general scheme is used for the ES and AE complexes of both PenG and CTX. RMSF and distance values were extracted from representative trajectories and average structures (n = 3-5, each 50-200 ns), respectively, from the WT MD simulations of TEM-1 and TEM-52 with PenG and CTX in the ES and AE states. Average values and standard deviations for these complexes are presented below in Tables S7-12; these were used to produce Figure 5-6 and Figure S17. Statistically significant ($p<0.05$) were determined from a two-tailed student's t-test (see t-values for given degrees of freedom (df)). Significant values were then utilized in the corresponding averages for each set of discretized regions or interactions in the active site based on proximity or known involvement in catalysis: K73/N132/E166/N170, oxyanion hole, β-lactam carboxylate, or β-lactam (Figure 5, 6, S12). Interactions referred to in the H-bond network of K73/N132/E166/N170 correspond to those side chains listed and those within H-bonding distance (≤3.5 Å) such as K73 to S70, the backbone of*

*N170 to the backbone of E240, among others. Oxyanion hole interactions are those that connect to S70 and the β-lactam C=O primarily via backbone interactions with S70 and A237. The carboxylate interactions refer to those made to the carboxylate functional group of PenG or CTX primarily from S130, K234 and R244, in addition to the atoms of the substrate itself (COO). Interactions to the β-lactam substrate (excluding β-lactam C=O and carboxylate) are those that link to the side chain amide (i.e. backbone amide of A237, N132), as well as the penam or cephem core (either bicyclic or acylated rings), or the respective side chains of PenG (phenylacetamido) and CTX (oxyimino/aminothiazolyl-ring).*

**Table S7. PenG-dependent average changes in active site RMSFs due to evolution.**

| PenG RMSF (Å) | ES | | | | AE | | | |
|---|---|---|---|---|---|---|---|---|
| | TEM-1 PenG | | TEM-52 PenG | | TEM-1 PenG | | TEM-52 PenG | |
| Atom [a,b] | Avg. | Std. Dev. | Avg. | Std. Dev. | Avg. | Std. Dev. | Avg. | Std. Dev. |
| S70bbN | **0.312** | **0.005** | **0.405** | **0.023** | **0.330** | **0.019** | **0.461** | **0.028** |
| S70bbH | **0.365** | **0.008** | **0.454** | **0.030** | **0.378** | **0.016** | **0.502** | **0.028** |
| S70O | **0.341** | **0.009** | **0.486** | **0.049** | **0.347** | **0.028** | **0.553** | **0.063** |
| S70bbO | **0.359** | **0.008** | **0.476** | **0.038** | **0.396** | **0.024** | **0.563** | **0.036** |
| K73bbN | **0.330** | **0.017** | **0.379** | **0.015** | **0.384** | **0.048** | **0.497** | **0.057** |
| K73bbH | **0.372** | **0.020** | **0.418** | **0.015** | **0.426** | **0.051** | **0.519** | **0.022** |
| K73N | **0.355** | **0.005** | **0.438** | **0.031** | **0.583** | **0.063** | **0.878** | **0.117** |
| X104bbO | **0.537** | **0.023** | **0.659** | **0.048** | **0.537** | **0.055** | **1.434** | **0.234** |
| Y105bbO | **0.460** | **0.011** | **0.555** | **0.048** | **0.491** | **0.043** | **1.814** | **0.519** |
| S130O | 0.453 | 0.015 | 0.573 | 0.117 | **0.494** | **0.100** | **1.078** | **0.131** |
| S130bbO | **0.400** | **0.009** | **0.488** | **0.052** | **0.457** | **0.049** | **0.970** | **0.149** |
| N132bbN | **0.322** | **0.011** | **0.407** | **0.039** | **0.334** | **0.037** | **0.667** | **0.089** |
| N132bbH | **0.392** | **0.011** | **0.465** | **0.031** | **0.408** | **0.031** | **0.713** | **0.055** |
| N132O | **0.438** | **0.014** | **0.572** | **0.051** | **0.449** | **0.040** | **1.446** | **0.290** |
| N132N | **0.553** | **0.022** | **0.679** | **0.052** | **0.473** | **0.056** | **1.278** | **0.164** |
| E166C | **0.442** | **0.029** | **0.687** | **0.127** | **0.477** | **0.028** | **0.803** | **0.135** |
| E166O1 | **0.561** | **0.090** | **0.985** | **0.239** | **0.968** | **0.250** | **1.321** | **0.159** |
| E166O2 | **0.587** | **0.101** | **0.991** | **0.232** | **0.969** | **0.261** | **1.358** | **0.122** |
| N170O | **0.517** | **0.040** | **1.019** | **0.379** | **0.662** | **0.231** | **1.296** | **0.533** |
| N170N | 0.535 | 0.031 | 0.848 | 0.187 | **0.688** | **0.247** | **1.203** | **0.360** |
| N170bbO | **0.510** | **0.027** | **1.082** | **0.445** | **0.580** | **0.074** | **1.049** | **0.355** |
| K234N | 0.688 | 0.137 | 0.840 | 0.171 | 0.793 | 0.161 | 1.037 | 0.689 |
| A237bbN | **0.325** | **0.011** | **0.379** | **0.029** | **0.333** | **0.023** | **0.509** | **0.054** |
| A237bbH | **0.373** | **0.009** | **0.432** | **0.029** | **0.394** | **0.023** | **0.576** | **0.063** |
| A237bbO | **0.391** | **0.010** | **0.535** | **0.042** | **0.403** | **0.025** | **0.634** | **0.108** |
| X238bbN | **0.362** | **0.009** | **0.442** | **0.028** | **0.382** | **0.027** | **0.588** | **0.087** |
| X238bbH | **0.418** | **0.009** | **0.473** | **0.035** | **0.432** | **0.023** | **0.592** | **0.064** |
| X238bbC | **0.409** | **0.010** | **0.479** | **0.026** | **0.420** | **0.028** | **0.645** | **0.135** |
| E240bbN | **0.416** | **0.013** | **0.521** | **0.043** | **0.428** | **0.034** | **0.718** | **0.190** |
| E240bbH | **0.457** | **0.011** | **0.571** | **0.062** | **0.470** | **0.028** | **0.776** | **0.217** |
| R244N1 | 0.411 | 0.005 | 0.436 | 0.032 | **0.428** | **0.038** | **1.316** | **0.639** |
| R244N2 | 0.461 | 0.008 | 0.483 | 0.034 | **0.481** | **0.052** | **1.281** | **0.524** |
| Water | **0.596** | **0.081** | **1.182** | **0.326** | **0.884** | **0.476** | **19.986** | **5.504** |
| BL_C | 0.318 | 0.114 | 0.349 | 0.077 | **0.330** | **0.030** | **0.530** | **0.065** |
| BL_O | 0.372 | 0.201 | 0.378 | 0.145 | **0.346** | **0.028** | **0.515** | **0.052** |
| PNM_C | 0.351 | 0.030 | 0.352 | 0.045 | **0.452** | **0.064** | **1.153** | **0.317** |
| PNM_C1 | 0.362 | 0.042 | 0.399 | 0.030 | **0.419** | **0.102** | **1.201** | **0.436** |
| PNM_C2 | 0.340 | 0.054 | 0.397 | 0.040 | **0.353** | **0.035** | **0.687** | **0.155** |

| | | | | | | | | |
|---|---|---|---|---|---|---|---|---|
| PNM_S | 0.427 | 0.109 | 0.416 | 0.066 | **0.492** | **0.046** | **1.155** | **0.334** |
| PNM_N | 0.323 | 0.051 | 0.363 | 0.035 | **0.390** | **0.066** | **1.040** | **0.356** |
| PNM_C3 | 0.320 | 0.085 | 0.394 | 0.063 | **0.348** | **0.038** | **0.612** | **0.108** |
| Amide_N | 0.409 | 0.166 | 0.423 | 0.067 | **0.384** | **0.044** | **0.647** | **0.104** |
| Amide_H | 0.502 | 0.240 | 0.461 | 0.107 | **0.434** | **0.040** | **0.646** | **0.092** |
| Amide_C | 0.448 | 0.184 | 0.463 | 0.103 | **0.448** | **0.055** | **0.779** | **0.137** |
| Amide_O | 0.543 | 0.192 | 0.565 | 0.118 | **0.482** | **0.059** | **0.919** | **0.148** |
| Phenyl_C6 | 0.538 | 0.322 | 0.547 | 0.165 | **0.637** | **0.060** | **0.882** | **0.167** |
| Phenyl_C7 | 0.568 | 0.294 | 0.675 | 0.239 | **0.824** | **0.117** | **1.110** | **0.207** |
| Phenyl_C8 | 0.986 | 0.568 | 1.079 | 0.301 | 1.561 | 0.110 | 1.769 | 0.214 |
| Phenyl_C9 | 0.711 | 0.301 | 1.021 | 0.373 | 1.829 | 0.222 | 2.161 | 0.413 |
| Phenyl_C10 | 1.120 | 0.575 | 1.411 | 0.408 | 1.582 | 0.354 | 2.068 | 0.589 |
| Phenyl_C11 | 0.833 | 0.299 | 1.313 | 0.454 | 1.841 | 0.245 | 2.188 | 0.409 |
| Phenyl_C12 | 0.890 | 0.310 | 1.393 | 0.413 | 1.567 | 0.128 | 1.799 | 0.210 |
| COO_C | 0.383 | 0.043 | 0.348 | 0.049 | **0.465** | **0.181** | **1.539** | **0.688** |
| COO_O1 | 0.405 | 0.034 | 0.351 | 0.062 | **0.541** | **0.250** | **1.896** | **0.853** |
| COO_O2 | 0.475 | 0.077 | 0.456 | 0.074 | **0.574** | **0.273** | **1.911** | **0.873** |

[a] *Rows are colored according to their interaction groupings.*

[b] *Bold rows indicate those with significant change altered ($p<0.05$ in two-tailed t-test) due to evolution (TEM-52 – TEM-1).*

*Abbreviations: BL= β-lactam substrate atom; bb = backbone atom; Amide = substrate amide side-chain atoms; COO = substrate carboxylate side-chain atoms; PNM = atoms of PenG corresponding to the penam core; SC = substrate phenyl side-chain, respectively. See Scheme S2.*

**Table S8. CTX-dependent average changes in active site RMSFs due to evolution.**

| CTX RMSF (Å) | ES | | | | AE | | | |
|---|---|---|---|---|---|---|---|---|
| | TEM-1 CTX | | TEM-52 CTX | | TEM-1 CTX | | TEM-52 CTX | |
| Atom [a,b] | Avg. | Std. Dev. | Avg. | Std. Dev. | Avg. | Std. Dev. | Avg. | Std. Dev. |
| S70bbN | 0.339 | 0.026 | 0.353 | 0.013 | 0.420 | 0.056 | 0.477 | 0.063 |
| S70bbH | 0.422 | 0.047 | 0.424 | 0.016 | 0.469 | 0.061 | 0.517 | 0.064 |
| S70O | 0.386 | 0.033 | 0.432 | 0.047 | 0.510 | 0.144 | 0.530 | 0.064 |
| S70bbO | **0.380** | **0.023** | **0.446** | **0.030** | 0.513 | 0.085 | 0.570 | 0.062 |
| K73bbN | 0.346 | 0.021 | 0.363 | 0.019 | 0.430 | 0.047 | 0.574 | 0.151 |
| K73bbH | **0.392** | **0.020** | **0.421** | **0.017** | 0.475 | 0.047 | 0.542 | 0.062 |
| K73N | **0.358** | **0.013** | **0.430** | **0.045** | **0.565** | **0.073** | **0.859** | **0.224** |
| X104bbO | **0.458** | **0.016** | **0.508** | **0.028** | 0.699 | 0.058 | 0.895 | 0.206 |
| Y105bbO | **0.417** | **0.011** | **0.487** | **0.031** | 0.674 | 0.103 | 0.889 | 0.197 |
| S130O | 0.381 | 0.010 | 0.430 | 0.054 | 0.719 | 0.061 | 0.811 | 0.339 |
| S130bbO | **0.366** | **0.010** | **0.412** | **0.025** | 0.573 | 0.097 | 0.662 | 0.257 |
| N132bbN | **0.301** | **0.007** | **0.335** | **0.022** | 0.439 | 0.041 | 0.522 | 0.133 |
| N132bbH | **0.370** | **0.011** | **0.408** | **0.020** | 0.512 | 0.044 | 0.597 | 0.126 |
| N132O | **0.401** | **0.018** | **0.435** | **0.026** | 0.664 | 0.098 | 1.006 | 0.377 |
| N132N | **0.409** | **0.017** | **0.467** | **0.026** | 0.732 | 0.067 | 0.910 | 0.211 |
| E166C | 0.448 | 0.058 | 0.442 | 0.045 | 0.698 | 0.124 | 0.789 | 0.109 |
| E166O1 | 0.571 | 0.181 | 0.747 | 0.269 | 1.183 | 0.299 | 1.272 | 0.118 |
| E166O2 | 0.551 | 0.215 | 0.772 | 0.260 | 1.054 | 0.171 | 1.297 | 0.189 |
| N170O | 0.766 | 0.245 | 0.860 | 0.237 | 1.236 | 0.291 | 1.001 | 0.346 |
| N170N | 0.643 | 0.276 | 0.923 | 0.284 | 0.884 | 0.226 | 0.930 | 0.256 |
| N170bbO | 0.752 | 0.184 | 0.795 | 0.215 | 1.170 | 0.319 | 1.205 | 0.488 |
| K234N | 0.525 | 0.026 | 0.503 | 0.051 | **1.048** | **0.159** | **0.700** | **0.130** |
| A237bbN | **0.331** | **0.021** | **0.361** | **0.020** | **0.402** | **0.019** | **0.484** | **0.058** |
| A237bbH | 0.385 | 0.018 | 0.414 | 0.025 | **0.457** | **0.016** | **0.541** | **0.059** |
| A237bbO | **0.403** | **0.025** | **0.465** | **0.028** | **0.501** | **0.065** | **0.625** | **0.066** |
| X238bbN | 0.379 | 0.023 | 0.380 | 0.012 | 0.471 | 0.049 | 0.527 | 0.067 |
| X238bbH | 0.431 | 0.024 | 0.419 | 0.012 | 0.529 | 0.039 | 0.518 | 0.069 |
| X238bbC | 0.405 | 0.022 | 0.383 | 0.019 | 0.525 | 0.054 | 0.613 | 0.107 |
| E240bbN | 0.417 | 0.021 | 0.439 | 0.025 | **0.526** | **0.057** | **0.715** | **0.123** |
| E240bbH | 0.479 | 0.022 | 0.506 | 0.042 | **0.581** | **0.058** | **0.813** | **0.139** |
| R244N1 | 0.595 | 0.421 | 0.513 | 0.043 | 0.507 | 0.023 | 1.323 | 0.851 |
| R244N2 | 0.618 | 0.384 | 0.614 | 0.075 | **0.587** | **0.017** | **1.467** | **0.770** |
| Water | **0.674** | **0.156** | **1.075** | **0.348** | 1.510 | 0.273 | 1.847 | 0.348 |
| BL_C | 0.410 | 0.035 | 0.462 | 0.082 | 0.464 | 0.084 | 0.536 | 0.082 |
| BL_O | 0.421 | 0.038 | 0.465 | 0.071 | 0.453 | 0.082 | 0.542 | 0.094 |
| CPM_C | 0.440 | 0.027 | 0.540 | 0.114 | 0.573 | 0.153 | 0.604 | 0.092 |
| CPM_C1 | **0.529** | **0.026** | **0.651** | **0.107** | 0.707 | 0.254 | 0.738 | 0.154 |
| CPM_C2 | 0.470 | 0.035 | 0.558 | 0.097 | 0.788 | 0.283 | 0.825 | 0.203 |

| | | | | | | | | |
|---|---|---|---|---|---|---|---|---|
| CPM_S | **0.522** | **0.030** | **0.610** | **0.082** | 1.028 | 0.330 | 1.099 | 0.439 |
| CPM_N | 0.431 | 0.032 | 0.507 | 0.097 | 1.327 | 0.392 | 1.372 | 0.378 |
| CPM_C3 | 0.441 | 0.037 | 0.509 | 0.087 | 1.134 | 0.413 | 1.183 | 0.259 |
| CPM_C7 | **0.592** | **0.032** | **0.718** | **0.095** | 0.852 | 0.395 | 0.897 | 0.185 |
| Amide_N | 0.439 | 0.037 | 0.496 | 0.079 | 0.664 | 0.109 | 0.699 | 0.117 |
| Amide_H | 0.499 | 0.051 | 0.541 | 0.068 | 0.816 | 0.089 | 0.846 | 0.136 |
| Amide_C | 0.422 | 0.024 | 0.472 | 0.082 | 0.801 | 0.169 | 0.816 | 0.132 |
| Amide_O | 0.487 | 0.022 | 0.533 | 0.085 | 1.024 | 0.237 | 0.951 | 0.160 |
| SC_C11 | 0.460 | 0.029 | 0.480 | 0.072 | 0.918 | 0.163 | 0.996 | 0.148 |
| SC_N2 | 0.503 | 0.040 | 0.488 | 0.064 | 0.927 | 0.110 | 1.054 | 0.144 |
| SC_O6 | **0.675** | **0.046** | **0.596** | **0.049** | 1.059 | 0.106 | 1.209 | 0.112 |
| SC_C12 | **0.781** | **0.066** | **0.630** | **0.045** | 1.297 | 0.175 | 1.321 | 0.162 |
| SC_C13 | 0.506 | 0.034 | 0.511 | 0.074 | 1.233 | 0.252 | 1.254 | 0.236 |
| SC_C14 | 0.575 | 0.043 | 0.585 | 0.073 | 1.498 | 0.330 | 1.528 | 0.272 |
| SC_C15 | 0.763 | 0.059 | 0.699 | 0.073 | 1.546 | 0.290 | 1.636 | 0.246 |
| SC_S1 | 0.684 | 0.065 | 0.666 | 0.080 | 1.965 | 0.409 | 1.859 | 0.413 |
| SC_N3 | 0.723 | 0.061 | 0.657 | 0.067 | 1.855 | 0.361 | 1.838 | 0.313 |
| SC_N4 | **1.048** | **0.096** | **0.923** | **0.071** | 2.239 | 0.417 | 2.307 | 0.382 |
| COO_C | 0.428 | 0.023 | 0.533 | 0.135 | 0.900 | 0.473 | 1.075 | 0.214 |
| COO_O1 | 0.479 | 0.042 | 0.617 | 0.150 | 1.062 | 0.560 | 1.339 | 0.300 |
| COO_O2 | 0.510 | 0.179 | 0.538 | 0.138 | 0.957 | 0.430 | 1.186 | 0.267 |

[a] *Rows are colored according to their interaction groupings.*

[b] *Bold rows indicate those with significant change altered ($p<0.05$ in two-tailed t-test) due to evolution (TEM-52 – TEM-1).*

*Abbreviations: BL= β-lactam substrate atom; bb = backbone atom; Amide = substrate amide side-chain atoms; COO = substrate carboxylate side-chain atoms; CPM = atoms of CTX corresponding to the cephem core; SC = substrate oxyimino/aminothiazolyl-ring, respectively. See Scheme S2.*

**Table S9. Collective RMSF changes in the evolution of TEM-1 to TEM-52 for PenG and CTX.**

| | TEM-1 RMSF (Å) | | | TEM-52 RMSF (Å) | | |
|---|---|---|---|---|---|---|
| **State & Substrate** | **Interaction[a]** | **Median** | **68% CI** | **Interaction[a]** | **Median** | **68% CI** |
| ES PenG | **Total** | **0.41** | **[0.33, 0.58]** | **Total** | **0.48** | **[0.38, 0.87]** |
| | **Oxyanion** | 0.34 | [0.31, 0.37] | **Oxyanion** | 0.41 | [0.37, 0.46] |
| | **Chemistry** | 0.49 | [0.35, 0.64] | **Chemistry** | 0.75 | [0.49, 1.10] |
| | **Supporting** | 0.41 | [0.35, 0.52] | **Supporting** | 0.51 | [0.41, 0.73] |
| | **COO** | | | **COO** | | |
| | **BL** | 0.44 | [0.38, 0.56] | **BL** | 0.60 | [0.51, 0.70] |
| ES CTX | **Total** | **0.44** | **[0.37, 0.66]** | **Total** | **0.49** | **[0.39, 0.68]** |
| | **Oxyanion** | 0.36 | [0.32, 0.39] | **Oxyanion** | 0.38 | [0.35, 0.43] |
| | **Chemistry** | 0.44 | [0.36, 0.75] | **Chemistry** | 0.51 | [0.40, 1.24] |
| | **Supporting** | 0.38 | [0.35, 0.42] | **Supporting** | 0.43 | [0.38, 0.49] |
| | **COO** | 0.42 | [0.38, 0.48] | **COO** | 0.50 | [0.39, 0.86] |
| | **BL** | 0.49 | [0.41, 0.73] | **BL** | 0.58 | [0.46, 0.73] |
| AE PenG | **Total** | **0.45** | **[0.35, 0.83]** | **Total** | **0.88** | **[0.53, 1.68]** |
| | **Oxyanion** | 0.35 | [0.32, 0.39] | **Oxyanion** | 0.51 | [0.45, 0.58] |
| | **Chemistry** | 0.65 | [0.36, 1.15] | **Chemistry** | 1.23 | [0.59, 1.62] |
| | **Supporting** | 0.46 | [0.38, 0.64] | **Supporting** | 0.87 | [0.54, 1.44] |
| | **COO** | 0.47 | [0.37, 0.65] | **COO** | 1.38 | [0.90, 2.29] |
| | **BL** | 0.44 | [0.35, 0.56] | **BL** | 0.84 | [0.60, 1.26] |
| AE CTX | **Total** | **0.70** | **[0.46, 1.31]** | **Total** | **0.85** | **[0.53, 1.42]** |
| | **Oxyanion** | 0.43 | [0.39, 0.46] | **Oxyanion** | 0.51 | [0.45, 0.58] |
| | **Chemistry** | | | **Chemistry** | | |
| | **Supporting** | 0.56 | [0.49, 0.62] | **Supporting** | 0.78 | [0.63, 0.96] |
| | **COO** | 0.63 | [0.58, 1.12] | **COO** | 0.82 | [0.60, 1.89] |
| | **BL** | 0.50 | [0.43, 0.56] | **BL** | 0.63 | [0.56, 0.69] |

[a] *Interaction groups and colors according to Scheme S2. COO and BL refer to binding interactions to the substrate carboxylate group and β-lactam core/amide side-chain, respectively.*

**Table S10. PenG-dependent average changes in active site distances due to evolution.**

| PenG Distance (Å) | ES | | | | AE | | | |
|---|---|---|---|---|---|---|---|---|
| | TEM-1 PenG | | TEM-52 PenG | | TEM-1 PenG | | TEM-52 PenG | |
| **Distance Pair [a,b]** | **Avg.** | **Std. Dev.** | **Avg.** | **Std. Dev.** | **Avg.** | **Std. Dev.** | **Avg.** | **Std. Dev.** |
| S70bbN – O | 3.100 | 0.100 | 3.240 | 0.100 | 2.740 | 0.100 | 2.800 | 0.100 |
| A237bbN – O | 2.800 | 0.100 | 2.800 | 0.100 | 2.900 | 0.100 | 2.900 | 0.100 |
| S70O – BL_O | 3.100 | 0.100 | 3.120 | 0.100 | | | | |
| S70O – BL_C | 3.100 | 0.100 | 3.160 | 0.100 | | | | |
| E166C – BL_C | 6.660 | 0.100 | 6.800 | 0.100 | 6.400 | 0.100 | 6.780 | 0.458 |
| Water1 – BL_C | **3.640** | **0.100** | **3.460** | **0.102** | **3.340** | **0.174** | **2.960** | **0.206** |
| Water1 – BL_O | 3.400 | 0.100 | 3.240 | 0.120 | 3.820 | 0.117 | 3.740 | 0.150 |
| Water1 – E166C | **3.400** | **0.100** | **3.780** | **0.117** | **3.360** | **0.120** | **4.040** | **0.609** |
| Water1 – S70O | 2.700 | 0.100 | 2.680 | 0.194 | **3.140** | **0.224** | **2.640** | **0.356** |
| S70O – E166C | 5.200 | 0.100 | 5.200 | 0.167 | 6.120 | 0.100 | 6.180 | 0.453 |
| S130O – K234N | 2.740 | 0.100 | 2.880 | 0.117 | **3.000** | **0.253** | **5.660** | **2.207** |
| S130O – COO_C | 3.700 | 0.100 | 3.780 | 0.160 | 4.040 | 0.480 | 5.300 | 1.134 |
| K234N – COO_C | 5.000 | 0.253 | 5.020 | 0.279 | **5.620** | **0.640** | **7.500** | **1.003** |
| R244N1 – COO_C | 3.900 | 0.100 | 3.900 | 0.100 | 2.800 | 0.100 | 2.850 | 0.100 |
| R244N2 – COO_C | 5.100 | 0.100 | 5.100 | 0.100 | **3.640** | **0.224** | **3.100** | **0.200** |
| N132N – AmideO | **3.140** | **0.102** | **3.860** | **0.372** | **2.980** | **0.100** | **4.600** | **0.982** |
| A237bbO – AmideN | 3.080 | 0.100 | 3.160 | 0.100 | 3.000 | 0.100 | 2.920 | 0.100 |
| S70O – S130O | 4.000 | 0.100 | 4.000 | 0.110 | **3.960** | **0.100** | **5.560** | **0.736** |
| K73N – N132O | 2.800 | 0.100 | 2.800 | 0.100 | **2.780** | **0.100** | **5.220** | **1.130** |
| K73N – S130bbO | 2.900 | 0.100 | 2.900 | 0.100 | **3.040** | **0.100** | **4.080** | **0.741** |
| K73N – S70O | 2.980 | 0.100 | 3.000 | 0.100 | **3.800** | **0.100** | **4.520** | **0.504** |
| K73N – E166C | 3.800 | 0.100 | 3.800 | 0.110 | 3.760 | 0.100 | 3.780 | 0.538 |
| X104bbO – N132N | 2.900 | 0.100 | 2.900 | 0.100 | **2.900** | **0.100** | **5.200** | **0.537** |
| Y105bbO – N132bbN | 3.200 | 0.100 | 3.200 | 0.100 | **3.240** | **0.100** | **4.980** | **1.139** |
| E166C – N170N | 3.820 | 0.100 | 4.080 | 0.248 | **4.060** | **0.100** | **5.300** | **0.965** |
| N170bbO – E240bbN | **3.000** | **0.100** | **3.340** | **0.224** | 3.040 | 0.100 | 4.380 | 1.411 |
| Water1 – K73N | **3.720** | **0.100** | **4.040** | **0.136** | 3.260 | 0.102 | 3.940 | 0.734 |
| Water1 – N170N | **3.680** | **0.100** | **4.040** | **0.150** | 3.520 | 0.100 | 3.780 | 0.515 |
| Water1 – N170O | **2.800** | **0.100** | **3.160** | **0.280** | 2.980 | 0.100 | 3.500 | 0.718 |

*[a] Abbreviations: BL= β-lactam substrate atom; bb = backbone atom; Amide = substrate amide side-chain atoms; COO = substrate carboxylate side-chain atoms. See Figure 5 and Scheme S2 for color coding of interactions.*

*[b] Bold rows entries indicate those that are significantly altered between TEM-1 and TEM-52 based on a two-tailed t-test ($p<0.05$ in two-tailed t-test) and represented in Figure 5-6.*

**Table S11. CTX-dependent average changes in active site distances due to evolution.**

| CTX Distance (Å) | ES | | | | AE | | | |
|---|---|---|---|---|---|---|---|---|
| | TEM-1 CTX | | TEM-52 CTX | | TEM-1 CTX | | TEM-52 CTX | |
| Distance Pair [a,b] | Avg. | Std. Dev. | Avg. | Std. Dev. | Avg. | Std. Dev. | Avg. | Std. Dev. |
| S70bbN – O | 3.445 | 0.223 | 3.220 | 0.117 | 2.820 | 0.100 | 2.860 | 0.100 |
| A237bbN – O | 2.873 | 0.045 | 2.900 | 0.100 | 2.880 | 0.100 | 2.900 | 0.100 |
| S70O – BL_O | 3.100 | 0.100 | 3.080 | 0.040 | | | | |
| S70O – BL_C | 3.445 | 0.078 | 3.380 | 0.040 | | | | |
| E166C – BL_C | **7.082** | **0.159** | **7.500** | **0.100** | 6.700 | 0.167 | 7.040 | 0.472 |
| Water1 – BL_C | 3.900 | 0.230 | 3.800 | 0.167 | 3.180 | 0.117 | 3.140 | 0.273 |
| Water1 – BL_O | 3.545 | 0.188 | 3.500 | 0.167 | 3.840 | 0.102 | 3.820 | 0.542 |
| Water1 – S70O | **2.691** | **0.100** | **2.940** | **0.162** | 3.060 | 0.250 | 3.020 | 0.223 |
| Water1 – E166C | **3.500** | **0.141** | **3.780** | **0.133** | 3.700 | 0.210 | 4.280 | 0.700 |
| S70O – E166C | **5.200** | **0.085** | **5.880** | **0.040** | 6.300 | 0.155 | 6.500 | 0.533 |
| S130O – K234N | 2.791 | 0.029 | 2.800 | 0.100 | 4.180 | 1.356 | 3.300 | 0.100 |
| S130O – COO_C | **3.482** | **0.039** | **3.640** | **0.049** | **5.800** | **0.253** | **4.900** | **0.100** |
| K234N – COO_C | 4.982 | 0.057 | 5.000 | 0.110 | **7.980** | **0.618** | **5.640** | **1.031** |
| R244N1 – COO_C | 4.236 | 1.113 | 3.920 | 0.075 | 3.400 | 0.179 | 6.160 | 2.968 |
| R244N2 – COO_C | 5.273 | 0.377 | 5.240 | 0.080 | 4.060 | 0.174 | 6.220 | 2.500 |
| N132N – AmideO | 2.809 | 0.029 | 2.820 | 0.040 | **3.760** | **0.615** | **5.020** | **0.926** |
| A237bbO – AmideN | 3.073 | 0.062 | 3.040 | 0.049 | 3.180 | 0.040 | 3.120 | 0.100 |
| S70O – S130O | **3.645** | **0.078** | **3.560** | **0.049** | 4.100 | 0.329 | 4.280 | 0.952 |
| K73N – N132O | **2.855** | **0.050** | **2.800** | **0.100** | **2.840** | **0.049** | **3.680** | **0.806** |
| K73N – S130bbO | 2.855 | 0.050 | 2.900 | 0.100 | **2.920** | **0.075** | **3.660** | **0.647** |
| K73N – S70O | 2.955 | 0.050 | 3.000 | 0.100 | 4.360 | 0.287 | 4.520 | 0.371 |
| K73N – E166C | **3.827** | **0.045** | **4.260** | **0.049** | **3.800** | **0.063** | **3.540** | **0.100** |
| X104bbO – N132N | 2.900 | 0.100 | 2.900 | 0.100 | 2.920 | 0.098 | 3.320 | 0.445 |
| Y105bbO – N132bbN | 3.109 | 0.029 | 3.100 | 0.100 | 3.160 | 0.100 | 3.200 | 0.100 |
| E166C – N170N | **4.691** | **0.760** | **5.620** | **0.098** | **6.240** | **1.174** | **4.680** | **0.100** |
| N170bbO – E240bbN | **3.309** | **0.320** | **5.060** | **0.136** | **4.120** | **0.584** | **4.940** | **0.273** |
| Water1 – K73N | **3.709** | **0.116** | **3.500** | **0.210** | 3.700 | 0.494 | 3.920 | 0.991 |
| Water1 – N170N | **3.400** | **0.113** | **4.640** | **0.174** | **4.240** | **0.393** | **3.660** | **0.314** |
| Water1 – N170O | **3.836** | **0.971** | **6.360** | **0.301** | **5.580** | **1.306** | **3.040** | **0.287** |

[a] *Abbreviations: BL= β-lactam substrate atom; bb = backbone atom; Amide = substrate amide side-chain atoms; COO = substrate carboxylate side-chain atoms. See Figure 5 and Scheme S2 for color coding of interactions.*

[b] *Bold rows entries indicate those that are significantly altered between TEM-1 and TEM-52 based on a two-tailed t-test (p<0.05 in two-tailed t-test) and represented in Figure 5-6.*

**Table S12. Collective distance changes in the evolution of TEM-1 to TEM-52 for PenG and CTX.**

| | TEM-1 Distances (Å) | | | TEM-52 Distance (Å) | | |
|---|---|---|---|---|---|---|
| **State & Substrate** | **Interaction[a]** | **Median** | **68% CI** | **Interaction[a]** | **Median** | **68% CI** |
| ES PenG | **Total** | 3.26 | [2.84, 4.01] | **Total** | 3.39 | [2.90, 4.23] |
| | **Oxyanion** | | | **Oxyanion** | | |
| | **Chemistry** | 3.52 | [3.35, 3.69] | **Chemistry** | 3.61 | [3.41, 3.83] |
| | **Supporting** | 3.34 | [2.82, 3.74] | **Supporting** | 3.76 | [3.13, 4.11] |
| | **COO** | | | **COO** | | |
| | **BL** | 3.14 | [3.04, 3.24] | **BL** | 3.86 | [3.49, 4.23] |
| ES CTX | **Total** | 3.44 | [2.84, 4.89] | **Total** | 3.56 | [2.90, 5.22] |
| | **Oxyanion** | | | **Oxyanion** | | |
| | **Chemistry** | 4.33 | [2.72, 7.03] | **Chemistry** | 4.39 | [2.99, 7.46] |
| | **Supporting** | 3.63 | [2.94, 4.00] | **Supporting** | 4.34 | [3.37, 5.67] |
| | **COO** | 3.48 | [3.44, 3.52] | **COO** | 3.64 | [3.59, 3.69] |
| | **BL** | | | **BL** | | |
| AE PenG | **Total** | 3.28 | [2.87, 4.05] | **Total** | 4.06 | [2.88, 6.05] |
| | **Oxyanion** | | | **Oxyanion** | | |
| | **Chemistry** | 3.3 | [3.08, 3.47] | **Chemistry** | 3.01 | [2.58, 4.08] |
| | **Supporting** | 3.25 | [2.85, 4.00] | **Supporting** | 4.96 | [4.03, 5.94] |
| | **COO** | 3.64 | [3.00, 5.66] | **COO** | 5.55 | [3.03, 7.96] |
| | **BL** | 2.98 | [2.88, 3.08] | **BL** | 4.6 | [3.62, 5.58] |
| AE CTX | **Total** | 3.78 | [2.93, 5.97] | **Total** | 3.77 | [3.00, 5.50] |
| | **Oxyanion** | | | **Oxyanion** | | |
| | **Chemistry** | | | **Chemistry** | | |
| | **Supporting** | 3.87 | [2.88, 5.75] | **Supporting** | 3.69 | [3.13, 4.75] |
| | **COO** | 6.43 | [5.68, 8.27] | **COO** | 4.97 | [4.78, 6.13] |
| | **BL** | 3.76 | [3.15, 4.37] | **BL** | 5.02 | [4.09, 5.94] |

[a] *Interaction groups and colors according to Scheme S2. COO and BL refer to binding interactions to the substrate carboxylate group and β-lactam core/amide side-chain, respectively.*

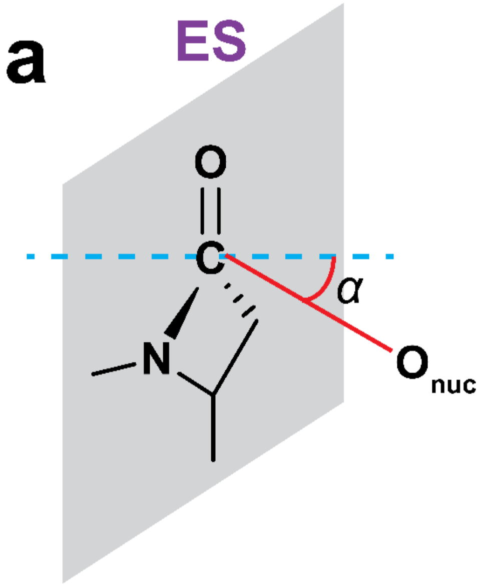

***Scheme S3. Angle of nucleophilic attack in the ES and AE complexes.*** *The angle of nucleophilic attack (α) for the* ***(a)*** *ES and* ***(b)*** *AE complexes are defined as normal to the plane of the four-atoms defining the sp*$^2$ *hybridized C in the <u>C</u>=O. An angle approaching zero degrees is expected to be indicative of better chemical positioning for nucleophilic addition at the C=O.*

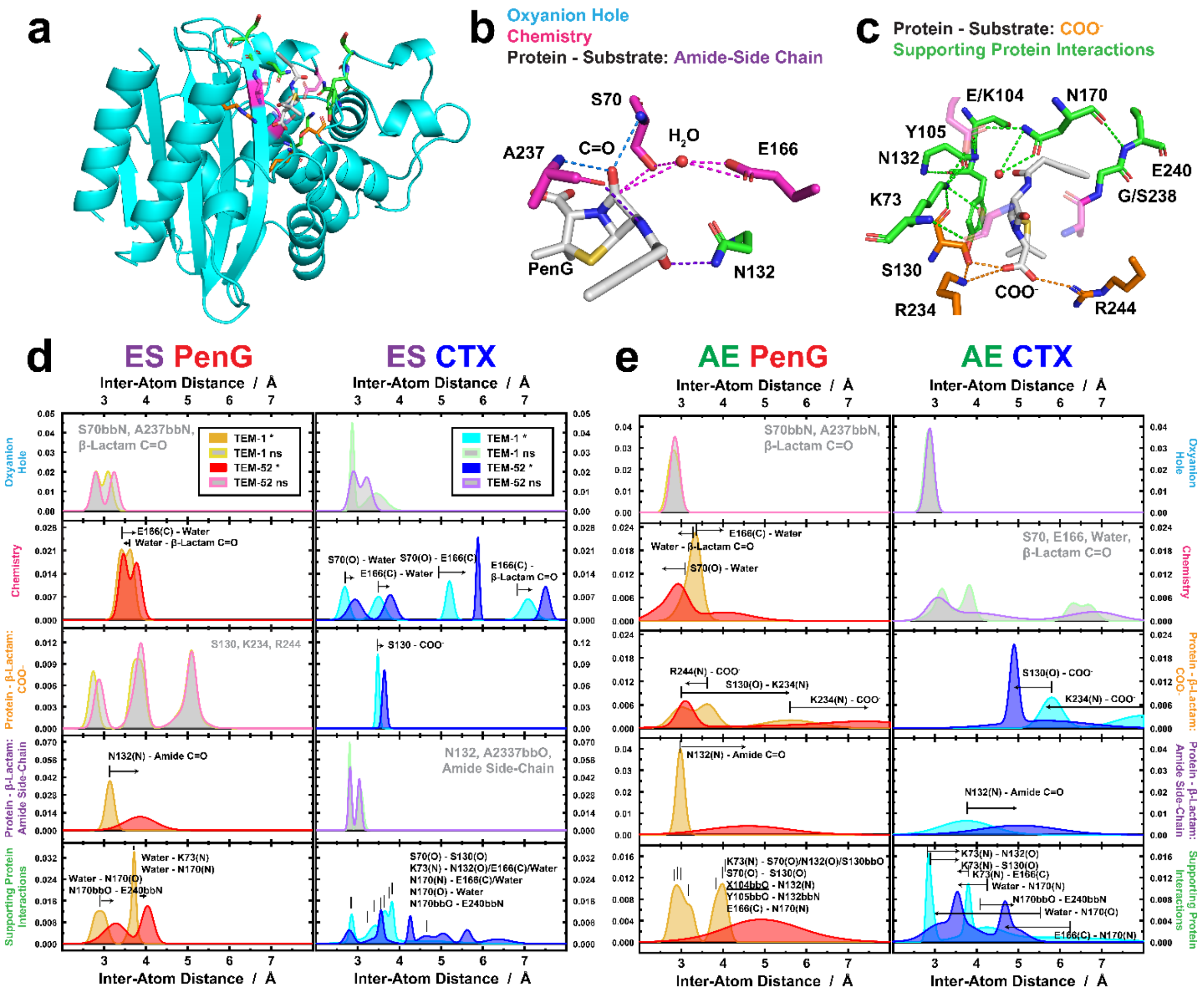

***Figure S15. Alternative probability density representation of significant active site distance changes over the course of evolution from TEM-1 to TEM-52 for PenG and CTX.*** ***(a)*** *Overall structure of the TEM-1 PenG ES complex with active site residues shown according to their broad classifications and role in catalysis.* ***(b,c)*** *Inter-atom interactions are classified according to their role in catalysis for key residues in the active site: oxyanion hole (blue), chemistry (pink), protein-substrate interactions (orange for the substrate COO<sup>-</sup> and purple for the amide side-chain), or supporting inter-amino acid interactions that participate in the extended H-bond network for chemical positioning (green). Protons removed for clarity.* ***(d,e)*** *Significant (*) average distances and corresponding heterogeneity of key active site non-covalent interactions shown as a normalized probability distributions for the (b) ES and (c) AE complexes of TEM-1 and -52 with PenG and CTX, respectively. Significant distance changes correspond to $p < 0.05$ in a two-tailed t-test (Scheme S2 and Tables S10-12). Vertical bars represent the average positions observed in TEM-1 for a given complex and arrows denote the*

*direction of change in TEM-52. In cases where no significant (ns) changes were observed among a given interaction group, the total probability densities are shown for all interactions and shaded in grey for TEM-1 (green outline) and TEM-52 (purple outline). Averages distances represent comparison across 3-5 average structures each from 50-200 ns of the MD trajectories.*

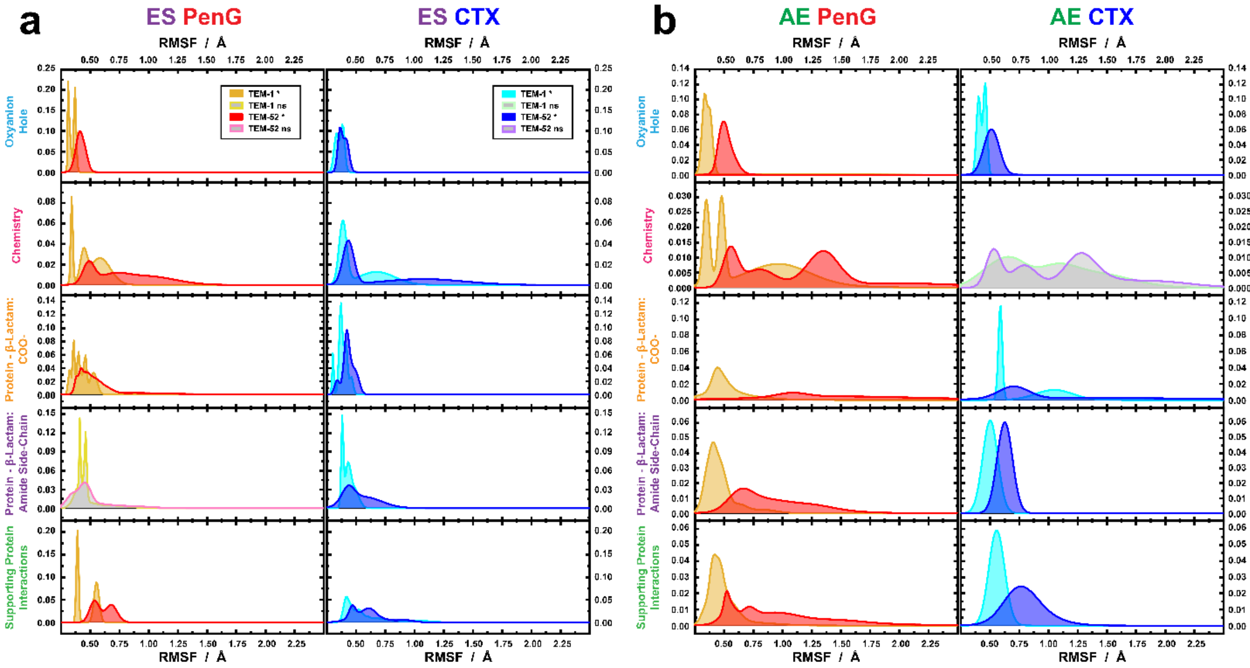

***Figure S16. Alternative probability density representation of significant active site RMSF changes over the course of evolution from TEM-1 to TEM-52 for PenG and CTX. (a,b)*** *Significant (*) average RMSF and corresponding heterogeneity of key active site atoms shown as a normalized probability distributions for the* ***(a)*** *ES and* ***(b)*** *AE complexes of TEM-1 and -52 with PenG (orange, red) and CTX (cyan, blue), respectively. Significant RMSF changes correspond to* $p < 0.05$ *in a two-tailed t-test (Scheme S2 and Tables S7-9). In cases where no significant (ns) changes were observed among a given interaction group, the total probability densities are shown for all interactions and shaded in grey for TEM-1 (pink outline) and TEM-52 (purple outline). Averages RMSFs and standard deviations represent comparison across 3-5 trajectories of 50-200 ns each from the MD trajectories.*

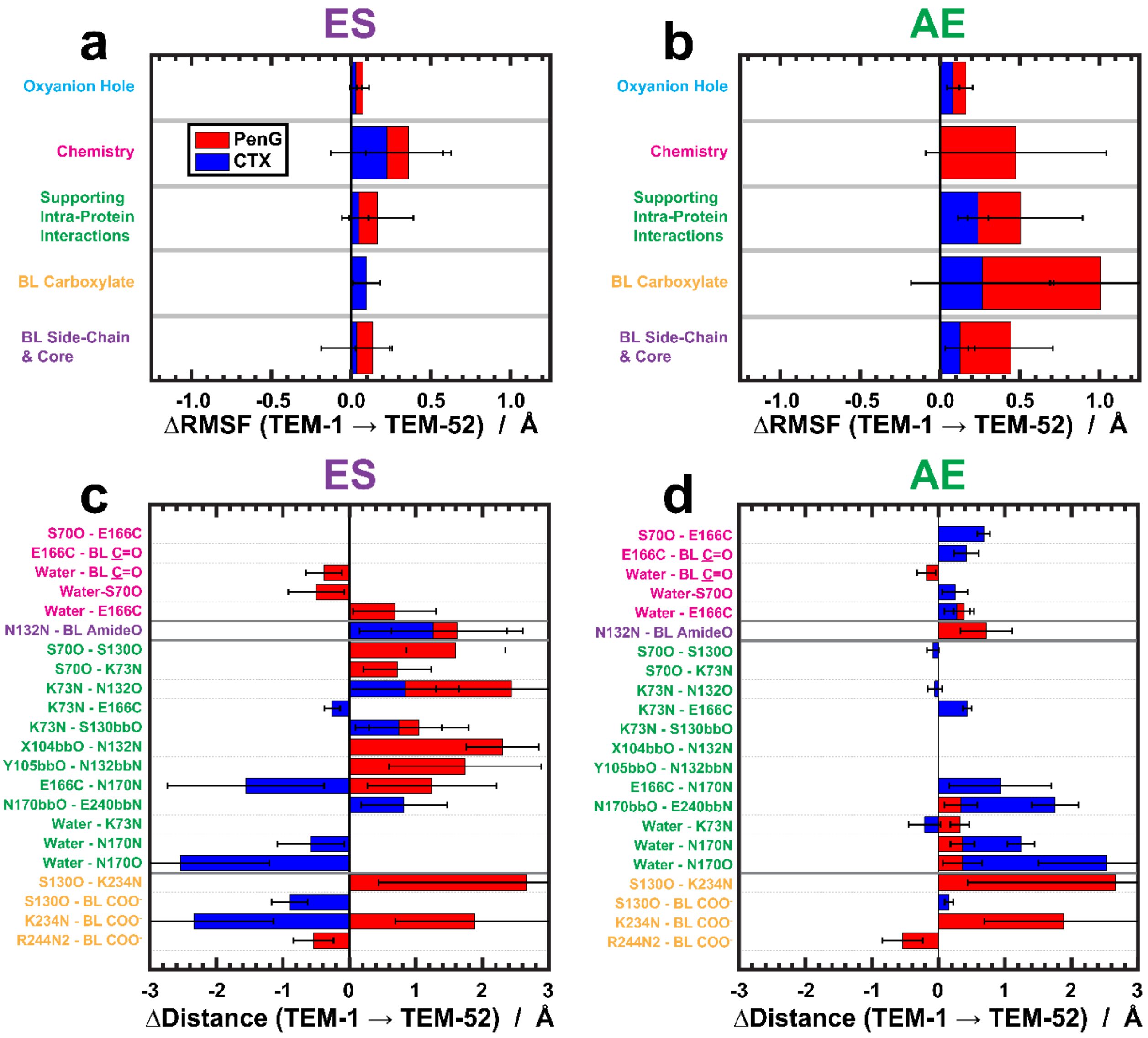

***Figure S17. Alternative representation of significant active site RMSF and distances changes over the course of evolution from TEM-1 to TEM-52 for PenG and CTX. (a,b)*** *Significant average change in RMSF (TEM-52 – TEM-1) and corresponding heterogeneity of key active site atoms for the* ***(a)*** *ES and* ***(b)*** *AE complexes with PenG (red) and CTX (blue), respectively.* ***(c,d)*** *Significant average change in distance (TEM-52 – TEM-1) between key active site atoms for the* ***(c)*** *ES and* ***(d)*** *AE complexes with PenG and CTX. Significant changes correspond to $p < 0.05$ in a two-tailed t-test for the interaction groups according to Scheme S2 (Tables S7-9). Averages RMSFs and standard deviations represent comparison across 3-5 trajectories of 50-200 ns each from the MD trajectories. Standard deviations correspond to*

$\sqrt{\sigma_{TEM1}^2 + \sigma_{TEM52}^2}$. *Abbreviations: BL = substrate β-lactam antibiotic, $COO^-$ = substrate's carboxylate functional group (C-atom), bb = backbone, Amide = substrate's amide side-chain.*

**Table S13. Average changes in active site RMSFs between TEM-1 and TEM-52 with PenG and CTX, respectively.** (E) and (S) refers to changes occurring on the enzyme or substrate, respectively.

| ΔRMSF [(TEM-1 (PenG) – TEM-52 (CTX)] | | |
|---|---|---|
| **Active Site Fluctuations** | **ES** | **AE** |
| Oxyanion Hole (E) | -0.06 ± 0.02 | -0.16 ± 0.02 |
| Oxyanion Hole (S) | | -0.20 ± 0.01 |
| Chemistry (E) | | |
| Chemistry (S) | | |
| β-Lactam Carboxylate (E) | -0.02 ± 0.15 | -0.94 ± 0.04 |
| β-Lactam Carboxylate (S) | -0.18 ± 0.03 | -0.67 ± 0.09 |
| β-Lactam Amide Side-Chain (E) | 0.01 ± 0.08 | -0.33 ± 0.11 |
| β-Lactam Amide Side-Chain (S) | | -0.39 ± 0.06 |
| Supporting Protein Interactions | | |

*Standard deviations correspond to the variance across all of the average RMSF changes for a given active site interaction type. See Scheme S2 for interaction groupings.*

**Table S14. Average changes in active site distances between TEM-1 and TEM-52 with PenG and CTX, respectively.**

| Δdist. [TEM-1 (PenG) – TEM-52 (CTX)] [a] | | |
|---|---|---|
| **Distance Pair** | **ES** | **AE** |
| S70O – [BL-C=O (C)] | -0.28 ± 0.11 | |
| S70O – E166C | -0.68 ± 0.11 | |
| S70 – Water | -0.24 ± 0.19 | |
| E166C – [BL-C=O (C)] | -0.84 ± 0.14 | -0.64 ± 0.48 |
| E166C – Water | -0.38 ± 0.17 | -0.92 ± 0.71 |
| S130O – K234N | | -0.30 ± 0.27 |
| S130O – COO(C) | | -0.86 ± 0.49 |
| R244N1 – COO(C) | -0.14 ± 0.13 | -3.36 ± 2.97 |
| N132N – [BL-AmideO] | 0.32 ± 0.11 | -2.04 ± 0.93 |
| S70O – S130O | 0.44 ± 0.11 | |
| S70O – K73N | | -0.72 ± 0.38 |
| K73N – N132O | | -0.90 ± 0.81 |
| K73N – E166C | -0.46 ± 0.11 | 0.22 ± 0.14 |
| E166C – N170N | -1.80 ± 0.14 | -0.62 ± 0.14 |
| N170bbO – E240bbN | -2.06 ± 0.17 | -1.90 ± 0.29 |
| N170N – Water | -0.96 ± 0.20 | |
| N170O – Water | -3.56 ± 0.32 | |

[a] *Standard deviations correspond to* $\sqrt{\sigma^2_{TEM1} + \sigma^2_{TEM52}}$. *Abbreviations: BL= β-lactam substrate atom; bb = backbone atom; Amide = substrate amide side-chain atoms; COO = substrate carboxylate side-chain atoms; core = atoms of substrate corresponding to the cephem or penam core; side-chain = substrate phenylacetamido or oxyimino/aminothiazolyl-ring, respectively. See Scheme S2 for interaction groupings.*

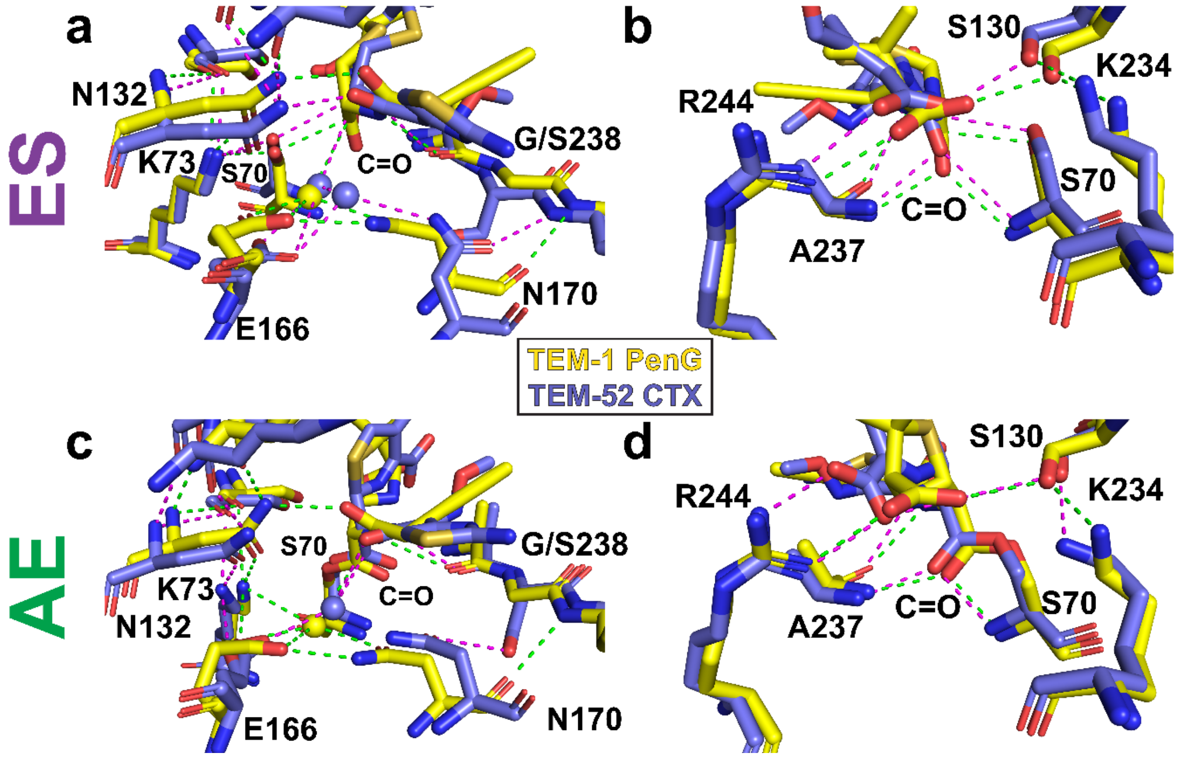

***Figure S18. Structural comparison of TEM-1 PenG versus TEM-52 CTX complexes.*** *Structural overlay of representative average structures of the* ***(a,b)*** *ES complex and* ***(c,d)*** *AE complex of TEM-1 PenG (yellow) relative to TEM-52 CTX (purple).* ***(a,c)*** *The changes in oxyanion hole and extended H-bond network around S70, E166, relative to the β-lactam C=O.* ***(b, d)*** *The substrate-induced changes in structure surrounding the carboxylate functional group of PenG and CTX of the ES and AE states, respectively. Dashed lines show non-covalent interactions ≤ 3.5 Å between nearby interacting functional groups for TEM-1 PenG (green) and TEM-52 CTX (magenta). Changes in structural displacement in terms of the*

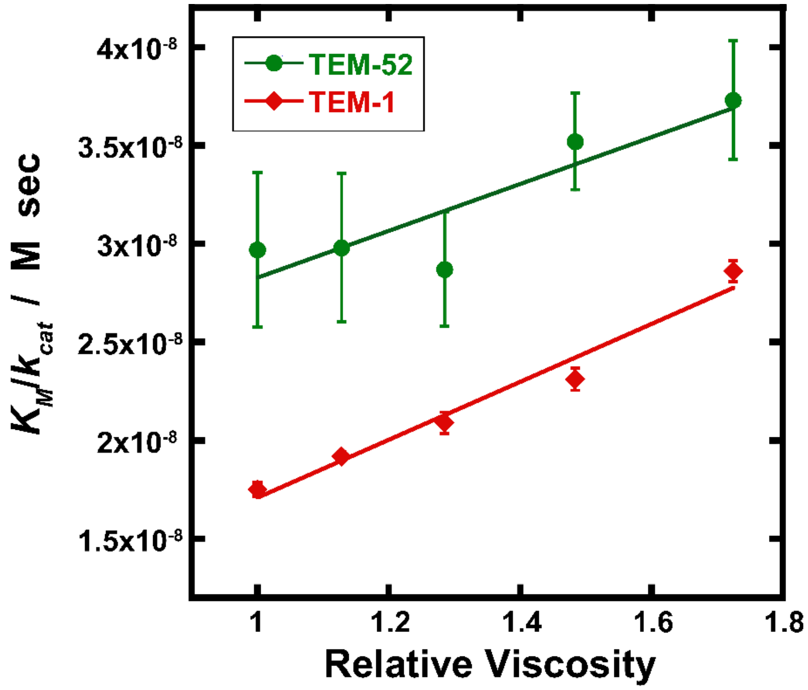

***Figure S19. Viscosity-dependence of PenG hydrolysis by TEM β-lactamases.*** *The change in $K_M/k_{cat}$ as a function of relative viscosity using glycerol as a viscogen (0-20% v/v) enables determination of kinetic parameters as shown in Table S15.*[9,10] *Error bars reflect the standard deviation across 10 measurements (see Methods section).*

**Table S15. Evolutionary change in on- and off-rates of PenG hydrolysis with TEM-1 and TEM-52.**

| **Enzyme with PenG** | **$k_1$ (μM$^{-1}$ s$^{-1}$)** | **$k_{-1}$ (s$^{-1}$)** [a] | **$k_2$ (s$^{-1}$)** | **$k_3$ (s$^{-1}$)** |
|---|---|---|---|---|
| **TEM-1** | 78.1 ± 3.6 | 553 | 5529 | 2702 |
| **TEM-52** | 84.0 ± 41.5 | 349 | 249 | 52 |

[a] *Propagated errors of $k_{-1}$ are of a similar magnitude as the values due to the uncertainty in the y-intercept of Figure S18.*

15. Frisch, M. J.; Trucks, G. W.; Schlegel, H. B.; Scuseria, G. E.; Robb, M. A.; Cheeseman, J. R.; Scalmani, G.; Barone, V.; Petersson, G. A.; Nakatsuji, H.; Li, X.; Caricato, M.; Marenich, A. V.; Bloino, J.; Janesko, B. G.; Gomperts, R.; Mennucci, B.; Hratchian, H. P.; Ortiz, J. V.; Izmaylov, A. F.; Sonnenberg, J. L.; Williams-Young, D.; Ding, F.; Lipparini, F.; Egidi, F.; Goings, J.; Peng, B.; Petrone, A.; Henderson, T.; Ranasinghe, D.; Zakrzewski, V. G.; Gao, J.; Rega, N.; Zheng, G.; Liang, W.; Hada, M.; Ehara, M.; Toyota, K.; Fukuda, R.; Hasegawa, J.; Ishida, M.; Nakajima, T.; Honda, Y.; Kitao, O.;

Nakai, H.; Vreven, T.; Throssell, K.; Montgomery, J. A., Jr.; Peralta, J. E.; Ogliaro, F.; Bearpark, M. J.; Heyd, J. J.; Brothers, E. N.; Kudin, K. N.; Staroverov, V. N.; Keith, T. A.; Kobayashi, R.; Normand, J.; Raghavachari, K.; Rendell, A. P.; Burant, J. C.; Iyengar, S. S.; Tomasi, J.; Cossi, M.; Millam, J. M.; Klene, M.; Adamo, C.; Cammi, R.; Ochterski, J. W.; Martin, R. L.; Morokuma, K.; Farkas, O.; Foresman, J. B.; Fox, D. J., Gaussian 09, Rev E.01. Gaussian 09, Rev E.01, Gaussian, Inc.: Wallingford, CT, 2016.

16. Lemkul, J., From Proteins to Perturbed Hamiltonians: A Suite of Tutorials for the GROMACS-2018 Molecular Simulation Package [Article v1.0]. *Living J. Comp. Mol. Sci* **2019,** *1* (1), 5068.

------------------------------------------------------------------------------------------------------------------------

### MD Files and Parameters:

##### PEN.gro

```
  289PEN      C 4061   4.665   3.745   3.526
  289PEN     C1 4062   4.812   3.780   3.558
  289PEN     C2 4063   4.653   3.974   3.635
  289PEN      H 4064   4.633   3.668   3.597
  289PEN     H1 4065   4.692   4.058   3.576
  289PEN      S 4066   4.791   3.877   3.711
  289PEN      N 4067   4.588   3.867   3.559
  289PEN     C3 4068   4.517   3.997   3.710
  289PEN     H2 4069   4.467   4.090   3.681
  289PEN     C4 4070   4.467   3.876   3.628
  289PEN      O 4071   4.370   3.803   3.631
  289PEN     N1 4072   4.507   3.983   3.854
  289PEN     H3 4073   4.472   3.895   3.892
  289PEN     C5 4074   4.546   4.075   3.947
  289PEN     O1 4075   4.589   4.185   3.915
  289PEN     C6 4076   4.537   4.026   4.091
  289PEN     H4 4077   4.474   3.937   4.094
  289PEN     H5 4078   4.485   4.104   4.148
  289PEN     C7 4079   4.673   3.995   4.153
  289PEN     C8 4080   4.684   3.992   4.293
  289PEN     C9 4081   4.788   3.974   4.074
  289PEN    C10 4082   4.810   3.978   4.353
  289PEN     H6 4083   4.596   4.006   4.355
  289PEN    C11 4084   4.914   3.957   4.135
  289PEN     H7 4085   4.783   3.974   3.966
  289PEN    C12 4086   4.924   3.962   4.274
  289PEN     H8 4087   4.819   3.982   4.461
  289PEN     H9 4088   5.003   3.943   4.074
  289PEN    H10 4089   5.022   3.955   4.321
  289PEN    C13 4090   4.644   3.685   3.388
  289PEN     O2 4091   4.582   3.748   3.303
  289PEN     O3 4092   4.681   3.570   3.370
  289PEN    C14 4093   4.881   3.861   3.445
  289PEN    H11 4094   4.984   3.883   3.472
  289PEN    H12 4095   4.884   3.803   3.352
  289PEN    H13 4096   4.829   3.955   3.425
  289PEN    C15 4097   4.897   3.657   3.594
  289PEN    H14 4098   4.852   3.601   3.676
  289PEN    H15 4099   4.909   3.589   3.508
  289PEN    H16 4100   4.998   3.688   3.626
```

##### PEN.top

```
[ defaults ]
; nbfunc        comb-rule       gen-pairs       fudgeLJ fudgeQQ
1               2               yes             0.5     0.8333

[ atomtypes ]
;name   bond_type     mass     charge   ptype   sigma         epsilon       Amb
 c3       c3          0.00000  0.00000   A     3.39967e-01   4.57730e-01 ; 1.91  0.1094
 cy       cy          0.00000  0.00000   A     3.39967e-01   3.59824e-01 ; 1.91  0.0860
 h1       h1          0.00000  0.00000   A     2.47135e-01   6.56888e-02 ; 1.39  0.0157
 h2       h2          0.00000  0.00000   A     2.29317e-01   6.56888e-02 ; 1.29  0.0157
 ss       ss          0.00000  0.00000   A     3.56359e-01   1.04600e+00 ; 2.00  0.2500
 nj       nj          0.00000  0.00000   A     3.25000e-01   7.11280e-01 ; 1.82  0.1700
 c        c           0.00000  0.00000   A     3.39967e-01   3.59824e-01 ; 1.91  0.0860
 o        o           0.00000  0.00000   A     2.95992e-01   8.78640e-01 ; 1.66  0.2100
 n        n           0.00000  0.00000   A     3.25000e-01   7.11280e-01 ; 1.82  0.1700
```

```
 hn      hn        0.00000  0.00000   A    1.06908e-01   6.56888e-02 ; 0.60  0.0157
 hc      hc        0.00000  0.00000   A    2.64953e-01   6.56888e-02 ; 1.49  0.0157
 ca      ca        0.00000  0.00000   A    3.39967e-01   3.59824e-01 ; 1.91  0.0860
 ha      ha        0.00000  0.00000   A    2.59964e-01   6.27600e-02 ; 1.46  0.0150

[ moleculetype ]
;name            nrexcl
 PEN             3

[ atoms ]
;   nr  type  resi  res  atom  cgnr     charge      mass       ; qtot   bond_type
     1   c3     1   PEN     C    1    -0.080300     12.01000 ; qtot -0.080
     2   c3     1   PEN    C1    2     0.091100     12.01000 ; qtot 0.011
     3   cy     1   PEN    C2    3     0.132800     12.01000 ; qtot 0.144
     4   h1     1   PEN     H    4     0.080700      1.00800 ; qtot 0.224
     5   h2     1   PEN    H1    5     0.121700      1.00800 ; qtot 0.346
     6   ss     1   PEN     S    6    -0.436200     32.06000 ; qtot -0.090
     7   nj     1   PEN     N    7    -0.337800     14.01000 ; qtot -0.428
     8   cy     1   PEN    C3    8    -0.015300     12.01000 ; qtot -0.443
     9   h1     1   PEN    H2    9     0.112700      1.00800 ; qtot -0.331
    10    c     1   PEN    C4   10     0.650501     12.01000 ; qtot 0.320
    11    o     1   PEN     O   11    -0.572501     16.00000 ; qtot -0.253
    12    n     1   PEN    N1   12    -0.534901     14.01000 ; qtot -0.788
    13   hn     1   PEN    H3   13     0.337500      1.00800 ; qtot -0.450
    14    c     1   PEN    C5   14     0.662101     12.01000 ; qtot 0.212
    15    o     1   PEN    O1   15    -0.623101     16.00000 ; qtot -0.411
    16   c3     1   PEN    C6   16    -0.112100     12.01000 ; qtot -0.523
    17   hc     1   PEN    H4   17     0.077200      1.00800 ; qtot -0.446
    18   hc     1   PEN    H5   18     0.077200      1.00800 ; qtot -0.369
    19   ca     1   PEN    C7   19    -0.088300     12.01000 ; qtot -0.457
    20   ca     1   PEN    C8   20    -0.120000     12.01000 ; qtot -0.577
    21   ca     1   PEN    C9   21    -0.120000     12.01000 ; qtot -0.697
    22   ca     1   PEN   C10   22    -0.129000     12.01000 ; qtot -0.826
    23   ha     1   PEN    H6   23     0.140000      1.00800 ; qtot -0.686
    24   ca     1   PEN   C11   24    -0.129000     12.01000 ; qtot -0.815
    25   ha     1   PEN    H7   25     0.140000      1.00800 ; qtot -0.675
    26   ca     1   PEN   C12   26    -0.131000     12.01000 ; qtot -0.806
    27   ha     1   PEN    H8   27     0.130500      1.00800 ; qtot -0.676
    28   ha     1   PEN    H9   28     0.130500      1.00800 ; qtot -0.545
    29   ha     1   PEN   H10   29     0.127000      1.00800 ; qtot -0.418
    30    c     1   PEN   C13   30     0.940604     12.01000 ; qtot 0.523
    31    o     1   PEN    O2   31    -0.811801     16.00000 ; qtot -0.289
    32    o     1   PEN    O3   32    -0.811801     16.00000 ; qtot -1.101
    33   c3     1   PEN   C14   33    -0.089100     12.01000 ; qtot -1.190
    34   hc     1   PEN   H11   34     0.046533      1.00800 ; qtot -1.144
    35   hc     1   PEN   H12   35     0.046533      1.00800 ; qtot -1.097
    36   hc     1   PEN   H13   36     0.046533      1.00800 ; qtot -1.050
    37   c3     1   PEN   C15   37    -0.089100     12.01000 ; qtot -1.140
    38   hc     1   PEN   H14   38     0.046533      1.00800 ; qtot -1.093
    39   hc     1   PEN   H15   39     0.046533      1.00800 ; qtot -1.047
    40   hc     1   PEN   H16   40     0.046533      1.00800 ; qtot -1.000

[ bonds ]
;   ai     aj funct   r             k
     1      2   1    1.5375e-01    2.5179e+05 ;      C - C1
     1      4   1    1.0969e-01    2.7665e+05 ;      C - H
     1      7   1    1.4520e-01    2.8359e+05 ;      C - N
     1     30   1    1.5241e-01    2.6192e+05 ;      C - C13
     2      6   1    1.8392e-01    1.8067e+05 ;     C1 - S
     2     33   1    1.5375e-01    2.5179e+05 ;     C1 - C14
     2     37   1    1.5375e-01    2.5179e+05 ;     C1 - C15
     3      5   1    1.0930e-01    2.8108e+05 ;     C2 - H1
     3      6   1    1.8500e-01    1.7598e+05 ;     C2 - S
     3      7   1    1.4750e-01    2.6426e+05 ;     C2 - N
     3      8   1    1.5580e-01    2.3732e+05 ;     C2 - C3
     7     10   1    1.4010e-01    3.3313e+05 ;      N - C4
     8      9   1    1.0950e-01    2.7874e+05 ;     C3 - H2
     8     10   1    1.5520e-01    2.4150e+05 ;     C3 - C4
     8     12   1    1.4330e-01    3.0091e+05 ;     C3 - N1
    10     11   1    1.2183e-01    5.3363e+05 ;     C4 - O
```

```
   12    13   1    1.0129e-01    3.3740e+05 ;     N1 - H3
   12    14   1    1.3789e-01    3.5782e+05 ;     N1 - C5
   14    15   1    1.2183e-01    5.3363e+05 ;     C5 - O1
   14    16   1    1.5241e-01    2.6192e+05 ;     C5 - C6
   16    17   1    1.0969e-01    2.7665e+05 ;     C6 - H4
   16    18   1    1.0969e-01    2.7665e+05 ;     C6 - H5
   16    19   1    1.5156e-01    2.6861e+05 ;     C6 - C7
   19    20   1    1.3984e-01    3.8585e+05 ;     C7 - C8
   19    21   1    1.3984e-01    3.8585e+05 ;     C7 - C9
   20    22   1    1.3984e-01    3.8585e+05 ;     C8 - C10
   20    23   1    1.0860e-01    2.8937e+05 ;     C8 - H6
   21    24   1    1.3984e-01    3.8585e+05 ;     C9 - C11
   21    25   1    1.0860e-01    2.8937e+05 ;     C9 - H7
   22    26   1    1.3984e-01    3.8585e+05 ;    C10 - C12
   22    27   1    1.0860e-01    2.8937e+05 ;    C10 - H8
   24    26   1    1.3984e-01    3.8585e+05 ;    C11 - C12
   24    28   1    1.0860e-01    2.8937e+05 ;    C11 - H9
   26    29   1    1.0860e-01    2.8937e+05 ;    C12 - H10
   30    31   1    1.2183e-01    5.3363e+05 ;    C13 - O2
   30    32   1    1.2183e-01    5.3363e+05 ;    C13 - O3
   33    34   1    1.0969e-01    2.7665e+05 ;    C14 - H11
   33    35   1    1.0969e-01    2.7665e+05 ;    C14 - H12
   33    36   1    1.0969e-01    2.7665e+05 ;    C14 - H13
   37    38   1    1.0969e-01    2.7665e+05 ;    C15 - H14
   37    39   1    1.0969e-01    2.7665e+05 ;    C15 - H15
   37    40   1    1.0969e-01    2.7665e+05 ;    C15 - H16

[ pairs ]
;   ai    aj   funct
    1     5     1 ;     C - H1
    1     8     1 ;     C - C3
    1    11     1 ;     C - O
    1    34     1 ;     C - H11
    1    35     1 ;     C - H12
    1    36     1 ;     C - H13
    1    38     1 ;     C - H14
    1    39     1 ;     C - H15
    1    40     1 ;     C - H16
    2     5     1 ;    C1 - H1
    2     8     1 ;    C1 - C3
    2    10     1 ;    C1 - C4
    2    31     1 ;    C1 - O2
    2    32     1 ;    C1 - O3
    3    11     1 ;    C2 - O
    3    13     1 ;    C2 - H3
    3    14     1 ;    C2 - C5
    3    33     1 ;    C2 - C14
    3    37     1 ;    C2 - C15
    4     3     1 ;    H - C2
    4     6     1 ;    H - S
    4    10     1 ;     H - C4
    4    31     1 ;     H - O2
    4    32     1 ;     H - O3
    4    33     1 ;     H - C14
    4    37     1 ;     H - C15
    5     9     1 ;    H1 - H2
    5    10     1 ;    H1 - C4
    5    12     1 ;    H1 - N1
    6     9     1 ;    S - H2
    6    10     1 ;     S - C4
    6    12     1 ;     S - N1
    6    34     1 ;     S - H11
    6    35     1 ;     S - H12
    6    36     1 ;     S - H13
    6    38     1 ;     S - H14
    6    39     1 ;     S - H15
    6    40     1 ;     S - H16
    7     9     1 ;    N - H2
    7    12     1 ;     N - N1
    7    31     1 ;     N - O2
```

```
    7    32     1 ;     N - O3
    7    33     1 ;     N - C14
    7    37     1 ;     N - C15
    8    15     1 ;    C3 - O1
    8    16     1 ;    C3 - C6
    9    11     1 ;    H2 - O
    9    13     1 ;    H2 - H3
    9    14     1 ;    H2 - C5
   10    13     1 ;    C4 - H3
   10    14     1 ;    C4 - C5
   11    12     1 ;     O - N1
   12    17     1 ;    N1 - H4
   12    18     1 ;    N1 - H5
   12    19     1 ;    N1 - C7
   13    15     1 ;    H3 - O1
   13    16     1 ;    H3 - C6
   14    20     1 ;    C5 - C8
   14    21     1 ;    C5 - C9
   15    17     1 ;    O1 - H4
   15    18     1 ;    O1 - H5
   15    19     1 ;    O1 - C7
   16    22     1 ;    C6 - C10
   16    23     1 ;    C6 - H6
   16    24     1 ;    C6 - C11
   16    25     1 ;    C6 - H7
   17    20     1 ;    H4 - C8
   17    21     1 ;    H4 - C9
   18    20     1 ;    H5 - C8
   18    21     1 ;    H5 - C9
   19    26     1 ;    C7 - C12
   19    27     1 ;    C7 - H8
   19    28     1 ;    C7 - H9
   20    24     1 ;    C8 - C11
   20    25     1 ;    C8 - H7
   20    29     1 ;    C8 - H10
   21    22     1 ;    C9 - C10
   21    23     1 ;    C9 - H6
   21    29     1 ;    C9 - H10
   22    28     1 ;   C10 - H9
   23    26     1 ;    H6 - C12
   23    27     1 ;    H6 - H8
   24    27     1 ;   C11 - H8
   25    26     1 ;    H7 - C12
   25    28     1 ;    H7 - H9
   27    29     1 ;    H8 - H10
   28    29     1 ;    H9 - H10
   30     3     1 ;   C13 - C2
   30     6     1 ;   C13 - S
   30    10     1 ;   C13 - C4
   30    33     1 ;   C13 - C14
   30    37     1 ;   C13 - C15
   33    38     1 ;   C14 - H14
   33    39     1 ;   C14 - H15
   33    40     1 ;   C14 - H16
   34    37     1 ;   H11 - C15
   35    37     1 ;   H12 - C15
   36    37     1 ;   H13 - C15

[ angles ]
;   ai    aj    ak   funct   theta        cth
    1     2     6     1    1.1027e+02    5.1296e+02 ;     C - C1     - S
    1     2    33     1    1.1151e+02    5.2635e+02 ;     C - C1     - C14
    1     2    37     1    1.1151e+02    5.2635e+02 ;     C - C1     - C15
    1     7     3     1    1.1718e+02    5.2300e+02 ;     C - N      - C2
    1     7    10     1    1.2760e+02    5.1463e+02 ;     C - N      - C4
    1    30    31     1    1.2320e+02    5.6400e+02 ;     C - C13    - O2
    1    30    32     1    1.2320e+02    5.6400e+02 ;     C - C13    - O3
    2     1     4     1    1.0956e+02    3.8828e+02 ;    C1 - C      - H
    2     1     7     1    1.0694e+02    5.6484e+02 ;    C1 - C      - N
    2     1    30     1    1.1104e+02    5.2969e+02 ;    C1 - C      - C13
```

```
    2     6     3     1   9.4240e+01    5.1547e+02 ;     C1 - S      - C2
    2    33    34     1   1.0980e+02    3.8744e+02 ;     C1 - C14    - H11
    2    33    35     1   1.0980e+02    3.8744e+02 ;     C1 - C14    - H12
    2    33    36     1   1.0980e+02    3.8744e+02 ;     C1 - C14    - H13
    2    37    38     1   1.0980e+02    3.8744e+02 ;     C1 - C15    - H14
    2    37    39     1   1.0980e+02    3.8744e+02 ;     C1 - C15    - H15
    2    37    40     1   1.0980e+02    3.8744e+02 ;     C1 - C15    - H16
    3     7    10     1   9.4020e+01    5.9413e+02 ;     C2 - N      - C4
    3     8     9     1   1.1317e+02    3.7740e+02 ;     C2 - C3     - H2
    3     8    10     1   8.5160e+01    5.9496e+02 ;     C2 - C3     - C4
    3     8    12     1   1.1987e+02    5.3220e+02 ;     C2 - C3     - N1
    4     1     7     1   1.0852e+02    4.2007e+02 ;     H - C       - N
    4     1    30     1   1.0822e+02    3.9330e+02 ;     H - C       - C13
    5     3     6     1   1.0966e+02    3.4811e+02 ;     H1 - C2     - S
    5     3     7     1   1.1454e+02    4.0334e+02 ;     H1 - C2     - N
    5     3     8     1   1.1679e+02    3.7154e+02 ;     H1 - C2     - C3
    6     2    33     1   1.1027e+02    5.1296e+02 ;     S - C1      - C14
    6     2    37     1   1.1027e+02    5.1296e+02 ;     S - C1      - C15
    6     3     7     1   1.0514e+02    5.4141e+02 ;     S - C2      - N
    6     3     8     1   1.1827e+02    4.9120e+02 ;     S - C2      - C3
    7     1    30     1   1.1019e+02    5.5982e+02 ;     N - C       - C13
    7     3     8     1   8.8640e+01    6.1170e+02 ;     N - C2      - C3
    7    10     8     1   9.1680e+01    6.1588e+02 ;     N - C4      - C3
    7    10    11     1   1.3306e+02    5.9078e+02 ;     N - C4      - O
    8    10    11     1   1.3523e+02    5.3053e+02 ;     C3 - C4     - O
    8    12    13     1   1.1885e+02    3.8828e+02 ;     C3 - N1     - H3
    8    12    14     1   1.2171e+02    5.3388e+02 ;     C3 - N1     - C5
    9     8    10     1   1.1297e+02    3.7907e+02 ;     H2 - C3     - C4
    9     8    12     1   1.0799e+02    4.2593e+02 ;     H2 - C3     - N1
   10     8    12     1   1.1738e+02    5.3974e+02 ;     C4 - C3     - N1
   12    14    15     1   1.2305e+02    6.2091e+02 ;     N1 - C5     - O1
   12    14    16     1   1.1518e+02    5.5898e+02 ;     N1 - C5     - C6
   13    12    14     1   1.1755e+02    4.0417e+02 ;     H3 - N1     - C5
   14    16    17     1   1.0877e+02    3.9246e+02 ;     C5 - C6     - H4
   14    16    18     1   1.0877e+02    3.9246e+02 ;     C5 - C6     - H5
   14    16    19     1   1.1101e+02    5.3304e+02 ;     C5 - C6     - C7
   15    14    16     1   1.2320e+02    5.6400e+02 ;     O1 - C5     - C6
   16    19    20     1   1.2077e+02    5.3137e+02 ;     C6 - C7     - C8
   16    19    21     1   1.2077e+02    5.3137e+02 ;     C6 - C7     - C9
   17    16    18     1   1.0758e+02    3.2970e+02 ;     H4 - C6     - H5
   17    16    19     1   1.1047e+02    3.9162e+02 ;     H4 - C6     - C7
   18    16    19     1   1.1047e+02    3.9162e+02 ;     H5 - C6     - C7
   19    20    22     1   1.2002e+02    5.5731e+02 ;     C7 - C8     - C10
   19    20    23     1   1.1988e+02    4.0334e+02 ;     C7 - C8     - H6
   19    21    24     1   1.2002e+02    5.5731e+02 ;     C7 - C9     - C11
   19    21    25     1   1.1988e+02    4.0334e+02 ;     C7 - C9     - H7
   20    19    21     1   1.2002e+02    5.5731e+02 ;     C8 - C7     - C9
   20    22    26     1   1.2002e+02    5.5731e+02 ;     C8 - C10    - C12
   20    22    27     1   1.1988e+02    4.0334e+02 ;     C8 - C10    - H8
   21    24    26     1   1.2002e+02    5.5731e+02 ;     C9 - C11    - C12
   21    24    28     1   1.1988e+02    4.0334e+02 ;     C9 - C11    - H9
   22    20    23     1   1.1988e+02    4.0334e+02 ;     C10 - C8    - H6
   22    26    24     1   1.2002e+02    5.5731e+02 ;     C10 - C12   - C11
   22    26    29     1   1.1988e+02    4.0334e+02 ;     C10 - C12   - H10
   24    21    25     1   1.1988e+02    4.0334e+02 ;     C11 - C9    - H7
   24    26    29     1   1.1988e+02    4.0334e+02 ;     C11 - C12   - H10
   26    22    27     1   1.1988e+02    4.0334e+02 ;     C12 - C10   - H8
   26    24    28     1   1.1988e+02    4.0334e+02 ;     C12 - C11   - H9
   31    30    32     1   1.3025e+02    6.5187e+02 ;     O2 - C13    - O3
   33     2    37     1   1.1151e+02    5.2635e+02 ;     C14 - C1    - C15
   34    33    35     1   1.0758e+02    3.2970e+02 ;     H11 - C14   - H12
   34    33    36     1   1.0758e+02    3.2970e+02 ;     H11 - C14   - H13
   35    33    36     1   1.0758e+02    3.2970e+02 ;     H12 - C14   - H13
   38    37    39     1   1.0758e+02    3.2970e+02 ;     H14 - C15   - H15
   38    37    40     1   1.0758e+02    3.2970e+02 ;     H14 - C15   - H16
   39    37    40     1   1.0758e+02    3.2970e+02 ;     H15 - C15   - H16

[ dihedrals ] ; propers
; treated as RBs in GROMACS to use combine multiple AMBER torsions per quartet
;    i      j      k      l   func     C0        C1        C2        C3        C4        C5
```

| | | | | | | | | | | | | | | | |
|---|---|---|---|---|---|---|---|---|---|---|---|---|---|---|---|
| 1 | 2 | 6 | 3 | 3 | 1.39467 | 4.18400 | 0.00000 | -5.57867 | 0.00000 | 0.00000 | ; | C- | C1- | S- | C2 |
| 1 | 2 | 33 | 34 | 3 | 0.66944 | 2.00832 | 0.00000 | -2.67776 | 0.00000 | 0.00000 | ; | C- | C1- | C14- | H11 |
| 1 | 2 | 33 | 35 | 3 | 0.66944 | 2.00832 | 0.00000 | -2.67776 | 0.00000 | 0.00000 | ; | C- | C1- | C14- | H12 |
| 1 | 2 | 33 | 36 | 3 | 0.66944 | 2.00832 | 0.00000 | -2.67776 | 0.00000 | 0.00000 | ; | C- | C1- | C14- | H13 |
| 1 | 2 | 37 | 38 | 3 | 0.66944 | 2.00832 | 0.00000 | -2.67776 | 0.00000 | 0.00000 | ; | C- | C1- | C15- | H14 |
| 1 | 2 | 37 | 39 | 3 | 0.66944 | 2.00832 | 0.00000 | -2.67776 | 0.00000 | 0.00000 | ; | C- | C1- | C15- | H15 |
| 1 | 2 | 37 | 40 | 3 | 0.66944 | 2.00832 | 0.00000 | -2.67776 | 0.00000 | 0.00000 | ; | C- | C1- | C15- | H16 |
| 1 | 7 | 3 | 5 | 3 | 0.00000 | 0.00000 | 0.00000 | 0.00000 | 0.00000 | 0.00000 | ; | C- | N- | C2- | H1 |
| 1 | 7 | 3 | 6 | 3 | 0.00000 | 0.00000 | 0.00000 | 0.00000 | 0.00000 | 0.00000 | ; | C- | N- | C2- | S |
| 1 | 7 | 3 | 8 | 3 | 0.00000 | 0.00000 | 0.00000 | 0.00000 | 0.00000 | 0.00000 | ; | C- | N- | C2- | C3 |
| 1 | 7 | 10 | 8 | 3 | 20.92000 | 0.00000 | -20.92000 | 0.00000 | 0.00000 | 0.00000 | ; | C- | N- | C4- | C3 |
| 1 | 7 | 10 | 11 | 3 | 20.92000 | 0.00000 | -20.92000 | 0.00000 | 0.00000 | 0.00000 | ; | C- | N- | C4- | O |
| 2 | 1 | 7 | 3 | 3 | 0.00000 | 0.00000 | 0.00000 | 0.00000 | 0.00000 | 0.00000 | ; | C1- | C- | N- | C2 |
| 2 | 1 | 7 | 10 | 3 | 2.84512 | -4.10032 | 16.73600 | 2.51040 | -16.73600 | 0.00000 | ; | C1- | C- | N- | C4 |
| 2 | 1 | 30 | 31 | 3 | 0.00000 | 0.00000 | 0.00000 | 0.00000 | 0.00000 | 0.00000 | ; | C1- | C- | C13- | O2 |
| 2 | 1 | 30 | 32 | 3 | 0.00000 | 0.00000 | 0.00000 | 0.00000 | 0.00000 | 0.00000 | ; | C1- | C- | C13- | O3 |
| 2 | 6 | 3 | 5 | 3 | 1.39467 | 4.18400 | 0.00000 | -5.57867 | 0.00000 | 0.00000 | ; | C1- | S- | C2- | H1 |
| 2 | 6 | 3 | 7 | 3 | 1.39467 | 4.18400 | 0.00000 | -5.57867 | 0.00000 | 0.00000 | ; | C1- | S- | C2- | N |
| 2 | 6 | 3 | 8 | 3 | 1.39467 | 4.18400 | 0.00000 | -5.57867 | 0.00000 | 0.00000 | ; | C1- | S- | C2- | C3 |
| 3 | 6 | 2 | 33 | 3 | 1.39467 | 4.18400 | 0.00000 | -5.57867 | 0.00000 | 0.00000 | ; | C2- | S- | C1- | C14 |
| 3 | 6 | 2 | 37 | 3 | 1.39467 | 4.18400 | 0.00000 | -5.57867 | 0.00000 | 0.00000 | ; | C2- | S- | C1- | C15 |
| 3 | 7 | 10 | 8 | 3 | 20.92000 | 0.00000 | -20.92000 | 0.00000 | 0.00000 | 0.00000 | ; | C2- | N- | C4- | C3 |
| 3 | 7 | 10 | 11 | 3 | 20.92000 | 0.00000 | -20.92000 | 0.00000 | 0.00000 | 0.00000 | ; | C2- | N- | C4- | O |
| 3 | 8 | 10 | 7 | 3 | 0.00000 | 0.00000 | 0.00000 | 0.00000 | 0.00000 | 0.00000 | ; | C2- | C3- | C4- | N |
| 3 | 8 | 10 | 11 | 3 | 0.00000 | 0.00000 | 0.00000 | 0.00000 | 0.00000 | 0.00000 | ; | C2- | C3- | C4- | O |
| 3 | 8 | 12 | 13 | 3 | 0.00000 | 0.00000 | 0.00000 | 0.00000 | 0.00000 | 0.00000 | ; | C2- | C3- | N1- | H3 |
| 3 | 8 | 12 | 14 | 3 | 0.00000 | 0.00000 | 0.00000 | 0.00000 | 0.00000 | 0.00000 | ; | C2- | C3- | N1- | C5 |
| 4 | 1 | 2 | 6 | 3 | 0.65084 | 1.95253 | 0.00000 | -2.60338 | 0.00000 | 0.00000 | ; | H- | C- | C1- | S |
| 4 | 1 | 2 | 33 | 3 | 0.65084 | 1.95253 | 0.00000 | -2.60338 | 0.00000 | 0.00000 | ; | H- | C- | C1- | C14 |
| 4 | 1 | 2 | 37 | 3 | 0.65084 | 1.95253 | 0.00000 | -2.60338 | 0.00000 | 0.00000 | ; | H- | C- | C1- | C15 |
| 4 | 1 | 7 | 3 | 3 | 0.00000 | 0.00000 | 0.00000 | 0.00000 | 0.00000 | 0.00000 | ; | H- | C- | N- | C2 |
| 4 | 1 | 7 | 10 | 3 | 0.00000 | 0.00000 | 0.00000 | 0.00000 | 0.00000 | 0.00000 | ; | H- | C- | N- | C4 |
| 4 | 1 | 30 | 31 | 3 | 3.68192 | -4.35136 | 0.00000 | 1.33888 | 0.00000 | 0.00000 | ; | H- | C- | C13- | O2 |
| 4 | 1 | 30 | 32 | 3 | 3.68192 | -4.35136 | 0.00000 | 1.33888 | 0.00000 | 0.00000 | ; | H- | C- | C13- | O3 |
| 5 | 3 | 7 | 10 | 3 | 0.00000 | 0.00000 | 0.00000 | 0.00000 | 0.00000 | 0.00000 | ; | H1- | C2- | N- | C4 |
| 5 | 3 | 8 | 9 | 3 | 0.65084 | 1.95253 | 0.00000 | -2.60338 | 0.00000 | 0.00000 | ; | H1- | C2- | C3- | H2 |
| 5 | 3 | 8 | 10 | 3 | 0.65084 | 1.95253 | 0.00000 | -2.60338 | 0.00000 | 0.00000 | ; | H1- | C2- | C3- | C4 |
| 5 | 3 | 8 | 12 | 3 | 0.65084 | 1.95253 | 0.00000 | -2.60338 | 0.00000 | 0.00000 | ; | H1- | C2- | C3- | N1 |
| 6 | 2 | 33 | 34 | 3 | 0.65084 | 1.95253 | 0.00000 | -2.60338 | 0.00000 | 0.00000 | ; | S- | C1- | C14- | H11 |
| 6 | 2 | 33 | 35 | 3 | 0.65084 | 1.95253 | 0.00000 | -2.60338 | 0.00000 | 0.00000 | ; | S- | C1- | C14- | H12 |
| 6 | 2 | 33 | 36 | 3 | 0.65084 | 1.95253 | 0.00000 | -2.60338 | 0.00000 | 0.00000 | ; | S- | C1- | C14- | H13 |
| 6 | 2 | 37 | 38 | 3 | 0.65084 | 1.95253 | 0.00000 | -2.60338 | 0.00000 | 0.00000 | ; | S- | C1- | C15- | H14 |
| 6 | 2 | 37 | 39 | 3 | 0.65084 | 1.95253 | 0.00000 | -2.60338 | 0.00000 | 0.00000 | ; | S- | C1- | C15- | H15 |
| 6 | 2 | 37 | 40 | 3 | 0.65084 | 1.95253 | 0.00000 | -2.60338 | 0.00000 | 0.00000 | ; | S- | C1- | C15- | H16 |
| 6 | 3 | 7 | 10 | 3 | 0.00000 | 0.00000 | 0.00000 | 0.00000 | 0.00000 | 0.00000 | ; | S- | C2- | N- | C4 |
| 6 | 3 | 8 | 9 | 3 | 0.65084 | 1.95253 | 0.00000 | -2.60338 | 0.00000 | 0.00000 | ; | S- | C2- | C3- | H2 |
| 6 | 3 | 8 | 10 | 3 | 0.65084 | 1.95253 | 0.00000 | -2.60338 | 0.00000 | 0.00000 | ; | S- | C2- | C3- | C4 |
| 6 | 3 | 8 | 12 | 3 | 0.65084 | 1.95253 | 0.00000 | -2.60338 | 0.00000 | 0.00000 | ; | S- | C2- | C3- | N1 |
| 7 | 1 | 2 | 6 | 3 | 0.65084 | 1.95253 | 0.00000 | -2.60338 | 0.00000 | 0.00000 | ; | N- | C- | C1- | S |
| 7 | 1 | 2 | 33 | 3 | 0.65084 | 1.95253 | 0.00000 | -2.60338 | 0.00000 | 0.00000 | ; | N- | C- | C1- | C14 |
| 7 | 1 | 2 | 37 | 3 | 0.65084 | 1.95253 | 0.00000 | -2.60338 | 0.00000 | 0.00000 | ; | N- | C- | C1- | C15 |
| 7 | 1 | 30 | 31 | 3 | 0.00000 | 0.00000 | 0.00000 | 0.00000 | 0.00000 | 0.00000 | ; | N- | C- | C13- | O2 |
| 7 | 1 | 30 | 32 | 3 | 0.00000 | 0.00000 | 0.00000 | 0.00000 | 0.00000 | 0.00000 | ; | N- | C- | C13- | O3 |
| 7 | 3 | 8 | 9 | 3 | 0.65084 | 1.95253 | 0.00000 | -2.60338 | 0.00000 | 0.00000 | ; | N- | C2- | C3- | H2 |
| 7 | 3 | 8 | 10 | 3 | 0.65084 | 1.95253 | 0.00000 | -2.60338 | 0.00000 | 0.00000 | ; | N- | C2- | C3- | C4 |
| 7 | 3 | 8 | 12 | 3 | 0.65084 | 1.95253 | 0.00000 | -2.60338 | 0.00000 | 0.00000 | ; | N- | C2- | C3- | N1 |
| 7 | 10 | 8 | 9 | 3 | 0.00000 | 0.00000 | 0.00000 | 0.00000 | 0.00000 | 0.00000 | ; | N- | C4- | C3- | H2 |
| 7 | 10 | 8 | 12 | 3 | 0.00000 | 0.00000 | 0.00000 | 0.00000 | 0.00000 | 0.00000 | ; | N- | C4- | C3- | N1 |
| 8 | 3 | 7 | 10 | 3 | 0.00000 | 0.00000 | 0.00000 | 0.00000 | 0.00000 | 0.00000 | ; | C3- | C2- | N- | C4 |
| 8 | 12 | 14 | 15 | 3 | 20.92000 | 0.00000 | -20.92000 | 0.00000 | 0.00000 | 0.00000 | ; | C3- | N1- | C5- | O1 |
| 8 | 12 | 14 | 16 | 3 | 20.92000 | 0.00000 | -20.92000 | 0.00000 | 0.00000 | 0.00000 | ; | C3- | N1- | C5- | C6 |
| 9 | 8 | 10 | 11 | 3 | 0.00000 | 0.00000 | 0.00000 | 0.00000 | 0.00000 | 0.00000 | ; | H2- | C3- | C4- | O |
| 9 | 8 | 12 | 13 | 3 | 0.00000 | 0.00000 | 0.00000 | 0.00000 | 0.00000 | 0.00000 | ; | H2- | C3- | N1- | H3 |
| 9 | 8 | 12 | 14 | 3 | 0.00000 | 0.00000 | 0.00000 | 0.00000 | 0.00000 | 0.00000 | ; | H2- | C3- | N1- | C5 |
| 10 | 8 | 12 | 13 | 3 | 0.00000 | 0.00000 | 0.00000 | 0.00000 | 0.00000 | 0.00000 | ; | C4- | C3- | N1- | H3 |
| 10 | 8 | 12 | 14 | 3 | 0.00000 | 0.00000 | 0.00000 | 0.00000 | 0.00000 | 0.00000 | ; | C4- | C3- | N1- | C5 |
| 11 | 10 | 8 | 12 | 3 | 0.00000 | 0.00000 | 0.00000 | 0.00000 | 0.00000 | 0.00000 | ; | O- | C4- | C3- | N1 |
| 12 | 14 | 16 | 17 | 3 | 0.00000 | 0.00000 | 0.00000 | 0.00000 | 0.00000 | 0.00000 | ; | N1- | C5- | C6- | H4 |
| 12 | 14 | 16 | 18 | 3 | 0.00000 | 0.00000 | 0.00000 | 0.00000 | 0.00000 | 0.00000 | ; | N1- | C5- | C6- | H5 |
| 12 | 14 | 16 | 19 | 3 | 0.00000 | 0.00000 | 0.00000 | 0.00000 | 0.00000 | 0.00000 | ; | N1- | C5- | C6- | C7 |

```
13 12 14 15 3 29.28800 -8.36800 -20.92000 0.00000 0.00000 0.00000 ; H3- N1- C5- O1
13 12 14 16 3 20.92000 0.00000 -20.92000 0.00000 0.00000 0.00000 ; H3- N1- C5- C6
14 16 19 20 3 0.00000 0.00000 0.00000 0.00000 0.00000 0.00000 ; C5- C6- C7- C8
14 16 19 21 3 0.00000 0.00000 0.00000 0.00000 0.00000 0.00000 ; C5- C6- C7- C9
15 14 16 17 3 3.68192 -4.35136 0.00000 1.33888 0.00000 0.00000 ; O1- C5- C6- H4
15 14 16 18 3 3.68192 -4.35136 0.00000 1.33888 0.00000 0.00000 ; O1- C5- C6- H5
15 14 16 19 3 0.00000 0.00000 0.00000 0.00000 0.00000 0.00000 ; O1- C5- C6- C7
16 19 20 22 3 30.33400 0.00000 -30.33400 0.00000 0.00000 0.00000 ; C6- C7- C8- C10
16 19 20 23 3 30.33400 0.00000 -30.33400 0.00000 0.00000 0.00000 ; C6- C7- C8- H6
16 19 21 24 3 30.33400 0.00000 -30.33400 0.00000 0.00000 0.00000 ; C6- C7- C9- C11
16 19 21 25 3 30.33400 0.00000 -30.33400 0.00000 0.00000 0.00000 ; C6- C7- C9- H7
17 16 19 20 3 0.00000 0.00000 0.00000 0.00000 0.00000 0.00000 ; H4- C6- C7- C8
17 16 19 21 3 0.00000 0.00000 0.00000 0.00000 0.00000 0.00000 ; H4- C6- C7- C9
18 16 19 20 3 0.00000 0.00000 0.00000 0.00000 0.00000 0.00000 ; H5- C6- C7- C8
18 16 19 21 3 0.00000 0.00000 0.00000 0.00000 0.00000 0.00000 ; H5- C6- C7- C9
19 20 22 26 3 30.33400 0.00000 -30.33400 0.00000 0.00000 0.00000 ; C7- C8- C10- C12
19 20 22 27 3 30.33400 0.00000 -30.33400 0.00000 0.00000 0.00000 ; C7- C8- C10- H8
19 21 24 26 3 30.33400 0.00000 -30.33400 0.00000 0.00000 0.00000 ; C7- C9- C11- C12
19 21 24 28 3 30.33400 0.00000 -30.33400 0.00000 0.00000 0.00000 ; C7- C9- C11- H9
20 19 21 24 3 30.33400 0.00000 -30.33400 0.00000 0.00000 0.00000 ; C8- C7- C9- C11
20 19 21 25 3 30.33400 0.00000 -30.33400 0.00000 0.00000 0.00000 ; C8- C7- C9- H7
20 22 26 24 3 30.33400 0.00000 -30.33400 0.00000 0.00000 0.00000 ; C8- C10- C12- C11
20 22 26 29 3 30.33400 0.00000 -30.33400 0.00000 0.00000 0.00000 ; C8- C10- C12- H10
21 19 20 22 3 30.33400 0.00000 -30.33400 0.00000 0.00000 0.00000 ; C9- C7- C8- C10
21 19 20 23 3 30.33400 0.00000 -30.33400 0.00000 0.00000 0.00000 ; C9- C7- C8- H6
21 24 26 22 3 30.33400 0.00000 -30.33400 0.00000 0.00000 0.00000 ; C9- C11- C12- C10
21 24 26 29 3 30.33400 0.00000 -30.33400 0.00000 0.00000 0.00000 ; C9- C11- C12- H10
22 26 24 28 3 30.33400 0.00000 -30.33400 0.00000 0.00000 0.00000 ; C10- C12- C11- H9
23 20 22 26 3 30.33400 0.00000 -30.33400 0.00000 0.00000 0.00000 ; H6- C8- C10- C12
23 20 22 27 3 30.33400 0.00000 -30.33400 0.00000 0.00000 0.00000 ; H6- C8- C10- H8
24 26 22 27 3 30.33400 0.00000 -30.33400 0.00000 0.00000 0.00000 ; C11- C12- C10- H8
25 21 24 26 3 30.33400 0.00000 -30.33400 0.00000 0.00000 0.00000 ; H7- C9- C11- C12
25 21 24 28 3 30.33400 0.00000 -30.33400 0.00000 0.00000 0.00000 ; H7- C9- C11- H9
27 22 26 29 3 30.33400 0.00000 -30.33400 0.00000 0.00000 0.00000 ; H8- C10- C12- H10
28 24 26 29 3 30.33400 0.00000 -30.33400 0.00000 0.00000 0.00000 ; H9- C11- C12- H10
30 1 2 6 3 0.65084 1.95253 0.00000 -2.60338 0.00000 0.00000 ; C13- C- C1- S
30 1 2 33 3 0.65084 1.95253 0.00000 -2.60338 0.00000 0.00000 ; C13- C- C1- C14
30 1 2 37 3 0.65084 1.95253 0.00000 -2.60338 0.00000 0.00000 ; C13- C- C1- C15
30 1 7 3 3 0.00000 0.00000 0.00000 0.00000 0.00000 0.00000 ; C13- C- N- C2
30 1 7 10 3 10.46000 -3.34720 -7.11280 0.00000 0.00000 0.00000 ; C13- C- N- C4
33 2 37 38 3 0.66944 2.00832 0.00000 -2.67776 0.00000 0.00000 ; C14- C1- C15- H14
33 2 37 39 3 0.66944 2.00832 0.00000 -2.67776 0.00000 0.00000 ; C14- C1- C15- H15
33 2 37 40 3 0.66944 2.00832 0.00000 -2.67776 0.00000 0.00000 ; C14- C1- C15- H16
34 33 2 37 3 0.66944 2.00832 0.00000 -2.67776 0.00000 0.00000 ; H11- C14- C1- C15
35 33 2 37 3 0.66944 2.00832 0.00000 -2.67776 0.00000 0.00000 ; H12- C14- C1- C15
36 33 2 37 3 0.66944 2.00832 0.00000 -2.67776 0.00000 0.00000 ; H13- C14- C1- C15

[ dihedrals ] ; impropers
; treated as propers in GROMACS to use correct AMBER analytical function
; i j k l func phase kd pn
1 31 30 32 1 180.00 4.60240 2 ; C- O2- C13- O3
8 7 10 11 1 180.00 43.93200 2 ; C3- N- C4- O
10 1 7 3 1 180.00 4.60240 2 ; C4- C- N- C2
14 8 12 13 1 180.00 4.60240 2 ; C5- C3- N1- H3
16 12 14 15 1 180.00 43.93200 2 ; C6- N1- C5- O1
19 22 20 23 1 180.00 4.60240 2 ; C7- C10- C8- H6
19 24 21 25 1 180.00 4.60240 2 ; C7- C11- C9- H7
20 21 19 16 1 180.00 4.60240 2 ; C8- C9- C7- C6
20 26 22 27 1 180.00 4.60240 2 ; C8- C12- C10- H8
21 26 24 28 1 180.00 4.60240 2 ; C9- C12- C11- H9
22 24 26 29 1 180.00 4.60240 2 ; C10- C11- C12- H10
```

#### SPG.gro

```
   70SPG      N  710   3.568   3.490   3.095
   70SPG      H  711   3.650   3.456   3.141
   70SPG     CA  712   3.582   3.552   2.964
   70SPG     HA  713   3.536   3.639   2.981
   70SPG     CB  714   3.727   3.577   2.925
   70SPG    HB1  715   3.733   3.624   2.837
   70SPG    HB2  716   3.778   3.491   2.921
   70SPG     OG  717   3.793   3.659   3.022
   70SPG     S1  718   4.192   3.726   3.176
   70SPG     C1  719   4.262   3.551   3.168
   70SPG     C2  720   4.142   3.464   3.118
   70SPG     N1  721   4.083   3.544   3.004
   70SPG     C3  722   4.059   3.687   3.055
   70SPG     C4  723   3.913   3.713   3.119
   70SPG     C5  724   3.803   3.658   3.022
   70SPG     O1  725   3.800   3.538   2.979
   70SPG     C6  726   4.378   3.549   3.063
   70SPG     C7  727   4.311   3.487   3.301
   70SPG     C8  728   4.189   3.302   3.085
   70SPG     O2  729   4.138   3.248   2.982
   70SPG     O3  730   4.269   3.255   3.178
   70SPG     N2  731   3.893   3.652   3.255
   70SPG     C9  732   3.904   3.745   3.366
   70SPG     O4  733   3.885   3.868   3.352
   70SPG    C10  734   3.934   3.671   3.502
   70SPG    C11  735   3.860   3.730   3.624
   70SPG    C12  736   3.830   3.649   3.736
   70SPG    C13  737   3.767   3.704   3.849
   70SPG    C14  738   3.732   3.841   3.852
   70SPG    C15  739   3.760   3.922   3.740
   70SPG    C16  740   3.824   3.867   3.627
   70SPG     H1  741   4.070   3.456   3.203
   70SPG     H2  742   3.986   3.505   2.982
   70SPG     H3  743   4.063   3.756   2.968
   70SPG     H4  744   3.896   3.822   3.133
   70SPG     H5  745   4.341   3.578   2.964
   70SPG     H6  746   4.459   3.618   3.092
   70SPG     H7  747   4.419   3.447   3.057
   70SPG     H8  748   4.411   3.522   3.330
   70SPG     H9  749   3.909   3.564   3.492
   70SPG    H10  750   4.043   3.677   3.519
   70SPG    H11  751   3.857   3.543   3.735
   70SPG    H12  752   3.744   3.640   3.936
   70SPG    H13  753   3.682   3.883   3.940
   70SPG    H14  754   3.732   4.028   3.741
   70SPG    H15  755   3.846   3.925   3.536
   70SPG    H16  756   3.949   3.564   3.267
   70SPG    H17  757   4.240   3.507   3.383
   70SPG    H18  758   4.306   3.375   3.268
   70SPG      C  759   3.505   3.484   2.850
   70SPG      O  760   3.494   3.539   2.741
```

#### SPG.top

```
[ defaults ]
; nbfunc      comb-rule      gen-pairs      fudgeLJ fudgeQQ
1             2              yes            0.5     0.8333
```

```
[ atomtypes ]
;name   bond_type     mass     charge   ptype   sigma         epsilon       Amb
 n       n          0.00000  0.00000   A     3.25000e-01   7.11280e-01 ; 1.82  0.1700
 hn      hn         0.00000  0.00000   A     1.06908e-01   6.56888e-02 ; 0.60  0.0157
 c3      c3         0.00000  0.00000   A     3.39967e-01   4.57730e-01 ; 1.91  0.1094
 h1      h1         0.00000  0.00000   A     2.47135e-01   6.56888e-02 ; 1.39  0.0157
 os      os         0.00000  0.00000   A     3.00001e-01   7.11280e-01 ; 1.68  0.1700
 ss      ss         0.00000  0.00000   A     3.56359e-01   1.04600e+00 ; 2.00  0.2500
 n3      n3         0.00000  0.00000   A     3.25000e-01   7.11280e-01 ; 1.82  0.1700
 c       c          0.00000  0.00000   A     3.39967e-01   3.59824e-01 ; 1.91  0.0860
 o       o          0.00000  0.00000   A     2.95992e-01   8.78640e-01 ; 1.66  0.2100
 ca      ca         0.00000  0.00000   A     3.39967e-01   3.59824e-01 ; 1.91  0.0860
 h2      h2         0.00000  0.00000   A     2.29317e-01   6.56888e-02 ; 1.29  0.0157
 hc      hc         0.00000  0.00000   A     2.64953e-01   6.56888e-02 ; 1.49  0.0157
 ha      ha         0.00000  0.00000   A     2.59964e-01   6.27600e-02 ; 1.46  0.0150

[ moleculetype ]
;name            nrexcl
 SPG              3

[ atoms ]
;   nr  type  resi  res  atom  cgnr     charge      mass       ; qtot   bond_type
710     N       70      SPG     N       1       -0.4157         14.01
711     H       70      SPG     H       2       0.2719          1.008
712     CT      70      SPG     CA      3       -0.0349         12.01
713     H1      70      SPG     HA      4       0.0853          1.008
714     CT      70      SPG     CB      5       0.1334          12.01
715     H1      70      SPG     HB1     6       0.0897          1.008
716     H1      70      SPG     HB2     7       0.0897          1.008
717     os      70      SPG     OG      8       -0.4439         16
718     ss      70      SPG     S1      9       -0.4702         32.06
719     c3      70      SPG     C1      10      0.0821          12.01
720     c3      70      SPG     C2      11      0.0145          12.01
721     n3      70      SPG     N1      12      -0.796201       14.01
722     c3      70      SPG     C3      13      0.2936          12.01
723     c3      70      SPG     C4      14      0.0877          12.01
724     c       70      SPG     C5      15      0.620101 12.01
725     o       70      SPG     O1      16      -0.554001       16
726     c3      70      SPG     C6      17      -0.0811         12.01
727     c3      70      SPG     C7      18      -0.0811         12.01
728     c       70      SPG     C8      19      0.935602 12.01
729     o       70      SPG     O2      20      -0.821301       16
730     o       70      SPG     O3      21      -0.821301       16
731     n       70      SPG     N2      22      -0.531901       14.01
732     c       70      SPG     C9      23      0.672101 12.01
733     o       70      SPG     O4      24      -0.660101       16
734     c3      70      SPG     C10     25      -0.0971   12.01
735     ca      70      SPG     C11     26      -0.1133   12.01
736     ca      70      SPG     C12     27      -0.133    12.01
737     ca      70      SPG     C13     28      -0.1215   12.01
738     ca      70      SPG     C14     29      -0.124    12.01
739     ca      70      SPG     C15     30      -0.1215   12.01
740     ca      70      SPG     C16     31      -0.133    12.01
741     h1      70      SPG     H1      32      0.0577    1.008
742     hn      70      SPG     H2      33      0.4038    1.008
743     h2      70      SPG     H3      34      0.0857    1.008
744     h1      70      SPG     H4      35      0.1137    1.008
745     hc      70      SPG     H5      36      0.0452    1.008
746     hc      70      SPG     H6      37      0.0452    1.008
747     hc      70      SPG     H7      38      0.0452    1.008
748     hc      70      SPG     H8      39      0.0452    1.008
749     hc      70      SPG     H9      40      0.0747    1.008
750     hc      70      SPG     H10     41      0.0747    1.008
751     ha      70      SPG     H11     42      0.132     1.008
752     ha      70      SPG     H12     43      0.1575    1.008
753     ha      70      SPG     H13     44      0.134     1.008
754     ha      70      SPG     H14     45      0.1575    1.008
755     ha      70      SPG     H15     46      0.132     1.008
756     hn      70      SPG     H16     47      0.3555    1.008
757     hc      70      SPG     H17     48      0.0452    1.008
```

| | | | | | | | |
|---|---|---|---|---|---|---|---|
| 758 | hc | 70 | SPG | H18 | 49 | 0.0452 | 1.008 |
| 759 | C | 70 | SPG | C | 50 | 0.5973 | 12.01 |
| 760 | O | 70 | SPG | O | 51 | -0.5679 | 16 |

[ bonds ]

; ai aj funct r k

| | | | | | | | | |
|---|---|---|---|---|---|---|---|---|
| 714 | 717 | 1 | 1.4316E-01 | 2.5824E+05 | ; | CB | - | OG |
| 717 | 724 | 1 | 1.3584E-01 | 3.2702E+05 | ; | OG | - | C5 |
| 718 | 719 | 1 | 1.8392E-01 | 1.8067E+05 | ; | S1 | - | C1 |
| 718 | 722 | 1 | 1.8392E-01 | 1.8067E+05 | ; | S1 | - | C3 |
| 719 | 720 | 1 | 1.5375E-01 | 2.5179E+05 | ; | C1 | - | C2 |
| 719 | 726 | 1 | 1.5375E-01 | 2.5179E+05 | ; | C1 | - | C6 |
| 719 | 727 | 1 | 1.5375E-01 | 2.5179E+05 | ; | C1 | - | C7 |
| 720 | 721 | 1 | 1.4647E-01 | 2.7271E+05 | ; | C2 | - | N1 |
| 720 | 728 | 1 | 1.5241E-01 | 2.6192E+05 | ; | C2 | - | C8 |
| 720 | 741 | 1 | 1.0969E-01 | 2.7665E+05 | ; | C2 | - | H1 |
| 721 | 722 | 1 | 1.4647E-01 | 2.7271E+05 | ; | N1 | - | C3 |
| 721 | 742 | 1 | 1.0190E-01 | 3.2836E+05 | ; | N1 | - | H2 |
| 722 | 723 | 1 | 1.5375E-01 | 2.5179E+05 | ; | C3 | - | C4 |
| 722 | 743 | 1 | 1.0961E-01 | 2.7757E+05 | ; | C3 | - | H3 |
| 723 | 724 | 1 | 1.5241E-01 | 2.6192E+05 | ; | C4 | - | C5 |
| 723 | 731 | 1 | 1.4619E-01 | 2.7506E+05 | ; | C4 | - | N2 |
| 723 | 744 | 1 | 1.0969E-01 | 2.7665E+05 | ; | C4 | - | H4 |
| 724 | 725 | 1 | 1.2183E-01 | 5.3363E+05 | ; | C5 | - | O1 |
| 726 | 745 | 1 | 1.0969E-01 | 2.7665E+05 | ; | C6 | - | H5 |
| 726 | 746 | 1 | 1.0969E-01 | 2.7665E+05 | ; | C6 | - | H6 |
| 726 | 747 | 1 | 1.0969E-01 | 2.7665E+05 | ; | C6 | - | H7 |
| 727 | 748 | 1 | 1.0969E-01 | 2.7665E+05 | ; | C7 | - | H8 |
| 727 | 757 | 1 | 1.0969E-01 | 2.7665E+05 | ; | C7 | - | H17 |
| 727 | 758 | 1 | 1.0969E-01 | 2.7665E+05 | ; | C7 | - | H18 |
| 728 | 729 | 1 | 1.2183E-01 | 5.3363E+05 | ; | C8 | - | O2 |
| 728 | 730 | 1 | 1.2183E-01 | 5.3363E+05 | ; | C8 | - | O3 |
| 731 | 732 | 1 | 1.3789E-01 | 3.5782E+05 | ; | N2 | - | C9 |
| 731 | 756 | 1 | 1.0129E-01 | 3.3740E+05 | ; | N2 | - | H16 |
| 732 | 733 | 1 | 1.2183E-01 | 5.3363E+05 | ; | C9 | - | O4 |
| 732 | 734 | 1 | 1.5241E-01 | 2.6192E+05 | ; | C9 | - | C10 |
| 734 | 735 | 1 | 1.5156E-01 | 2.6861E+05 | ; | C10 | - | C11 |
| 734 | 749 | 1 | 1.0969E-01 | 2.7665E+05 | ; | C10 | - | H9 |
| 734 | 750 | 1 | 1.0969E-01 | 2.7665E+05 | ; | C10 | - | H10 |
| 735 | 736 | 1 | 1.3984E-01 | 3.8585E+05 | ; | C11 | - | C12 |
| 735 | 740 | 1 | 1.3984E-01 | 3.8585E+05 | ; | C11 | - | C16 |
| 736 | 737 | 1 | 1.3984E-01 | 3.8585E+05 | ; | C12 | - | C13 |
| 736 | 751 | 1 | 1.0860E-01 | 2.8937E+05 | ; | C12 | - | H11 |
| 737 | 738 | 1 | 1.3984E-01 | 3.8585E+05 | ; | C13 | - | C14 |
| 737 | 752 | 1 | 1.0860E-01 | 2.8937E+05 | ; | C13 | - | H12 |
| 738 | 739 | 1 | 1.3984E-01 | 3.8585E+05 | ; | C14 | - | C15 |
| 738 | 753 | 1 | 1.0860E-01 | 2.8937E+05 | ; | C14 | - | H13 |
| 739 | 740 | 1 | 1.3984E-01 | 3.8585E+05 | ; | C15 | - | C16 |
| 739 | 754 | 1 | 1.0860E-01 | 2.8937E+05 | ; | C15 | - | H14 |
| 740 | 755 | 1 | 1.0860E-01 | 2.8937E+05 | ; | C16 | - | H15 |

[ pairs ]

; ai aj funct

| | | | | | | |
|---|---|---|---|---|---|---|
| 710 | 717 | 1 | ; | N | - | OG |
| 712 | 724 | 1 | ; | CA | - | C5 |
| 713 | 717 | 1 | ; | HA | - | OG |
| 714 | 723 | 1 | ; | CB | - | C4 |
| 714 | 725 | 1 | ; | CB | - | O1 |
| 715 | 724 | 1 | ; | HB1 | - | C5 |
| 716 | 724 | 1 | ; | HB2 | - | C5 |
| 717 | 722 | 1 | ; | OG | - | C3 |
| 717 | 731 | 1 | ; | OG | - | N2 |
| 717 | 744 | 1 | ; | OG | - | H4 |
| 717 | 759 | 1 | ; | OG | - | C |
| 718 | 724 | 1 | ; | S1 | - | C5 |
| 718 | 728 | 1 | ; | S1 | - | C8 |
| 718 | 731 | 1 | ; | S1 | - | N2 |
| 718 | 741 | 1 | ; | S1 | - | H1 |
| 718 | 742 | 1 | ; | S1 | - | H2 |

| | | | | | | |
|---|---|---|---|---|---|---|
| 718 | 744 | 1 | ; | S1 | - | H4 |
| 718 | 745 | 1 | ; | S1 | - | H5 |
| 718 | 746 | 1 | ; | S1 | - | H6 |
| 718 | 747 | 1 | ; | S1 | - | H7 |
| 718 | 748 | 1 | ; | S1 | - | H8 |
| 718 | 757 | 1 | ; | S1 | - | H17 |
| 718 | 758 | 1 | ; | S1 | - | H18 |
| 719 | 723 | 1 | ; | C1 | - | C4 |
| 719 | 729 | 1 | ; | C1 | - | O2 |
| 719 | 730 | 1 | ; | C1 | - | O3 |
| 719 | 742 | 1 | ; | C1 | - | H2 |
| 719 | 743 | 1 | ; | C1 | - | H3 |
| 720 | 723 | 1 | ; | C2 | - | C4 |
| 720 | 743 | 1 | ; | C2 | - | H3 |
| 720 | 745 | 1 | ; | C2 | - | H5 |
| 720 | 746 | 1 | ; | C2 | - | H6 |
| 720 | 747 | 1 | ; | C2 | - | H7 |
| 720 | 748 | 1 | ; | C2 | - | H8 |
| 720 | 757 | 1 | ; | C2 | - | H17 |
| 720 | 758 | 1 | ; | C2 | - | H18 |
| 721 | 724 | 1 | ; | N1 | - | C5 |
| 721 | 726 | 1 | ; | N1 | - | C6 |
| 721 | 727 | 1 | ; | N1 | - | C7 |
| 721 | 729 | 1 | ; | N1 | - | O2 |
| 721 | 730 | 1 | ; | N1 | - | O3 |
| 721 | 731 | 1 | ; | N1 | - | N2 |
| 721 | 744 | 1 | ; | N1 | - | H4 |
| 722 | 725 | 1 | ; | C3 | - | O1 |
| 722 | 726 | 1 | ; | C3 | - | C6 |
| 722 | 727 | 1 | ; | C3 | - | C7 |
| 722 | 728 | 1 | ; | C3 | - | C8 |
| 722 | 732 | 1 | ; | C3 | - | C9 |
| 722 | 741 | 1 | ; | C3 | - | H1 |
| 722 | 756 | 1 | ; | C3 | - | H16 |
| 723 | 733 | 1 | ; | C4 | - | O4 |
| 723 | 734 | 1 | ; | C4 | - | C10 |
| 723 | 742 | 1 | ; | C4 | - | H2 |
| 724 | 732 | 1 | ; | C5 | - | C9 |
| 724 | 743 | 1 | ; | C5 | - | H3 |
| 724 | 756 | 1 | ; | C5 | - | H16 |
| 725 | 731 | 1 | ; | O1 | - | N2 |
| 725 | 744 | 1 | ; | O1 | - | H4 |
| 726 | 728 | 1 | ; | C6 | - | C8 |
| 726 | 741 | 1 | ; | C6 | - | H1 |
| 726 | 748 | 1 | ; | C6 | - | H8 |
| 726 | 757 | 1 | ; | C6 | - | H17 |
| 726 | 758 | 1 | ; | C6 | - | H18 |
| 727 | 728 | 1 | ; | C7 | - | C8 |
| 727 | 741 | 1 | ; | C7 | - | H1 |
| 727 | 745 | 1 | ; | C7 | - | H5 |
| 727 | 746 | 1 | ; | C7 | - | H6 |
| 727 | 747 | 1 | ; | C7 | - | H7 |
| 728 | 742 | 1 | ; | C8 | - | H2 |
| 729 | 741 | 1 | ; | O2 | - | H1 |
| 730 | 741 | 1 | ; | O3 | - | H1 |
| 731 | 735 | 1 | ; | N2 | - | C11 |
| 731 | 743 | 1 | ; | N2 | - | H3 |
| 731 | 749 | 1 | ; | N2 | - | H9 |
| 731 | 750 | 1 | ; | N2 | - | H10 |
| 732 | 736 | 1 | ; | C9 | - | C12 |
| 732 | 740 | 1 | ; | C9 | - | C16 |
| 732 | 744 | 1 | ; | C9 | - | H4 |
| 733 | 735 | 1 | ; | O4 | - | C11 |
| 733 | 749 | 1 | ; | O4 | - | H9 |
| 733 | 750 | 1 | ; | O4 | - | H10 |
| 733 | 756 | 1 | ; | O4 | - | H16 |
| 734 | 737 | 1 | ; | C10 | - | C13 |
| 734 | 739 | 1 | ; | C10 | - | C15 |
| 734 | 751 | 1 | ; | C10 | - | H11 |
| 734 | 755 | 1 | ; | C10 | - | H15 |

| | | | | | | |
|---|---|---|---|---|---|---|
| 734 | 756 | 1 | ; | C10 | - | H16 |
| 735 | 738 | 1 | ; | C11 | - | C14 |
| 735 | 752 | 1 | ; | C11 | - | H12 |
| 735 | 754 | 1 | ; | C11 | - | H14 |
| 736 | 739 | 1 | ; | C12 | - | C15 |
| 736 | 749 | 1 | ; | C12 | - | H9 |
| 736 | 750 | 1 | ; | C12 | - | H10 |
| 736 | 753 | 1 | ; | C12 | - | H13 |
| 736 | 755 | 1 | ; | C12 | - | H15 |
| 737 | 740 | 1 | ; | C13 | - | C16 |
| 737 | 754 | 1 | ; | C13 | - | H14 |
| 738 | 751 | 1 | ; | C14 | - | H11 |
| 738 | 755 | 1 | ; | C14 | - | H15 |
| 739 | 752 | 1 | ; | C15 | - | H12 |
| 740 | 749 | 1 | ; | C16 | - | H9 |
| 740 | 750 | 1 | ; | C16 | - | H10 |
| 740 | 751 | 1 | ; | C16 | - | H11 |
| 740 | 753 | 1 | ; | C16 | - | H13 |
| 741 | 742 | 1 | ; | H1 | - | H2 |
| 742 | 743 | 1 | ; | H2 | - | H3 |
| 743 | 744 | 1 | ; | H3 | - | H4 |
| 744 | 756 | 1 | ; | H4 | - | H16 |
| 751 | 752 | 1 | ; | H11 | - | H12 |
| 752 | 753 | 1 | ; | H12 | - | H13 |
| 753 | 754 | 1 | ; | H13 | - | H14 |
| 754 | 755 | 1 | ; | H14 | - | H15 |

[ angles ]

| ; ai | aj | ak | funct | theta | cth | | | | | | |
|---|---|---|---|---|---|---|---|---|---|---|---|
| 712 | 714 | 717 | 1 | 1.0797E+02 | 5.6902E+02 | ; | CA | - | CB | - | OG |
| 714 | 717 | 724 | 1 | 1.1598E+02 | 5.2969E+02 | ; | CB | - | OG | - | C5 |
| 715 | 714 | 717 | 1 | 1.0978E+02 | 4.2509E+02 | ; | HB1 | - | CB | - | OG |
| 716 | 714 | 717 | 1 | 1.0978E+02 | 4.2509E+02 | ; | HB2 | - | CB | - | OG |
| 717 | 724 | 723 | 1 | 1.1072E+02 | 5.7656E+02 | ; | OG | - | C5 | - | C4 |
| 717 | 724 | 725 | 1 | 1.2325E+02 | 6.3011E+02 | ; | OG | - | C5 | - | O1 |
| 718 | 719 | 720 | 1 | 1.1027E+02 | 5.1296E+02 | ; | S1 | - | C1 | - | C2 |
| 718 | 719 | 726 | 1 | 1.1027E+02 | 5.1296E+02 | ; | S1 | - | C1 | - | C6 |
| 718 | 719 | 727 | 1 | 1.1027E+02 | 5.1296E+02 | ; | S1 | - | C1 | - | C7 |
| 718 | 722 | 721 | 1 | 1.0738E+02 | 5.3890E+02 | ; | S1 | - | C3 | - | N1 |
| 718 | 722 | 723 | 1 | 1.1027E+02 | 5.1296E+02 | ; | S1 | - | C3 | - | C4 |
| 718 | 722 | 743 | 1 | 1.0833E+02 | 3.5229E+02 | ; | S1 | - | C3 | - | H3 |
| 719 | 718 | 722 | 1 | 9.9240E+01 | 5.0375E+02 | ; | C1 | - | S1 | - | C3 |
| 719 | 720 | 721 | 1 | 1.1104E+02 | 5.5229E+02 | ; | C1 | - | C2 | - | N1 |
| 719 | 720 | 728 | 1 | 1.1104E+02 | 5.2969E+02 | ; | C1 | - | C2 | - | C8 |
| 719 | 720 | 741 | 1 | 1.0956E+02 | 3.8828E+02 | ; | C1 | - | C2 | - | H1 |
| 719 | 726 | 745 | 1 | 1.0980E+02 | 3.8744E+02 | ; | C1 | - | C6 | - | H5 |
| 719 | 726 | 746 | 1 | 1.0980E+02 | 3.8744E+02 | ; | C1 | - | C6 | - | H6 |
| 719 | 726 | 747 | 1 | 1.0980E+02 | 3.8744E+02 | ; | C1 | - | C6 | - | H7 |
| 719 | 727 | 748 | 1 | 1.0980E+02 | 3.8744E+02 | ; | C1 | - | C7 | - | H8 |

| | | | | | | | | | | | |
|---|---|---|---|---|---|---|---|---|---|---|---|
| 719 | 727 | 757 | 1 | 1.0980E+02 | 3.8744E+02 | ; | C1 | - | C7 | - | H17 |
| 719 | 727 | 758 | 1 | 1.0980E+02 | 3.8744E+02 | ; | C1 | - | C7 | - | H18 |
| 720 | 719 | 726 | 1 | 1.1151E+02 | 5.2635E+02 | ; | C2 | - | C1 | - | C6 |
| 720 | 719 | 727 | 1 | 1.1151E+02 | 5.2635E+02 | ; | C2 | - | C1 | - | C7 |
| 720 | 721 | 722 | 1 | 1.1235E+02 | 5.3388E+02 | ; | C2 | - | N1 | - | C3 |
| 720 | 721 | 742 | 1 | 1.0929E+02 | 3.9664E+02 | ; | C2 | - | N1 | - | H2 |
| 720 | 728 | 729 | 1 | 1.2320E+02 | 5.6400E+02 | ; | C2 | - | C8 | - | O2 |
| 720 | 728 | 730 | 1 | 1.2320E+02 | 5.6400E+02 | ; | C2 | - | C8 | - | O3 |
| 721 | 720 | 728 | 1 | 1.1114E+02 | 5.5480E+02 | ; | N1 | - | C2 | - | C8 |
| 721 | 720 | 741 | 1 | 1.0988E+02 | 4.1422E+02 | ; | N1 | - | C2 | - | H1 |
| 721 | 722 | 723 | 1 | 1.1104E+02 | 5.5229E+02 | ; | N1 | - | C3 | - | C4 |
| 721 | 722 | 743 | 1 | 1.0935E+02 | 4.1505E+02 | ; | N1 | - | C3 | - | H3 |
| 722 | 721 | 742 | 1 | 1.0929E+02 | 3.9664E+02 | ; | C3 | - | N1 | - | H2 |
| 722 | 723 | 724 | 1 | 1.1104E+02 | 5.2969E+02 | ; | C3 | - | C4 | - | C5 |
| 722 | 723 | 731 | 1 | 1.1161E+02 | 5.5145E+02 | ; | C3 | - | C4 | - | N2 |
| 722 | 723 | 744 | 1 | 1.0956E+02 | 3.8828E+02 | ; | C3 | - | C4 | - | H4 |
| 723 | 722 | 743 | 1 | 1.1022E+02 | 3.8660E+02 | ; | C4 | - | C3 | - | H3 |
| 723 | 724 | 725 | 1 | 1.2320E+02 | 5.6400E+02 | ; | C4 | - | C5 | - | O1 |
| 723 | 731 | 732 | 1 | 1.2069E+02 | 5.3053E+02 | ; | C4 | - | N2 | - | C9 |
| 723 | 731 | 756 | 1 | 1.1768E+02 | 3.8325E+02 | ; | C4 | - | N2 | - | H16 |
| 724 | 723 | 731 | 1 | 1.0906E+02 | 5.6066E+02 | ; | C5 | - | C4 | - | N2 |
| 724 | 723 | 744 | 1 | 1.0822E+02 | 3.9330E+02 | ; | C5 | - | C4 | - | H4 |
| 726 | 719 | 727 | 1 | 1.1151E+02 | 5.2635E+02 | ; | C6 | - | C1 | - | C7 |
| 728 | 720 | 741 | 1 | 1.0822E+02 | 3.9330E+02 | ; | C8 | - | C2 | - | H1 |
| 729 | 728 | 730 | 1 | 1.3025E+02 | 6.5187E+02 | ; | O2 | - | C8 | - | O3 |
| 731 | 723 | 744 | 1 | 1.0888E+02 | 4.1673E+02 | ; | N2 | - | C4 | - | H4 |
| 731 | 732 | 733 | 1 | 1.2305E+02 | 6.2091E+02 | ; | N2 | - | C9 | - | O4 |
| 731 | 732 | 734 | 1 | 1.1518E+02 | 5.5898E+02 | ; | N2 | - | C9 | - | C10 |
| 732 | 731 | 756 | 1 | 1.1755E+02 | 4.0417E+02 | ; | C9 | - | N2 | - | H16 |
| 732 | 734 | 735 | 1 | 1.1101E+02 | 5.3304E+02 | ; | C9 | - | C10 | - | C11 |
| 732 | 734 | 749 | 1 | 1.0877E+02 | 3.9246E+02 | ; | C9 | - | C10 | - | H9 |
| 732 | 734 | 750 | 1 | 1.0877E+02 | 3.9246E+02 | ; | C9 | - | C10 | - | H10 |
| 733 | 732 | 734 | 1 | 1.2320E+02 | 5.6400E+02 | ; | O4 | - | C9 | - | C10 |
| 734 | 735 | 736 | 1 | 1.2077E+02 | 5.3137E+02 | ; | C10 | - | C11 | - | C12 |
| 734 | 735 | 740 | 1 | 1.2077E+02 | 5.3137E+02 | ; | C10 | - | C11 | - | C16 |

| | | | | | | | |
|---|---|---|---|---|---|---|---|
| 735 | 734 | 749 | 1 | 1.1047E+02 | 3.9162E+02 | ; | C11 - C10 - H9 |
| 735 | 734 | 750 | 1 | 1.1047E+02 | 3.9162E+02 | ; | C11 - C10 - H10 |
| 735 | 736 | 737 | 1 | 1.2002E+02 | 5.5731E+02 | ; | C11 - C12 - C13 |
| 735 | 736 | 751 | 1 | 1.1988E+02 | 4.0334E+02 | ; | C11 - C12 - H11 |
| 735 | 740 | 739 | 1 | 1.2002E+02 | 5.5731E+02 | ; | C11 - C16 - C15 |
| 735 | 740 | 755 | 1 | 1.1988E+02 | 4.0334E+02 | ; | C11 - C16 - H15 |
| 736 | 735 | 740 | 1 | 1.2002E+02 | 5.5731E+02 | ; | C12 - C11 - C16 |
| 736 | 737 | 738 | 1 | 1.2002E+02 | 5.5731E+02 | ; | C12 - C13 - C14 |
| 736 | 737 | 752 | 1 | 1.1988E+02 | 4.0334E+02 | ; | C12 - C13 - H12 |
| 737 | 736 | 751 | 1 | 1.1988E+02 | 4.0334E+02 | ; | C13 - C12 - H11 |
| 737 | 738 | 739 | 1 | 1.2002E+02 | 5.5731E+02 | ; | C13 - C14 - C15 |
| 737 | 738 | 753 | 1 | 1.1988E+02 | 4.0334E+02 | ; | C13 - C14 - H13 |
| 738 | 737 | 752 | 1 | 1.1988E+02 | 4.0334E+02 | ; | C14 - C13 - H12 |
| 738 | 739 | 740 | 1 | 1.2002E+02 | 5.5731E+02 | ; | C14 - C15 - C16 |
| 738 | 739 | 754 | 1 | 1.1988E+02 | 4.0334E+02 | ; | C14 - C15 - H14 |
| 739 | 738 | 753 | 1 | 1.1988E+02 | 4.0334E+02 | ; | C15 - C14 - H13 |
| 739 | 740 | 755 | 1 | 1.1988E+02 | 4.0334E+02 | ; | C15 - C16 - H15 |
| 740 | 739 | 754 | 1 | 1.1988E+02 | 4.0334E+02 | ; | C16 - C15 - H14 |
| 745 | 726 | 746 | 1 | 1.0758E+02 | 3.2970E+02 | ; | H5 - C6 - H6 |
| 745 | 726 | 747 | 1 | 1.0758E+02 | 3.2970E+02 | ; | H5 - C6 - H7 |
| 746 | 726 | 747 | 1 | 1.0758E+02 | 3.2970E+02 | ; | H6 - C6 - H7 |
| 748 | 727 | 757 | 1 | 1.0758E+02 | 3.2970E+02 | ; | H8 - C7 - H17 |
| 748 | 727 | 758 | 1 | 1.0758E+02 | 3.2970E+02 | ; | H8 - C7 - H18 |
| 749 | 734 | 750 | 1 | 1.0758E+02 | 3.2970E+02 | ; | H9 - C10 - H10 |
| 757 | 727 | 758 | 1 | 1.0758E+02 | 3.2970E+02 | ; | H17 - C7 - H18 |

[ dihedrals ] ; propers
; treated as RBs in GROMACS to use combine multiple AMBER torsions per quartet

| ; i | j | k | l | func | C0 | C1 | C2 | C3 | C4 | C5 | | |
|---|---|---|---|---|---|---|---|---|---|---|---|---|
| 710 | 712 | 714 | 717 | 3 | 0.65084 | 1.95253 | 0.00000 | -2.60338 | 0.00000 | 0.00000 | ; | N- CA- CB- OG |
| 712 | 714 | 717 | 724 | 3 | 4.94967 | 8.15462 | 0.00000 | -6.40989 | 0.00000 | 0.00000 | ; | CA- CB- OG- C5 |
| 713 | 712 | 714 | 717 | 3 | 1.04600 | -1.04600 | 0.00000 | 0.00000 | 0.00000 | 0.00000 | ; | HA- CA- CB- OG |
| 714 | 717 | 724 | 723 | 3 | 27.40520 | 14.43480 | -22.59360 | -19.24640 | 0.00000 | 0.00000 | ; | CB- OG- C5- C4 |
| 714 | 717 | 724 | 725 | 3 | 28.45120 | 5.85760 | -22.59360 | 0.00000 | 0.00000 | 0.00000 | ; | CB- OG- C5- O1 |
| 715 | 714 | 717 | 724 | 3 | 1.60387 | 4.81160 | 0.00000 | -6.41547 | 0.00000 | 0.00000 | ; | HB1- CB- OG- C5 |
| 716 | 714 | 717 | 724 | 3 | 1.60387 | 4.81160 | 0.00000 | -6.41547 | 0.00000 | 0.00000 | ; | HB2- CB- OG- C5 |

| | | | | | | | | | | | | |
|---|---|---|---|---|---|---|---|---|---|---|---|---|
| 717 | 724 C5- | 723 C4- | 722 C3 | 3 | 0.00000 | 0.00000 | 0.00000 | 0.00000 | 0.00000 | 0.00000 | ; | OG- |
| 717 | 724 C5- | 723 C4- | 731 N2 | 3 | 0.00000 | 0.00000 | 0.00000 | 0.00000 | 0.00000 | 0.00000 | ; | OG- |
| 717 | 724 C5- | 723 C4- | 744 H4 | 3 | 0.00000 | 0.00000 | 0.00000 | 0.00000 | 0.00000 | 0.00000 | ; | OG- |
| 718 | 719 C1- | 720 C2- | 721 N1 | 3 | 0.65084 | 1.95253 | 0.00000 | -2.60338 | 0.00000 | 0.00000 | ; | S1- |
| 718 | 719 C1- | 720 C2- | 728 C8 | 3 | 0.65084 | 1.95253 | 0.00000 | -2.60338 | 0.00000 | 0.00000 | ; | S1- |
| 718 | 719 C1- | 720 C2- | 741 H1 | 3 | 0.65084 | 1.95253 | 0.00000 | -2.60338 | 0.00000 | 0.00000 | ; | S1- |
| 718 | 719 C1- | 726 C6- | 745 H5 | 3 | 0.65084 | 1.95253 | 0.00000 | -2.60338 | 0.00000 | 0.00000 | ; | S1- |
| 718 | 719 C1- | 726 C6- | 746 H6 | 3 | 0.65084 | 1.95253 | 0.00000 | -2.60338 | 0.00000 | 0.00000 | ; | S1- |
| 718 | 719 C1- | 726 C6- | 747 H7 | 3 | 0.65084 | 1.95253 | 0.00000 | -2.60338 | 0.00000 | 0.00000 | ; | S1- |
| 718 | 719 C1- | 727 C7- | 748 H8 | 3 | 0.65084 | 1.95253 | 0.00000 | -2.60338 | 0.00000 | 0.00000 | ; | S1- |
| 718 | 719 C1- | 727 C7- | 757 H17 | 3 | 0.65084 | 1.95253 | 0.00000 | -2.60338 | 0.00000 | 0.00000 | ; | S1- |
| 718 | 719 C1- | 727 C7- | 758 H18 | 3 | 0.65084 | 1.95253 | 0.00000 | -2.60338 | 0.00000 | 0.00000 | ; | S1- |
| 718 | 722 C3- | 721 N1- | 720 C2 | 3 | 1.25520 | 3.76560 | 0.00000 | -5.02080 | 0.00000 | 0.00000 | ; | S1- |
| 718 | 722 C3- | 721 N1- | 742 H2 | 3 | 1.25520 | 3.76560 | 0.00000 | -5.02080 | 0.00000 | 0.00000 | ; | S1- |
| 718 | 722 C3- | 723 C4- | 724 C5 | 3 | 0.65084 | 1.95253 | 0.00000 | -2.60338 | 0.00000 | 0.00000 | ; | S1- |
| 718 | 722 C3- | 723 C4- | 731 N2 | 3 | 0.65084 | 1.95253 | 0.00000 | -2.60338 | 0.00000 | 0.00000 | ; | S1- |
| 718 | 722 C3- | 723 C4- | 744 H4 | 3 | 0.65084 | 1.95253 | 0.00000 | -2.60338 | 0.00000 | 0.00000 | ; | S1- |
| 719 | 718 S1- | 722 C3- | 721 N1 | 3 | 1.39467 | 4.18400 | 0.00000 | -5.57867 | 0.00000 | 0.00000 | ; | C1- |
| 719 | 718 S1- | 722 C3- | 723 C4 | 3 | 1.39467 | 4.18400 | 0.00000 | -5.57867 | 0.00000 | 0.00000 | ; | C1- |
| 719 | 718 S1- | 722 C3- | 743 H3 | 3 | 1.39467 | 4.18400 | 0.00000 | -5.57867 | 0.00000 | 0.00000 | ; | C1- |
| 719 | 720 C2- | 721 N1- | 722 C3 | 3 | 5.27184 | 3.76560 | -4.01664 | -5.02080 | 0.00000 | 0.00000 | ; | C1- |
| 719 | 720 C2- | 721 N1- | 742 H2 | 3 | 1.25520 | 3.76560 | 0.00000 | -5.02080 | 0.00000 | 0.00000 | ; | C1- |
| 719 | 720 C2- | 728 C8- | 729 O2 | 3 | 0.00000 | 0.00000 | 0.00000 | 0.00000 | 0.00000 | 0.00000 | ; | C1- |
| 719 | 720 C2- | 728 C8- | 730 O3 | 3 | 0.00000 | 0.00000 | 0.00000 | 0.00000 | 0.00000 | 0.00000 | ; | C1- |
| 720 | 719 C1- | 718 S1- | 722 C3 | 3 | 1.39467 | 4.18400 | 0.00000 | -5.57867 | 0.00000 | 0.00000 | ; | C2- |
| 720 | 719 C1- | 726 C6- | 745 H5 | 3 | 0.66944 | 2.00832 | 0.00000 | -2.67776 | 0.00000 | 0.00000 | ; | C2- |
| 720 | 719 C1- | 726 C6- | 746 H6 | 3 | 0.66944 | 2.00832 | 0.00000 | -2.67776 | 0.00000 | 0.00000 | ; | C2- |
| 720 | 719 C1- | 726 C6- | 747 H7 | 3 | 0.66944 | 2.00832 | 0.00000 | -2.67776 | 0.00000 | 0.00000 | ; | C2- |
| 720 | 719 C1- | 727 C7- | 748 H8 | 3 | 0.66944 | 2.00832 | 0.00000 | -2.67776 | 0.00000 | 0.00000 | ; | C2- |
| 720 | 719 C1- | 727 C7- | 757 H17 | 3 | 0.66944 | 2.00832 | 0.00000 | -2.67776 | 0.00000 | 0.00000 | ; | C2- |
| 720 | 719 C1- | 727 C7- | 758 H18 | 3 | 0.66944 | 2.00832 | 0.00000 | -2.67776 | 0.00000 | 0.00000 | ; | C2- |
| 720 | 721 N1- | 722 C3- | 723 C4 | 3 | 5.27184 | 3.76560 | -4.01664 | -5.02080 | 0.00000 | 0.00000 | ; | C2- |
| 720 | 721 N1- | 722 C3- | 743 H3 | 3 | 1.25520 | 3.76560 | 0.00000 | -5.02080 | 0.00000 | 0.00000 | ; | C2- |
| 721 | 720 C2- | 719 C1- | 726 C6 | 3 | 0.65084 | 1.95253 | 0.00000 | -2.60338 | 0.00000 | 0.00000 | ; | N1- |
| 721 | 720 C2- | 719 C1- | 727 C7 | 3 | 0.65084 | 1.95253 | 0.00000 | -2.60338 | 0.00000 | 0.00000 | ; | N1- |

| | | | | | | | | | | | | |
|---|---|---|---|---|---|---|---|---|---|---|---|---|
| 721 | 720 C2- | 728 C8- | 729 O2 | 3 | 0.00000 | 0.00000 | 0.00000 | 0.00000 | 0.00000 | 0.00000 | ; | N1- |
| 721 | 720 C2- | 728 C8- | 730 O3 | 3 | 0.00000 | 0.00000 | 0.00000 | 0.00000 | 0.00000 | 0.00000 | ; | N1- |
| 721 | 722 C3- | 723 C4- | 724 C5 | 3 | 0.65084 | 1.95253 | 0.00000 | -2.60338 | 0.00000 | 0.00000 | ; | N1- |
| 721 | 722 C3- | 723 C4- | 731 N2 | 3 | 0.65084 | 1.95253 | 0.00000 | -2.60338 | 0.00000 | 0.00000 | ; | N1- |
| 721 | 722 C3- | 723 C4- | 744 H4 | 3 | 0.65084 | 1.95253 | 0.00000 | -2.60338 | 0.00000 | 0.00000 | ; | N1- |
| 722 | 718 S1- | 719 C1- | 726 C6 | 3 | 1.39467 | 4.18400 | 0.00000 | -5.57867 | 0.00000 | 0.00000 | ; | C3- |
| 722 | 718 S1- | 719 C1- | 727 C7 | 3 | 1.39467 | 4.18400 | 0.00000 | -5.57867 | 0.00000 | 0.00000 | ; | C3- |
| 722 | 721 N1- | 720 C2- | 728 C8 | 3 | 1.25520 | 3.76560 | 0.00000 | -5.02080 | 0.00000 | 0.00000 | ; | C3- |
| 722 | 721 N1- | 720 C2- | 741 H1 | 3 | 1.25520 | 3.76560 | 0.00000 | -5.02080 | 0.00000 | 0.00000 | ; | C3- |
| 722 | 723 C4- | 724 C5- | 725 O1 | 3 | 0.00000 | 0.00000 | 0.00000 | 0.00000 | 0.00000 | 0.00000 | ; | C3- |
| 722 | 723 C3- | 731 C4- | 732 N2- | 3 C9 | 2.84512 | -4.10032 | 16.73600 | 2.51040 | -16.73600 | | 0.00000 | ; |
| 722 | 723 C4- | 731 N2- | 756 H16 | 3 | 0.00000 | 0.00000 | 0.00000 | 0.00000 | 0.00000 | 0.00000 | ; | C3- |
| 723 | 722 C3- | 721 N1- | 742 H2 | 3 | 1.25520 | 3.76560 | 0.00000 | -5.02080 | 0.00000 | 0.00000 | ; | C4- |
| 723 | 731 C4- | 732 N2- | 733 C9- | 3 O4 | 20.92000 | 0.00000 | -20.92000 | | 0.00000 | 0.00000 | 0.00000 | ; |
| 723 | 731 N2- | 732 C9- | 734 C10 | 3 | 6.27600 | 6.27600 | 0.00000 | 0.00000 | 0.00000 | 0.00000 | ; | C4- |
| 724 | 723 C4- | 722 C3- | 743 H3 | 3 | 0.65084 | 1.95253 | 0.00000 | -2.60338 | 0.00000 | 0.00000 | ; | C5- |
| 724 | 723 C4- | 731 N2- | 732 C9 | 3 | 10.46000 | -3.34720 | -7.11280 | 0.00000 | 0.00000 | 0.00000 | ; | C5- |
| 724 | 723 C4- | 731 N2- | 756 H16 | 3 | 0.00000 | 0.00000 | 0.00000 | 0.00000 | 0.00000 | 0.00000 | ; | C5- |
| 725 | 724 C5- | 723 C4- | 731 N2 | 3 | 0.00000 | 0.00000 | 0.00000 | 0.00000 | 0.00000 | 0.00000 | ; | O1- |
| 725 | 724 C5- | 723 C4- | 744 H4 | 3 | 3.68192 | -4.35136 | 0.00000 | 1.33888 | 0.00000 | 0.00000 | ; | O1- |
| 726 | 719 C1- | 720 C2- | 728 C8 | 3 | 0.65084 | 1.95253 | 0.00000 | -2.60338 | 0.00000 | 0.00000 | ; | C6- |
| 726 | 719 C1- | 720 C2- | 741 H1 | 3 | 0.65084 | 1.95253 | 0.00000 | -2.60338 | 0.00000 | 0.00000 | ; | C6- |
| 726 | 719 C1- | 727 C7- | 748 H8 | 3 | 0.66944 | 2.00832 | 0.00000 | -2.67776 | 0.00000 | 0.00000 | ; | C6- |
| 726 | 719 C1- | 727 C7- | 757 H17 | 3 | 0.66944 | 2.00832 | 0.00000 | -2.67776 | 0.00000 | 0.00000 | ; | C6- |
| 726 | 719 C1- | 727 C7- | 758 H18 | 3 | 0.66944 | 2.00832 | 0.00000 | -2.67776 | 0.00000 | 0.00000 | ; | C6- |
| 727 | 719 C1- | 720 C2- | 728 C8 | 3 | 0.65084 | 1.95253 | 0.00000 | -2.60338 | 0.00000 | 0.00000 | ; | C7- |
| 727 | 719 C1- | 720 C2- | 741 H1 | 3 | 0.65084 | 1.95253 | 0.00000 | -2.60338 | 0.00000 | 0.00000 | ; | C7- |
| 727 | 719 C1- | 726 C6- | 745 H5 | 3 | 0.66944 | 2.00832 | 0.00000 | -2.67776 | 0.00000 | 0.00000 | ; | C7- |
| 727 | 719 C1- | 726 C6- | 746 H6 | 3 | 0.66944 | 2.00832 | 0.00000 | -2.67776 | 0.00000 | 0.00000 | ; | C7- |
| 727 | 719 C1- | 726 C6- | 747 H7 | 3 | 0.66944 | 2.00832 | 0.00000 | -2.67776 | 0.00000 | 0.00000 | ; | C7- |
| 728 | 720 C2- | 721 N1- | 742 H2 | 3 | 1.25520 | 3.76560 | 0.00000 | -5.02080 | 0.00000 | 0.00000 | ; | C8- |
| 729 | 728 C8- | 720 C2- | 741 H1 | 3 | 3.68192 | -4.35136 | 0.00000 | 1.33888 | 0.00000 | 0.00000 | ; | O2- |
| 730 | 728 C8- | 720 C2- | 741 H1 | 3 | 3.68192 | -4.35136 | 0.00000 | 1.33888 | 0.00000 | 0.00000 | ; | O3- |
| 731 | 723 C4- | 722 C3- | 743 H3 | 3 | 0.65084 | 1.95253 | 0.00000 | -2.60338 | 0.00000 | 0.00000 | ; | N2- |
| 731 | 732 C9- | 734 C10- | 735 C11 | 3 | 0.00000 | 0.00000 | 0.00000 | 0.00000 | 0.00000 | 0.00000 | ; | N2- |

| | | | | | | | | | | | | |
|---|---|---|---|---|---|---|---|---|---|---|---|---|
| 731 | 732 C9- | 734 C10- | 749 H9 | 3 | 0.00000 | 0.00000 | 0.00000 | 0.00000 | 0.00000 | 0.00000 | ; | N2- |
| 731 | 732 C9- | 734 C10- | 750 H10 | 3 | 0.00000 | 0.00000 | 0.00000 | 0.00000 | 0.00000 | 0.00000 | ; | N2- |
| 732 | 731 N2- | 723 C4- | 744 H4 | 3 | 0.00000 | 0.00000 | 0.00000 | 0.00000 | 0.00000 | 0.00000 | ; | C9- |
| 732 | 734 C10- | 735 C11- | 736 C12 | 3 | 0.00000 | 0.00000 | 0.00000 | 0.00000 | 0.00000 | 0.00000 | ; | C9- |
| 732 | 734 C10- | 735 C11- | 740 C16 | 3 | 0.00000 | 0.00000 | 0.00000 | 0.00000 | 0.00000 | 0.00000 | ; | C9- |
| 733 | 732 O4- | 731 C9- | 756 N2- | 3 H16 | 29.28800 | -8.36800 | -20.92000 | | 0.00000 | 0.00000 | 0.00000 | ; |
| 733 | 732 C9- | 734 C10- | 735 C11 | 3 | 0.00000 | 0.00000 | 0.00000 | 0.00000 | 0.00000 | 0.00000 | ; | O4- |
| 733 | 732 C9- | 734 C10- | 749 H9 | 3 | 3.68192 | -4.35136 | 0.00000 | 1.33888 | 0.00000 | 0.00000 | ; | O4- |
| 733 | 732 C9- | 734 C10- | 750 H10 | 3 | 3.68192 | -4.35136 | 0.00000 | 1.33888 | 0.00000 | 0.00000 | ; | O4- |
| 734 | 732 C10- | 731 C9- | 756 N2- | 3 H16 | 20.92000 | 0.00000 | -20.92000 | | 0.00000 | 0.00000 | 0.00000 | ; |
| 734 | 735 C10- | 736 C11- | 737 C12- | 3 C13 | 30.33400 | 0.00000 | -30.33400 | | 0.00000 | 0.00000 | 0.00000 | ; |
| 734 | 735 C10- | 736 C11- | 751 C12- | 3 H11 | 30.33400 | 0.00000 | -30.33400 | | 0.00000 | 0.00000 | 0.00000 | ; |
| 734 | 735 C10- | 740 C11- | 739 C16- | 3 C15 | 30.33400 | 0.00000 | -30.33400 | | 0.00000 | 0.00000 | 0.00000 | ; |
| 734 | 735 C10- | 740 C11- | 755 C16- | 3 H15 | 30.33400 | 0.00000 | -30.33400 | | 0.00000 | 0.00000 | 0.00000 | ; |
| 735 | 736 C11- | 737 C12- | 738 C13- | 3 C14 | 30.33400 | 0.00000 | -30.33400 | | 0.00000 | 0.00000 | 0.00000 | ; |
| 735 | 736 C11- | 737 C12- | 752 C13- | 3 H12 | 30.33400 | 0.00000 | -30.33400 | | 0.00000 | 0.00000 | 0.00000 | ; |
| 735 | 740 C11- | 739 C16- | 738 C15- | 3 C14 | 30.33400 | 0.00000 | -30.33400 | | 0.00000 | 0.00000 | 0.00000 | ; |
| 735 | 740 C11- | 739 C16- | 754 C15- | 3 H14 | 30.33400 | 0.00000 | -30.33400 | | 0.00000 | 0.00000 | 0.00000 | ; |
| 736 | 735 C11- | 734 C10- | 749 H9 | 3 | 0.00000 | 0.00000 | 0.00000 | 0.00000 | 0.00000 | 0.00000 | ; | C12- |
| 736 | 735 C11- | 734 C10- | 750 H10 | 3 | 0.00000 | 0.00000 | 0.00000 | 0.00000 | 0.00000 | 0.00000 | ; | C12- |
| 736 | 735 C12- | 740 C11- | 739 C16- | 3 C15 | 30.33400 | 0.00000 | -30.33400 | | 0.00000 | 0.00000 | 0.00000 | ; |
| 736 | 735 C12- | 740 C11- | 755 C16- | 3 H15 | 30.33400 | 0.00000 | -30.33400 | | 0.00000 | 0.00000 | 0.00000 | ; |
| 736 | 737 C12- | 738 C13- | 739 C14- | 3 C15 | 30.33400 | 0.00000 | -30.33400 | | 0.00000 | 0.00000 | 0.00000 | ; |
| 736 | 737 C12- | 738 C13- | 753 C14- | 3 H13 | 30.33400 | 0.00000 | -30.33400 | | 0.00000 | 0.00000 | 0.00000 | ; |
| 737 | 736 C13- | 735 C12- | 740 C11- | 3 C16 | 30.33400 | 0.00000 | -30.33400 | | 0.00000 | 0.00000 | 0.00000 | ; |
| 737 | 738 C13- | 739 C14- | 740 C15- | 3 C16 | 30.33400 | 0.00000 | -30.33400 | | 0.00000 | 0.00000 | 0.00000 | ; |
| 737 | 738 C13- | 739 C14- | 754 C15- | 3 H14 | 30.33400 | 0.00000 | -30.33400 | | 0.00000 | 0.00000 | 0.00000 | ; |
| 738 | 737 C14- | 736 C13- | 751 C12- | 3 H11 | 30.33400 | 0.00000 | -30.33400 | | 0.00000 | 0.00000 | 0.00000 | ; |
| 738 | 739 C14- | 740 C15- | 755 C16- | 3 H15 | 30.33400 | 0.00000 | -30.33400 | | 0.00000 | 0.00000 | 0.00000 | ; |
| 739 | 738 C15- | 737 C14- | 752 C13- | 3 H12 | 30.33400 | 0.00000 | -30.33400 | | 0.00000 | 0.00000 | 0.00000 | ; |
| 740 | 735 C11- | 734 C10- | 749 H9 | 3 | 0.00000 | 0.00000 | 0.00000 | 0.00000 | 0.00000 | 0.00000 | ; | C16- |
| 740 | 735 C11- | 734 C10- | 750 H10 | 3 | 0.00000 | 0.00000 | 0.00000 | 0.00000 | 0.00000 | 0.00000 | ; | C16- |
| 740 | 735 C16- | 736 C11- | 751 C12- | 3 H11 | 30.33400 | 0.00000 | -30.33400 | | 0.00000 | 0.00000 | 0.00000 | ; |
| 740 | 739 C16- | 738 C15- | 753 C14- | 3 H13 | 30.33400 | 0.00000 | -30.33400 | | 0.00000 | 0.00000 | 0.00000 | ; |
| 741 | 720 C2- | 721 N1- | 742 H2 | 3 | 1.25520 | 3.76560 | 0.00000 | -5.02080 | 0.00000 | 0.00000 | ; | H1- |

| | | | | | | | | | | | | |
|---|---|---|---|---|---|---|---|---|---|---|---|---|
| 742 | 721 | 722 | 743 | 3 | 1.25520 | 3.76560 | 0.00000 | -5.02080 | 0.00000 | 0.00000 | ; | H2- |
| | N1- | C3- | H3 | | | | | | | | | |
| 743 | 722 | 723 | 744 | 3 | 0.65084 | 1.95253 | 0.00000 | -2.60338 | 0.00000 | 0.00000 | ; | H3- |
| | C3- | C4- | H4 | | | | | | | | | |
| 744 | 723 | 731 | 756 | 3 | 0.00000 | 0.00000 | 0.00000 | 0.00000 | 0.00000 | 0.00000 | ; | H4- |
| | C4- | N2- | H16 | | | | | | | | | |
| 751 | 736 | 737 | 752 | 3 | 30.33400 | 0.00000 | -30.33400 | | 0.00000 | 0.00000 | 0.00000 | ; |
| | H11- | C12- | C13- | H12 | | | | | | | | |
| 752 | 737 | 738 | 753 | 3 | 30.33400 | 0.00000 | -30.33400 | | 0.00000 | 0.00000 | 0.00000 | ; |
| | H12- | C13- | C14- | H13 | | | | | | | | |
| 753 | 738 | 739 | 754 | 3 | 30.33400 | 0.00000 | -30.33400 | | 0.00000 | 0.00000 | 0.00000 | ; |
| | H13- | C14- | C15- | H14 | | | | | | | | |
| 754 | 739 | 740 | 755 | 3 | 30.33400 | 0.00000 | -30.33400 | | 0.00000 | 0.00000 | 0.00000 | ; |
| | H14- | C15- | C16- | H15 | | | | | | | | |

[ dihedrals ] ; impropers
; treated as propers in GROMACS to use correct AMBER analytical function
; i j k l func phase kd pn

| | | | | | | | | | | | | |
|---|---|---|---|---|---|---|---|---|---|---|---|---|
| 720 | 729 | 728 | 730 | 1 | 180.00000 | | 4.60240 | 2 | ; | C2- | O2- | C8- |
| | O3 | | | | | | | | | | | |
| 723 | 725 | 724 | 717 | 1 | 180.00000 | | 43.93200 | 2 | ; | C4- | O1- | C5- |
| | OG | | | | | | | | | | | |
| 732 | 723 | 731 | 756 | 1 | 180.00000 | | 4.60240 | 2 | ; | C9- | C4- | N2- |
| | H16 | | | | | | | | | | | |
| 734 | 731 | 732 | 733 | 1 | 180.00000 | | 43.93200 | 2 | ; | C10- | N2- | C9- |
| | O4 | | | | | | | | | | | |
| 735 | 737 | 736 | 751 | 1 | 180.00000 | | 4.60240 | 2 | ; | C11- | C13- | C12- |
| | H11 | | | | | | | | | | | |
| 735 | 739 | 740 | 755 | 1 | 180.00000 | | 4.60240 | 2 | ; | C11- | C15- | C16- |
| | H15 | | | | | | | | | | | |
| 736 | 738 | 737 | 752 | 1 | 180.00000 | | 4.60240 | 2 | ; | C12- | C14- | C13- |
| | H12 | | | | | | | | | | | |
| 736 | 740 | 735 | 734 | 1 | 180.00000 | | 4.60240 | 2 | ; | C12- | C16- | C11- |
| | C10 | | | | | | | | | | | |
| 737 | 739 | 738 | 753 | 1 | 180.00000 | | 4.60240 | 2 | ; | C13- | C15- | C14- |
| | H13 | | | | | | | | | | | |
| 738 | 740 | 739 | 754 | 1 | 180.00000 | | 4.60240 | 2 | ; | C14- | C16- | C15- |
| | H14 | | | | | | | | | | | |

#### CTX.gro

```
 289CTX      C 4061   4.669   3.785   3.531
 289CTX     C1 4062   4.792   3.756   3.587
 289CTX     C2 4063   4.635   3.964   3.709
 289CTX      H 4064   4.708   4.037   3.673
 289CTX      S 4065   4.705   3.850   3.834
 289CTX      N 4066   4.582   3.871   3.609
 289CTX     C3 4067   4.488   4.012   3.734
 289CTX     H1 4068   4.463   4.102   3.676
 289CTX     C4 4069   4.449   3.882   3.652
 289CTX      O 4070   4.350   3.814   3.633
 289CTX     N1 4071   4.451   4.038   3.873
 289CTX     C5 4072   4.619   3.732   3.399
 289CTX     O1 4073   4.670   3.633   3.349
 289CTX     O2 4074   4.526   3.788   3.343
 289CTX     C6 4075   4.899   3.673   3.515
 289CTX     H2 4076   4.887   3.569   3.546
 289CTX     H3 4077   4.888   3.674   3.406
 289CTX     C7 4078   4.842   3.797   3.725
 289CTX     H4 4079   4.913   3.880   3.717
 289CTX     H5 4080   4.893   3.714   3.773
 289CTX     O3 4081   5.033   3.715   3.550
 289CTX     C8 4082   5.103   3.815   3.485
 289CTX     C9 4083   5.030   3.889   3.374
 289CTX     O4 4084   5.219   3.839   3.512
 289CTX     H6 4085   5.100   3.960   3.327
 289CTX     H7 4086   4.995   3.820   3.298
 289CTX     H8 4087   4.944   3.944   3.413
 289CTX    C10 4088   4.527   4.121   3.954
 289CTX     O5 4089   4.616   4.189   3.904
 289CTX     H9 4090   4.381   3.981   3.919
 289CTX    C11 4091   4.507   4.118   4.103
 289CTX     N2 4092   4.427   4.028   4.154
 289CTX     O6 4093   4.390   4.064   4.285
 289CTX    C12 4094   4.392   3.955   4.376
 289CTX    H10 4095   4.321   3.877   4.346
 289CTX    H11 4096   4.493   3.912   4.380
 289CTX    H12 4097   4.364   3.990   4.477
 289CTX    C13 4098   4.619   4.158   4.189
 289CTX    C14 4099   4.737   4.087   4.198
 289CTX    H13 4100   4.774   4.012   4.129
 289CTX    C15 4101   4.684   4.218   4.393
 289CTX     S1 4102   4.811   4.104   4.356
 289CTX     N3 4103   4.597   4.241   4.298
 289CTX     N4 4104   4.653   4.263   4.517
 289CTX    H14 4105   4.554   4.291   4.517
 289CTX    H15 4106   4.674   4.198   4.592
```

#### CTX.top

```
[ defaults ]
; nbfunc        comb-rule       gen-pairs       fudgeLJ fudgeQQ
1               2               yes             0.5     0.8333

[ atomtypes ]
;name   bond_type     mass     charge   ptype   sigma         epsilon       Amb
 ce       ce          0.00000  0.00000   A     3.39967e-01   3.59824e-01 ; 1.91  0.0860
 c2       c2          0.00000  0.00000   A     3.39967e-01   3.59824e-01 ; 1.91  0.0860
```

```
 cy      cy         0.00000  0.00000   A     3.39967e-01   3.59824e-01 ; 1.91  0.0860
 h2      h2         0.00000  0.00000   A     2.29317e-01   6.56888e-02 ; 1.29  0.0157
 ss      ss         0.00000  0.00000   A     3.56359e-01   1.04600e+00 ; 2.00  0.2500
 nj      nj         0.00000  0.00000   A     3.25000e-01   7.11280e-01 ; 1.82  0.1700
 h1      h1         0.00000  0.00000   A     2.47135e-01   6.56888e-02 ; 1.39  0.0157
 c       c          0.00000  0.00000   A     3.39967e-01   3.59824e-01 ; 1.91  0.0860
 o       o          0.00000  0.00000   A     2.95992e-01   8.78640e-01 ; 1.66  0.2100
 n       n          0.00000  0.00000   A     3.25000e-01   7.11280e-01 ; 1.82  0.1700
 c3      c3         0.00000  0.00000   A     3.39967e-01   4.57730e-01 ; 1.91  0.1094
 os      os         0.00000  0.00000   A     3.00001e-01   7.11280e-01 ; 1.68  0.1700
 hc      hc         0.00000  0.00000   A     2.64953e-01   6.56888e-02 ; 1.49  0.0157
 hn      hn         0.00000  0.00000   A     1.06908e-01   6.56888e-02 ; 0.60  0.0157
 n2      n2         0.00000  0.00000   A     3.25000e-01   7.11280e-01 ; 1.82  0.1700
 cc      cc         0.00000  0.00000   A     3.39967e-01   3.59824e-01 ; 1.91  0.0860
 cd      cd         0.00000  0.00000   A     3.39967e-01   3.59824e-01 ; 1.91  0.0860
 h4      h4         0.00000  0.00000   A     2.51055e-01   6.27600e-02 ; 1.41  0.0150
 na      na         0.00000  0.00000   A     3.25000e-01   7.11280e-01 ; 1.82  0.1700
 nh      nh         0.00000  0.00000   A     3.25000e-01   7.11280e-01 ; 1.82  0.1700
 nc      nc         0.00000  0.00000   A     3.25000e-01   7.11280e-01 ; 1.82  0.1700

[ moleculetype ]
;name            nrexcl
 CTX             3

[ atoms ]
;   nr  type  resi  res  atom  cgnr     charge      mass       ; qtot   bond_type
     1   ce    1   CTX     C    1     0.007400     12.01000 ; qtot 0.007
     2   c2    1   CTX    C1    2    -0.299400     12.01000 ; qtot -0.292
     3   cy    1   CTX    C2    3     0.104800     12.01000 ; qtot -0.187
     4   h2    1   CTX     H    4     0.091700      1.00800 ; qtot -0.096
     5   ss    1   CTX     S    5    -0.360200     32.06000 ; qtot -0.456
     6   nj    1   CTX     N    6    -0.333000     14.01000 ; qtot -0.789
     7   cy    1   CTX    C3    7     0.003700     12.01000 ; qtot -0.785
     8   h1    1   CTX    H1    8     0.114700      1.00800 ; qtot -0.670
     9   c     1   CTX    C4    9     0.700501     12.01000 ; qtot 0.030
    10   o     1   CTX     O   10    -0.502501     16.00000 ; qtot -0.472
    11   n     1   CTX    N1   11    -0.518901     14.01000 ; qtot -0.991
    12   c     1   CTX    C5   12     0.927802     12.01000 ; qtot -0.063
    13   o     1   CTX    O1   13    -0.790801     16.00000 ; qtot -0.854
    14   o     1   CTX    O2   14    -0.790801     16.00000 ; qtot -1.645
    15   c3    1   CTX    C6   15     0.198600     12.01000 ; qtot -1.446
    16   h1    1   CTX    H2   16     0.072700      1.00800 ; qtot -1.374
    17   h1    1   CTX    H3   17     0.072700      1.00800 ; qtot -1.301
    18   c3    1   CTX    C7   18     0.086900     12.01000 ; qtot -1.214
    19   h1    1   CTX    H4   19     0.072700      1.00800 ; qtot -1.141
    20   h1    1   CTX    H5   20     0.072700      1.00800 ; qtot -1.069
    21   os    1   CTX    O3   21    -0.441900     16.00000 ; qtot -1.511
    22   c     1   CTX    C8   22     0.638101     12.01000 ; qtot -0.872
    23   c3    1   CTX    C9   23    -0.200100     12.01000 ; qtot -1.073
    24   o     1   CTX    O4   24    -0.557001     16.00000 ; qtot -1.630
    25   hc    1   CTX    H6   25     0.089367      1.00800 ; qtot -1.540
    26   hc    1   CTX    H7   26     0.089367      1.00800 ; qtot -1.451
    27   hc    1   CTX    H8   27     0.089367      1.00800 ; qtot -1.361
    28   c     1   CTX   C10   28     0.609801     12.01000 ; qtot -0.752
    29   o     1   CTX    O5   29    -0.609101     16.00000 ; qtot -1.361
    30   hn    1   CTX    H9   30     0.346500      1.00800 ; qtot -1.014
    31   ce    1   CTX   C11   31     0.284300     12.01000 ; qtot -0.730
    32   n2    1   CTX    N2   32    -0.387200     14.01000 ; qtot -1.117
    33   os    1   CTX    O6   33    -0.163300     16.00000 ; qtot -1.280
    34   c3    1   CTX   C12   34     0.127700     12.01000 ; qtot -1.153
    35   h1    1   CTX   H10   35     0.050033      1.00800 ; qtot -1.103
    36   h1    1   CTX   H11   36     0.050033      1.00800 ; qtot -1.053
    37   h1    1   CTX   H12   37     0.050033      1.00800 ; qtot -1.003
    38   cc    1   CTX   C13   38     0.296000     12.01000 ; qtot -0.707
    39   cd    1   CTX   C14   39    -0.227400     12.01000 ; qtot -0.934
    40   h4    1   CTX   H13   40     0.211000      1.00800 ; qtot -0.723
    41   cd    1   CTX   C15   41     0.560301     12.01000 ; qtot -0.163
    42   ss    1   CTX    S1   42    -0.081300     32.06000 ; qtot -0.244
    43   nc    1   CTX    N3   43    -0.641001     14.01000 ; qtot -0.885
    44   nh    1   CTX    N4   44    -0.876501     14.01000 ; qtot -1.762
```

```
   45   hn     1   CTX   H14   45     0.380800      1.00800 ; qtot -1.381
   46   hn     1   CTX   H15   46     0.380800      1.00800 ; qtot -1.000

[ bonds ]
;   ai     aj funct   r             k
    1      2   1    1.3461e-01    4.5798e+05 ;      C - C1
    1      6   1    1.4250e-01    3.0861e+05 ;      C - N
    1     12   1    1.4825e-01    2.9665e+05 ;      C - C5
    2     15   1    1.5095e-01    2.7347e+05 ;     C1 - C6
    2     18   1    1.5095e-01    2.7347e+05 ;     C1 - C7
    3      4   1    1.0930e-01    2.8108e+05 ;     C2 - H
    3      5   1    1.8500e-01    1.7598e+05 ;     C2 - S
    3      6   1    1.4750e-01    2.6426e+05 ;     C2 - N
    3      7   1    1.5580e-01    2.3732e+05 ;     C2 - C3
    5     18   1    1.8392e-01    1.8067e+05 ;      S - C7
    6      9   1    1.4010e-01    3.3313e+05 ;      N - C4
    7      8   1    1.0950e-01    2.7874e+05 ;     C3 - H1
    7      9   1    1.5520e-01    2.4150e+05 ;     C3 - C4
    7     11   1    1.4330e-01    3.0091e+05 ;     C3 - N1
    9     10   1    1.2183e-01    5.3363e+05 ;     C4 - O
   11     28   1    1.3789e-01    3.5782e+05 ;     N1 - C10
   11     30   1    1.0129e-01    3.3740e+05 ;     N1 - H9
   12     13   1    1.2183e-01    5.3363e+05 ;     C5 - O1
   12     14   1    1.2183e-01    5.3363e+05 ;     C5 - O2
   15     16   1    1.0969e-01    2.7665e+05 ;     C6 - H2
   15     17   1    1.0969e-01    2.7665e+05 ;     C6 - H3
   15     21   1    1.4316e-01    2.5824e+05 ;     C6 - O3
   18     19   1    1.0969e-01    2.7665e+05 ;     C7 - H4
   18     20   1    1.0969e-01    2.7665e+05 ;     C7 - H5
   21     22   1    1.3584e-01    3.2702e+05 ;     O3 - C8
   22     23   1    1.5241e-01    2.6192e+05 ;     C8 - C9
   22     24   1    1.2183e-01    5.3363e+05 ;     C8 - O4
   23     25   1    1.0969e-01    2.7665e+05 ;     C9 - H6
   23     26   1    1.0969e-01    2.7665e+05 ;     C9 - H7
   23     27   1    1.0969e-01    2.7665e+05 ;     C9 - H8
   28     29   1    1.2183e-01    5.3363e+05 ;    C10 - O5
   28     31   1    1.4825e-01    2.9665e+05 ;    C10 - C11
   31     32   1    1.2874e-01    4.8735e+05 ;    C11 - N2
   31     38   1    1.4540e-01    3.2376e+05 ;    C11 - C13
   32     33   1    1.4015e-01    3.3999e+05 ;     N2 - O6
   33     34   1    1.4316e-01    2.5824e+05 ;     O6 - C12
   34     35   1    1.0969e-01    2.7665e+05 ;    C12 - H10
   34     36   1    1.0969e-01    2.7665e+05 ;    C12 - H11
   34     37   1    1.0969e-01    2.7665e+05 ;    C12 - H12
   38     39   1    1.3729e-01    4.1915e+05 ;    C13 - C14
   38     43   1    1.3694e-01    3.6911e+05 ;    C13 - N3
   39     40   1    1.0817e-01    2.9455e+05 ;    C14 - H13
   39     42   1    1.7562e-01    2.2242e+05 ;    C14 - S1
   41     42   1    1.7562e-01    2.2242e+05 ;    C15 - S1
   41     43   1    1.3172e-01    4.3965e+05 ;    C15 - N3
   41     44   1    1.3735e-01    3.6418e+05 ;    C15 - N4
   44     45   1    1.0121e-01    3.3857e+05 ;     N4 - H14
   44     46   1    1.0121e-01    3.3857e+05 ;     N4 - H15

[ pairs ]
;   ai     aj    funct
    1      4      1 ;      C - H
    1      5      1 ;      C - S
    1      7      1 ;      C - C3
    1     10      1 ;      C - O
    1     16      1 ;      C - H2
    1     17      1 ;      C - H3
    1     19      1 ;      C - H4
    1     20      1 ;      C - H5
    1     21      1 ;      C - O3
    2      3      1 ;     C1 - C2
    2      9      1 ;     C1 - C4
    2     13      1 ;     C1 - O1
    2     14      1 ;     C1 - O2
    2     22      1 ;     C1 - C8
```

| | | | |
|---|---|---|---|
| 3 | 10 | 1 ; | C2 - O |
| 3 | 19 | 1 ; | C2 - H4 |
| 3 | 20 | 1 ; | C2 - H5 |
| 3 | 28 | 1 ; | C2 - C10 |
| 3 | 30 | 1 ; | C2 - H9 |
| 4 | 8 | 1 ; | H - H1 |
| 4 | 9 | 1 ; | H - C4 |
| 4 | 11 | 1 ; | H - N1 |
| 4 | 18 | 1 ; | H - C7 |
| 5 | 8 | 1 ; | S - H1 |
| 5 | 9 | 1 ; | S - C4 |
| 5 | 11 | 1 ; | S - N1 |
| 5 | 15 | 1 ; | S - C6 |
| 6 | 8 | 1 ; | N - H1 |
| 6 | 11 | 1 ; | N - N1 |
| 6 | 13 | 1 ; | N - O1 |
| 6 | 14 | 1 ; | N - O2 |
| 6 | 15 | 1 ; | N - C6 |
| 6 | 18 | 1 ; | N - C7 |
| 7 | 18 | 1 ; | C3 - C7 |
| 7 | 29 | 1 ; | C3 - O5 |
| 7 | 31 | 1 ; | C3 - C11 |
| 8 | 10 | 1 ; | H1 - O |
| 8 | 28 | 1 ; | H1 - C10 |
| 8 | 30 | 1 ; | H1 - H9 |
| 9 | 28 | 1 ; | C4 - C10 |
| 9 | 30 | 1 ; | C4 - H9 |
| 10 | 11 | 1 ; | O - N1 |
| 11 | 32 | 1 ; | N1 - N2 |
| 11 | 38 | 1 ; | N1 - C13 |
| 12 | 3 | 1 ; | C5 - C2 |
| 12 | 9 | 1 ; | C5 - C4 |
| 12 | 15 | 1 ; | C5 - C6 |
| 12 | 18 | 1 ; | C5 - C7 |
| 15 | 19 | 1 ; | C6 - H4 |
| 15 | 20 | 1 ; | C6 - H5 |
| 15 | 23 | 1 ; | C6 - C9 |
| 15 | 24 | 1 ; | C6 - O4 |
| 16 | 18 | 1 ; | H2 - C7 |
| 16 | 22 | 1 ; | H2 - C8 |
| 17 | 18 | 1 ; | H3 - C7 |
| 17 | 22 | 1 ; | H3 - C8 |
| 18 | 21 | 1 ; | C7 - O3 |
| 21 | 25 | 1 ; | O3 - H6 |
| 21 | 26 | 1 ; | O3 - H7 |
| 21 | 27 | 1 ; | O3 - H8 |
| 24 | 25 | 1 ; | O4 - H6 |
| 24 | 26 | 1 ; | O4 - H7 |
| 24 | 27 | 1 ; | O4 - H8 |
| 28 | 33 | 1 ; | C10 - O6 |
| 28 | 39 | 1 ; | C10 - C14 |
| 28 | 43 | 1 ; | C10 - N3 |
| 29 | 30 | 1 ; | O5 - H9 |
| 29 | 32 | 1 ; | O5 - N2 |
| 29 | 38 | 1 ; | O5 - C13 |
| 30 | 31 | 1 ; | H9 - C11 |
| 31 | 34 | 1 ; | C11 - C12 |
| 31 | 40 | 1 ; | C11 - H13 |
| 31 | 41 | 1 ; | C11 - C15 |
| 31 | 42 | 1 ; | C11 - S1 |
| 32 | 35 | 1 ; | N2 - H10 |
| 32 | 36 | 1 ; | N2 - H11 |
| 32 | 37 | 1 ; | N2 - H12 |
| 32 | 39 | 1 ; | N2 - C14 |
| 32 | 43 | 1 ; | N2 - N3 |
| 33 | 38 | 1 ; | O6 - C13 |
| 38 | 44 | 1 ; | C13 - N4 |
| 39 | 44 | 1 ; | C14 - N4 |
| 40 | 41 | 1 ; | H13 - C15 |
| 40 | 43 | 1 ; | H13 - N3 |

```
   42    45     1 ;     S1 - H14
   42    46     1 ;     S1 - H15
   43    45     1 ;     N3 - H14
   43    46     1 ;     N3 - H15

[ angles ]
;   ai    aj    ak   funct   theta         cth
    1     2    15     1   1.2315e+02   5.3555e+02 ;     C - C1     - C6
    1     2    18     1   1.2315e+02   5.3555e+02 ;     C - C1     - C7
    1     6     3     1   1.1165e+02   5.4057e+02 ;     C - N      - C2
    1     6     9     1   1.3156e+02   5.1128e+02 ;     C - N      - C4
    1    12    13     1   1.2320e+02   5.7572e+02 ;     C - C5     - O1
    1    12    14     1   1.2320e+02   5.7572e+02 ;     C - C5     - O2
    2     1     6     1   1.1033e+02   5.9999e+02 ;    C1 - C      - N
    2     1    12     1   1.2042e+02   5.4810e+02 ;    C1 - C      - C5
    2    15    16     1   1.0996e+02   3.9413e+02 ;    C1 - C6     - H2
    2    15    17     1   1.0996e+02   3.9413e+02 ;    C1 - C6     - H3
    2    15    21     1   1.0856e+02   5.7321e+02 ;    C1 - C6     - O3
    2    18     5     1   1.0497e+02   5.2802e+02 ;    C1 - C7     - S
    2    18    19     1   1.0996e+02   3.9413e+02 ;    C1 - C7     - H4
    2    18    20     1   1.0996e+02   3.9413e+02 ;    C1 - C7     - H5
    3     5    18     1   9.4240e+01   5.1547e+02 ;    C2 - S      - C7
    3     6     9     1   9.4020e+01   5.9413e+02 ;    C2 - N      - C4
    3     7     8     1   1.1317e+02   3.7740e+02 ;    C2 - C3     - H1
    3     7     9     1   8.5160e+01   5.9496e+02 ;    C2 - C3     - C4
    3     7    11     1   1.1987e+02   5.3220e+02 ;    C2 - C3     - N1
    4     3     5     1   1.0966e+02   3.4811e+02 ;     H - C2     - S
    4     3     6     1   1.1454e+02   4.0334e+02 ;     H - C2     - N
    4     3     7     1   1.1679e+02   3.7154e+02 ;     H - C2     - C3
    5     3     6     1   1.0514e+02   5.4141e+02 ;     S - C2     - N
    5     3     7     1   1.1827e+02   4.9120e+02 ;     S - C2     - C3
    5    18    19     1   1.0876e+02   3.5229e+02 ;     S - C7     - H4
    5    18    20     1   1.0876e+02   3.5229e+02 ;     S - C7     - H5
    6     1    12     1   1.1853e+02   5.5229e+02 ;     N - C      - C5
    6     3     7     1   8.8640e+01   6.1170e+02 ;     N - C2     - C3
    6     9     7     1   9.1680e+01   6.1588e+02 ;     N - C4     - C3
    6     9    10     1   1.3306e+02   5.9078e+02 ;     N - C4     - O
    7     9    10     1   1.3523e+02   5.3053e+02 ;    C3 - C4     - O
    7    11    28     1   1.2171e+02   5.3388e+02 ;    C3 - N1     - C10
    7    11    30     1   1.1885e+02   3.8828e+02 ;    C3 - N1     - H9
    8     7     9     1   1.1297e+02   3.7907e+02 ;    H1 - C3     - C4
    8     7    11     1   1.0799e+02   4.2593e+02 ;    H1 - C3     - N1
    9     7    11     1   1.1738e+02   5.3974e+02 ;    C4 - C3     - N1
   11    28    29     1   1.2305e+02   6.2091e+02 ;    N1 - C10    - O5
   11    28    31     1   1.1522e+02   5.6819e+02 ;    N1 - C10    - C11
   13    12    14     1   1.3025e+02   6.5187e+02 ;    O1 - C5     - O2
   15     2    18     1   1.1565e+02   5.2635e+02 ;    C6 - C1     - C7
   15    21    22     1   1.1598e+02   5.2969e+02 ;    C6 - O3     - C8
   16    15    17     1   1.0846e+02   3.2803e+02 ;    H2 - C6     - H3
   16    15    21     1   1.0978e+02   4.2509e+02 ;    H2 - C6     - O3
   17    15    21     1   1.0978e+02   4.2509e+02 ;    H3 - C6     - O3
   19    18    20     1   1.0846e+02   3.2803e+02 ;    H4 - C7     - H5
   21    22    23     1   1.1072e+02   5.7656e+02 ;    O3 - C8     - C9
   21    22    24     1   1.2325e+02   6.3011e+02 ;    O3 - C8     - O4
   22    23    25     1   1.0877e+02   3.9246e+02 ;    C8 - C9     - H6
   22    23    26     1   1.0877e+02   3.9246e+02 ;    C8 - C9     - H7
   22    23    27     1   1.0877e+02   3.9246e+02 ;    C8 - C9     - H8
   23    22    24     1   1.2320e+02   5.6400e+02 ;    C9 - C8     - O4
   25    23    26     1   1.0758e+02   3.2970e+02 ;    H6 - C9     - H7
   25    23    27     1   1.0758e+02   3.2970e+02 ;    H6 - C9     - H8
   26    23    27     1   1.0758e+02   3.2970e+02 ;    H7 - C9     - H8
   28    11    30     1   1.1755e+02   4.0417e+02 ;   C10 - N1     - H9
   28    31    32     1   1.1441e+02   5.8492e+02 ;   C10 - C11    - N2
   28    31    38     1   1.1782e+02   5.3555e+02 ;   C10 - C11    - C13
   29    28    31     1   1.2320e+02   5.7572e+02 ;    O5 - C10    - C11
   31    32    33     1   1.1279e+02   5.9580e+02 ;   C11 - N2     - O6
   31    38    39     1   1.2805e+02   5.3304e+02 ;   C11 - C13    - C14
   31    38    43     1   1.2110e+02   5.6233e+02 ;   C11 - C13    - N3
   32    31    38     1   1.2096e+02   5.7572e+02 ;    N2 - C11    - C13
   32    33    34     1   1.0923e+02   5.5145e+02 ;    N2 - O6     - C12
```

```
   33    34    35     1    1.0978e+02    4.2509e+02 ;     O6 - C12    - H10
   33    34    36     1    1.0978e+02    4.2509e+02 ;     O6 - C12    - H11
   33    34    37     1    1.0978e+02    4.2509e+02 ;     O6 - C12    - H12
   35    34    36     1    1.0846e+02    3.2803e+02 ;    H10 - C12    - H11
   35    34    37     1    1.0846e+02    3.2803e+02 ;    H10 - C12    - H12
   36    34    37     1    1.0846e+02    3.2803e+02 ;    H11 - C12    - H12
   38    39    40     1    1.2848e+02    3.9581e+02 ;    C13 - C14    - H13
   38    39    42     1    1.1155e+02    5.4225e+02 ;    C13 - C14    - S1
   38    43    41     1    1.0549e+02    6.0082e+02 ;    C13 - N3     - C15
   39    38    43     1    1.1165e+02    6.0417e+02 ;    C14 - C13    - N3
   39    42    41     1    9.0240e+01    5.5396e+02 ;    C14 - S1     - C15
   40    39    42     1    1.1997e+02    3.5229e+02 ;    H13 - C14    - S1
   41    44    45     1    1.1563e+02    4.0920e+02 ;    C15 - N4     - H14
   41    44    46     1    1.1563e+02    4.0920e+02 ;    C15 - N4     - H15
   42    41    43     1    1.1451e+02    5.5229e+02 ;     S1 - C15    - N3
   42    41    44     1    1.2181e+02    5.3220e+02 ;     S1 - C15    - N4
   43    41    44     1    1.2065e+02    6.0584e+02 ;     N3 - C15    - N4
   45    44    46     1    1.1512e+02    3.3556e+02 ;    H14 - N4     - H15

[ dihedrals ] ; propers
; treated as RBs in GROMACS to use combine multiple AMBER torsions per quartet
;    i     j     k     l   func    C0         C1         C2         C3         C4         C5
     1     2    15    16     3    0.00000    0.00000    0.00000    0.00000    0.00000    0.00000 ;     C-    C1-    C6-    H2
     1     2    15    17     3    0.00000    0.00000    0.00000    0.00000    0.00000    0.00000 ;     C-    C1-    C6-    H3
     1     2    15    21     3    0.00000    0.00000    0.00000    0.00000    0.00000    0.00000 ;     C-    C1-    C6-    O3
     1     2    18     5     3    0.00000    0.00000    0.00000    0.00000    0.00000    0.00000 ;     C-    C1-    C7-    S
     1     2    18    19     3    0.00000    0.00000    0.00000    0.00000    0.00000    0.00000 ;     C-    C1-    C7-    H4
     1     2    18    20     3    0.00000    0.00000    0.00000    0.00000    0.00000    0.00000 ;     C-    C1-    C7-    H5
     1     6     3     4     3    0.00000    0.00000    0.00000    0.00000    0.00000    0.00000 ;     C-    N-    C2-    H
     1     6     3     5     3    0.00000    0.00000    0.00000    0.00000    0.00000    0.00000 ;     C-    N-    C2-    S
     1     6     3     7     3    0.00000    0.00000    0.00000    0.00000    0.00000    0.00000 ;     C-    N-    C2-    C3
     1     6     9     7     3   20.92000    0.00000  -20.92000    0.00000    0.00000    0.00000 ;     C-    N-    C4-    C3
     1     6     9    10     3   20.92000    0.00000  -20.92000    0.00000    0.00000    0.00000 ;     C-    N-    C4-    O
     2     1     6     3     3   13.80720    0.00000  -13.80720    0.00000    0.00000    0.00000 ;    C1-    C-    N-    C2
     2     1     6     9     3   10.46000    5.02080   -5.43920    0.00000    0.00000    0.00000 ;    C1-    C-    N-    C4
     2     1    12    13     3   18.20040    0.00000  -18.20040    0.00000    0.00000    0.00000 ;    C1-    C-    C5-    O1
     2     1    12    14     3   18.20040    0.00000  -18.20040    0.00000    0.00000    0.00000 ;    C1-    C-    C5-    O2
     2    15    21    22     3    1.60387    4.81160    0.00000   -6.41547    0.00000    0.00000 ;    C1-    C6-    O3-    C8
     2    18     5     3     3    1.39467    4.18400    0.00000   -5.57867    0.00000    0.00000 ;    C1-    C7-    S-    C2
     3     5    18    19     3    1.39467    4.18400    0.00000   -5.57867    0.00000    0.00000 ;    C2-    S-    C7-    H4
     3     5    18    20     3    1.39467    4.18400    0.00000   -5.57867    0.00000    0.00000 ;    C2-    S-    C7-    H5
     3     6     9     7     3   20.92000    0.00000  -20.92000    0.00000    0.00000    0.00000 ;    C2-    N-    C4-    C3
     3     6     9    10     3   20.92000    0.00000  -20.92000    0.00000    0.00000    0.00000 ;    C2-    N-    C4-    O
     3     7     9     6     3    0.00000    0.00000    0.00000    0.00000    0.00000    0.00000 ;    C2-    C3-    C4-    N
     3     7     9    10     3    0.00000    0.00000    0.00000    0.00000    0.00000    0.00000 ;    C2-    C3-    C4-    O
     3     7    11    28     3    0.00000    0.00000    0.00000    0.00000    0.00000    0.00000 ;    C2-    C3-    N1-    C10
     3     7    11    30     3    0.00000    0.00000    0.00000    0.00000    0.00000    0.00000 ;    C2-    C3-    N1-    H9
     4     3     5    18     3    1.39467    4.18400    0.00000   -5.57867    0.00000    0.00000 ;    H-    C2-    S-    C7
     4     3     6     9     3    0.00000    0.00000    0.00000    0.00000    0.00000    0.00000 ;    H-    C2-    N-    C4
     4     3     7     8     3    0.65084    1.95253    0.00000   -2.60338    0.00000    0.00000 ;    H-    C2-    C3-    H1
     4     3     7     9     3    0.65084    1.95253    0.00000   -2.60338    0.00000    0.00000 ;    H-    C2-    C3-    C4
     4     3     7    11     3    0.65084    1.95253    0.00000   -2.60338    0.00000    0.00000 ;    H-    C2-    C3-    N1
     5     3     6     9     3    0.00000    0.00000    0.00000    0.00000    0.00000    0.00000 ;    S-    C2-    N-    C4
     5     3     7     8     3    0.65084    1.95253    0.00000   -2.60338    0.00000    0.00000 ;    S-    C2-    C3-    H1
     5     3     7     9     3    0.65084    1.95253    0.00000   -2.60338    0.00000    0.00000 ;    S-    C2-    C3-    C4
     5     3     7    11     3    0.65084    1.95253    0.00000   -2.60338    0.00000    0.00000 ;    S-    C2-    C3-    N1
     5    18     2    15     3    0.00000    0.00000    0.00000    0.00000    0.00000    0.00000 ;    S-    C7-    C1-    C6
     6     1     2    15     3   55.64720    0.00000  -55.64720    0.00000    0.00000    0.00000 ;    N-    C-    C1-    C6
     6     1     2    18     3   55.64720    0.00000  -55.64720    0.00000    0.00000    0.00000 ;    N-    C-    C1-    C7
     6     1    12    13     3   18.20040    0.00000  -18.20040    0.00000    0.00000    0.00000 ;    N-    C-    C5-    O1
     6     1    12    14     3   18.20040    0.00000  -18.20040    0.00000    0.00000    0.00000 ;    N-    C-    C5-    O2
     6     3     5    18     3    1.39467    4.18400    0.00000   -5.57867    0.00000    0.00000 ;    N-    C2-    S-    C7
     6     3     7     8     3    0.65084    1.95253    0.00000   -2.60338    0.00000    0.00000 ;    N-    C2-    C3-    H1
     6     3     7     9     3    0.65084    1.95253    0.00000   -2.60338    0.00000    0.00000 ;    N-    C2-    C3-    C4
     6     3     7    11     3    0.65084    1.95253    0.00000   -2.60338    0.00000    0.00000 ;    N-    C2-    C3-    N1
     6     9     7     8     3    0.00000    0.00000    0.00000    0.00000    0.00000    0.00000 ;    N-    C4-    C3-    H1
     6     9     7    11     3    0.00000    0.00000    0.00000    0.00000    0.00000    0.00000 ;    N-    C4-    C3-    N1
     7     3     5    18     3    1.39467    4.18400    0.00000   -5.57867    0.00000    0.00000 ;    C3-    C2-    S-    C7
     7     3     6     9     3    0.00000    0.00000    0.00000    0.00000    0.00000    0.00000 ;    C3-    C2-    N-    C4
     7    11    28    29     3   20.92000    0.00000  -20.92000    0.00000    0.00000    0.00000 ;    C3-    N1-    C10-    O5
```

```
  7   11   28   31    3  20.92000   0.00000  -20.92000   0.00000   0.00000   0.00000 ;   C3-   N1-  C10-  C11
  8    7    9   10    3   0.00000   0.00000    0.00000   0.00000   0.00000   0.00000 ;   H1-   C3-   C4-   O
  8    7   11   28    3   0.00000   0.00000    0.00000   0.00000   0.00000   0.00000 ;   H1-   C3-   N1-  C10
  8    7   11   30    3   0.00000   0.00000    0.00000   0.00000   0.00000   0.00000 ;   H1-   C3-   N1-   H9
  9    7   11   28    3   0.00000   0.00000    0.00000   0.00000   0.00000   0.00000 ;   C4-   C3-   N1-  C10
  9    7   11   30    3   0.00000   0.00000    0.00000   0.00000   0.00000   0.00000 ;   C4-   C3-   N1-   H9
 10    9    7   11    3   0.00000   0.00000    0.00000   0.00000   0.00000   0.00000 ;   O-   C4-   C3-   N1
 11   28   31   32    3  18.20040   0.00000  -18.20040   0.00000   0.00000   0.00000 ;   N1-  C10-  C11-   N2
 11   28   31   38    3  18.20040   0.00000  -18.20040   0.00000   0.00000   0.00000 ;   N1-  C10-  C11-  C13
 12    1    2   15    3  55.64720   0.00000  -55.64720   0.00000   0.00000   0.00000 ;   C5-   C-   C1-   C6
 12    1    2   18    3  55.64720   0.00000  -55.64720   0.00000   0.00000   0.00000 ;   C5-   C-   C1-   C7
 12    1    6    3    3  13.80720   0.00000  -13.80720   0.00000   0.00000   0.00000 ;   C5-   C-   N-   C2
 12    1    6    9    3  10.46000   5.02080   -5.43920   0.00000   0.00000   0.00000 ;   C5-   C-   N-   C4
 15    2   18   19    3   0.00000   0.00000    0.00000   0.00000   0.00000   0.00000 ;   C6-   C1-   C7-   H4
 15    2   18   20    3   0.00000   0.00000    0.00000   0.00000   0.00000   0.00000 ;   C6-   C1-   C7-   H5
 15   21   22   23    3  27.40520  14.43480  -22.59360 -19.24640   0.00000   0.00000 ;   C6-   O3-   C8-   C9
 15   21   22   24    3  28.45120   5.85760  -22.59360   0.00000   0.00000   0.00000 ;   C6-   O3-   C8-   O4
 16   15    2   18    3   0.00000   0.00000    0.00000   0.00000   0.00000   0.00000 ;   H2-   C6-   C1-   C7
 16   15   21   22    3   1.60387   4.81160    0.00000  -6.41547   0.00000   0.00000 ;   H2-   C6-   O3-   C8
 17   15    2   18    3   0.00000   0.00000    0.00000   0.00000   0.00000   0.00000 ;   H3-   C6-   C1-   C7
 17   15   21   22    3   1.60387   4.81160    0.00000  -6.41547   0.00000   0.00000 ;   H3-   C6-   O3-   C8
 18    2   15   21    3   0.00000   0.00000    0.00000   0.00000   0.00000   0.00000 ;   C7-   C1-   C6-   O3
 21   22   23   25    3   0.00000   0.00000    0.00000   0.00000   0.00000   0.00000 ;   O3-   C8-   C9-   H6
 21   22   23   26    3   0.00000   0.00000    0.00000   0.00000   0.00000   0.00000 ;   O3-   C8-   C9-   H7
 21   22   23   27    3   0.00000   0.00000    0.00000   0.00000   0.00000   0.00000 ;   O3-   C8-   C9-   H8
 24   22   23   25    3   3.68192  -4.35136    0.00000   1.33888   0.00000   0.00000 ;   O4-   C8-   C9-   H6
 24   22   23   26    3   3.68192  -4.35136    0.00000   1.33888   0.00000   0.00000 ;   O4-   C8-   C9-   H7
 24   22   23   27    3   3.68192  -4.35136    0.00000   1.33888   0.00000   0.00000 ;   O4-   C8-   C9-   H8
 28   31   32   33    3   6.69440   0.00000   -6.69440   0.00000   0.00000   0.00000 ;  C10-  C11-   N2-   O6
 28   31   38   39    3   8.36800   0.00000   -8.36800   0.00000   0.00000   0.00000 ;  C10-  C11-  C13-  C14
 28   31   38   43    3   8.36800   0.00000   -8.36800   0.00000   0.00000   0.00000 ;  C10-  C11-  C13-   N3
 29   28   11   30    3  29.28800  -8.36800  -20.92000   0.00000   0.00000   0.00000 ;   O5-  C10-   N1-   H9
 29   28   31   32    3  18.20040   0.00000  -18.20040   0.00000   0.00000   0.00000 ;   O5-  C10-  C11-   N2
 29   28   31   38    3  18.20040   0.00000  -18.20040   0.00000   0.00000   0.00000 ;   O5-  C10-  C11-  C13
 30   11   28   31    3  20.92000   0.00000  -20.92000   0.00000   0.00000   0.00000 ;   H9-   N1-  C10-  C11
 31   32   33   34    3  25.10400   0.00000  -25.10400   0.00000   0.00000   0.00000 ;  C11-   N2-   O6-  C12
 31   38   39   40    3  33.47200   0.00000  -33.47200   0.00000   0.00000   0.00000 ;  C11-  C13-  C14-  H13
 31   38   39   42    3  33.47200   0.00000  -33.47200   0.00000   0.00000   0.00000 ;  C11-  C13-  C14-   S1
 31   38   43   41    3  39.74800   0.00000  -39.74800   0.00000   0.00000   0.00000 ;  C11-  C13-   N3-  C15
 32   31   38   39    3   8.36800   0.00000   -8.36800   0.00000   0.00000   0.00000 ;   N2-  C11-  C13-  C14
 32   31   38   43    3   8.36800   0.00000   -8.36800   0.00000   0.00000   0.00000 ;   N2-  C11-  C13-   N3
 32   33   34   35    3   1.60387   4.81160    0.00000  -6.41547   0.00000   0.00000 ;   N2-   O6-  C12-  H10
 32   33   34   36    3   1.60387   4.81160    0.00000  -6.41547   0.00000   0.00000 ;   N2-   O6-  C12-  H11
 32   33   34   37    3   1.60387   4.81160    0.00000  -6.41547   0.00000   0.00000 ;   N2-   O6-  C12-  H12
 33   32   31   38    3   6.69440   0.00000   -6.69440   0.00000   0.00000   0.00000 ;   O6-   N2-  C11-  C13
 38   39   42   41    3   9.20480   0.00000   -9.20480   0.00000   0.00000   0.00000 ;  C13-  C14-   S1-  C15
 38   43   41   42    3  39.74800   0.00000  -39.74800   0.00000   0.00000   0.00000 ;  C13-   N3-  C15-   S1
 38   43   41   44    3  39.74800   0.00000  -39.74800   0.00000   0.00000   0.00000 ;  C13-   N3-  C15-   N4
 39   38   43   41    3  39.74800   0.00000  -39.74800   0.00000   0.00000   0.00000 ;  C14-  C13-   N3-  C15
 39   42   41   43    3   9.20480   0.00000   -9.20480   0.00000   0.00000   0.00000 ;  C14-   S1-  C15-   N3
 39   42   41   44    3   9.20480   0.00000   -9.20480   0.00000   0.00000   0.00000 ;  C14-   S1-  C15-   N4
 40   39   38   43    3  33.47200   0.00000  -33.47200   0.00000   0.00000   0.00000 ;  H13-  C14-  C13-   N3
 40   39   42   41    3   9.20480   0.00000   -9.20480   0.00000   0.00000   0.00000 ;  H13-  C14-   S1-  C15
 42   39   38   43    3  33.47200   0.00000  -33.47200   0.00000   0.00000   0.00000 ;   S1-  C14-  C13-   N3
 42   41   44   45    3   8.78640   0.00000   -8.78640   0.00000   0.00000   0.00000 ;   S1-  C15-   N4-  H14
 42   41   44   46    3   8.78640   0.00000   -8.78640   0.00000   0.00000   0.00000 ;   S1-  C15-   N4-  H15
 43   41   44   45    3   8.78640   0.00000   -8.78640   0.00000   0.00000   0.00000 ;   N3-  C15-   N4-  H14
 43   41   44   46    3   8.78640   0.00000   -8.78640   0.00000   0.00000   0.00000 ;   N3-  C15-   N4-  H15

[ dihedrals ] ; impropers
; treated as propers in GROMACS to use correct AMBER analytical function
;   i    j    k    l  func  phase     kd    pn
    1    2   18   15     1  180.00   4.60240   2 ;    C-   C1-   C7-   C6
    1   13   12   14     1  180.00   4.60240   2 ;    C-   O1-   C5-   O2
    6    1    2   12     1  180.00   4.60240   2 ;    N-   C-   C1-   C5
    7    6    9   10     1  180.00  43.93200   2 ;   C3-   N-   C4-   O
    9    1    6    3     1  180.00   4.60240   2 ;   C4-   C-   N-   C2
   23   24   22   21     1  180.00  43.93200   2 ;   C9-   O4-   C8-   O3
   28    7   11   30     1  180.00   4.60240   2 ;  C10-   C3-   N1-   H9
   28   38   31   32     1  180.00   4.60240   2 ;  C10-  C13-  C11-   N2
```

| | | | | | | | | | | | |
|---|---|---|---|---|---|---|---|---|---|---|---|
| 31 | 11 | 28 | 29 | 1 | 180.00 | 43.93200 | 2 ; | C11- | N1- | C10- | O5 |
| 38 | 40 | 39 | 42 | 1 | 180.00 | 4.60240 | 2 ; | C13- | H13- | C14- | S1 |
| 39 | 31 | 38 | 43 | 1 | 180.00 | 4.60240 | 2 ; | C14- | C11- | C13- | N3 |
| 41 | 45 | 44 | 46 | 1 | 180.00 | 4.60240 | 2 ; | C15- | H14- | N4- | H15 |
| 43 | 44 | 41 | 42 | 1 | 180.00 | 4.60240 | 2 ; | N3- | N4- | C15- | S1 |

**SCX.gro**

```
70SCX   N  710  3.863  4.146  3.708
70SCX   H  711  3.949  4.125  3.758
70SCX  CA  712  3.871  4.207  3.573
70SCX  HA  713  3.831  4.308  3.582
70SCX  CB  714  4.018  4.219  3.527
70SCX HB1  715  4.019  4.263  3.428
70SCX HB2  716  4.062  4.119  3.520
70SCX  OG  717  4.098  4.303  3.614
70SCX  C2  718  4.159  4.248  3.724
70SCX  O1  719  4.145  4.130  3.753
70SCX  C3  720  4.267  4.339  3.788
70SCX  C4  721  4.461  4.211  3.561
70SCX  C5  722  4.605  4.234  3.579
70SCX  C6  723  4.362  4.402  3.681
70SCX  H4  724  4.314  4.493  3.644
70SCX  S2  725  4.524  4.453  3.749
70SCX  N1  726  4.382  4.315  3.564
70SCX  H5  727  4.213  4.421  3.837
70SCX  N2  728  4.343  4.264  3.891
70SCX  C7  729  4.403  4.086  3.492
70SCX  O2  730  4.464  3.979  3.489
70SCX  O3  731  4.296  4.092  3.432
70SCX  C8  732  4.699  4.171  3.505
70SCX  H6  733  4.805  4.195  3.516
70SCX  H7  734  4.674  4.097  3.429
70SCX  C9  735  4.652  4.344  3.674
70SCX  H8  736  4.722  4.409  3.620
70SCX  H9  737  4.710  4.300  3.755
70SCX C10  738  4.389  4.316  4.010
70SCX  O4  739  4.362  4.431  4.042
70SCX H10  740  4.354  4.164  3.881
70SCX C11  741  4.470  4.223  4.097
70SCX  N3  742  4.537  4.125  4.042
70SCX  O5  743  4.611  4.157  3.928
70SCX C12  744  4.732  4.079  3.927
70SCX H11  745  4.798  4.113  4.008
70SCX H12  746  4.708  3.974  3.943
70SCX H13  747  4.784  4.091  3.832
70SCX C13  748  4.469  4.237  4.243
70SCX C14  749  4.394  4.324  4.319
70SCX H14  750  4.322  4.397  4.283
70SCX C15  751  4.532  4.179  4.447
70SCX  S1  752  4.423  4.309  4.490
70SCX  N4  753  4.547  4.152  4.321
70SCX  N5  754  4.607  4.107  4.534
70SCX H15  755  4.684  4.059  4.491
70SCX H16  756  4.616  4.141  4.629
70SCX   C  757  3.789  4.138  3.462
70SCX   O  758  3.778  4.191  3.351
```

**SCX.top**

```
[ moleculetype ]
; Name            nrexcl
SCX     3
```

```
[ defaults ]
; nbfunc        comb-rule       gen-pairs       fudgeLJ fudgeQQ
1               2               yes             0.5     0.8333

[ atomtypes ]
;name   bond_type     mass     charge   ptype   sigma         epsilon       Amb
 n        n          0.00000  0.00000   A     3.25000e-01   7.11280e-01 ; 1.82  0.1700
 hn       hn         0.00000  0.00000   A     1.06908e-01   6.56888e-02 ; 0.60  0.0157
 c3       c3         0.00000  0.00000   A     3.39967e-01   4.57730e-01 ; 1.91  0.1094
 h1       h1         0.00000  0.00000   A     2.47135e-01   6.56888e-02 ; 1.39  0.0157
 os       os         0.00000  0.00000   A     3.00001e-01   7.11280e-01 ; 1.68  0.1700
 c        c          0.00000  0.00000   A     3.39967e-01   3.59824e-01 ; 1.91  0.0860
 o        o          0.00000  0.00000   A     2.95992e-01   8.78640e-01 ; 1.66  0.2100
 ce       ce         0.00000  0.00000   A     3.39967e-01   3.59824e-01 ; 1.91  0.0860
 h2       h2         0.00000  0.00000   A     2.29317e-01   6.56888e-02 ; 1.29  0.0157
 ss       ss         0.00000  0.00000   A     3.56359e-01   1.04600e+00 ; 2.00  0.2500
 n2       n2         0.00000  0.00000   A     3.25000e-01   7.11280e-01 ; 1.82  0.1700
 c2       c2         0.00000  0.00000   A     3.39967e-01   3.59824e-01 ; 1.91  0.0860
 ha       ha         0.00000  0.00000   A     2.59964e-01   6.27600e-02 ; 1.46  0.0150
 cc       cc         0.00000  0.00000   A     3.39967e-01   3.59824e-01 ; 1.91  0.0860
 cd       cd         0.00000  0.00000   A     3.39967e-01   3.59824e-01 ; 1.91  0.0860
 h4       h4         0.00000  0.00000   A     2.51055e-01   6.27600e-02 ; 1.41  0.0150
 nc       nc         0.00000  0.00000   A     3.25000e-01   7.11280e-01 ; 1.82  0.1700
 nh       nh         0.00000  0.00000   A     3.25000e-01   7.11280e-01 ; 1.82  0.1700
 hc       hc         0.00000  0.00000   A     2.64953e-01   6.56888e-02 ; 1.49  0.0157

[ atoms ]
;   nr  type  resi  res  atom  cgnr     charge      mass       ; qtot   bond_type
710     N       70      SCX     N       710     -0.4157         14.01
711     H       70      SCX     H       711     0.2719          1.008
712     CT      70      SCX     CA      712     -0.0349         12.01
713     H1      70      SCX     HA      713     0.0853          1.008
714     CT      70      SCX     CB      714     0.1414          12.01
715     H1      70      SCX     HB1     715     0.085205        1.008
716     H1      70      SCX     HB2     716     0.085205        1.008
717     os      70      SCX     OG      717     -0.3949         16
718     c       70      SCX     C2      718     0.621101        12.01
719     o       70      SCX     O1      719     -0.572001       16
720     c3      70      SCX     C3      720     0.0717          12.01
721     ce      70      SCX     C4      721     0.4466          12.01
722     ce      70      SCX     C5      722     -0.2338         12.01
723     c3      70      SCX     C6      723     0.2556          12.01
724     h2      70      SCX     H4      724     0.0987          1.008
725     ss      70      SCX     S2      725     -0.4452         32.06
726     n2      70      SCX     N1      726     -0.609901       14.01
727     h1      70      SCX     H5      727     0.1287          1.008
728     n       70      SCX     N2      728     -0.548901       14.01
729     c       70      SCX     C7      729     0.855301        12.01
730     o       70      SCX     O2      730     -0.806301       16
731     o       70      SCX     O3      731     -0.806301       16
732     c2      70      SCX     C8      732     -0.156          12.01
733     ha      70      SCX     H6      733     0.128           1.008
734     ha      70      SCX     H7      734     0.128           1.008
735     c3      70      SCX     C9      735     0.0599          12.01
736     h1      70      SCX     H8      736     0.0762          1.008
737     h1      70      SCX     H9      737     0.0762          1.008
738     c       70      SCX     C10     738     0.568801        12.01
739     o       70      SCX     O4      739     -0.575101       16
740     hn      70      SCX     H10     740     0.3395          1.008
741     ce      70      SCX     C11     741     0.2883          12.01
742     n2      70      SCX     N3      742     -0.3782         14.01
743     os      70      SCX     O5      743     -0.1793         16
744     c3      70      SCX     C12     744     0.1337          12.01
745     h1      70      SCX     H11     745     0.0477          1.008
746     h1      70      SCX     H12     746     0.0477          1.008
747     h1      70      SCX     H13     747     0.0477          1.008
748     cc      70      SCX     C13     748     0.317           12.01
749     cd      70      SCX     C14     749     -0.2464         12.01
750     h4      70      SCX     H14     750     0.217           1.008
751     cd      70      SCX     C15     751     0.532301        12.01
```

| | | | | | | | |
|---|---|---|---|---|---|---|---|
| 752 | ss | 70 | SCX | S1 | 752 | -0.0223 | 32.06 |
| 753 | nc | 70 | SCX | N4 | 753 | -0.640001 | 14.01 |
| 754 | nh | 70 | SCX | N5 | 754 | -0.871502 | 14.01 |
| 755 | hn | 70 | SCX | H15 | 755 | 0.3763 | 1.008 |
| 756 | hn | 70 | SCX | H16 | 756 | 0.3763 | 1.008 |
| 757 | C | 70 | SCX | C | 757 | 0.5973 | 12.01 |
| 758 | O | 70 | SCX | O | 758 | -0.5679 | 16 |

[ bonds ]

; ai aj funct r k

| | | | | | | | | |
|---|---|---|---|---|---|---|---|---|
| 714 | 717 | 1 | 1.4316E-01 | 2.5824E+05 | ; | C1 | - | O |
| 717 | 718 | 1 | 1.3584E-01 | 3.2702E+05 | ; | O | - | C2 |
| 718 | 719 | 1 | 1.2183E-01 | 5.3363E+05 | ; | C2 | - | O1 |
| 718 | 720 | 1 | 1.5241E-01 | 2.6192E+05 | ; | C2 | - | C3 |
| 720 | 723 | 1 | 1.5375E-01 | 2.5179E+05 | ; | C3 | - | C6 |
| 720 | 727 | 1 | 1.0969E-01 | 2.7665E+05 | ; | C3 | - | H5 |
| 720 | 728 | 1 | 1.4619E-01 | 2.7506E+05 | ; | C3 | - | N2 |
| 721 | 722 | 1 | 1.4574E-01 | 3.2033E+05 | ; | C4 | - | C5 |
| 721 | 726 | 1 | 1.2874E-01 | 4.8735E+05 | ; | C4 | - | N1 |
| 721 | 729 | 1 | 1.4825E-01 | 2.9665E+05 | ; | C4 | - | C7 |
| 722 | 732 | 1 | 1.3461E-01 | 4.5798E+05 | ; | C5 | - | C8 |
| 722 | 735 | 1 | 1.5159E-01 | 2.6836E+05 | ; | C5 | - | C9 |
| 723 | 724 | 1 | 1.0961E-01 | 2.7757E+05 | ; | C6 | - | H4 |
| 723 | 725 | 1 | 1.8392E-01 | 1.8067E+05 | ; | C6 | - | S |
| 723 | 726 | 1 | 1.4661E-01 | 2.7154E+05 | ; | C6 | - | N1 |
| 725 | 735 | 1 | 1.8392E-01 | 1.8067E+05 | ; | S | - | C9 |
| 728 | 738 | 1 | 1.3789E-01 | 3.5782E+05 | ; | N2 | - | C10 |
| 728 | 740 | 1 | 1.0129E-01 | 3.3740E+05 | ; | N2 | - | H10 |
| 729 | 730 | 1 | 1.2183E-01 | 5.3363E+05 | ; | C7 | - | O2 |
| 729 | 731 | 1 | 1.2183E-01 | 5.3363E+05 | ; | C7 | - | O3 |
| 732 | 733 | 1 | 1.0879E-01 | 2.8711E+05 | ; | C8 | - | H6 |
| 732 | 734 | 1 | 1.0879E-01 | 2.8711E+05 | ; | C8 | - | H7 |
| 735 | 736 | 1 | 1.0969E-01 | 2.7665E+05 | ; | C9 | - | H8 |
| 735 | 737 | 1 | 1.0969E-01 | 2.7665E+05 | ; | C9 | - | H9 |
| 738 | 739 | 1 | 1.2183E-01 | 5.3363E+05 | ; | C10 | - | O4 |
| 738 | 741 | 1 | 1.4825E-01 | 2.9665E+05 | ; | C10 | - | C11 |
| 741 | 742 | 1 | 1.2874E-01 | 4.8735E+05 | ; | C11 | - | N3 |
| 741 | 748 | 1 | 1.4540E-01 | 3.2376E+05 | ; | C11 | - | C13 |
| 742 | 743 | 1 | 1.4015E-01 | 3.3999E+05 | ; | N3 | - | O5 |
| 743 | 744 | 1 | 1.4316E-01 | 2.5824E+05 | ; | O5 | - | C12 |
| 744 | 745 | 1 | 1.0969E-01 | 2.7665E+05 | ; | C12 | - | H11 |
| 744 | 746 | 1 | 1.0969E-01 | 2.7665E+05 | ; | C12 | - | H12 |
| 744 | 747 | 1 | 1.0969E-01 | 2.7665E+05 | ; | C12 | - | H13 |
| 748 | 749 | 1 | 1.3729E-01 | 4.1915E+05 | ; | C13 | - | C14 |
| 748 | 753 | 1 | 1.3694E-01 | 3.6911E+05 | ; | C13 | - | N4 |
| 749 | 750 | 1 | 1.0817E-01 | 2.9455E+05 | ; | C14 | - | H14 |
| 749 | 752 | 1 | 1.7562E-01 | 2.2242E+05 | ; | C14 | - | S1 |
| 751 | 752 | 1 | 1.7562E-01 | 2.2242E+05 | ; | C15 | - | S1 |
| 751 | 753 | 1 | 1.3172E-01 | 4.3965E+05 | ; | C15 | - | N4 |
| 751 | 754 | 1 | 1.3735E-01 | 3.6418E+05 | ; | C15 | - | N5 |
| 754 | 755 | 1 | 1.0121E-01 | 3.3857E+05 | ; | N5 | - | H15 |
| 754 | 756 | 1 | 1.0121E-01 | 3.3857E+05 | ; | N5 | - | H16 |

[ pairs ]

; ai aj funct

| | | | | | | |
|---|---|---|---|---|---|---|
| 710 | 715 | 1 | ; | N | - | H2 |
| 710 | 716 | 1 | ; | N | - | H3 |
| 710 | 717 | 1 | ; | N | - | O |
| 710 | 758 | 1 | ; | N | - | O6 |
| 711 | 713 | 1 | ; | H | - | H1 |
| 711 | 714 | 1 | ; | H | - | C1 |
| 711 | 757 | 1 | ; | H | - | C16 |
| 712 | 718 | 1 | ; | C | - | C2 |
| 713 | 715 | 1 | ; | H1 | - | H2 |
| 713 | 716 | 1 | ; | H1 | - | H3 |
| 713 | 717 | 1 | ; | H1 | - | O |
| 713 | 758 | 1 | ; | H1 | - | O6 |
| 714 | 719 | 1 | ; | C1 | - | O1 |
| 714 | 720 | 1 | ; | C1 | - | C3 |
| 714 | 758 | 1 | ; | C1 | - | O6 |

| | | | | | | |
|---|---|---|---|---|---|---|
| 715 | 718 | 1 | ; | H2 | - | C2 |
| 715 | 757 | 1 | ; | H2 | - | C16 |
| 716 | 718 | 1 | ; | H3 | - | C2 |
| 716 | 757 | 1 | ; | H3 | - | C16 |
| 717 | 723 | 1 | ; | O | - | C6 |
| 717 | 727 | 1 | ; | O | - | H5 |
| 717 | 728 | 1 | ; | O | - | N2 |
| 717 | 757 | 1 | ; | O | - | C16 |
| 718 | 724 | 1 | ; | C2 | - | H4 |
| 718 | 725 | 1 | ; | C2 | - | S |
| 718 | 726 | 1 | ; | C2 | - | N1 |
| 718 | 738 | 1 | ; | C2 | - | C10 |
| 718 | 740 | 1 | ; | C2 | - | H10 |
| 719 | 723 | 1 | ; | O1 | - | C6 |
| 719 | 727 | 1 | ; | O1 | - | H5 |
| 719 | 728 | 1 | ; | O1 | - | N2 |
| 720 | 721 | 1 | ; | C3 | - | C4 |
| 720 | 735 | 1 | ; | C3 | - | C9 |
| 720 | 739 | 1 | ; | C3 | - | O4 |
| 720 | 741 | 1 | ; | C3 | - | C11 |
| 721 | 724 | 1 | ; | C4 | - | H4 |
| 721 | 725 | 1 | ; | C4 | - | S |
| 721 | 733 | 1 | ; | C4 | - | H6 |
| 721 | 734 | 1 | ; | C4 | - | H7 |
| 721 | 736 | 1 | ; | C4 | - | H8 |
| 721 | 737 | 1 | ; | C4 | - | H9 |
| 722 | 723 | 1 | ; | C5 | - | C6 |
| 722 | 730 | 1 | ; | C5 | - | O2 |
| 722 | 731 | 1 | ; | C5 | - | O3 |
| 723 | 729 | 1 | ; | C6 | - | C7 |
| 723 | 736 | 1 | ; | C6 | - | H8 |
| 723 | 737 | 1 | ; | C6 | - | H9 |
| 723 | 738 | 1 | ; | C6 | - | C10 |
| 723 | 740 | 1 | ; | C6 | - | H10 |
| 724 | 727 | 1 | ; | H4 | - | H5 |
| 724 | 728 | 1 | ; | H4 | - | N2 |
| 724 | 735 | 1 | ; | H4 | - | C9 |
| 725 | 727 | 1 | ; | S | - | H5 |
| 725 | 728 | 1 | ; | S | - | N2 |
| 725 | 732 | 1 | ; | S | - | C8 |
| 726 | 727 | 1 | ; | N1 | - | H5 |
| 726 | 728 | 1 | ; | N1 | - | N2 |
| 726 | 730 | 1 | ; | N1 | - | O2 |
| 726 | 731 | 1 | ; | N1 | - | O3 |
| 726 | 732 | 1 | ; | N1 | - | C8 |
| 726 | 735 | 1 | ; | N1 | - | C9 |
| 727 | 738 | 1 | ; | H5 | - | C10 |
| 727 | 740 | 1 | ; | H5 | - | H10 |
| 728 | 742 | 1 | ; | N2 | - | N3 |
| 728 | 748 | 1 | ; | N2 | - | C13 |
| 729 | 732 | 1 | ; | C7 | - | C8 |
| 729 | 735 | 1 | ; | C7 | - | C9 |
| 732 | 736 | 1 | ; | C8 | - | H8 |
| 732 | 737 | 1 | ; | C8 | - | H9 |
| 733 | 735 | 1 | ; | H6 | - | C9 |
| 734 | 735 | 1 | ; | H7 | - | C9 |
| 738 | 743 | 1 | ; | C10 | - | O5 |
| 738 | 749 | 1 | ; | C10 | - | C14 |
| 738 | 753 | 1 | ; | C10 | - | N4 |
| 739 | 740 | 1 | ; | O4 | - | H10 |
| 739 | 742 | 1 | ; | O4 | - | N3 |
| 739 | 748 | 1 | ; | O4 | - | C13 |
| 740 | 741 | 1 | ; | H10 | - | C11 |
| 741 | 744 | 1 | ; | C11 | - | C12 |
| 741 | 750 | 1 | ; | C11 | - | H14 |
| 741 | 751 | 1 | ; | C11 | - | C15 |
| 741 | 752 | 1 | ; | C11 | - | S1 |
| 742 | 745 | 1 | ; | N3 | - | H11 |
| 742 | 746 | 1 | ; | N3 | - | H12 |
| 742 | 747 | 1 | ; | N3 | - | H13 |

| | | | | | | |
|---|---|---|---|---|---|---|
| 742 | 749 | 1 | ; | N3 | - | C14 |
| 742 | 753 | 1 | ; | N3 | - | N4 |
| 743 | 748 | 1 | ; | O5 | - | C13 |
| 748 | 754 | 1 | ; | C13 | - | N5 |
| 749 | 754 | 1 | ; | C14 | - | N5 |
| 750 | 751 | 1 | ; | H14 | - | C15 |
| 750 | 753 | 1 | ; | H14 | - | N4 |
| 752 | 755 | 1 | ; | S1 | - | H15 |
| 752 | 756 | 1 | ; | S1 | - | H16 |
| 753 | 755 | 1 | ; | N4 | - | H15 |
| 753 | 756 | 1 | ; | N4 | - | H16 |

[ angles ]

| ; ai | aj | ak | funct | theta | cth | | | | | | |
|---|---|---|---|---|---|---|---|---|---|---|---|
| 712 | 714 | 715 | 1 | 1.0956E+02 | 3.8828E+02 | ; | C | - | C1 | - | H2 |
| 712 | 714 | 716 | 1 | 1.0956E+02 | 3.8828E+02 | ; | C | - | C1 | - | H3 |
| 712 | 714 | 717 | 1 | 1.0797E+02 | 5.6902E+02 | ; | C | - | C1 | - | O |
| 712 | 757 | 758 | 1 | 1.2320E+02 | 5.6400E+02 | ; | C | - | C16 | - | O6 |
| 713 | 712 | 714 | 1 | 1.0956E+02 | 3.8828E+02 | ; | H1 | - | C | - | C1 |
| 713 | 712 | 757 | 1 | 1.0822E+02 | 3.9330E+02 | ; | H1 | - | C | - | C16 |
| 714 | 712 | 757 | 1 | 1.1104E+02 | 5.2969E+02 | ; | C1 | - | C | - | C16 |
| 714 | 717 | 718 | 1 | 1.1598E+02 | 5.2969E+02 | ; | C1 | - | O | - | C2 |
| 715 | 714 | 716 | 1 | 1.0846E+02 | 3.2803E+02 | ; | H2 | - | C1 | - | H3 |
| 715 | 714 | 717 | 1 | 1.0978E+02 | 4.2509E+02 | ; | H2 | - | C1 | - | O |
| 716 | 714 | 717 | 1 | 1.0978E+02 | 4.2509E+02 | ; | H3 | - | C1 | - | O |
| 717 | 718 | 719 | 1 | 1.2325E+02 | 6.3011E+02 | ; | O | - | C2 | - | O1 |
| 717 | 718 | 720 | 1 | 1.1072E+02 | 5.7656E+02 | ; | O | - | C2 | - | C3 |
| 718 | 720 | 723 | 1 | 1.1104E+02 | 5.2969E+02 | ; | C2 | - | C3 | - | C6 |
| 718 | 720 | 727 | 1 | 1.0822E+02 | 3.9330E+02 | ; | C2 | - | C3 | - | H5 |
| 718 | 720 | 728 | 1 | 1.0906E+02 | 5.6066E+02 | ; | C2 | - | C3 | - | N2 |
| 719 | 718 | 720 | 1 | 1.2320E+02 | 5.6400E+02 | ; | O1 | - | C2 | - | C3 |
| 720 | 723 | 724 | 1 | 1.1022E+02 | 3.8660E+02 | ; | C3 | - | C6 | - | H4 |
| 720 | 723 | 725 | 1 | 1.1027E+02 | 5.1296E+02 | ; | C3 | - | C6 | - | S |
| 720 | 723 | 726 | 1 | 1.0880E+02 | 5.5815E+02 | ; | C3 | - | C6 | - | N1 |
| 720 | 728 | 738 | 1 | 1.2069E+02 | 5.3053E+02 | ; | C3 | - | N2 | - | C10 |
| 720 | 728 | 740 | 1 | 1.1768E+02 | 3.8325E+02 | ; | C3 | - | N2 | - | H10 |
| 721 | 722 | 732 | 1 | 1.2326E+02 | 5.4727E+02 | ; | C4 | - | C5 | - | C8 |
| 721 | 722 | 735 | 1 | 1.1712E+02 | 5.3053E+02 | ; | C4 | - | C5 | - | C9 |
| 721 | 726 | 723 | 1 | 1.1867E+02 | 5.4810E+02 | ; | C4 | - | N1 | - | C6 |
| 721 | 729 | 730 | 1 | 1.2320E+02 | 5.7572E+02 | ; | C4 | - | C7 | - | O2 |
| 721 | 729 | 731 | 1 | 1.2320E+02 | 5.7572E+02 | ; | C4 | - | C7 | - | O3 |
| 722 | 721 | 726 | 1 | 1.1893E+02 | 5.7990E+02 | ; | C5 | - | C4 | - | N1 |

| | | | | | | | | | | | |
|---|---|---|---|---|---|---|---|---|---|---|---|
| 722 | 721 | 729 | 1 | 1.2098E+02 | 5.2802E+02 | ; | C5 | - | C4 | - | C7 |
| 722 | 732 | 733 | 1 | 1.2045E+02 | 4.1422E+02 | ; | C5 | - | C8 | - | H6 |
| 722 | 732 | 734 | 1 | 1.2045E+02 | 4.1422E+02 | ; | C5 | - | C8 | - | H7 |
| 722 | 735 | 725 | 1 | 1.1072E+02 | 5.1380E+02 | ; | C5 | - | C9 | - | S |
| 722 | 735 | 736 | 1 | 1.0954E+02 | 3.9330E+02 | ; | C5 | - | C9 | - | H8 |
| 722 | 735 | 737 | 1 | 1.0954E+02 | 3.9330E+02 | ; | C5 | - | C9 | - | H9 |
| 723 | 720 | 727 | 1 | 1.0956E+02 | 3.8828E+02 | ; | C6 | - | C3 | - | H5 |
| 723 | 720 | 728 | 1 | 1.1161E+02 | 5.5145E+02 | ; | C6 | - | C3 | - | N2 |
| 723 | 725 | 735 | 1 | 9.9240E+01 | 5.0375E+02 | ; | C6 | - | S | - | C9 |
| 724 | 723 | 725 | 1 | 1.0833E+02 | 3.5229E+02 | ; | H4 | - | C6 | - | S |
| 724 | 723 | 726 | 1 | 1.1020E+02 | 4.1338E+02 | ; | H4 | - | C6 | - | N1 |
| 725 | 723 | 726 | 1 | 1.0939E+02 | 5.3388E+02 | ; | S | - | C6 | - | N1 |
| 725 | 735 | 736 | 1 | 1.0876E+02 | 3.5229E+02 | ; | S | - | C9 | - | H8 |
| 725 | 735 | 737 | 1 | 1.0876E+02 | 3.5229E+02 | ; | S | - | C9 | - | H9 |
| 726 | 721 | 729 | 1 | 1.1441E+02 | 5.8492E+02 | ; | N1 | - | C4 | - | C7 |
| 727 | 720 | 728 | 1 | 1.0888E+02 | 4.1673E+02 | ; | H5 | - | C3 | - | N2 |
| 728 | 738 | 739 | 1 | 1.2305E+02 | 6.2091E+02 | ; | N2 | - | C10 | - | O4 |
| 728 | 738 | 741 | 1 | 1.1522E+02 | 5.6819E+02 | ; | N2 | - | C10 | - | C11 |
| 730 | 729 | 731 | 1 | 1.3025E+02 | 6.5187E+02 | ; | O2 | - | C7 | - | O3 |
| 732 | 722 | 735 | 1 | 1.2253E+02 | 5.3555E+02 | ; | C8 | - | C5 | - | C9 |
| 733 | 732 | 734 | 1 | 1.1690E+02 | 3.1882E+02 | ; | H6 | - | C8 | - | H7 |
| 736 | 735 | 737 | 1 | 1.0846E+02 | 3.2803E+02 | ; | H8 | - | C9 | - | H9 |
| 738 | 728 | 740 | 1 | 1.1755E+02 | 4.0417E+02 | ; | C10 | - | N2 | - | H10 |
| 738 | 741 | 742 | 1 | 1.1441E+02 | 5.8492E+02 | ; | C10 | - | C11 | - | N3 |
| 738 | 741 | 748 | 1 | 1.1782E+02 | 5.3555E+02 | ; | C10 | - | C11 | - | C13 |
| 739 | 738 | 741 | 1 | 1.2320E+02 | 5.7572E+02 | ; | O4 | - | C10 | - | C11 |
| 741 | 742 | 743 | 1 | 1.1279E+02 | 5.9580E+02 | ; | C11 | - | N3 | - | O5 |
| 741 | 748 | 749 | 1 | 1.2805E+02 | 5.3304E+02 | ; | C11 | - | C13 | - | C14 |
| 741 | 748 | 753 | 1 | 1.2110E+02 | 5.6233E+02 | ; | C11 | - | C13 | - | N4 |
| 742 | 741 | 748 | 1 | 1.2096E+02 | 5.7572E+02 | ; | N3 | - | C11 | - | C13 |
| 742 | 743 | 744 | 1 | 1.0923E+02 | 5.5145E+02 | ; | N3 | - | O5 | - | C12 |
| 743 | 744 | 745 | 1 | 1.0978E+02 | 4.2509E+02 | ; | O5 | - | C12 | - | H11 |
| 743 | 744 | 746 | 1 | 1.0978E+02 | 4.2509E+02 | ; | O5 | - | C12 | - | H12 |
| 743 | 744 | 747 | 1 | 1.0978E+02 | 4.2509E+02 | ; | O5 | - | C12 | - | H13 |
| 745 | 744 | 746 | 1 | 1.0846E+02 | 3.2803E+02 | ; | H11 | - | C12 | - | H12 |

| | | | | | | | | | | |
|---|---|---|---|---|---|---|---|---|---|---|
| 745 | 744 | 747 | 1 | 1.0846E+02 | 3.2803E+02 | ; | H11 | - | C12 | - |
| | H13 | | | | | | | | | |
| 746 | 744 | 747 | 1 | 1.0846E+02 | 3.2803E+02 | ; | H12 | - | C12 | - |
| | H13 | | | | | | | | | |
| 748 | 749 | 750 | 1 | 1.2848E+02 | 3.9581E+02 | ; | C13 | - | C14 | - |
| | H14 | | | | | | | | | |
| 748 | 749 | 752 | 1 | 1.1155E+02 | 5.4225E+02 | ; | C13 | - | C14 | - |
| | S1 | | | | | | | | | |
| 748 | 753 | 751 | 1 | 1.0549E+02 | 6.0082E+02 | ; | C13 | - | N4 | - |
| | C15 | | | | | | | | | |
| 749 | 748 | 753 | 1 | 1.1165E+02 | 6.0417E+02 | ; | C14 | - | C13 | - |
| | N4 | | | | | | | | | |
| 749 | 752 | 751 | 1 | 9.0240E+01 | 5.5396E+02 | ; | C14 | - | S1 | - |
| | C15 | | | | | | | | | |
| 750 | 749 | 752 | 1 | 1.1997E+02 | 3.5229E+02 | ; | H14 | - | C14 | - |
| | S1 | | | | | | | | | |
| 751 | 754 | 755 | 1 | 1.1563E+02 | 4.0920E+02 | ; | C15 | - | N5 | - |
| | H15 | | | | | | | | | |
| 751 | 754 | 756 | 1 | 1.1563E+02 | 4.0920E+02 | ; | C15 | - | N5 | - |
| | H16 | | | | | | | | | |
| 752 | 751 | 753 | 1 | 1.1451E+02 | 5.5229E+02 | ; | S1 | - | C15 | - |
| | N4 | | | | | | | | | |
| 752 | 751 | 754 | 1 | 1.2181E+02 | 5.3220E+02 | ; | S1 | - | C15 | - |
| | N5 | | | | | | | | | |
| 753 | 751 | 754 | 1 | 1.2065E+02 | 6.0584E+02 | ; | N4 | - | C15 | - |
| | N5 | | | | | | | | | |
| 755 | 754 | 756 | 1 | 1.1512E+02 | 3.3556E+02 | ; | H15 | - | N5 | - |
| | H16 | | | | | | | | | |

[ dihedrals ] ; propers
; treated as RBs in GROMACS to use combine multiple AMBER torsions per quartet

| ; i | j | k | l | func | C0 | C1 | C2 | C3 | C4 | C5 | |
|---|---|---|---|---|---|---|---|---|---|---|---|
| 710 | 712 | 714 | 717 | 3 | 6.5084E-01 | 1.95253 | 0 | -2.60338 | 0 | 0 | ; |
| | N- | C- | C1- | O | | | | | | | |
| 711 | 710 | 712 | 713 | 3 | 0.0000E+00 | 0 | 0 | 0 | 0 | 0 | ; |
| | H- | N- | C- | H1 | | | | | | | |
| 711 | 710 | 712 | 714 | 3 | 0.0000E+00 | 0 | 0 | 0 | 0 | 0 | ; |
| | H- | N- | C- | C1 | | | | | | | |
| 712 | 714 | 717 | 718 | 3 | 4.9497E+00 | 8.15462 | 0 | -6.40989 | 0 | 0 | ; |
| | C- | C1- | O- | C2 | | | | | | | |
| 713 | 712 | 714 | 715 | 3 | 6.5084E-01 | 1.95253 | 0 | -2.60338 | 0 | 0 | ; |
| | H1- | C- | C1- | H2 | | | | | | | |
| 713 | 712 | 714 | 716 | 3 | 6.5084E-01 | 1.95253 | 0 | -2.60338 | 0 | 0 | ; |
| | H1- | C- | C1- | H3 | | | | | | | |
| 713 | 712 | 714 | 717 | 3 | 1.0460E+00 | -1.046 | 0 | 0 | 0 | 0 | ; |
| | H1- | C- | C1- | O | | | | | | | |
| 714 | 717 | 718 | 719 | 3 | 2.8451E+01 | 5.8576 | -22.5936 | 0 | 0 | 0 | ; |
| | C1- | O- | C2- | O1 | | | | | | | |
| 714 | 717 | 718 | 720 | 3 | 2.7405E+01 | 14.4348 | -22.5936 | -19.2464 | 0 | 0 | ; |
| | C1- | O- | C2- | C3 | | | | | | | |
| 715 | 714 | 717 | 718 | 3 | 1.6039E+00 | 4.8116 | 0 | -6.41547 | 0 | 0 | ; |
| | H2- | C1- | O- | C2 | | | | | | | |
| 716 | 714 | 717 | 718 | 3 | 1.6039E+00 | 4.8116 | 0 | -6.41547 | 0 | 0 | ; |
| | H3- | C1- | O- | C2 | | | | | | | |
| 717 | 718 | 720 | 723 | 3 | 0.0000E+00 | 0 | 0 | 0 | 0 | 0 | ; |
| | O- | C2- | C3- | C6 | | | | | | | |
| 717 | 718 | 720 | 727 | 3 | 0.0000E+00 | 0 | 0 | 0 | 0 | 0 | ; |
| | O- | C2- | C3- | H5 | | | | | | | |
| 717 | 718 | 720 | 728 | 3 | 0.0000E+00 | 0 | 0 | 0 | 0 | 0 | ; |
| | O- | C2- | C3- | N2 | | | | | | | |
| 718 | 720 | 723 | 724 | 3 | 6.5084E-01 | 1.95253 | 0 | -2.60338 | 0 | 0 | ; |
| | C2- | C3- | C6- | H4 | | | | | | | |
| 718 | 720 | 723 | 725 | 3 | 6.5084E-01 | 1.95253 | 0 | -2.60338 | 0 | 0 | ; |
| | C2- | C3- | C6- | S | | | | | | | |
| 718 | 720 | 723 | 726 | 3 | 6.5084E-01 | 1.95253 | 0 | -2.60338 | 0 | 0 | ; |
| | C2- | C3- | C6- | N1 | | | | | | | |
| 718 | 720 | 728 | 738 | 3 | 1.0460E+01 | -3.3472 | -7.1128 | 0 | 0 | 0 | ; |
| | C2- | C3- | N2- | C10 | | | | | | | |
| 718 | 720 | 728 | 740 | 3 | 0.0000E+00 | 0 | 0 | 0 | 0 | 0 | ; |
| | C2- | C3- | N2- | H10 | | | | | | | |

| | | | | | | | | | | | |
|---|---|---|---|---|---|---|---|---|---|---|---|
| 719 | 718 | 720 | 723 | 3 | 0.0000E+00 | 0 | 0 | 0 | 0 | 0 | ; |
| | O1- | C2- | C3- | C6 | | | | | | | |
| 719 | 718 | 720 | 727 | 3 | 3.6819E+00 | -4.35136 | 0 | 1.33888 | 0 | 0 | ; |
| | O1- | C2- | C3- | H5 | | | | | | | |
| 719 | 718 | 720 | 728 | 3 | 0.0000E+00 | 0 | 0 | 0 | 0 | 0 | ; |
| | O1- | C2- | C3- | N2 | | | | | | | |
| 720 | 723 | 725 | 735 | 3 | 1.3947E+00 | 4.184 | 0 | -5.57867 | 0 | 0 | ; |
| | C3- | C6- | S- | C9 | | | | | | | |
| 720 | 723 | 726 | 721 | 3 | 0.0000E+00 | 0 | 0 | 0 | 0 | 0 | ; |
| | C3- | C6- | N1- | C4 | | | | | | | |
| 720 | 728 | 738 | 739 | 3 | 2.0920E+01 | 0 | -20.92 | 0 | 0 | 0 | ; |
| | C3- | N2- | C10- | O4 | | | | | | | |
| 720 | 728 | 738 | 741 | 3 | 2.0920E+01 | 0 | -20.92 | 0 | 0 | 0 | ; |
| | C3- | N2- | C10- | C11 | | | | | | | |
| 721 | 722 | 732 | 733 | 3 | 5.5647E+01 | 0 | -55.6472 | 0 | 0 | 0 | ; |
| | C4- | C5- | C8- | H6 | | | | | | | |
| 721 | 722 | 732 | 734 | 3 | 5.5647E+01 | 0 | -55.6472 | 0 | 0 | 0 | ; |
| | C4- | C5- | C8- | H7 | | | | | | | |
| 721 | 722 | 735 | 725 | 3 | 0.0000E+00 | 0 | 0 | 0 | 0 | 0 | ; |
| | C4- | C5- | C9- | S | | | | | | | |
| 721 | 722 | 735 | 736 | 3 | 6.4015E+00 | -9.58136 | 0 | 6.35968 | 0 | 0 | ; |
| | C4- | C5- | C9- | H8 | | | | | | | |
| 721 | 722 | 735 | 737 | 3 | 6.4015E+00 | -9.58136 | 0 | 6.35968 | 0 | 0 | ; |
| | C4- | C5- | C9- | H9 | | | | | | | |
| 721 | 726 | 723 | 724 | 3 | 0.0000E+00 | 0 | 0 | 0 | 0 | 0 | ; |
| | C4- | N1- | C6- | H4 | | | | | | | |
| 721 | 726 | 723 | 725 | 3 | 0.0000E+00 | 0 | 0 | 0 | 0 | 0 | ; |
| | C4- | N1- | C6- | S | | | | | | | |
| 722 | 721 | 726 | 723 | 3 | 6.6944E+00 | 0 | -6.6944 | 0 | 0 | 0 | ; |
| | C5- | C4- | N1- | C6 | | | | | | | |
| 722 | 721 | 729 | 730 | 3 | 1.8200E+01 | 0 | -18.2004 | 0 | 0 | 0 | ; |
| | C5- | C4- | C7- | O2 | | | | | | | |
| 722 | 721 | 729 | 731 | 3 | 1.8200E+01 | 0 | -18.2004 | 0 | 0 | 0 | ; |
| | C5- | C4- | C7- | O3 | | | | | | | |
| 722 | 735 | 725 | 723 | 3 | 1.3947E+00 | 4.184 | 0 | -5.57867 | 0 | 0 | ; |
| | C5- | C9- | S- | C6 | | | | | | | |
| 723 | 720 | 728 | 738 | 3 | 2.8451E+00 | -4.10032 | 16.736 | 2.5104 | -16.736 | 0 | ; |
| | C6- | C3- | N2- | C10 | | | | | | | |
| 723 | 720 | 728 | 740 | 3 | 0.0000E+00 | 0 | 0 | 0 | 0 | 0 | ; |
| | C6- | C3- | N2- | H10 | | | | | | | |
| 723 | 725 | 735 | 736 | 3 | 1.3947E+00 | 4.184 | 0 | -5.57867 | 0 | 0 | ; |
| | C6- | S- | C9- | H8 | | | | | | | |
| 723 | 725 | 735 | 737 | 3 | 1.3947E+00 | 4.184 | 0 | -5.57867 | 0 | 0 | ; |
| | C6- | S- | C9- | H9 | | | | | | | |
| 723 | 726 | 721 | 729 | 3 | 6.6944E+00 | 0 | -6.6944 | 0 | 0 | 0 | ; |
| | C6- | N1- | C4- | C7 | | | | | | | |
| 724 | 723 | 720 | 727 | 3 | 6.5084E-01 | 1.95253 | 0 | -2.60338 | 0 | 0 | ; |
| | H4- | C6- | C3- | H5 | | | | | | | |
| 724 | 723 | 720 | 728 | 3 | 6.5084E-01 | 1.95253 | 0 | -2.60338 | 0 | 0 | ; |
| | H4- | C6- | C3- | N2 | | | | | | | |
| 724 | 723 | 725 | 735 | 3 | 1.3947E+00 | 4.184 | 0 | -5.57867 | 0 | 0 | ; |
| | H4- | C6- | S- | C9 | | | | | | | |
| 725 | 723 | 720 | 727 | 3 | 6.5084E-01 | 1.95253 | 0 | -2.60338 | 0 | 0 | ; |
| | S- | C6- | C3- | H5 | | | | | | | |
| 725 | 723 | 720 | 728 | 3 | 6.5084E-01 | 1.95253 | 0 | -2.60338 | 0 | 0 | ; |
| | S- | C6- | C3- | N2 | | | | | | | |
| 725 | 735 | 722 | 732 | 3 | 0.0000E+00 | 0 | 0 | 0 | 0 | 0 | ; |
| | S- | C9- | C5- | C8 | | | | | | | |
| 726 | 721 | 722 | 732 | 3 | 8.3680E+00 | 0 | -8.368 | 0 | 0 | 0 | ; |
| | N1- | C4- | C5- | C8 | | | | | | | |
| 726 | 721 | 722 | 735 | 3 | 8.3680E+00 | 0 | -8.368 | 0 | 0 | 0 | ; |
| | N1- | C4- | C5- | C9 | | | | | | | |
| 726 | 721 | 729 | 730 | 3 | 1.8200E+01 | 0 | -18.2004 | 0 | 0 | 0 | ; |
| | N1- | C4- | C7- | O2 | | | | | | | |
| 726 | 721 | 729 | 731 | 3 | 1.8200E+01 | 0 | -18.2004 | 0 | 0 | 0 | ; |
| | N1- | C4- | C7- | O3 | | | | | | | |
| 726 | 723 | 720 | 727 | 3 | 6.5084E-01 | 1.95253 | 0 | -2.60338 | 0 | 0 | ; |
| | N1- | C6- | C3- | H5 | | | | | | | |
| 726 | 723 | 720 | 728 | 3 | 6.5084E-01 | 1.95253 | 0 | -2.60338 | 0 | 0 | ; |
| | N1- | C6- | C3- | N2 | | | | | | | |

| | | | | | | | | | | | |
|---|---|---|---|---|---|---|---|---|---|---|---|
| 726 | 723 | 725 | 735 | 3 | 1.3947E+00 | 4.184 | 0 | -5.57867 | 0 | 0 | ; |
| | N1- | C6- | S- | C9 | | | | | | | |
| 727 | 720 | 728 | 738 | 3 | 0.0000E+00 | 0 | 0 | 0 | 0 | 0 | ; |
| | H5- | C3- | N2- | C10 | | | | | | | |
| 727 | 720 | 728 | 740 | 3 | 0.0000E+00 | 0 | 0 | 0 | 0 | 0 | ; |
| | H5- | C3- | N2- | H10 | | | | | | | |
| 728 | 738 | 741 | 742 | 3 | 1.8200E+01 | 0 | -18.2004 | 0 | 0 | 0 | ; |
| | N2- | C10- | C11- | N3 | | | | | | | |
| 728 | 738 | 741 | 748 | 3 | 1.8200E+01 | 0 | -18.2004 | 0 | 0 | 0 | ; |
| | N2- | C10- | C11- | C13 | | | | | | | |
| 729 | 721 | 722 | 732 | 3 | 8.3680E+00 | 0 | -8.368 | 0 | 0 | 0 | ; |
| | C7- | C4- | C5- | C8 | | | | | | | |
| 729 | 721 | 722 | 735 | 3 | 8.3680E+00 | 0 | -8.368 | 0 | 0 | 0 | ; |
| | C7- | C4- | C5- | C9 | | | | | | | |
| 732 | 722 | 735 | 736 | 3 | 6.4015E+00 | -9.58136 | 0 | 6.35968 | 0 | 0 | ; |
| | C8- | C5- | C9- | H8 | | | | | | | |
| 732 | 722 | 735 | 737 | 3 | 6.4015E+00 | -9.58136 | 0 | 6.35968 | 0 | 0 | ; |
| | C8- | C5- | C9- | H9 | | | | | | | |
| 733 | 732 | 722 | 735 | 3 | 5.5647E+01 | 0 | -55.6472 | 0 | 0 | 0 | ; |
| | H6- | C8- | C5- | C9 | | | | | | | |
| 734 | 732 | 722 | 735 | 3 | 5.5647E+01 | 0 | -55.6472 | 0 | 0 | 0 | ; |
| | H7- | C8- | C5- | C9 | | | | | | | |
| 738 | 741 | 742 | 743 | 3 | 6.6944E+00 | 0 | -6.6944 | 0 | 0 | 0 | ; |
| | C10- | C11- | N3- | O5 | | | | | | | |
| 738 | 741 | 748 | 749 | 3 | 8.3680E+00 | 0 | -8.368 | 0 | 0 | 0 | ; |
| | C10- | C11- | C13- | C14 | | | | | | | |
| 738 | 741 | 748 | 753 | 3 | 8.3680E+00 | 0 | -8.368 | 0 | 0 | 0 | ; |
| | C10- | C11- | C13- | N4 | | | | | | | |
| 739 | 738 | 728 | 740 | 3 | 2.9288E+01 | -8.368 | -20.92 | 0 | 0 | 0 | ; |
| | O4- | C10- | N2- | H10 | | | | | | | |
| 739 | 738 | 741 | 742 | 3 | 1.8200E+01 | 0 | -18.2004 | 0 | 0 | 0 | ; |
| | O4- | C10- | C11- | N3 | | | | | | | |
| 739 | 738 | 741 | 748 | 3 | 1.8200E+01 | 0 | -18.2004 | 0 | 0 | 0 | ; |
| | O4- | C10- | C11- | C13 | | | | | | | |
| 740 | 728 | 738 | 741 | 3 | 2.0920E+01 | 0 | -20.92 | 0 | 0 | 0 | ; |
| | H10- | N2- | C10- | C11 | | | | | | | |
| 741 | 742 | 743 | 744 | 3 | 2.5104E+01 | 0 | -25.104 | 0 | 0 | 0 | ; |
| | C11- | N3- | O5- | C12 | | | | | | | |
| 741 | 748 | 749 | 750 | 3 | 3.3472E+01 | 0 | -33.472 | 0 | 0 | 0 | ; |
| | C11- | C13- | C14- | H14 | | | | | | | |
| 741 | 748 | 749 | 752 | 3 | 3.3472E+01 | 0 | -33.472 | 0 | 0 | 0 | ; |
| | C11- | C13- | C14- | S1 | | | | | | | |
| 741 | 748 | 753 | 751 | 3 | 3.9748E+01 | 0 | -39.748 | 0 | 0 | 0 | ; |
| | C11- | C13- | N4- | C15 | | | | | | | |
| 742 | 741 | 748 | 749 | 3 | 8.3680E+00 | 0 | -8.368 | 0 | 0 | 0 | ; |
| | N3- | C11- | C13- | C14 | | | | | | | |
| 742 | 741 | 748 | 753 | 3 | 8.3680E+00 | 0 | -8.368 | 0 | 0 | 0 | ; |
| | N3- | C11- | C13- | N4 | | | | | | | |
| 742 | 743 | 744 | 745 | 3 | 1.6039E+00 | 4.8116 | 0 | -6.41547 | 0 | 0 | ; |
| | N3- | O5- | C12- | H11 | | | | | | | |
| 742 | 743 | 744 | 746 | 3 | 1.6039E+00 | 4.8116 | 0 | -6.41547 | 0 | 0 | ; |
| | N3- | O5- | C12- | H12 | | | | | | | |
| 742 | 743 | 744 | 747 | 3 | 1.6039E+00 | 4.8116 | 0 | -6.41547 | 0 | 0 | ; |
| | N3- | O5- | C12- | H13 | | | | | | | |
| 743 | 742 | 741 | 748 | 3 | 6.6944E+00 | 0 | -6.6944 | 0 | 0 | 0 | ; |
| | O5- | N3- | C11- | C13 | | | | | | | |
| 748 | 749 | 752 | 751 | 3 | 9.2048E+00 | 0 | -9.2048 | 0 | 0 | 0 | ; |
| | C13- | C14- | S1- | C15 | | | | | | | |
| 748 | 753 | 751 | 752 | 3 | 3.9748E+01 | 0 | -39.748 | 0 | 0 | 0 | ; |
| | C13- | N4- | C15- | S1 | | | | | | | |
| 748 | 753 | 751 | 754 | 3 | 3.9748E+01 | 0 | -39.748 | 0 | 0 | 0 | ; |
| | C13- | N4- | C15- | N5 | | | | | | | |
| 749 | 748 | 753 | 751 | 3 | 3.9748E+01 | 0 | -39.748 | 0 | 0 | 0 | ; |
| | C14- | C13- | N4- | C15 | | | | | | | |
| 749 | 752 | 751 | 753 | 3 | 9.2048E+00 | 0 | -9.2048 | 0 | 0 | 0 | ; |
| | C14- | S1- | C15- | N4 | | | | | | | |
| 749 | 752 | 751 | 754 | 3 | 9.2048E+00 | 0 | -9.2048 | 0 | 0 | 0 | ; |
| | C14- | S1- | C15- | N5 | | | | | | | |
| 750 | 749 | 748 | 753 | 3 | 3.3472E+01 | 0 | -33.472 | 0 | 0 | 0 | ; |
| | H14- | C14- | C13- | N4 | | | | | | | |

| | | | | | | | | | | | |
|---|---|---|---|---|---|---|---|---|---|---|---|
| 750 | 749 | 752 | 751 | 3 | 9.2048E+00 | 0 | -9.2048 | 0 | 0 | 0 | ; |
| | H14- | C14- | S1- | C15 | | | | | | | |
| 752 | 749 | 748 | 753 | 3 | 3.3472E+01 | 0 | -33.472 | 0 | 0 | 0 | ; |
| | S1- | C14- | C13- | N4 | | | | | | | |
| 752 | 751 | 754 | 755 | 3 | 8.7864E+00 | 0 | -8.7864 | 0 | 0 | 0 | ; |
| | S1- | C15- | N5- | H15 | | | | | | | |
| 752 | 751 | 754 | 756 | 3 | 8.7864E+00 | 0 | -8.7864 | 0 | 0 | 0 | ; |
| | S1- | C15- | N5- | H16 | | | | | | | |
| 753 | 751 | 754 | 755 | 3 | 8.7864E+00 | 0 | -8.7864 | 0 | 0 | 0 | ; |
| | N4- | C15- | N5- | H15 | | | | | | | |
| 753 | 751 | 754 | 756 | 3 | 8.7864E+00 | 0 | -8.7864 | 0 | 0 | 0 | ; |
| | N4- | C15- | N5- | H16 | | | | | | | |

[ dihedrals ] ; impropers
; treated as propers in GROMACS to use correct AMBER analytical function
; i j k l func phase kd pn

| | | | | | | | | | | | | |
|---|---|---|---|---|---|---|---|---|---|---|---|---|
| 720 | 719 | 718 | 717 | 1 | 180 | 4.6024 | 2 | ; | C3- | O1- | C2- | O |
| 721 | 730 | 729 | 731 | 1 | 180 | 4.6024 | 2 | ; | C4- | O2- | C7- | O3 |
| 722 | 733 | 732 | 734 | 1 | 180 | 4.6024 | 2 | ; | C5- | H6- | C8- | H7 |
| 729 | 722 | 721 | 726 | 1 | 180 | 4.6024 | 2 | ; | C7- | C5- | C4- | N1 |
| 732 | 735 | 722 | 721 | 1 | 180 | 4.6024 | 2 | ; | C8- | C9- | C5- | C4 |
| 738 | 720 | 728 | 740 | 1 | 180 | 4.6024 | 2 | ; | C10- | C3- | N2- | H10 |
| 738 | 748 | 741 | 742 | 1 | 180 | 4.6024 | 2 | ; | C10- | C13- | C11- | N3 |
| 741 | 728 | 738 | 739 | 1 | 180 | 43.932 | 2 | ; | C11- | N2- | C10- | O4 |
| 748 | 750 | 749 | 752 | 1 | 180 | 4.6024 | 2 | ; | C13- | H14- | C14- | S1 |
| 749 | 741 | 748 | 753 | 1 | 180 | 4.6024 | 2 | ; | C14- | C11- | C13- | N4 |
| 751 | 755 | 754 | 756 | 1 | 180 | 4.6024 | 2 | ; | C15- | H15- | N5- | H16 |
| 753 | 754 | 751 | 752 | 1 | 180 | 4.6024 | 2 | ; | N4- | N5- | C15- | S1 |

### Matlab Scripts –

---

*"MMLam.m"*

---

```
function [Km,kcat,y,residual,StErr]=MMLam(Data);
%"Data" is a 2-column matrix of x-values (1st column; sec), and y-values (2nd
%column; uM)for analytical fitting of steady-state time traces.
    t=Data(:,1); %Time (sec), t
    St=Data(:,2); %Substrate concentration (uM) at each time, t
    function W=Lambert(beta,t,St)
    S0=100.00; %Starting substrate concentration (uM)
    E0=1.0; %Enzyme concentration (uM)
    Km=beta(1);
    kcat=beta(2);

    W=Km*lambertw((exp((S0-kcat*E0*t)./Km)).*(S0/Km)); %General form for MM kinetics
    end
fun=@Lambert;
beta0=[50, 50]; %initial guess for KM (uM) and kcat (sec-1)
options=statset('FunValCheck','off');
[beta,R,J,CovB]=nlinfit(t,St,fun,beta0,options);

y=beta(1)*lambertw((exp((S0-beta(2)*E0*t)./beta(1))).*(S0/beta(1)));
format long
Km=beta(1);
kcat=beta(2);
residual=R;
plot(t,St,t,y,t,St-y)
StErr=(sqrt(diag(CovB)));
end
```

---

---

*"KineticEstimator.m"*

---

```
function [t,U]=KineticEstimator(KM,kcat,S0,E0);
%Analytic simulation of a progress curve for steady-state enzymatic catalysis given a
set of kinetic parameters. Useful for predicting the duration of various kinetic
regimes. Input parameters as follows: KM=Michaelis Constant (uM), kcat (sec-1),
%S0=time-zero substrate concentration (uM), E0=time-zero enzyme
%concentration (uM)
t=[0:1:600]; %time domain to simulate (sec)
F=(S0/KM)*(exp((S0-(E0*kcat*t))/KM));
U=KM*lambertw(F);
plot(t,U)
xlabel('Time (sec)')
```

```
ylabel('[Substrate] (uM)')
xlim([min(t), max(t)])
ylim([-50 max(U)])
```

---

*"MassAnalysis_Gaussian.m"*

---

```
%% MassAnalysis_Gaussian
% Script for analyzing steady-state MS data of protein charge envelope.
% Input intact mass of free and covalently-bound enzyme, in addition to x (Da), y
% (counts) data for analysis. Input is a .txt file with two column format,
% x-data is in the first column, y-data in the second. This version implements
% a Gaussian baselining protocol to be implemented in the fitting, see
% lines for details. Written by Sam Schneider (Revised Jan 2020) for
% evaluation of acyl-enzyme/Michaelis complex ratios in TEM B-lactamases.
%% Load Data and Input Nominal Masses and Desired Baseline
% load MS data (x,y) from .txt file
[fname,pname]=uigetfile('*.*','Load MS Spectrum','MultiSelect','on');
cd(pname);
Data=readtable(fname,'Delimiter',' ','Format','%f %f','ReadVariableNames',false);
Da=Data; %x-data
Counts=Data; %y-data

%input mass of free (E) and bound enzyme (EA)
dlgtitle='Input Information';
numLines=1;
prompt1={'E mass (Da)','EA mass (Da)'};
def1={'31500','31914'};
answer1=inputdlg(prompt1,dlgtitle,numLines,def1);
drawnow

%Input the approximate masses of E and EA complex for guiding the fits. If
%too far off (+-30Da) may yield errors.
E_mass=str2num(char(answer1(1)));
EA_mass=str2num(char(answer1(2)));

%% Initial Fitting Parameter Initialization
Intensities=zeros(20,8); %Initialize matrix containing the n, E/n, area_1, area_2,
EA/n, area_1, area_2 values
E=zeros(20,1); % starting conditions for mass_fracs for enzyme
EA=zeros(20,1); %starting conditions for mass_fracs for EA complex

%Determine the starting guesses/bounds for the peak position searches to
%bound the fits
for n=1:20;
    E(n,1)=E_mass/(40-n);
    EA(n,1)=EA_mass/(40-n);
    Intensities(n,1)=(40-n);
end

offset=1.5;
amplb=0.9*min(Counts);
```

```
ampub=1.1*max(Counts);
ampstart=max(Counts);

%% Gaussian Baselining
% Fit 1 - Fit Main Peak (bounds of shoulder set to 0)
fun=@(a,x)a(2)*exp(-((x-a(3)).^2)/(2*a(1)^2))+a(4)*exp(-((x-
a(5)).^2)/(2*a(1)^2))+a(6)*exp(-((x-a(7)).^2)/(2*a(1)^2))+a(8)*exp(-((x-
a(9)).^2)/(2*a(1)^2))+a(10)*exp(-((x-a(11)).^2)/(2*a(1)^2))+a(12)*exp(-((x-
a(13)).^2)/(2*a(1)^2))+a(14)*exp(-((x-a(15)).^2)/(2*a(1)^2))+a(16)*exp(-((x-
a(17)).^2)/(2*a(1)^2))+a(18)*exp(-((x-a(19)).^2)/(2*a(1)^2))+a(20)*exp(-((x-
a(21)).^2)/(2*a(1)^2))+a(22)*exp(-((x-a(23)).^2)/(2*a(1)^2))+a(24)*exp(-((x-
a(25)).^2)/(2*a(1)^2))+a(26)*exp(-((x-a(27)).^2)/(2*a(1)^2))+a(28)*exp(-((x-
a(29)).^2)/(2*a(1)^2))+a(30)*exp(-((x-a(31)).^2)/(2*a(1)^2))+a(32)*exp(-((x-
a(33)).^2)/(2*a(1)^2))+a(34)*exp(-((x-a(35)).^2)/(2*a(1)^2))+a(36)*exp(-((x-
a(37)).^2)/(2*a(1)^2))+a(38)*exp(-((x-a(39)).^2)/(2*a(1)^2))+a(40)*exp(-((x-
a(41)).^2)/(2*a(1)^2))+a(42)*exp(-((x-a(43)).^2)/(2*a(1)^2))+a(44)*exp(-((x-
a(45)).^2)/(2*a(1)^2))+a(46)*exp(-((x-a(47)).^2)/(2*a(1)^2))+a(48)*exp(-((x-
a(49)).^2)/(2*a(1)^2))+a(50)*exp(-((x-a(51)).^2)/(2*a(1)^2))+a(52)*exp(-((x-
a(53)).^2)/(2*a(1)^2))+a(54)*exp(-((x-a(55)).^2)/(2*a(1)^2))+a(56)*exp(-((x-
a(57)).^2)/(2*a(1)^2))+a(58)*exp(-((x-a(59)).^2)/(2*a(1)^2))+a(60)*exp(-((x-
a(61)).^2)/(2*a(1)^2))+a(62)*exp(-((x-a(63)).^2)/(2*a(1)^2))+a(64)*exp(-((x-
a(65)).^2)/(2*a(1)^2))+a(66)*exp(-((x-a(67)).^2)/(2*a(1)^2))+a(68)*exp(-((x-
a(69)).^2)/(2*a(1)^2))+a(70)*exp(-((x-a(71)).^2)/(2*a(1)^2))+a(72)*exp(-((x-
a(73)).^2)/(2*a(1)^2))+a(74)*exp(-((x-a(75)).^2)/(2*a(1)^2))+a(76)*exp(-((x-
a(77)).^2)/(2*a(1)^2))+a(78)*exp(-((x-a(79)).^2)/(2*a(1)^2))+a(80)*exp(-((x-
a(81)).^2)/(2*a(1)^2))+a(82)*exp(-((x-a(3)+a(122)).^2)/(2*a(1)^2))+a(83)*exp(-((x-
a(5)+a(122)).^2)/(2*a(1)^2))+a(84)*exp(-((x-a(7)+a(122)).^2)/(2*a(1)^2))+a(85)*exp(-
((x-a(9)+a(122)).^2)/(2*a(1)^2))+a(86)*exp(-((x-
a(11)+a(122)).^2)/(2*a(1)^2))+a(87)*exp(-((x-a(13)+a(122)).^2)/(2*a(1)^2))+a(88)*exp(-
((x-a(15)+a(122)).^2)/(2*a(1)^2))+a(89)*exp(-((x-
a(17)+a(122)).^2)/(2*a(1)^2))+a(90)*exp(-((x-a(19)+a(122)).^2)/(2*a(1)^2))+a(91)*exp(-
((x-a(21)+a(122)).^2)/(2*a(1)^2))+a(92)*exp(-((x-
a(23)+a(122)).^2)/(2*a(1)^2))+a(93)*exp(-((x-a(25)+a(122)).^2)/(2*a(1)^2))+a(94)*exp(-
((x-a(27)+a(122)).^2)/(2*a(1)^2))+a(95)*exp(-((x-
a(29)+a(122)).^2)/(2*a(1)^2))+a(96)*exp(-((x-a(31)+a(122)).^2)/(2*a(1)^2))+a(97)*exp(-
((x-a(33)+a(122)).^2)/(2*a(1)^2))+a(98)*exp(-((x-
a(35)+a(122)).^2)/(2*a(1)^2))+a(99)*exp(-((x-
a(37)+a(122)).^2)/(2*a(1)^2))+a(100)*exp(-((x-
a(39)+a(122)).^2)/(2*a(1)^2))+a(101)*exp(-((x-
a(41)+a(122)).^2)/(2*a(1)^2))+a(102)*exp(-((x-
a(43)+a(122)).^2)/(2*a(1)^2))+a(103)*exp(-((x-
a(45)+a(122)).^2)/(2*a(1)^2))+a(104)*exp(-((x-
a(47)+a(122)).^2)/(2*a(1)^2))+a(105)*exp(-((x-
a(49)+a(122)).^2)/(2*a(1)^2))+a(106)*exp(-((x-
a(51)+a(122)).^2)/(2*a(1)^2))+a(107)*exp(-((x-
a(53)+a(122)).^2)/(2*a(1)^2))+a(108)*exp(-((x-
a(55)+a(122)).^2)/(2*a(1)^2))+a(109)*exp(-((x-
a(57)+a(122)).^2)/(2*a(1)^2))+a(110)*exp(-((x-
a(59)+a(122)).^2)/(2*a(1)^2))+a(111)*exp(-((x-
a(61)+a(122)).^2)/(2*a(1)^2))+a(112)*exp(-((x-
a(63)+a(122)).^2)/(2*a(1)^2))+a(113)*exp(-((x-
a(65)+a(122)).^2)/(2*a(1)^2))+a(114)*exp(-((x-
a(67)+a(122)).^2)/(2*a(1)^2))+a(115)*exp(-((x-
a(69)+a(122)).^2)/(2*a(1)^2))+a(116)*exp(-((x-
a(71)+a(122)).^2)/(2*a(1)^2))+a(117)*exp(-((x-
a(73)+a(122)).^2)/(2*a(1)^2))+a(118)*exp(-((x-
a(75)+a(122)).^2)/(2*a(1)^2))+a(119)*exp(-((x-
a(77)+a(122)).^2)/(2*a(1)^2))+a(120)*exp(-((x-
a(79)+a(122)).^2)/(2*a(1)^2))+a(121)*exp(-((x-
a(81)+a(122)).^2)/(2*a(1)^2))+a(123)*exp(-((x-a(124)).^2)/(2*a(125)^2)); %a(1)=width
of all peaks, a(122)=shoulder offset, a(123-126) are for underlying polynomial (3rd-
order) baseline
```

```
a0=[0.3 ampstart E(1,1) ampstart EA(1,1) ampstart E(2,1) ampstart EA(2,1) ampstart
E(3,1) ampstart EA(3,1) ampstart E(4,1) ampstart EA(4,1) ampstart E(5,1) ampstart
EA(5,1) ampstart E(6,1) ampstart EA(6,1) ampstart E(7,1) ampstart EA(7,1) ampstart
E(8,1) ampstart EA(8,1) ampstart E(9,1) ampstart EA(9,1) ampstart E(10,1) ampstart
EA(10,1) ampstart E(11,1) ampstart EA(11,1) ampstart E(12,1) ampstart EA(12,1)
ampstart E(13,1) ampstart EA(13,1) ampstart E(14,1) ampstart EA(14,1) ampstart E(15,1)
ampstart EA(15,1)  ampstart E(16,1) ampstart EA(16,1)  ampstart E(17,1) ampstart
EA(17,1) ampstart E(18,1) ampstart EA(18,1) ampstart E(19,1) ampstart EA(19,1)
ampstart E(20,1) ampstart EA(20,1) 0 0 0 0 0 0 0 0 0 0 0 0 0 0 0 0 0 0 0 0 0 0 0 0 0 0
0 0 0 0 0 0 0 0 0 0 0 0 0 0 -1 2E5 950 200 ];
lb=[0.2 amplb E(1,1)-offset amplb EA(1,1)-offset amplb E(2,1)-offset amplb EA(2,1)-
offset amplb E(3,1)-offset amplb EA(3,1)-offset amplb E(4,1)-offset amplb EA(4,1)-
offset amplb E(5,1)-offset amplb EA(5,1)-offset amplb E(6,1)-offset amplb EA(6,1)-
offset amplb E(7,1)-offset amplb EA(7,1)-offset amplb E(8,1)-offset amplb EA(8,1)-
offset amplb E(9,1)-offset amplb EA(9,1)-offset amplb E(10,1)-offset amplb EA(10,1)-
offset amplb E(11,1)-offset amplb EA(11,1)-offset amplb E(12,1)-offset amplb EA(12,1)-
offset amplb E(13,1)-offset amplb EA(13,1)-offset amplb E(14,1)-offset amplb EA(14,1)-
offset amplb E(15,1)-offset amplb EA(15,1)-offset  amplb E(16,1)-offset amplb
EA(16,1)-offset  amplb E(17,1)-offset amplb EA(17,1)-offset amplb E(18,1)-offset amplb
EA(18,1)-offset amplb E(19,1)-offset amplb EA(19,1)-offset amplb E(20,1)-offset amplb
EA(20,1)-offset 0 0 0 0 0 0 0 0 0 0 0 0 0 0 0 0 0 0 0 0 0 0 0 0 0 0 0 0 0 0 0 0 0 0 0
0 0 0 0 0 -2.0 -Inf -Inf -Inf ];
ub=[0.3 ampub E(1,1)+offset ampub EA(1,1)+offset ampub E(2,1)+offset ampub
EA(2,1)+offset ampub E(3,1)+offset ampub EA(3,1)+offset ampub E(4,1)+offset ampub
EA(4,1)+offset ampub E(5,1)+offset ampub EA(5,1)+offset ampub E(6,1)+offset ampub
EA(6,1)+offset ampub E(7,1)+offset ampub EA(7,1)+offset ampub E(8,1)+offset ampub
EA(8,1)+offset ampub E(9,1)+offset ampub EA(9,1)+offset ampub E(10,1)+offset ampub
EA(10,1)+offset ampub E(11,1)+offset ampub EA(11,1)+offset ampub E(12,1)+offset ampub
EA(12,1)+offset ampub E(13,1)+offset ampub EA(13,1)+offset ampub E(14,1)+offset ampub
EA(14,1)+offset ampub E(15,1)+offset ampub EA(15,1)+offset  ampub E(16,1)+offset ampub
EA(16,1)+offset  ampub E(17,1)+offset ampub EA(17,1)+offset ampub E(18,1)+offset ampub
EA(18,1)+offset ampub E(19,1)+offset ampub EA(19,1)+offset ampub E(20,1)+offset ampub
EA(20,1)+offset 0 0 0 0 0 0 0 0 0 0 0 0 0 0 0 0 0 0 0 0 0 0 0 0 0 0 0 0 0 0 0 0 0 0 0
0 0 0 0 0 -0.5 Inf Inf Inf ];

options=optimset('MaxFunEvals',5000000,'TolFun',1e-10,'MaxIter',20000,'TolX', 1E-8);
%options parameters for fitting
[a,resnorm,residual,exitflag,output,lambda,jacobian] = lsqcurvefit(fun, a0, Da,
Counts,lb,ub,options);

% Fit 2 - Fit shoulder (fix main peak)
fun=@(a,x)a(2)*exp(-((x-a(3)).^2)/(2*a(1)^2))+a(4)*exp(-((x-
a(5)).^2)/(2*a(1)^2))+a(6)*exp(-((x-a(7)).^2)/(2*a(1)^2))+a(8)*exp(-((x-
a(9)).^2)/(2*a(1)^2))+a(10)*exp(-((x-a(11)).^2)/(2*a(1)^2))+a(12)*exp(-((x-
a(13)).^2)/(2*a(1)^2))+a(14)*exp(-((x-a(15)).^2)/(2*a(1)^2))+a(16)*exp(-((x-
a(17)).^2)/(2*a(1)^2))+a(18)*exp(-((x-a(19)).^2)/(2*a(1)^2))+a(20)*exp(-((x-
a(21)).^2)/(2*a(1)^2))+a(22)*exp(-((x-a(23)).^2)/(2*a(1)^2))+a(24)*exp(-((x-
a(25)).^2)/(2*a(1)^2))+a(26)*exp(-((x-a(27)).^2)/(2*a(1)^2))+a(28)*exp(-((x-
a(29)).^2)/(2*a(1)^2))+a(30)*exp(-((x-a(31)).^2)/(2*a(1)^2))+a(32)*exp(-((x-
a(33)).^2)/(2*a(1)^2))+a(34)*exp(-((x-a(35)).^2)/(2*a(1)^2))+a(36)*exp(-((x-
a(37)).^2)/(2*a(1)^2))+a(38)*exp(-((x-a(39)).^2)/(2*a(1)^2))+a(40)*exp(-((x-
a(41)).^2)/(2*a(1)^2))+a(42)*exp(-((x-a(43)).^2)/(2*a(1)^2))+a(44)*exp(-((x-
a(45)).^2)/(2*a(1)^2))+a(46)*exp(-((x-a(47)).^2)/(2*a(1)^2))+a(48)*exp(-((x-
a(49)).^2)/(2*a(1)^2))+a(50)*exp(-((x-a(51)).^2)/(2*a(1)^2))+a(52)*exp(-((x-
a(53)).^2)/(2*a(1)^2))+a(54)*exp(-((x-a(55)).^2)/(2*a(1)^2))+a(56)*exp(-((x-
a(57)).^2)/(2*a(1)^2))+a(58)*exp(-((x-a(59)).^2)/(2*a(1)^2))+a(60)*exp(-((x-
a(61)).^2)/(2*a(1)^2))+a(62)*exp(-((x-a(63)).^2)/(2*a(1)^2))+a(64)*exp(-((x-
a(65)).^2)/(2*a(1)^2))+a(66)*exp(-((x-a(67)).^2)/(2*a(1)^2))+a(68)*exp(-((x-
a(69)).^2)/(2*a(1)^2))+a(70)*exp(-((x-a(71)).^2)/(2*a(1)^2))+a(72)*exp(-((x-
a(73)).^2)/(2*a(1)^2))+a(74)*exp(-((x-a(75)).^2)/(2*a(1)^2))+a(76)*exp(-((x-
a(77)).^2)/(2*a(1)^2))+a(78)*exp(-((x-a(79)).^2)/(2*a(1)^2))+a(80)*exp(-((x-
a(81)).^2)/(2*a(1)^2))+a(82)*exp(-((x-a(3)+a(122)).^2)/(2*a(1)^2))+a(83)*exp(-((x-
a(5)+a(122)).^2)/(2*a(1)^2))+a(84)*exp(-((x-a(7)+a(122)).^2)/(2*a(1)^2))+a(85)*exp(-
```

```
((x-a(9)+a(122)).^2)/(2*a(1)^2))+a(86)*exp(-((x-
a(11)+a(122)).^2)/(2*a(1)^2))+a(87)*exp(-((x-a(13)+a(122)).^2)/(2*a(1)^2))+a(88)*exp(-
((x-a(15)+a(122)).^2)/(2*a(1)^2))+a(89)*exp(-((x-
a(17)+a(122)).^2)/(2*a(1)^2))+a(90)*exp(-((x-a(19)+a(122)).^2)/(2*a(1)^2))+a(91)*exp(-
((x-a(21)+a(122)).^2)/(2*a(1)^2))+a(92)*exp(-((x-
a(23)+a(122)).^2)/(2*a(1)^2))+a(93)*exp(-((x-a(25)+a(122)).^2)/(2*a(1)^2))+a(94)*exp(-
((x-a(27)+a(122)).^2)/(2*a(1)^2))+a(95)*exp(-((x-
a(29)+a(122)).^2)/(2*a(1)^2))+a(96)*exp(-((x-a(31)+a(122)).^2)/(2*a(1)^2))+a(97)*exp(-
((x-a(33)+a(122)).^2)/(2*a(1)^2))+a(98)*exp(-((x-
a(35)+a(122)).^2)/(2*a(1)^2))+a(99)*exp(-((x-
a(37)+a(122)).^2)/(2*a(1)^2))+a(100)*exp(-((x-
a(39)+a(122)).^2)/(2*a(1)^2))+a(101)*exp(-((x-
a(41)+a(122)).^2)/(2*a(1)^2))+a(102)*exp(-((x-
a(43)+a(122)).^2)/(2*a(1)^2))+a(103)*exp(-((x-
a(45)+a(122)).^2)/(2*a(1)^2))+a(104)*exp(-((x-
a(47)+a(122)).^2)/(2*a(1)^2))+a(105)*exp(-((x-
a(49)+a(122)).^2)/(2*a(1)^2))+a(106)*exp(-((x-
a(51)+a(122)).^2)/(2*a(1)^2))+a(107)*exp(-((x-
a(53)+a(122)).^2)/(2*a(1)^2))+a(108)*exp(-((x-
a(55)+a(122)).^2)/(2*a(1)^2))+a(109)*exp(-((x-
a(57)+a(122)).^2)/(2*a(1)^2))+a(110)*exp(-((x-
a(59)+a(122)).^2)/(2*a(1)^2))+a(111)*exp(-((x-
a(61)+a(122)).^2)/(2*a(1)^2))+a(112)*exp(-((x-
a(63)+a(122)).^2)/(2*a(1)^2))+a(113)*exp(-((x-
a(65)+a(122)).^2)/(2*a(1)^2))+a(114)*exp(-((x-
a(67)+a(122)).^2)/(2*a(1)^2))+a(115)*exp(-((x-
a(69)+a(122)).^2)/(2*a(1)^2))+a(116)*exp(-((x-
a(71)+a(122)).^2)/(2*a(1)^2))+a(117)*exp(-((x-
a(73)+a(122)).^2)/(2*a(1)^2))+a(118)*exp(-((x-
a(75)+a(122)).^2)/(2*a(1)^2))+a(119)*exp(-((x-
a(77)+a(122)).^2)/(2*a(1)^2))+a(120)*exp(-((x-
a(79)+a(122)).^2)/(2*a(1)^2))+a(121)*exp(-((x-
a(81)+a(122)).^2)/(2*a(1)^2))+a(123)*exp(-((x-a(124)).^2)/(2*a(125)^2)); %a(1)=width
of all peaks, a(122)=shoulder offset, a(123-126) are for underlying polynomial (3rd-
order) baseline
a0=a; %Uses the previously fit parameters as starting conditions for Fit 2, lb/ub for
1st peak fixed
lb=[0.3 a(2) a(3) a(4) a(5) a(6) a(7) a(8) a(9) a(10) a(11) a(12) a(13) a(14) a(15)
a(16) a(17) a(18) a(19) a(20) a(21) a(22) a(23) a(24) a(25) a(26) a(27) a(28) a(29)
a(30) a(31) a(32) a(33) a(34) a(35) a(36) a(37) a(38) a(39) a(40) a(41) a(42) a(43)
a(44) a(45) a(46) a(47) a(48) a(49) a(50) a(51) a(52) a(53) a(54) a(55) a(56) a(57)
a(58) a(59) a(60) a(61) a(62) a(63) a(64) a(65) a(66) a(67) a(68) a(69) a(70) a(71)
a(72) a(73) a(74) a(75) a(76) a(77) a(78) a(79) a(80) a(81) amplb amplb amplb amplb
amplb amplb amplb amplb amplb amplb amplb amplb amplb amplb amplb amplb amplb amplb
amplb amplb amplb amplb amplb amplb amplb amplb amplb amplb amplb amplb amplb amplb
amplb amplb amplb amplb amplb amplb amplb amplb -2.0 -Inf -Inf -Inf ];
ub=[0.3 a(2) a(3) a(4) a(5) a(6) a(7) a(8) a(9) a(10) a(11) a(12) a(13) a(14) a(15)
a(16) a(17) a(18) a(19) a(20) a(21) a(22) a(23) a(24) a(25) a(26) a(27) a(28) a(29)
a(30) a(31) a(32) a(33) a(34) a(35) a(36) a(37) a(38) a(39) a(40) a(41) a(42) a(43)
a(44) a(45) a(46) a(47) a(48) a(49) a(50) a(51) a(52) a(53) a(54) a(55) a(56) a(57)
a(58) a(59) a(60) a(61) a(62) a(63) a(64) a(65) a(66) a(67) a(68) a(69) a(70) a(71)
a(72) a(73) a(74) a(75) a(76) a(77) a(78) a(79) a(80) a(81) ampub ampub ampub ampub
ampub ampub ampub ampub ampub ampub ampub ampub ampub ampub ampub ampub ampub ampub
ampub ampub ampub ampub ampub ampub ampub ampub ampub ampub ampub ampub ampub ampub
ampub ampub ampub ampub ampub ampub ampub ampub -0.5 Inf Inf Inf ];

options=optimset('MaxFunEvals',5000000,'TolFun',1e-4,'MaxIter',20000,'TolX', 1E-4);
%options parameters for fitting
[a,resnorm,residual,exitflag,output,lambda,jacobian] = lsqcurvefit(fun, a0, Da,
Counts,lb,ub,options);

% Fit 3 - Final Fit, Relaxed Parameters
```

```
fun=@(a,x)a(2)*exp(-((x-a(3)).^2)/(2*a(1)^2))+a(4)*exp(-((x-
a(5)).^2)/(2*a(1)^2))+a(6)*exp(-((x-a(7)).^2)/(2*a(1)^2))+a(8)*exp(-((x-
a(9)).^2)/(2*a(1)^2))+a(10)*exp(-((x-a(11)).^2)/(2*a(1)^2))+a(12)*exp(-((x-
a(13)).^2)/(2*a(1)^2))+a(14)*exp(-((x-a(15)).^2)/(2*a(1)^2))+a(16)*exp(-((x-
a(17)).^2)/(2*a(1)^2))+a(18)*exp(-((x-a(19)).^2)/(2*a(1)^2))+a(20)*exp(-((x-
a(21)).^2)/(2*a(1)^2))+a(22)*exp(-((x-a(23)).^2)/(2*a(1)^2))+a(24)*exp(-((x-
a(25)).^2)/(2*a(1)^2))+a(26)*exp(-((x-a(27)).^2)/(2*a(1)^2))+a(28)*exp(-((x-
a(29)).^2)/(2*a(1)^2))+a(30)*exp(-((x-a(31)).^2)/(2*a(1)^2))+a(32)*exp(-((x-
a(33)).^2)/(2*a(1)^2))+a(34)*exp(-((x-a(35)).^2)/(2*a(1)^2))+a(36)*exp(-((x-
a(37)).^2)/(2*a(1)^2))+a(38)*exp(-((x-a(39)).^2)/(2*a(1)^2))+a(40)*exp(-((x-
a(41)).^2)/(2*a(1)^2))+a(42)*exp(-((x-a(43)).^2)/(2*a(1)^2))+a(44)*exp(-((x-
a(45)).^2)/(2*a(1)^2))+a(46)*exp(-((x-a(47)).^2)/(2*a(1)^2))+a(48)*exp(-((x-
a(49)).^2)/(2*a(1)^2))+a(50)*exp(-((x-a(51)).^2)/(2*a(1)^2))+a(52)*exp(-((x-
a(53)).^2)/(2*a(1)^2))+a(54)*exp(-((x-a(55)).^2)/(2*a(1)^2))+a(56)*exp(-((x-
a(57)).^2)/(2*a(1)^2))+a(58)*exp(-((x-a(59)).^2)/(2*a(1)^2))+a(60)*exp(-((x-
a(61)).^2)/(2*a(1)^2))+a(62)*exp(-((x-a(63)).^2)/(2*a(1)^2))+a(64)*exp(-((x-
a(65)).^2)/(2*a(1)^2))+a(66)*exp(-((x-a(67)).^2)/(2*a(1)^2))+a(68)*exp(-((x-
a(69)).^2)/(2*a(1)^2))+a(70)*exp(-((x-a(71)).^2)/(2*a(1)^2))+a(72)*exp(-((x-
a(73)).^2)/(2*a(1)^2))+a(74)*exp(-((x-a(75)).^2)/(2*a(1)^2))+a(76)*exp(-((x-
a(77)).^2)/(2*a(1)^2))+a(78)*exp(-((x-a(79)).^2)/(2*a(1)^2))+a(80)*exp(-((x-
a(81)).^2)/(2*a(1)^2))+a(82)*exp(-((x-a(3)+a(122)).^2)/(2*a(1)^2))+a(83)*exp(-((x-
a(5)+a(122)).^2)/(2*a(1)^2))+a(84)*exp(-((x-a(7)+a(122)).^2)/(2*a(1)^2))+a(85)*exp(-
((x-a(9)+a(122)).^2)/(2*a(1)^2))+a(86)*exp(-((x-
a(11)+a(122)).^2)/(2*a(1)^2))+a(87)*exp(-((x-a(13)+a(122)).^2)/(2*a(1)^2))+a(88)*exp(-
((x-a(15)+a(122)).^2)/(2*a(1)^2))+a(89)*exp(-((x-
a(17)+a(122)).^2)/(2*a(1)^2))+a(90)*exp(-((x-a(19)+a(122)).^2)/(2*a(1)^2))+a(91)*exp(-
((x-a(21)+a(122)).^2)/(2*a(1)^2))+a(92)*exp(-((x-
a(23)+a(122)).^2)/(2*a(1)^2))+a(93)*exp(-((x-a(25)+a(122)).^2)/(2*a(1)^2))+a(94)*exp(-
((x-a(27)+a(122)).^2)/(2*a(1)^2))+a(95)*exp(-((x-
a(29)+a(122)).^2)/(2*a(1)^2))+a(96)*exp(-((x-a(31)+a(122)).^2)/(2*a(1)^2))+a(97)*exp(-
((x-a(33)+a(122)).^2)/(2*a(1)^2))+a(98)*exp(-((x-
a(35)+a(122)).^2)/(2*a(1)^2))+a(99)*exp(-((x-
a(37)+a(122)).^2)/(2*a(1)^2))+a(100)*exp(-((x-
a(39)+a(122)).^2)/(2*a(1)^2))+a(101)*exp(-((x-
a(41)+a(122)).^2)/(2*a(1)^2))+a(102)*exp(-((x-
a(43)+a(122)).^2)/(2*a(1)^2))+a(103)*exp(-((x-
a(45)+a(122)).^2)/(2*a(1)^2))+a(104)*exp(-((x-
a(47)+a(122)).^2)/(2*a(1)^2))+a(105)*exp(-((x-
a(49)+a(122)).^2)/(2*a(1)^2))+a(106)*exp(-((x-
a(51)+a(122)).^2)/(2*a(1)^2))+a(107)*exp(-((x-
a(53)+a(122)).^2)/(2*a(1)^2))+a(108)*exp(-((x-
a(55)+a(122)).^2)/(2*a(1)^2))+a(109)*exp(-((x-
a(57)+a(122)).^2)/(2*a(1)^2))+a(110)*exp(-((x-
a(59)+a(122)).^2)/(2*a(1)^2))+a(111)*exp(-((x-
a(61)+a(122)).^2)/(2*a(1)^2))+a(112)*exp(-((x-
a(63)+a(122)).^2)/(2*a(1)^2))+a(113)*exp(-((x-
a(65)+a(122)).^2)/(2*a(1)^2))+a(114)*exp(-((x-
a(67)+a(122)).^2)/(2*a(1)^2))+a(115)*exp(-((x-
a(69)+a(122)).^2)/(2*a(1)^2))+a(116)*exp(-((x-
a(71)+a(122)).^2)/(2*a(1)^2))+a(117)*exp(-((x-
a(73)+a(122)).^2)/(2*a(1)^2))+a(118)*exp(-((x-
a(75)+a(122)).^2)/(2*a(1)^2))+a(119)*exp(-((x-
a(77)+a(122)).^2)/(2*a(1)^2))+a(120)*exp(-((x-
a(79)+a(122)).^2)/(2*a(1)^2))+a(121)*exp(-((x-
a(81)+a(122)).^2)/(2*a(1)^2))+a(123)*exp(-((x-a(124)).^2)/(2*a(125)^2)); %a(1)=width
of all peaks, a(122)=shoulder offset, a(123-126) are for underlying polynomial (3rd-
order) baseline
a0=a;
lb=[0.2 amplb E(1,1)-offset amplb EA(1,1)-offset amplb E(2,1)-offset amplb EA(2,1)-
offset amplb E(3,1)-offset amplb EA(3,1)-offset amplb E(4,1)-offset amplb EA(4,1)-
offset amplb E(5,1)-offset amplb EA(5,1)-offset amplb E(6,1)-offset amplb EA(6,1)-
offset amplb E(7,1)-offset amplb EA(7,1)-offset amplb E(8,1)-offset amplb EA(8,1)-
offset amplb E(9,1)-offset amplb EA(9,1)-offset amplb E(10,1)-offset amplb EA(10,1)-
```

```
offset amplb E(11,1)-offset amplb EA(11,1)-offset amplb E(12,1)-offset amplb EA(12,1)-
offset amplb E(13,1)-offset amplb EA(13,1)-offset amplb E(14,1)-offset amplb EA(14,1)-
offset amplb E(15,1)-offset amplb EA(15,1)-offset  amplb E(16,1)-offset amplb
EA(16,1)-offset  amplb E(17,1)-offset amplb EA(17,1)-offset amplb E(18,1)-offset amplb
EA(18,1)-offset amplb E(19,1)-offset amplb EA(19,1)-offset amplb E(20,1)-offset amplb
EA(20,1)-offset amplb amplb amplb amplb amplb amplb amplb amplb amplb amplb amplb
amplb amplb amplb amplb amplb amplb amplb amplb amplb amplb amplb amplb amplb amplb
amplb amplb amplb amplb amplb amplb amplb amplb amplb amplb amplb amplb amplb amplb
amplb -2 -Inf -Inf -Inf ];
ub=[0.5 ampub E(1,1)+offset ampub EA(1,1)+offset ampub E(2,1)+offset ampub
EA(2,1)+offset ampub E(3,1)+offset ampub EA(3,1)+offset ampub E(4,1)+offset ampub
EA(4,1)+offset ampub E(5,1)+offset ampub EA(5,1)+offset ampub E(6,1)+offset ampub
EA(6,1)+offset ampub E(7,1)+offset ampub EA(7,1)+offset ampub E(8,1)+offset ampub
EA(8,1)+offset ampub E(9,1)+offset ampub EA(9,1)+offset ampub E(10,1)+offset ampub
EA(10,1)+offset ampub E(11,1)+offset ampub EA(11,1)+offset ampub E(12,1)+offset ampub
EA(12,1)+offset ampub E(13,1)+offset ampub EA(13,1)+offset ampub E(14,1)+offset ampub
EA(14,1)+offset ampub E(15,1)+offset ampub EA(15,1)+offset  ampub E(16,1)+offset ampub
EA(16,1)+offset  ampub E(17,1)+offset ampub EA(17,1)+offset ampub E(18,1)+offset ampub
EA(18,1)+offset ampub E(19,1)+offset ampub EA(19,1)+offset ampub E(20,1)+offset ampub
EA(20,1)+offset ampub ampub ampub ampub ampub ampub ampub ampub ampub ampub ampub
ampub ampub ampub ampub ampub ampub ampub ampub ampub ampub ampub ampub ampub ampub
ampub ampub ampub ampub ampub ampub ampub ampub ampub ampub ampub ampub ampub ampub
ampub -0.5 Inf Inf Inf ];

options=optimset('MaxFunEvals',5000000,'TolFun',1e-8,'MaxIter',20000,'TolX', 1E-8);
%options parameters for fitting
[a,resnorm,residual,exitflag,output,lambda,jacobian] = lsqcurvefit(fun, a0, Da,
Counts,lb,ub,options);
%% Store and Report Fit Data
%Store the individual mass fraction peaks and areas for the various ES and
%E peaks
Intensities(:,2)=a(3:4:79);%E_mass
Intensities(:,3)=a(2:4:78);%E_mass areas main peak
Intensities(:,4)=a(82:2:120);%E_mass areas shoulder
Intensities(:,5)=a(5:4:81);%EA_mass
Intensities(:,6)=a(4:4:80);%EA_mass areas main peak
Intensities(:,7)=a(83:2:121);%EA_mass areas shoulder

Ratio=(Intensities(:,6)+Intensities(:,7))./(Intensities(:,3)+Intensities(:,4)); %Ratio
EA/E
Intensities(:,8)=Ratio;
MeanRatio=mean(Ratio(2:19));%Average EA/ES ratio
StdRatio=std(Ratio(2:19)); %Std.dev of ratio
Resid=Counts-fun(a,Da);

%Direct to the variables where data is stored and display the avg and std.
%dev of EA/ES ratio
disp('Variables stored in "Intensities", "Ratio (EA/ES)", "MeanRatio" and "StdRatio"')
disp('Average EA/ES ratio is ...')
disp(MeanRatio)
disp('Std. dev of EA/ES ratio is ...')
disp(StdRatio)

%Plot the data and fits
plot(Da,Counts,Da,fun(a,Da),Da,Resid)
```

---

---

*"RateDistribution.m"*

-------------------------------------------------------------------------------------------------------------------

```
%% RateDistribution: Use EA/ES ratios determined from steady-state MS and kcat (and
errors) to construct avg. and std dev. for k2 and k3. Written by Sam Schneider in June
2018 for utilization with TEM B-lactamases.
% load Ratio (EA/ES) data from .txt file (single column of ratios; no
% header)
[fname,pname]=uigetfile('*.*','Load MS Ratios','MultiSelect','on');
cd(pname);
Data=readtable(fname,'Delimiter',' ','Format','%f %f','ReadVariableNames',false);
x=Data{:,1}; %single column of Ratios from Steady-State MS analysis (MassAnalysis.m)

%input kcat (s-1), kcat std dev, and the # of bootstrap iterations to perform
dlgtitle='Input Information';
numLines=1;
prompt1={'kcat (sec^-1)','kcat Error','Iterations'};
def1={'4.2','1.3','1000000'};
answer1=inputdlg(prompt1,dlgtitle,numLines,def1);
drawnow

kcat=str2num(char(answer1(1))); %average value of kcat (s^-1)
kcat_error=str2num(char(answer1(2))); %standard deviations of kcat (s^-1)
Iter=str2num(char(answer1(3))); %number of iterations to sample

%% Determine k2 values
k2=zeros(length(x),Iter);
for i = 1:Iter;
kcatE=(kcat+(kcat_error*randn(1)));
k2_iter=kcatE.*(1+x);
k2(1:length(x),i)=k2_iter;
end
k2=k2(:);%convert matrix of k2 to single column vector

k2_avg=mean2(k2); %average over all elements of k2 matrix
k2_stddev=std2(k2); %std dev over all elements of k2 matrix
k2_median=median(k2);
yk2=quantile(k2,[0.01 0.025 0.05 0.15865 0.25 0.50 0.75 0.84135 0.95 0.975 0.99]);

disp('*********************************************************************************
****************************')
AVG_k2 = sprintf('  The MEAN k2 (s-1) is   %d   with std. dev of %d (68CI) [  %d  ,
%d  ], 95CI of [  %d  ,  %d  ]', k2_avg,k2_stddev,(k2_avg-
k2_stddev),(k2_avg+k2_stddev),(k2_avg-2*k2_stddev),(k2_avg+2*k2_stddev));
disp(AVG_k2);
MEDIAN_k2 = sprintf('The MEDIAN k2 (s-1) is   %d   with std. dev of (68CI) [  %d  ,
%d  ], 95CI of [  %d  ,  %d  ]', k2_median,yk2(4),yk2(8),yk2(2),yk2(10));
disp(MEDIAN_k2);
disp(' ------------------------------------  k2 Quantiles  ----------------------------
--------- ')
disp('The 1.0, 2.5, 5, 15.865, 25, 50, 75, 84.135, 95, 97.5, and 99% quantiles:')
disp(yk2)
disp(' --------------------------------------------------------------------------------
--------- ')
disp('*********************************************************************************
****************************')

%% Determine k3 values (independent of k2)
k3=zeros(length(x),Iter);
for i = 1:Iter;
k3(1:length(x),i)=(kcat+(kcat_error*randn(1))).*(1+(1./x));
```

```
end
k3=k3(:);%convert matrix of k3 to single column vector

k3_avg=mean2(k3); %average over all elements of k2 matrix
k3_stddev=std2(k3); %std dev over all elements of k2 matrix
k3_median=median(k3);
yk3=quantile(k3,[0.01 0.025 0.05 0.15865 0.25 0.50 0.75 0.84135 0.95 0.975 0.99]);

disp('*******************************************************************************
****************************')
disp(' FOR k3 INDEPENDENT OF k2 ')
AVG_k3 = sprintf('  The MEAN k3 (s-1) is   %d   with std. dev of %d [  %d  ,  %d  ],
95CI of [  %d  ,  %d  ]', k3_avg,k3_stddev,(k3_avg-
k3_stddev),(k3_avg+k3_stddev),(k3_avg-2*k3_stddev),(k3_avg+2*k3_stddev));
disp(AVG_k3);
MEDIAN_k3 = sprintf('The MEDIAN k3 (s-1) is   %d   with std. dev of [  %d  ,  %d  ],
95CI of [  %d  ,  %d  ]', k3_median,yk3(4),yk3(8),yk3(2),yk3(10));
disp(MEDIAN_k3);
disp(' ------------------------------------  k3 (k2 INDPENDENT) Quantiles  -----------
------------------------- ')
disp('The 1.0, 2.5, 5, 15.865, 25, 50, 75, 84.135, 95, 97.5, and 99% quantiles:')
disp(yk3)
disp(' --------------------------------------------------------------------------------
--------- ')
disp('*******************************************************************************
****************************')

%% Histogram data of Normalized Probability for k2 and k3
%Plot the Normalized probability distribution of k2. Lines indicate 25, 50
%and 75% percentile
subplot(1,3,1)
j=histogram(k2,'Normalization','probability');
EdgesJ=j.BinEdges;
WidthJ=j.BinWidth;
LimitsJ=j.BinLimits;
BinsJX=[(LimitsJ(1)+(WidthJ/2)):WidthJ:(LimitsJ(2)-(WidthJ/2))];
CountsJ=j.Values;
title('k2 Distribution (s-1)')
xlabel('k2 (sec-1)')
ylabel('Norm. Prob.')
%xlim([-2,12])

line([k2_median, k2_median], [0, max(CountsJ)*1.1],'Color','red','LineStyle','-')
line([yk2(4), yk2(4)], [0, max(CountsJ)*1.1],'Color','red','LineStyle','--')
line([yk2(8), yk2(8)], [0, max(CountsJ)]*1.1,'Color','red','LineStyle','--')
line([yk2(2), yk2(2)], [0, max(CountsJ)]*1.1,'Color','red','LineStyle','-.')
line([yk2(10), yk2(10)], [0, max(CountsJ)]*1.1,'Color','red','LineStyle','-.')
line([k2_avg, k2_avg], [0, max(CountsJ)]*1.1,'Color','green','LineStyle','-')
legend('k2','Median','-1StdDev','+1StdDev','-2StdDev','+2StdDev','Average')

%Plot the Normalized probability distribution of k3. Lines indicate 25, 50
%and 75% percentile
subplot(1,3,2)
h=histogram(k3,'Normalization','probability');
Edges=h.BinEdges;
Width=h.BinWidth;
Limits=h.BinLimits;
BinsX=[(Limits(1)+(Width/2)):Width:(Limits(2)-(Width/2))];
Counts=h.Values;
hold on
%k=histogram(k3d,'Normalization','probability');
%EdgesK=k.BinEdges;
%WidthK=k.BinWidth;
```

```
%LimitsK=k.BinLimits;
%BinsKX=[(LimitsK(1)+(WidthK/2)):WidthK:(LimitsK(2)-(WidthK/2))];
%CountsK=k.Values;
hold off
title('k3 Distribution (s-1)')
xlabel('k3 (sec-1)')
ylabel('Norm. Prob.')
%legend('k3-Ind','k3-Dep')
xlim([-100,(yk3(10)*1.2)])
%xlim([-100,1200])

line([k3_median, k3_median], [0, max(Counts)*1.1],'Color','red','LineStyle','-')
line([yk3(4), yk3(4)], [0, max(Counts)*1.1],'Color','red','LineStyle','--')
line([yk3(8), yk3(8)], [0, max(Counts)]*1.1,'Color','red','LineStyle','--')
line([yk3(2), yk3(2)], [0, max(Counts)]*1.1,'Color','red','LineStyle','-.')
line([yk3(10), yk3(10)], [0, max(Counts)]*1.1,'Color','red','LineStyle','-.')
line([k3_avg, k3_avg], [0, max(Counts)]*1.1,'Color','green','LineStyle','-')

legend('k3-Ind','Median','-1StdDev','+1StdDev','-2StdDev','+2StdDev','Average')

%Plot the BoxPlot of the k3 distribution
subplot(1,3,3)
boxplot(k3,'Notch','on','Symbol','','DataLim',[-100,yk3(10)*1.2])
ylim([-100,(yk3(10)*1.2)])
%boxplot([k3,k3d],'Notch','on','Symbol','','DataLim',[0,yk3(10)*1.2],'Labels',{'k3-
Ind','k3-Dep'})
ylabel('k3 (sec-1)')
title('k3 Distributions (s-1)')

disp('&&&&&&&&&&&&&&&&&&&&&&&&&&&&&&&&&&&&&&&&&&&&&&&&&&&&&&&&&&&&&&&&&&&&&&&&&&&&&&&&
&&&&&&&&&&&&&&&&&&&&&&&&&&&&&&')
disp('(x,y) plot of k2 probability distribution can be made from
"plot(BinsJX,CountsJ)"')
disp('(x,y) plot of k3 probability distribution can be made from
"plot(BinsX,Counts)"')
disp('For reference, in a normal distribution, 1 std. dev contains 68.27% of the data
(2 is 95.45%, 3 is 99.73%)')
disp('&&&&&&&&&&&&&&&&&&&&&&&&&&&&&&&&&&&&&&&&&&&&&&&&&&&&&&&&&&&&&&&&&&&&&&&&&&&&&&&&
&&&&&&&&&&&&&&&&&&&&&&&&&&&&&&')
```
